# Immunocompetent Cell Targeting by Food-Additive Titanium Dioxide

**DOI:** 10.1101/2024.04.16.589772

**Authors:** John W. Wills, Alicja Dabrowska, Jack Robertson, Michelle Miniter, Sebastian Riedle, Huw D. Summers, Rachel E. Hewitt, Adeeba Fathima, Alessandra Barreto da Silva, Carlos A. P. Bastos, Stuart Micklethwaite, Åsa V. Keita, Johan D. Söderholm, Nicole C. Roy, Don Otter, Ravin Jugdaohsingh, Pietro Mastroeni, Andy P. Brown, Paul Rees, Jonathan J. Powell

## Abstract

Food-grade titanium dioxide (fgTiO_2_) is a bio-persistent particle under intense regulatory scrutiny. Paradoxically, meaningful *in vivo* cellular accumulation has never been demonstrated: the only known cell reservoirs for fgTiO_2_ are ‘graveyard’ intestinal pigment cells which are metabolically and immunologically quiescent. Here we identify major new immunocompetent cell targets of fgTiO_2_ in humans, most notably in the subepithelial dome region of intestinal Peyer’s patches. Using multimodal microscopy techniques with single-particle detection and per-cell / vesicle image analysis we achieved correlative dosimetry, quantitatively recapitulating human cellular exposures in the terminal ileum of mice fed a fgTiO_2_-containing diet. Epithelial microfold cells selectively funneled fgTiO_2_ into LysoMac and LysoDC cells with ensuing accumulation. Notwithstanding, proximity extension analyses for 92 protein targets revealed no measureable perturbation of cell signalling pathways. When chased with oral Δ*aroA*-*Salmonella*, pro-inflammatory signalling was confirmed, but no augmentation by fgTiO_2_ was revealed despite marked same-cell loading. Interestingly, *Salmonella* caused the fgTiO_2_-recipient cells to migrate basolaterally in the patch and, sporadically, to the lamina propria, thereby fully recreating the intestinal tissue distribution of fgTiO_2_ in humans. Immunocompetent cells that accumulate fgTiO_2_ *in vivo* are now identified and we demonstrate a mouse model that finally enables human-relevant risk assessments of ingested, bio-persistent (nano)particles.

## INTRODUCTION

Titanium dioxide (TiO_2_) is a highly versatile nano- and micro-particulate mineral with widespread use in consumer products and manufacturing processes, including as a food and pharmaceutical additive. Although, for many years^1, 2^, there have been question marks over a potential role of ingested TiO_2_ in initiating or augmenting an inflammatory response in the intestine, the global industry, valued at >$17 billion per annum^3^, is now facing a whole new ‘bio-nano’ challenge. In May 2021, the European Food Safety Authority (EFSA) questioned the safety of food-grade TiO_2_ (fgTiO_2_; also termed E171) primarily citing unresolved genotoxicity concerns^4^. Based on this, European Commission member states agreed to remove fgTiO_2_ as an additive in foods. This represented a seemingly pragmatic decision balancing need for this additive (*e.g.,* aesthetics) against hypothetical risk. However, the European Commission also requested that the European Medicines Agency (EMA) evaluate the impact of removal of fgTiO_2_ from the list of authorised excipients that are permitted in medicinal products. In this case, fgTiO_2_ is *not* merely about aesthetics: it delivers opacity and protects actives from UV light degradation whilst aiding moisture control and patient compliance. The EMA concluded that any foreseeable ban, impacting the ∼91,000 medicinal products that contain this pigment in Europe, risks causing pharmaceutical shortages^5^. Moreover, they estimated that a period of 7-12 years would likely be required for implementation of suitable alternatives^5^. Meanwhile, consumer concerns over the safety of fgTiO_2_ have been expressed beyond Europe. Notably, in 2022, a class action suit was filed against the multinational food company Mars Incorporated in the USA alleging that consumers of the hard-shell candy ‘Skittles’ are at heightened risk of health effects stemming from genotoxicity^6^.

Whilst regulators in other continents / territories have not followed EFSA’s 2021 decision, there is widespread agreement that the available data addressing the uptake, biological fate and effects of fgTiO_2_ is insufficient^7, 8, 9^. In fact, direct toxicity of fgTiO_2_ in cell culture experiments has not been consistently demonstrated, especially when conditions are carefully controlled to avoid artefactual *in vitro* cell gorging^10^. In contrast, it is recognised *in vitro* that fgTiO_2_ can modestly augment cellular responses to primary danger signals (*e.g.,* microbe-associated molecular patterns) although cell loading of the particles needs to be relatively high^1^. Similarly, Kirkland *et al.,*^11^ identified any potential genotoxic activity of various titanium dioxide polymorphs as secondary to cell stress which, certainly for fgTiO_2,_ requires marked cell loading^1^.

Aside from specific cells of the small intestine, there are no known circumstances under which fgTiO_2_ markedly accumulates *in vivo*. Systemic exposure to fgTiO_2_ appears to be very low^12^. Even allowing for lifetime accumulation in humans it appears that anticipated target organs (liver and spleen) contain levels of Ti and Ti-rich particulates that are just above the tissue-free background, and the origin of this slightly enhanced signal is unknown^13^. Consistent with this, many animal feeding studies show almost no detectable systemic accumulation of fgTiO_2_^12, 14^ and, overall, there are no convincing *ex vivo* imaging studies showing multiple, heavily-loaded cells. The exception, as noted above, is the small intestine, in both humans and rodent models^15, 16^, and specifically their large lymphoid follicles termed Peyer’s patches. In absence of afferent lymphatics, the apical surfaces of these follicles possess a unique population of microfold (M) cells that avidly sample the lumen for bulky macromolecules including small micro- and large nano-particles (herein referred to as ‘particles’). This material is passed to underlying immune cells in the apical, ‘subepithelial dome’ (SED) region of the follicle which contains recognised populations of monocyte-derived, lysozyme-expressing cells termed ‘LysoMacs’ and ‘LysoDCs’ that are specialised for anti-microbial defence^17^. Overall, the major role of Peyer’s patches is initiating processes for the generation of mucosal IgA and systemic IgG in response to intestinal antigen thereby orchestrating the sequence of events that leads to gut-derived humoral immunity. Over time, fgTiO_2_ is known to accumulate in Peyer’s patches, alongside other persistent particles, in basally located tissue-fixed macrophages termed ‘pigment cells’^15, 16^. These cells are of low immunological and metabolic activity and appear to act as lifetime ‘graveyards’ for persistent particles, presumably limiting systemic absorption through local sequestration in the same way that tissue-fixed macrophages sequester particulate tattoo inks^15, 16, 18, 19^.

So the paradox remains: whilst fgTiO_2_ is under intense regulatory scrutiny, there is no known site of accumulation in active (*e.g.,* immuno-competent) cells that could plausibly give cause for concern or that allows informed risk assessment. Given that the intestine is the only known target of fgTiO_2_ accumulation we have addressed whether similar, selective cell loading occurs elsewhere (*i.e.,* away from the ‘safe house’ base-of-patch pigment cells) and, if so, what cells are targeted, at what frequency and with what impact? The approach used quantitative image analysis of light and electron microanalytical data, correlative mouse-human cell dosimetry and broad-target interrogation of fgTiO_2_^’^s capability to augment a primary cellular danger signal *in vivo*.

## RESULTS

### Titanium dioxide in human intestinal tissues

High-resolution reflectance confocal microscopy of human small intestinal tissue samples identified bright foci at the base of Peyer’s patches (**Figure 1a)** consistent with previous reports of pigment cells in this region^16, 18, 20^. These cells are known to mostly accumulate ingested engineered aluminosilicates and fgTiO_2_^16, 20^. Using Al and Ti as biomarkers for these particles, correlative analyses using scanning electron microscopy (SEM) and energy dispersive X-ray spectroscopy (EDX) confirmed the presence and co-localisation of these elements (**Figure 1b**; top row). Reflectant foci were found entirely attributable to Ti-rich particles, as areas with Al but no Ti signals were not reflectant, whereas all detectable Ti foci yielded reflectance (**Figure 1b**; rows 2-5). Alongside further checks using mouse tissues known to contain fgTiO_2_ or not (shown, **Supplementary Figure 1**), the reflectance imaging approach was confirmed as a robust, high-throughput means to detect fgTiO_2_ in tissue^15^.

**Figure 1.**
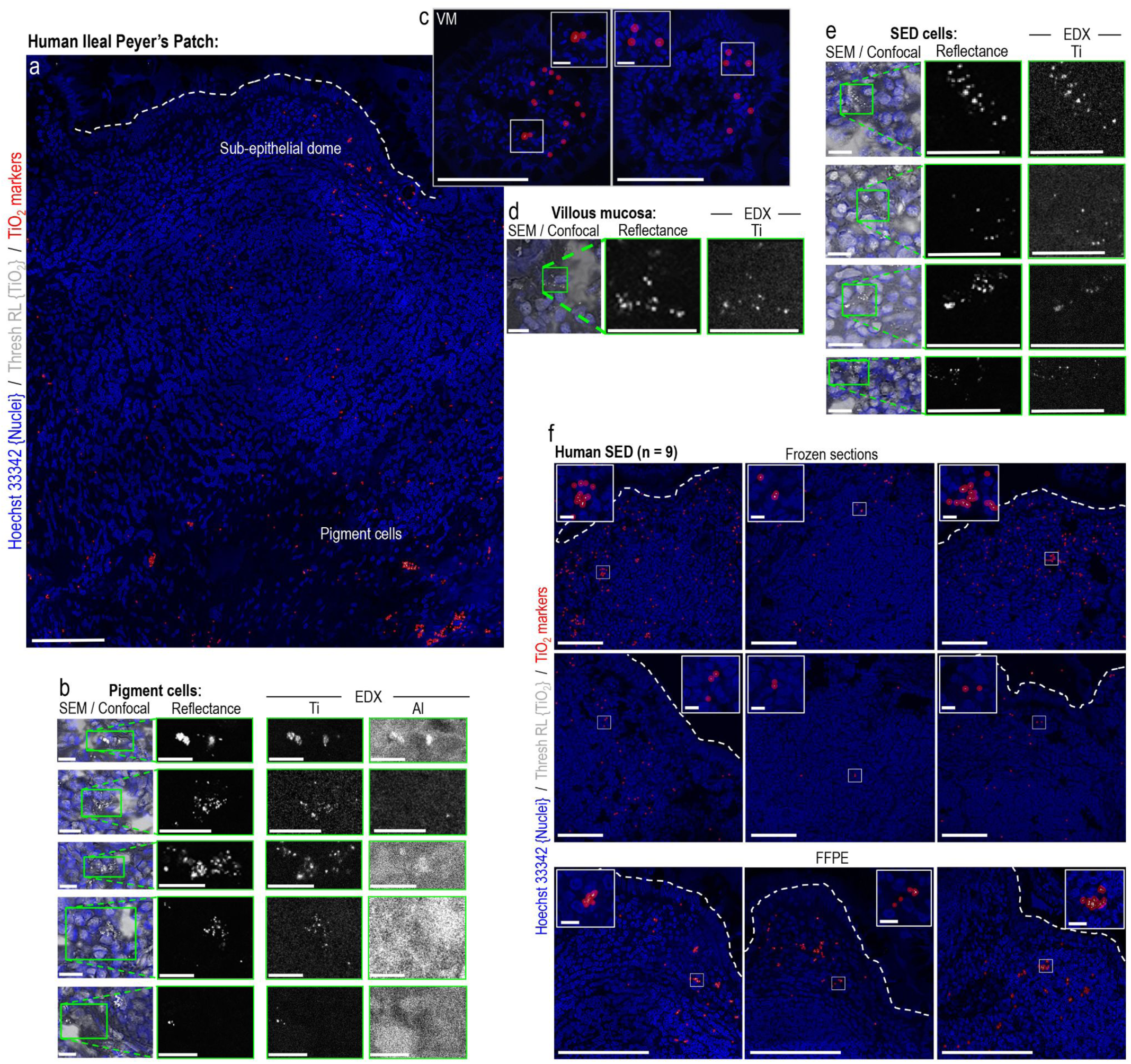
Titanium dioxide in human intestinal tissues. **a**, Tilescanned image acquired by confocal reflectance microscopy. Reflectant foci were detected along the follicle base consistent with previously-reported mineral particle-containing “pigment cells”. Of note, similar reflectant foci were also present apically in the immuno-active, subepithelial dome tissue-region, and away from the patch in the (**c**) lamina propria of the regular villous mucosa (VM). Translucent red circle-markers are placed on the reflectant foci to aid visualisation. **b**/**d**/**e**, Correlative SEM/EDX analyses performed on the same tissue section shown in (**a/c**). The pigment cell region is shown in (**b**) where X-ray signal for both Al (attributable to aluminosilicates) and Ti (attributable to titanium dioxide) was found. EDX analyses in the (**d**) villous mucosa and (**e**) subepithelial dome regions showed reflectant foci were attributable to Ti. **b/d/e**, In all instances the elemental data for Ti were a near-exact match for the reflectant foci, whereas (**b**) areas with Al signal alone did not reflect. **f**, Reflectance microscopical analyses of nine, randomly-drawn human samples (six frozen, three formalin-fixed paraffin embedded (FFPE)) showed reflectant foci consistent with TiO_2_ in the subepithelial dome of *every* sample. *Scale bars: **a** = 50 µm; **b** = 50 µm with 10 µm insets; **c-e** = 10 µm; **f** = 50 µm with 10 µm insets*.

Of significant note, previously unreported populations of human cells containing similar, reflective material were also identified (i) sporadically in the regular lamina propria (**Figure 1c**) and (ii) densely in the immuno-active subepithelial dome (SED) region of Peyer’s patches (**Figure 1a**). These cell regions were also subjected to correlative SEM-EDX analyses and, again, reflectant foci were a match for the Ti signal (**Figure 1d/e**). Given the unexpected density of the newly-identified Peyer’s patch population, whole SED regions from a randomly selected series of nine human subjects were imaged using the analytically-validated confocal microscopy approach. Albeit with marked inter-individual variability, reflectant signal was found in the SED of every human case (**Figure 1f**). Importantly, the recipient cells often exhibited multiple foci per-cell (*i.e.,* high cellular loading), suggesting specific cellular targeting and accumulation of fgTiO_2_ (**Figure 1f** – insets) in a non-pigment cell population.

### Recapitulation in a mouse model

To investigate the pathways giving rise to the above observations, and to phenotype the newly-identified population of fgTiO_2_-accumulating cells, in “Mouse Study 1”, we turned to a recently reported murine model for human-relevant, oral exposure to fgTiO_2_^15^. Initially we tilescanned a complete ileal transverse section using high-resolution confocal reflectance microscopy and used image analysis to segment all cells (cell segmentation accuracies assessed, **Supplementary Figure 2**). This allowed the cellular dose of fgTiO_2_ to be measured in each cell-object (*i.e.,* the integrated intensity of the thresholded reflectance signal per cell) enabling the subdivision and display of the cellular loading of fgTiO_2_ in three groups: low, intermediate and high (**Figure 2a**). As in the human samples (**Figure 1**), the patch region was decorated with reflectance signal, notably in the SED and at the base, and cellular loading was often high – especially in the SED (**Figure 2a**). However, unlike in human tissue, signal was entirely absent away from the patch except for two small, isolated foci in the villous epithelium. The resolution of the imaging process allowed detailed investigation of these latter events confirming that this signal was not *inside* the tissue, but derived from material within goblet cell invaginations that were open to the intestinal lumen (**Figure 2a** – inset).

**Figure 2.**
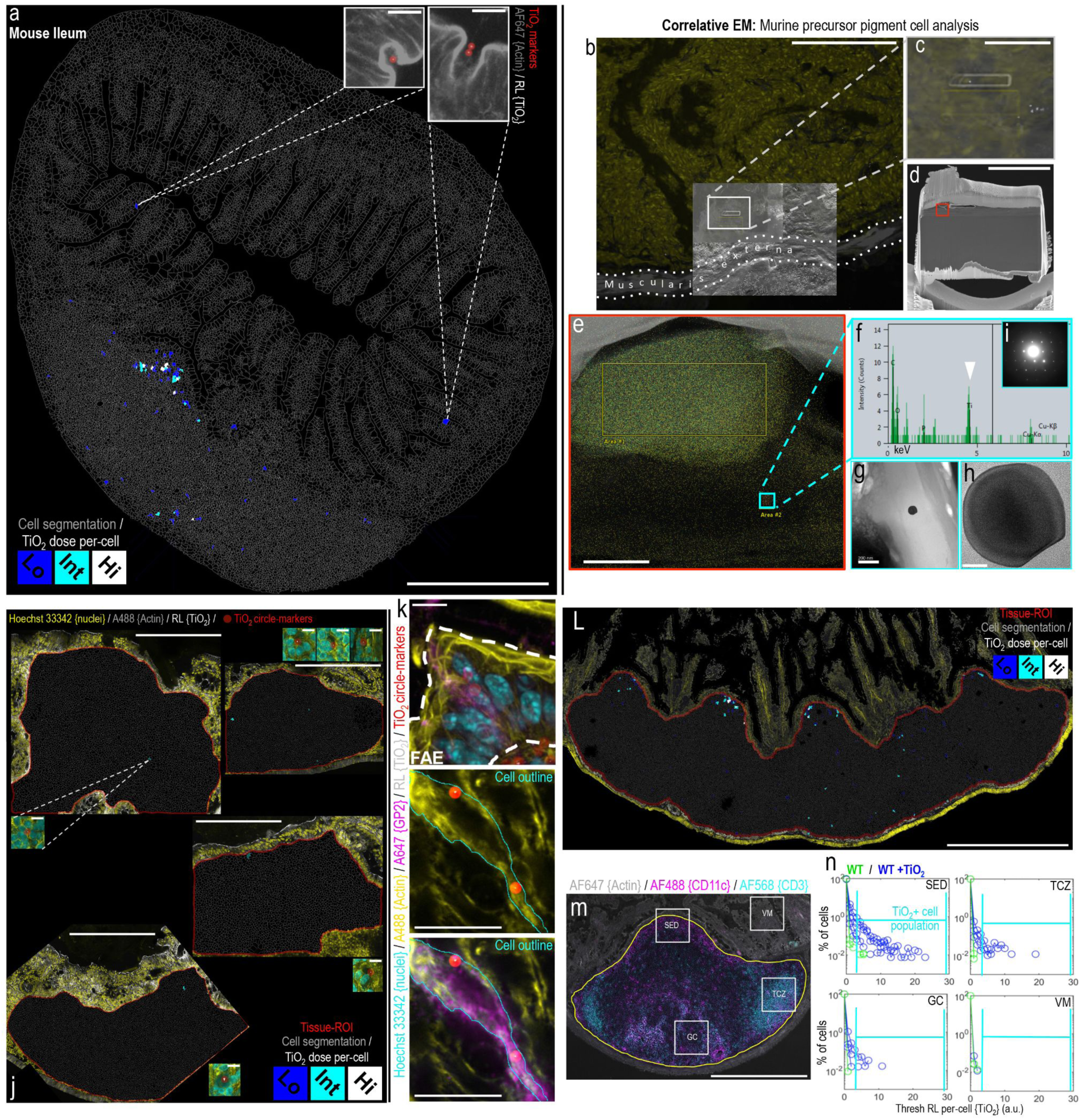
A mouse feeding study demonstrates fgTiO_2_ specificity for Peyer’s patches. **a,** Single-cell image analysis of a transverse mouse ileal tissue section imaged by tilescanning confocal reflectance microscopy. The cellular dose of fgTiO_2_ – measured from the thresholded reflectance (RL) signal – is colour-coded at low (lo), intermediate (int) and high (hi) levels. As in humans (Figure 1a**/f**), fgTiO_2_ loaded into the subepithelial dome of the Peyer’s patch and there was evidence of ‘precursor pigment cell’ formation along the follicle base. Away from the Peyer’s patch, two positive cells (**a**, insets) were found in the overlying villous mucosa. In contrast to the human findings (Figure 1c) however, both were caused by luminal fgTiO_2_ trapped in the invaginated space of goblet cells. **b-d**, To investigate the sensitivity of fgTiO_2_ detection by confocal reflectance microscopy, a tissue-region containing a single, isolated reflectant foci was milled out under correlative SEM using a focussed ion beam (**b/c** shows the registered SEM images transparently overlaid onto the confocal reflectance data before lift-out. The white ‘rectangle’ in (**c**) is caused by electron charging of the platinum strap laid down on the tissue region to be lifted out; (**d**) shows a side-view of the lifted-out lamella). **e-i**, Transfer of the lamella to a transmission electron microscope enabled high resolution imaging and X-ray (EDX) and electron diffraction analysis. A (**g/h**) single particle of fgTiO_2_ with an (**i**) anatase diffraction pattern (*i.e.,* as added to the diet) was confirmed responsible for the single reflectant foci observed. **j**, Again, colour-coding the cellular dose of fgTiO_2_ into lo, int and hi levels, tissue sections from adjacent caecal patches at the top of the colon were found to be almost completely devoid of fgTiO_2_ (n = 4 animals). **k**, Immunofluorescence microscopy showed fgTiO_2_ access to Peyer’s patches was via GP2-positive microfold (M) cells (translucent red circle-markers are placed on thresholded reflectant foci to aid fgTiO_2_ visualisation / the outline of the M-cell was manually drawn by tracing the actin-delineated cell boundary). **L**, Single-cell analysis of a longitudinal section of mouse ileum consistently showed the close association between an M-cell rich, follicle-associated epithelium and fgTiO_2_ loading into Peyer’s patches. **m/n**, Guided by CD11c and CD3 immunofluorescence labelling for phagocytic mononuclear cells and T-lymphocytes (respectively) sets of images were collected from the subepithelial dome (SED), germinal centre (GC) T-cell zone (TCZ) or overlying villous mucosa (VM) tissue regions from four mice fed the fgTiO_2_-supplemented diet (WT + TiO_2_) and four control-diet mice (WT). Some fgTiO_2_ signal was observed in the GC and TCZ regions of the murine Peyer’s patches but the majority was in the SED. In keeping with (**a**), and in contrast to the human findings (Figure 1c), in all four mice, no uptake was seen in the villous mucosa demonstrating specificity for M-cell mediated, Peyer’s patch targeting. *Scale bars: **a** = 500 µm with insets 5 µm; **b** = 50 µm; **c** = 10 µm; **d** = 5 µm; **e** = 500 nm; **g** = 200 nm; **h** = 20 nm; **j** = 500 µm; **k** = 10 µm; **L/m** = 500 µm*.

To confirm the apparent selectivity of fgTiO_2_ for the lymphoid follicles – rather than the regular villous mucosa – we questioned whether a single particle in intestinal tissue would be missed by our reflectance microscopy approach. To do this, a tissue region containing a single, small yet clearly visible reflectant foci was milled out under correlative SEM, using a focussed ion beam (**Figure 2b-d**). The milled-out lamella was then transferred to a transmission electron microscope for imaging with analysis by EDX and electron diffraction. A single particle of fgTiO_2_, with a diameter around 100 nm and an anatase diffraction pattern consistent with dietary exposure, was identified (**Figure 2e-i**). Since single, disperse particles of fgTiO_2_ were therefore detectable by the reflectance imaging strategy we could confirm that ingested fgTiO_2_ had not been missed with our imaging approach in the regular villous mucosa and thus was not taken up in this region (**Figure 2a**).

Following from this, we next considered the caecal patches (*i.e.,* large lymphoid follicles of the first part of the large intestine that adjoins the terminal ileum where Peyer’s patches are located). Here, transverse tissue sections revealed almost no reflectant foci showing that, unlike Peyer’s patches, the neighbouring caecal patch is spared from particle targeting (**Figure 2j**). In spite of the marked similarity of these secondary lymphoid sites, the microfold (M) cell density of their overlying epithelium differs greatly: caecal patches have very few (like the villous epithelium) with little capacity for particle uptake, whereas Peyer’s patch epithelium is enriched with mature M-cells that exhibit distinct particle uptake capability for small particulates in general^21, 22^. Indeed, cells of the Peyer’s patch epithelium stained positively for the mature M-cell marker (GP2) and, here-and-there, these M-cells were ‘caught in the act’ with co-signal for fgTiO_2_ (**Figure 2k**) as had long been assumed for fgTiO_2_ but not actually demonstrated^18^. Indeed, longitudinal sections of ileal tissue containing Peyer’s patches consistently demonstrated the close association between fgTiO_2_ signal and an overlying, M-cell rich, follicle-associated epithelium (**Figure 2L**). In this way, the above findings provide a rationale for the specificity of fgTiO_2_ uptake by the Peyer’s patches of the small intestine. It would appear, therefore, that to explain the presence of fgTiO_2_ in the human intestinal lamina propria a mechanism other than direct uptake by the regular villous epithelium must exist (this is addressed below in “Mouse Study 2”). First, however, we quantified how cell loading with fgTiO_2_ varied amongst the major different functional zones of the Peyer’s patch.

Sets of confocal reflectance images were collected from the SED, germinal centre (GC), T-cell zone (TCZ) and overlying villous mucosa (VM) tissue regions (indicated, **Figure 2m**) using tissue sections taken from four mice fed with the fgTiO_2_-supplemented diet (+fgTiO_2_), alongside four control mice from the same experiment and ingesting the same diet but without added fgTiO_2_. Single-cell analysis, using flow cytometry-type gating informed by the negative control tissues, demonstrated some fgTiO_2_ signal in the TCZ and GC but established the dominance of the signal, and the highest cell loading, in the biologically-active SED (**Figure 2n**). As per the whole ileal cross-section data (**Figure 2a**), in all sections analysed across the four mice, the villous mucosa was entirely devoid of fgTiO_2_-positive cells and consistently exhibited the same reflectance distribution as the negative control mice (**Figure 2n**).

### Establishing correlative human-mouse dosimetry

Rodent models provide pragmatic access to healthy pristine tissue (*i.e.,* rapidly flash frozen at necropsy) from fully controlled experiments (*i.e.,* population-matched with availability of fgTiO_2_-negative tissues) in a way that human surgical specimens do not. Nonetheless, it is critical that a murine feeding model appropriately reports on the human situation. Next, therefore, we exploited the window-of-insight provided by precision image analysis to consider how the cellular dosimetry of fgTiO_2_ in the SED cells of the murine model reflected SED-cell exposure in humans. To do this, the snap-frozen Peyer’s patch tissue sections, taken from six randomly-drawn human samples, were compared with six murine samples taken from the fgTiO_2_-fed group. Visually, in both species, there was marked inter-individual variability in fgTiO_2_ concentration in the SED (**Figure 3a-L**). To quantify this, we analysed the thresholded reflectance signal in the tissue sections in four ways, namely: thresholded reflectance per unit tissue area (*i.e.,* fgTiO_2_ abundance; **Figure 3m),** thresholded reflectance per cell (*i.e.,* fgTiO_2_ load per cell; **Figure 3n**), the number of reflectant foci per cell (*i.e.,* cellular count of fgTiO_2_-loaded vesicles; **Figure 3o**), and finally the amount of reflectance signal per foci (*i.e.,* fgTiO_2_ load per vesicle; **Figure 3p)**. The strategy is schematically explained in **Supplementary Figure 3**; cell segmentation accuracy assessments are presented in **Supplementary Figure 2**; background reflectance comparisons in mice and humans are shown in **Supplementary Figure 4** and examples of the raw reflectance signal, resulting circle-marker placements and fgTiO_2_-loaded vesicle segmentations are shown in **Supplementary Figure 5**. By all metrics, the data from the two groups showed remarkable overlap and under statistical comparison none of the measured fgTiO_2_ distributions were found to be entirely unique to either species (presented, **Supplementary Figure 6**). Notably when Peyer’s patch loading of fgTiO_2_ from mice on lower-dose-fgTiO_2_ diets was analysed, the cellular accumulation of particles in the SED was several orders of magnitude lower compared to the human situation (**Supplementary Figure 7**). The above findings therefore establish the apparent ability of the murine model to provide human-relevant data – quantitatively as well as qualitatively – in terms of fgTiO_2_ cell targeting and dosimetry in the biologically-active SED region of the Peyer’s patch.

**Figure 3.**
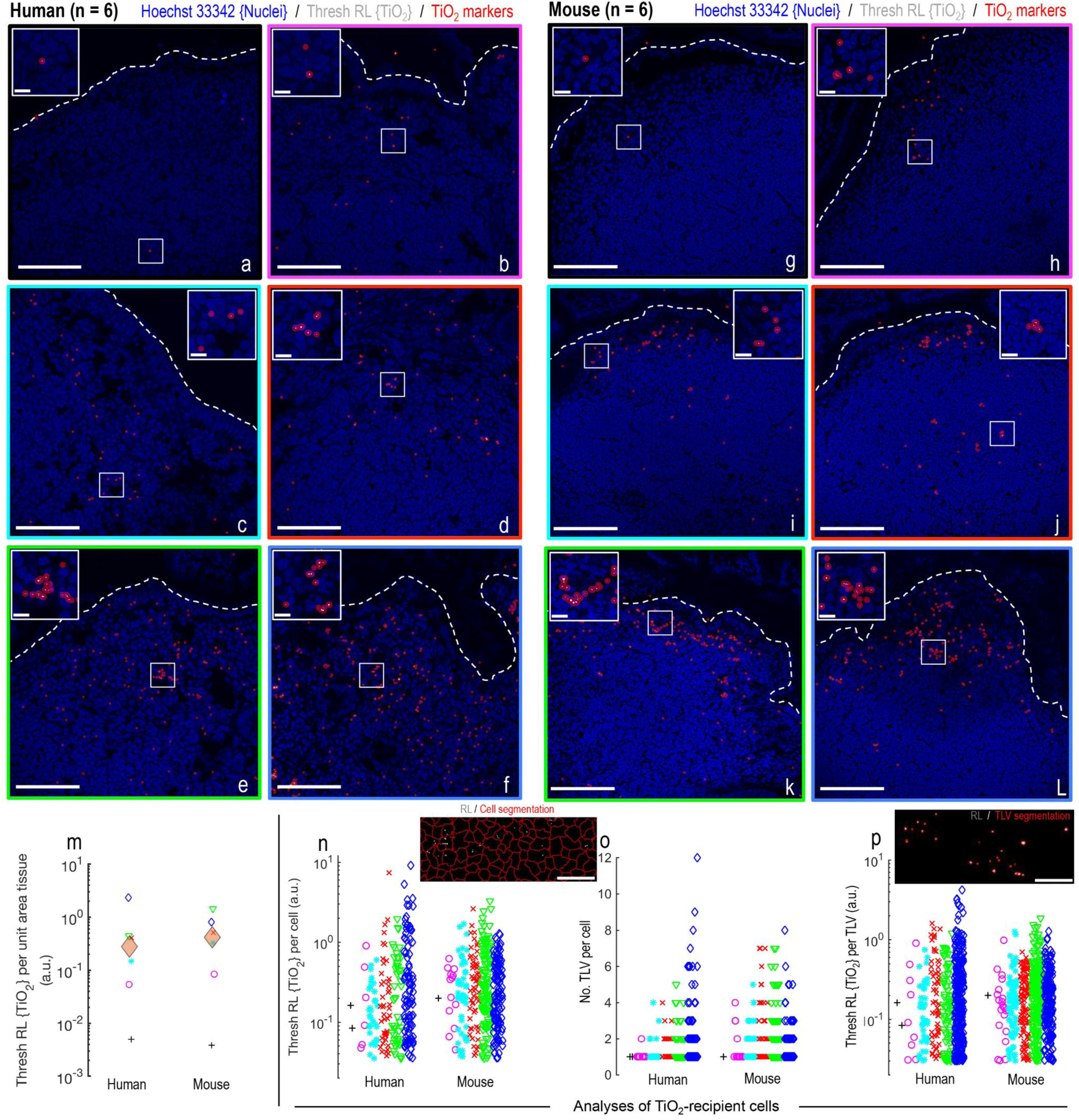
Establishing correlative human-mouse dosimetry. **a-L**, Confocal reflectance microscopy showing the range of cellular loading in the subepithelial dome tissue region of (**a-f**), six randomly-drawn human and (**g-L**) six randomly-drawn mouse samples. After acquisition, the image-fields per specimen were manually laid out in order of lowest-to-highest fgTiO_2_ SED accumulation to visually present the wide variation in fgTiO_2_ cellular loading. Translucent red circle-markers were placed on thresholded reflectant foci to aid visualisation. **m-p**, Quantitative analysis of the image-data shown in **a-L**. **m**, Thresholded reflectance per unit tissue area (*i.e.,* amount of fgTiO_2_ per unit tissue area). **n**, Thresholded reflectance per cell (*i.e.,* fgTiO_2_ dose per cell). **o**, Number of thresholded reflectant foci per cell (*i.e.,* number of fgTiO_2_-loaded vesicles (TLV) per cell). **p**, Thresholded reflectance signal per foci (*i.e.,* fgTiO_2_ dose per TLV). Statistical comparison of the distributions is presented in **Supplementary Figure 6** (two-sided Wilcoxon rank-sum analysis). None of the measured fgTiO_2_ distributions were found to be entirely unique to either species. **m-p**, By all of the quantitative measures established, feeding a murine diet supplemented with 0.0625% (w/w) fgTiO_2_ for eighteen weeks provides significant overlap with measured, real-world human exposures. *Scale bars: **a-L** = 50 µm with insets 10 µm; **n** = 20 µm; **p** = 5 µm*.

### fgTiO_2_ selectively targets immunocompetant cells

The fgTiO_2_-targeted cells in the SED were, unsurprisingly, CD11c^+^ (*i.e.,* phagocytic mononuclear cells) and not only was the frequency of such cells unaffected by the presence of fgTiO_2_ particles so was their expression of the CD11c integrin biomarker (**Figure 4a-h**). Neighbouring cells were chiefly CD3^+^ (T cells) and B220^+^ (B cells) and, similarly, their expression patterns and frequencies were not significantly affected by the presence of fgTiO_2_ (P ≥ 0.26, two-sided unpaired samples T-tests) (**Figure 4a-h**). Within the Peyer’s patch, CD11c^+^ cells are known to exhibit a number of sub-phenotypes in distinct follicular locations^17^. In this regard, the SED localisation and autofluorescence of the fgTiO_2_^+^ cells were entirely consistent with ‘LysoMacs’ and/or ‘LysoDCs’ which represent extensively-described macrophage and dendritic cell subsets, respectively, that express high levels of lysozyme alongside a unique autofluorescence signature^17, 23^ (see **Figure 4i-s** and **Supplementary Figure 8** for the autofluorescence spectra collected). Indeed, phenotypic markers that help differentiate LysoDCs from LysoMacs (cell surface CD4 and MHCII) demonstrated a mixed population of these two cell types for the fgTiO_2_^+^ cells (**Figure 4t/u**).

**Figure 4.**
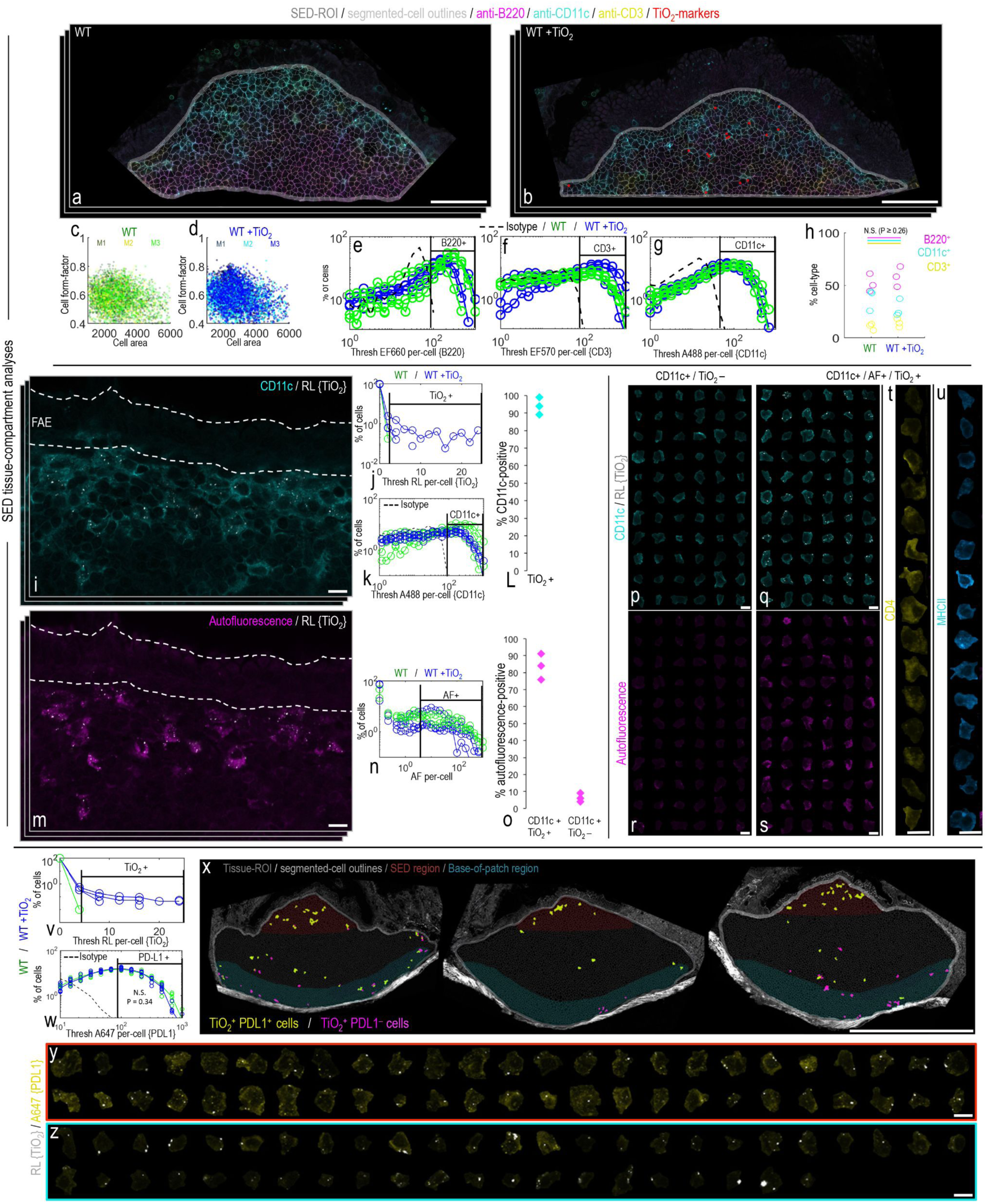
fgTiO selectively targets PD-L1^+^ LysoMac and LysoDC cells. **a-h**, *In situ*, single-cell analysis of key immune-cell subtypes of the mouse subepithelial dome. **a/b**, Example images immunofluorescently labelled for B220, CD3 and CD11c (*i.e.,* identifying B-lymphocytes, T-lymphocytes and phagocytic mononuclear cells, respectively) from wild-type (WT) mice fed (**a**) without (WT) or (**b**) with (WT + TiO_2_) fgTiO_2_ dietary supplementation (0.0625% w/w of diet). fgTiO_2_ was measured from thresholded reflectance images. Translucent red circle-markers are placed on reflectant foci to aid visualisation. **c/d**, Flow cytometry-type plots showing cell area / cell aspect ratio, (**e-g)** immunofluorescence distributions and (**h**) cell counts for B220^+^, CD11c^+^ and CD3^+^ cells (n = 3 animals). All measures remained similar regardless of fgTiO_2_ feeding (cell count comparison (**h**), P = ≥0.26, two-sided unpaired samples T-tests). **i-L**, fgTiO_2_-recipient cells of the subepithelial dome were CD11c^+^ in ∼ 95% of cases. **m-o**, These cells also exhibited a unique, highly autofluorescent signature as previously-described for subepithelial dome ‘lysomac’ and ‘lysoDC’ mononuclear phagocyte cells. **o**, Again, this was true for almost all fgTiO2^+^/CD11c^+^ cells and, equally, these autofluorescent cells were similarly present in (**n**) controls without fgTiO_2_ feeding (further data shown, **Supplementary Figure 8**). **p-u**, Montaged CD11c^+^/fgTiO_2_ ^+^ or CD11c^+^/fgTiO_2_ ^-^ cell populations from the SEDs of mice fed the fgTiO_2_ supplemented diet visualised in terms of **(r/s)** autofluorescence signature. fgTiO_2_ is seen to both selectively and specifically target the autofluorescent cells and (**t/u**) the diversity of CD4 and MHCII expression confirms that both lysomac and lysoDCs take up fgTiO_2_. **v-z**, fgTiO_2_ ^+^ cells of the SED expressed the immunotolerance marker programmed death ligand 1 (PD-L1) whereas fgTiO_2_ ^+^ pigment cells at the base of the Peyer’s patch tended to lose PD-L1 expression (**y** versus **z**, respectively). **w**, PD-L1 expression in the Peyer’s patches was not significantly perturbed by feeding the fgTiO_2_-supplemented diet (P = 0.34, z = -0.96, two-sided Wilcoxon rank-sum analysis, n = 6 animals). *Scale bars: **a/b** = 50 µm; **i/m**, **p-u** = 10 µm; **x** = 500 µm; **y/z** = 10 µm*.

Whilst similar populations of autofluorescent^hi^ SED cells were observed in all mice regardless of diet, these cells were strongly particle-positive in mice receiving the dietary fgTiO_2_ (**Figure 4m/s, Supplementary Figure 9** and **Supplementary Video 1**). Upon quantification (**Figure 4o**), only ∼7% of autofluorescent cells showed no clear particle signal for the +fgTiO_2_ group (**Figure 4i-s**). These cells are known to be important in the initial uptake and processing of luminal antigen which other work suggests may be coated in calcium phosphate to facilitate enzyme resistance during the early phases of the lumen-to-phagocyte sampling process^24^. Except in Crohn’s disease, these cells have been shown previously to be positive for the immunoregulatory cell surface ligand PD-L1^25^. Again, consistent with this prior work, our quantitative single-cell tissue analyses showed that the fgTiO_2_^+^ (and therefore CD11c^+^, autofluorescence^hi^) cells of the SED were PD-L1^+^ (**Figure 4v-y**, antibody controls presented **Supplementary Figure 10**). This contrasted with precursor pigment cells at the base of the Peyer’s patch which were PD-L1^lo^, despite their fgTiO_2_ content (**Figure 4v-z**). In other words, based on this finding, and that mice unexposed to fgTiO_2_ show similar PD-L1^+^ distributions (**Figure 4w)**, we can conclude that expression of this immunoregulatory molecule is unaffected by fgTiO_2_ uptake (P = 0.34, z = -0.96, two-sided Wilcoxon rank-sum analysis) and, rather, fgTiO_2_ is selectively and specifically shuttled into already-PD-L1^+^ LysoMacs/LysoDCs in the SED of Peyer’s patches in the small intestine. As such, for the first time, we show that there *is* an immunocompetent cell population that accumulates fgTiO_2_ in humans, and in a rodent model, which will now allow the proposed adverse effects of the particulates to be tested.

### Potential for fgTiO_2_ – *Salmonella* interactions

As noted above, in earlier *in vitro* work, we have shown that the phagocyte proinflammatory response can be augmented by cellular loading with fgTiO_2_^1^. As such “Mouse Study 2” considered the potential for fgTiO_2_ to initiate or augment an inflammatory response, precisely in the Peyer’s patch-rich, terminal ileal region where the particles accumulate. Initiation was measured directly whilst, for augmentation, we investigated the synergy between fgTiO_2_ and an orally-delivered auxotrophic *Salmonella* strain, namely Δ*aroA*-deficient *Salmonella enterica* serovar Typhimurium (Δ*aroA*-*Salmonella*). The choice of bacterial infection was because (i) again, it is human-relevant (*i.e.,* commonly experienced^26^) (ii) it targets LysoDCs and LysoMacs^27^ resulting in a modest immuno-inflammatory response (unlike bacterial fragments, to which the intestine is uniquely tolerant) and (iii) this response can be augmented leaving headroom for any interactions to be observed^28, 29^. Regarding this latter point, it is known that same-cell concurrence of an amplification signal, such as over-expressed bacterial flagellin or co-expressed vectorised cytokines, is required to augment the *Salmonella-*induced immuno-inflammatory response^29, 30^. For this reason, we first demonstrated substantial same-cell occurrence of the particles and bacteria, in the SED of Peyer’s patches, when fgTiO_2_-fed mice (for 16 weeks) were immediately chased with a single oral dose of Δ*aroA*-*Salmonella* and tissues collected 3 days later (+ 3 days; **Figure 5a-c**). *Salmonella*-specific faecal IgA was undetectable at this time-point but was clearly demonstrated in a second group at + 28 days following the *Salmonella* challenge. This pattern was irrespective of fgTiO_2_ ingestion (**Figure 5d**) and is consistent with a latent immuno-inflammatory response following oral exposure to the pathogen^31, 32^.

**Figure 5.**
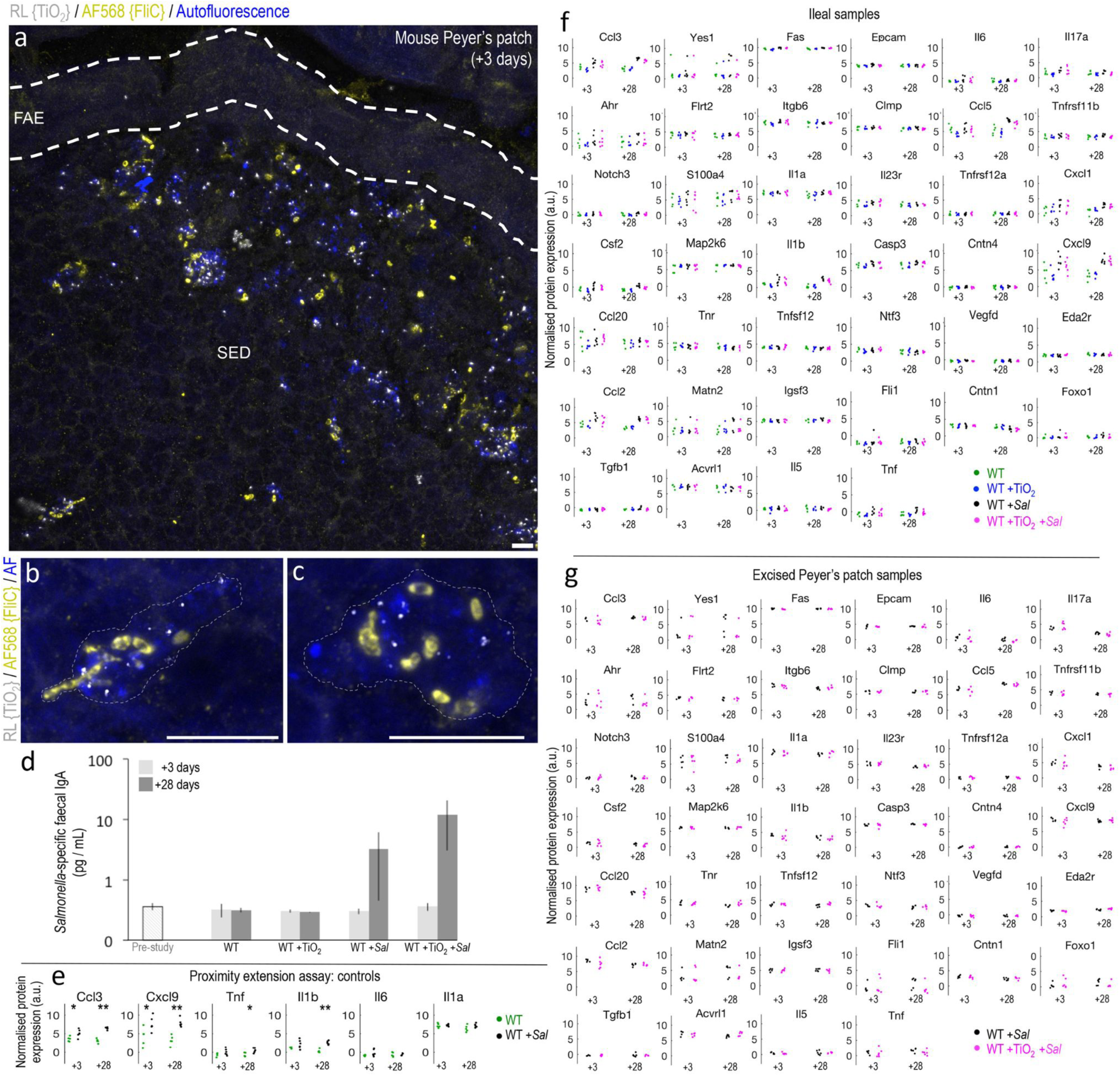
fgTiO_2_-*Salmonella* interactions. To enable studies of particle adjuvanticity and cell migration, in “Mouse Study 2”, after the 16-week fgTiO_2_ feeding period all animals were switched to a normal laboratory chow diet (*i.e.,* without fgTiO_2_-supplementation) then half were orally inoculated with attenuated, Δ*aroA*-*Salmonella*. Tissues were harvested +3 or +28 days after infection. **a-c**, Confocal reflectance micrographs displayed as z-stack maximum projections showing immunofluorescent labelling for flagellin (FliC) to detect the Δ*aroA*-*Salmonella* in the Peyer’s patch SED region. Marked accumulation of both fgTiO_2_ and Δ*aroA*-*Salmonella* was observed in autofluorescent, phagocytic mononuclear cells at the +3 timepoint. **d**, Faecal ELISA studies confirmed a marked, *Salmonella*-specific IgA response at the +28 timepoint (bars represent medians with interquartile range error-bars, n = 3-6 samples per treatment group). **e-g**, Protein expression analyses of ileal tissue digests by proximity extension assay (OLINK mouse exploratory panel). **e,** Comparison of ileal tissues taken from wild-type (WT) mice unexposed to fgTiO_2_ but with (WT *+Sal*) or without (WT) Δ*aroA*-*Salmonella* infection confirmed significant increases in expression of key cytokines / chemokines associated with a Th1 immune response (P ≤ 0.026, two-sided Wilcoxon rank-sum test, n = 5-6 tissues per group). **f**, Inflammation-associated protein expression analyses of ileal tissue digests in the WT, WT +TiO_2_, WT +*Sal* and WT +TiO_2_ *+Sal* treatment groups at the +3 and +28-day timepoints. No significant differences were observed between the WT and WT +TiO_2_ (P ≥ 0.53, two-sided Wilcoxon rank-sum test, n = 3 animals per group) or WT + *Sal* and WT +TiO_2_ +*Sal* groups (P ≥ 0.79, two-sided Wilcoxon rank-sum test, n = 6 animals per group). Full dataset is shown in **Supplementary Figure 12**. **g**, Peyer’s patch-focussed inflammation-associated protein expression analyses using carefully-excised patch digests in mice treated with either Δ*aroA*-*Salmonella* alone (WT *+Sal*) or with fgTiO_2_ and Δ*aroA*-*Salmonella* (WT +TiO_2_ *+Sal*) at +3 or +28 timepoints. No significant differences were observed between the two groups (P ≥ 0.2, two-sided Wilcoxon rank-sum test, n = 6 animals per group) at either timepoint demonstrating no interaction of fgTiO_2_ and Δ*aroA*-*Salmonella* despite (**a-c**) heavy loading into the same cells (full dataset shown in **Supplementary Figure 13)**. OLINK protein abbreviations are defined in **Supplementary Table 1**. *Scale bars: **a-c** = 10 µm*.

To investigate both initiation and augmentation of an inflammatory response, and to cover a broad array of potential outcomes, we undertook protein quantification using the sensitive and accurate (*i.e.,* dual antibodies per target) proximity extension assay (Olink Mouse Exploratory Panel) which provided data for 92 different protein biomarkers (repeat-sample and dilution controls presented, **Supplementary Figure 11**; full protein names shown in **Supplementary Table 1)**. Since *Salmonella* Typhimurium induces a Th1 profile in the ileum^33^ we first confirmed, using this analytical approach, that associated cytokines/chemokines were enhanced in ileal tissue from mice that had been challenged with Δ*aroA*-*Salmonella*. Of Th1 relevance, the proximity extension panel detects IL-1 (α&β), TNFα, IL-6, CXCL9 and CCL3^33, 34^ and four of these markers (CCL3, CXCL9, IL1β and TNFα) showed significantly (P ≤ 0.026, two-sided Wilcoxon rank-sum tests) increased protein expression in the ileum of Δ*aroA*-*Salmonella-*exposed mice versus unexposed matched controls at the +28 day time-point (**Figure 5e**). Next we considered the effect on the ileum of chronic oral exposure to fgTiO_2_ (*i.e.,* mimicking the Peyer’s patch loading seen in humans) in the absence of *Salmonella.* The proximity extension assay data showed remarkable consistency between the two groups for all measurable proteins and, therefore, no suggestion that long-term fgTiO_2_ ingestion initiates inflammatory signalling in the ileum of wild-type mice (**Figure 5f** and **Supplementary Figure 12**). In a similar fashion, we considered the effect on the ileum of chronic oral exposure to fgTiO_2_ plus *Salmonella* infection. Again, not one protein differed significantly in its expression between the groups receiving *Salmonella* and *Salmonella* + fgTiO_2_ (**Figure 5f** and **Supplementary Figure 12**) indicating no measurable augmentation of fgTiO_2_ on *Salmonella*-induced immuno-inflammatory signalling in the ileum of wild type mice, in spite of substantial same-cell uptake of the bacteria and the fgTiO_2_ particles. To ensure that some small effect, localised to the Peyer’s patch as the site of particle accumulation, had not been missed (*e.g.,* ‘diluted-out’ by surrounding, unaffected tissue) we made the same analyses for a potential augmentation effect using ileal tissue samples that were trimmed tightly to either side of Peyer’s patch such that each sample primarily consisted of only the patch itself. But, again, no significant differences in protein expression were measureable by the Olink array (**Figure 5g** and **Supplementary Figure 13**).

### Cellular dosimetry and ileal tissue distribution

Finally, we considered what factors influenced fgTiO_2_ distribution in the Peyer’s patch. In the absence of *Salmonella* challenge we observed fgTiO_2_ in autofluorescent cells of the Peyer’s patch SED region, as well as some further towards the base of the patch, at both the +3 and +28 day time-points (**Figure 6a/b**), consistent with observations in Mouse Study 1. Quantitatively, however, between the two time points, there was a measurable increase in the number of fgTiO_2_-loaded vesicles per cell, without a change in load per vesicle, in the mid-region of the follicle (**Figure 6c-j**), suggesting a natural history of vesicular inheritance by certain cells and gradual development towards the well-reported basal pigment cells^15, 16, 20^. Along this pathway, cells maintained the same autofluorescent profile and mixed MHCII and CD4 expression as described in Mouse Study 1 suggesting that any such process does not give rise to a new population of cells. Visually however, with increasing distance from the SED, an increase in CD4 expression in conjunction with a decrease in MHCII expression was observed, suggesting an shift to sequestration in cells with the longer-lived, LysoMac phenotype (shown, **Supplementary Figure 14)**. A more remarkable influence on the spatial distribution of fgTiO_2_ was caused by the single oral dose of Δ*aroA*-*Salmonella*. SEDs were almost entirely devoid of fgTiO_2_ and autofluorescence-positive cells at +28 days (**Figure 6k**), while some cells of the regular villous lamina propria, as well as of the basal aspect of the Peyer’s patch, were now positive for fgTiO_2_ (**Figure 7a-c**; enlarged display with counterstaining context, **Supplementary Figure 15**). Consistently, it has been reported that oral danger signals, including *Salmonella*, deplete the Peyer’s patch SED region of prior-absorbed plastic microparticles^35^. The pathway that then enables appearance of fgTiO_2_ in sporadic cells of the regular mucosa, as we show here, is not clear but (i) fgTiO_2_ loading per cell was low versus most Peyer’s patch cells (**Figure 7b/c**) and (ii) many of these cells showed an autofluorescent signature (**Figure 7d**). As such, LysoDC migration from the patch with re-homing to the gut mucosa should be considered.

**Figure 6.**
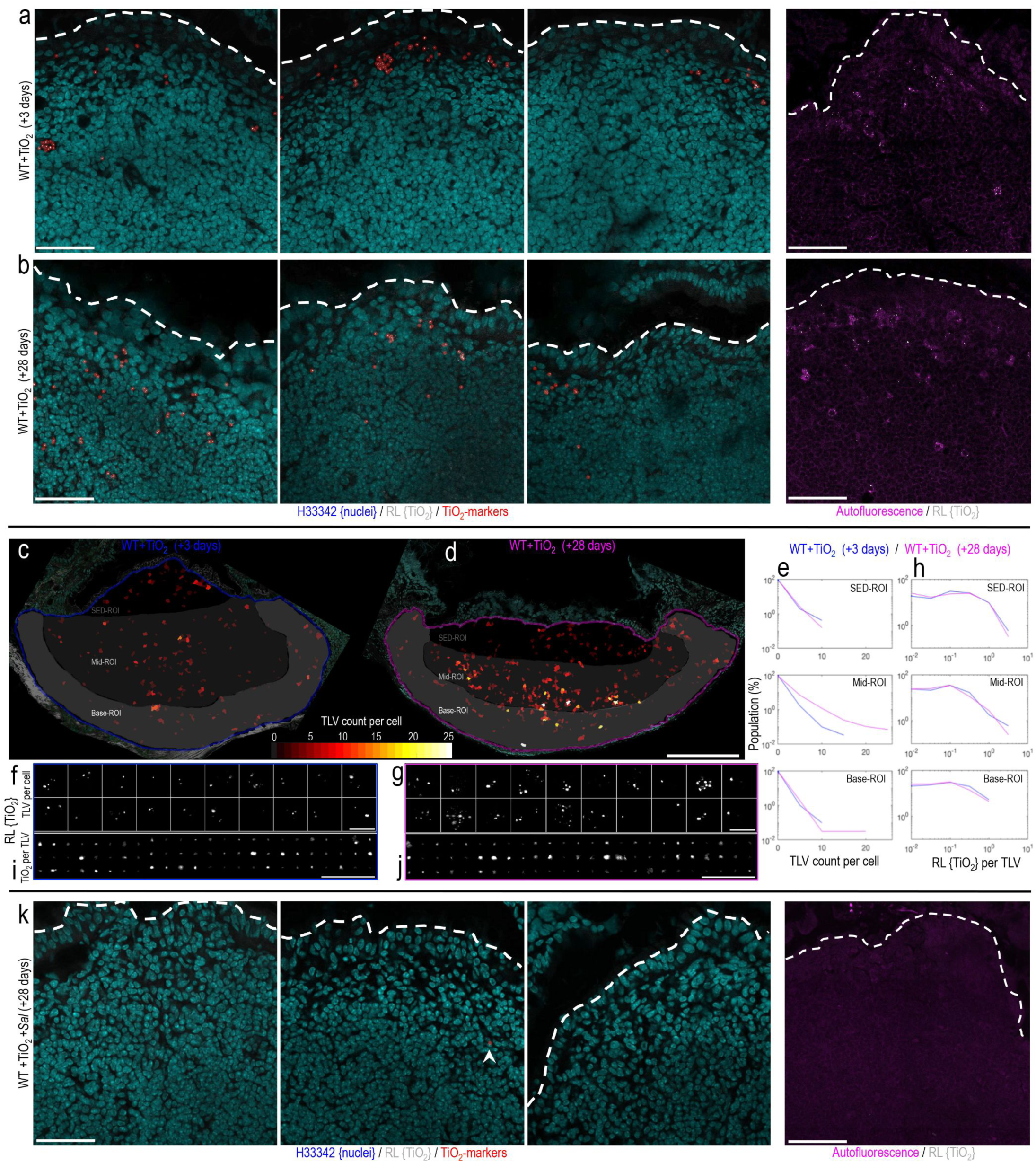
Cellular dosimetry of fgTiO_2_ across timepoints and *Salmonella*-induced migration. **a/b/k**, Confocal reflectance micrographs from Mouse Study 2 showing the subepithelial dome (SED) tissue region (n = 3 animals per group). After the 16-week fgTiO_2_ feeding period, all animals were switched to normal laboratory chow diet (without fgTiO_2_ supplementation) and half were orally dosed with Δ*aroA*-*Salmonella* before tissue harvest at +3 or +28-day timepoints. **a/b**, In the animals fed the fgTiO_2_-supplemented diet without *Salmonella* exposure (WT +TiO_2_), fgTiO_2_-loaded cells were present in the SEDs at both +3 and +28 timepoints. **c-j**, To gain insights into cellular dosimetry changes across the two timepoints, quantitative image analysis of complete Peyer’s patch transverse sections was used to precisely measure the number of reflectant foci per-cell (*i.e.,* the fgTiO_2_-loaded vesicle (TLV) count per-cell) and the amount of reflectance per foci (*i.e.,* equivalent to the fgTiO_2_ dose per-vesicle). **d**/**e**, At the +28 timepoint, the TLV count per-cell was markedly elevated with a new population of cells with TLV counts >13 forming in the mid-to-base regions of the follicle. **f/g**, Randomly sampling and montaging the reflectance (RL) information in cells from each of the two timepoints visually demonstrates this, showing (**g**) more heavily-loaded cells at the +28 timepoint. **h**, Moving from cell to vesicle information, in contrast, the amount of reflected light per individual foci remained near-identical across the two timepoints and this was again borne out visually when (**i/j**) individual TLVs were randomly montaged. Collectively, the results suggest that once fgTiO2-loaded cells move below the SED, there is a process of vesicular inheritance by certain cells over time, providing a route to the formation of heavily-loaded pigment cells at the follicle base – as is well-described in humans (Figure 1). **k**, Challenge with Δ*aroA*-*Salmonella* (WT +TiO_2_ +*Sal*) markedly changed this picture (n = 4 animals per group). At the +28 timepoint, SEDs were almost completely devoid of both fgTiO_2_ and autofluorescence-positive cells. Out of four SEDs imaged, only one reflectant foci was detectable (indicated by arrow) showing marked migration of fgTiO_2_ from the SED tissue compartment after Δ*aroA*-*Salmonella* exposure. *Scale bars: **a**/**b**/**k** = 50 µm, **c**/**d** = 250 µm, **f**/**g**/**i**/**j**, = 10 µm*.

**Figure 7.**
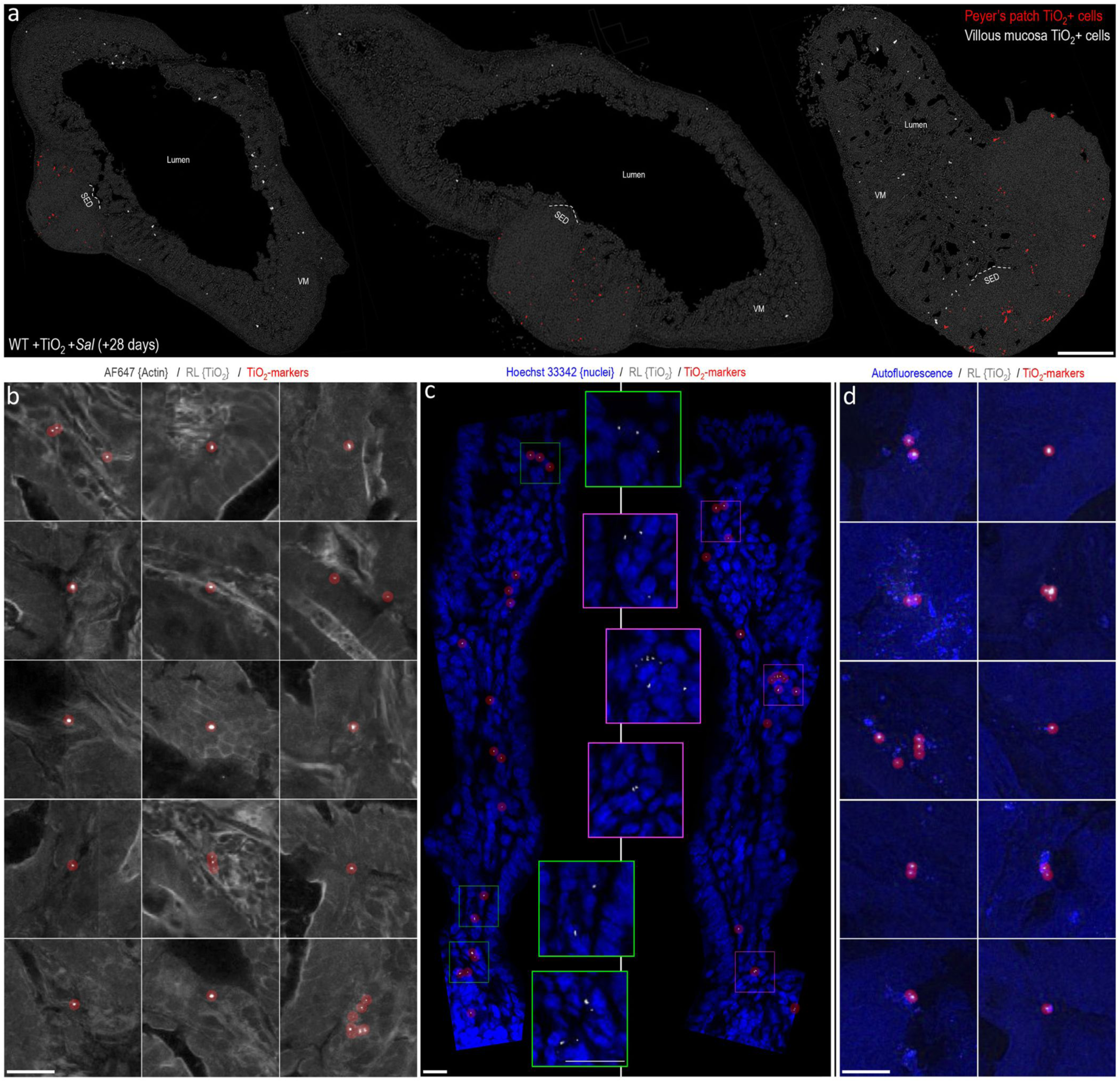
Ileal distribution of fgTiO_2_ following *Salmonella* challenge. **a**, Single-cell image analysis of tilescanned confocal reflectance images collected from transverse sections of mouse ileal tissue from the fgTiO_2_-exposed, Δ*aroA*-*Salmonella*-infected treatment group (+28-day timepoint, n = 3 animals). fgTiO_2_-positive cells in the Peyer’s patches are displayed in red. As in Figure 6, the subepithelial dome (SED) regions were devoid of fgTiO_2_ with the majority of the positive cells located basally around the follicle margins. In contrast to all previous data collected without Δ*aroA*-*Salmonella* exposure, fgTiO_2_-positive cells (displayed in white) were now also present in the villous mucosa. Bigger versions of the single-cell analyses presented in (**a**) are available in **Supplementary Figure 15**. **b**, Montaging positive events from villous mucosa regions showed that particle loading was low compared to Peyer’s patch cells with the majority containing single reflectant foci. **c**, High resolution imaging of optically cross-sectioned villi confirmed reflectant foci deep in the lamina propria in the same manner as observed in humans (Figure 1c). **d**, The majority of the fgTiO_2_-positive events in the villous mucosa also displayed the same autofluorescence signature as was observed in the particle-recipient cells of the SED. *Scale bars: **a** = 250 µm; **b**-**d** = 20 µm*.

## DISCUSSION

In this work, we have exploited the growing accessibility of machine learning methods^36^ to enable precise analyses of complex images yielding *in situ* single-cell and single-vesicle data from tissue samples with full locational data retained. Particle selectivity of cellular targets in distinct tissue regions can thus be assessed^37, 38^. Access to quantitative single-cell and vesicle measurements has provided an unprecedented opportunity to compare and establish cell-dose equivalence between humans and mice. The findings move the field forward in four important areas of persistent particle exposure via the oral route, namely: (i) human relevance (ii) mechanism of absorption and intestinal distribution (iii) role in intestinal inflammation and (iv) provision of a model for cogent risk assessment.

### Human relevance

The human small intestine is the only known site of cellular accumulation of dietary microparticles, as exemplified by fgTiO_2_^16, 18, 20^ (**Figure 1**). We show here that all features of this can be qualitatively and quantitatively recapitulated in a murine model, but full achieving this requires exposure to human-relevant microbial ‘danger signals’. Consistently, in this work, murine dosing with fgTiO_2_ was undertaken in a manner that reflects real-life exposures (*i.e.,* via the diet at relatively low levels for a long period of time). Our findings contrast with a prior study, which used bolus dosing via oral gavage or surgical intervention in fasted animals, where direct fgTiO_2_ uptake via the epithelium was reported^39^. Future work may wish to consider the necessity of natural ingestion and concomitant normal microbial exposures, versus non-physiological dosing in specific pathogen free environments, to recapitulate the human situation and address risk accurately for oral particle exposures.

### Mechanism of absorption and intestinal distribution

Seminal early work from Frey and Mantis^40, 41^ showed that the mucus-rich glycocalyx markedly impairs model particle uptake by the regular villous epithelium, whereas the near-absence of these features overlying Peyer’s patches allows direct access of particulates to M-cells. These cells have an extraordinary ability to engulf luminal particles and macromolecules, passing them through to abutting cells at their basolateral membrane. We first confirmed, unequivocally, that this uptake mechanism holds true for fgTiO_2_ and, next, demonstrated that specific SED immuno-competent phagocytes (LysoMacs and LysoDCs) are the direct recipient cells that accumulate cargo from the M-cell. Taking all of our data together, we believe that there is an on-going process of slow re-arrangement, typified by vesicular inheritance by some of the cells and gradual particle-loaded-cell movement to the base of the patch. This migration (presumed to occur in LysoMacs), which must ultimately result in mature pigment cell formation^16, 18^, is greatly accelerated by oral exposure to the bacterial pathogen *S.* Typhimurium. In addition, some particulate migration out of the patch, to the villous lamina propria, also occurs (typically low-loaded cells, presumed LysoDCs). Again, building on formative work on the fate of reporter particles of fluorescent plastic in the Peyer’s patch^35^, it is likely that the ‘danger signal effect’ holds true for oral exposure to soluble enterotoxin adjuvants as well as pathogenic bacteria (*i.e.* multiple human-relevant danger signals). Continuous exposure to fgTiO_2_, as happens for many human populations, would then re-load the apical (SED) phagocytes until another danger signal event occurs. Thus, for the first time, we describe the natural history of persistent particle routing through – and beyond – the Peyer’s patch following oral exposure. This explains basal pigment cell accumulation of multiple particle types, including titanium dioxide, silica and aluminosilicates, in human Peyer’s patches as well as the occurrence of sporadic particles elsewhere in cells of the small intestine^16, 18, 20^.

### Role in intestinal pathology

By extension of the above observations, our findings also imply that fgTiO_2_ is not in itself perceived as a danger signal in the intestine of wild-type mice since a secondary (microbial) signal is still required for the ‘acute’ migration. Moreover, using highly sensitive and selective protein analyses, that captured a broad array of biomarkers of immuno-inflammatory signalling, we were unable to identify either direct or augmented perturbation of tissue homeostasis at the site of particle accumulation. Indeed, mammals have very likely evolved with oral exposure to other fine particles (*e.g.,* geologically-derived and soot). Certainly aluminosilicates, in abundance, and some forms of silica are reported in human Peyer’s patches alongside titanium dioxide^16, 18^. Perhaps therefore, effective compensatory mechanisms generally exist, which may include constitutive cell expression of immuno-inhibitory PD-L1 (**Figure 4v-z**).

Notwithstanding, the potential for fgTiO_2_-induction or augmentation of intestinal pathology still deserves scrutiny. First, in subjects with Crohn’s disease, fine-particle-targeted macrophages of the Peyer’s patch do *not* express PD-L1^25^. Interestingly, Crohn’s disease is a relatively modern inflammatory disorder and there is good evidence that it may initiate in the Peyer’s patches^42^. The most commonly-associated single gene mutation with the disease is NOD2 and there is *in vitro* evidence that when NOD2 is perturbed fgTiO_2_ augments enhanced inflammatory activity with microbial fragments^1^. So it should now be carefully considered what impact the ingestion of fgTiO_2_ has on at-risk genotypes and, especially, where NOD2 functionality is diminished or removed. Secondly, it is important to note that conclusions from this work refer to the distal small intestine where there is uptake of fgTiO_2_. As noted previously^12^, however, it does not shed light on potential effects of fgTiO_2_ where it is *not* absorbed such as in the large intestine. Direct luminal effects (on microbiome or apical enterocytes for example) cannot be precluded, especially given that this is where exposure will be by far the greatest. Indeed, both Urrutia-Ortega *et al.,*^43^ and Bettini *et al.,*^44^ have provided evidence for direct, pro-tumorigenic effects of fgTiO_2_ on the colon which deserves further attention^12^.

### A model for cogent risk assessment

Finally, we note that the particulate-accumulating population of apical SED cells are clearly (i) immunocompetent (*i.e.,* non-quiescent) based upon their migration characteristics, expression of cell surface immunoregulatory PD-L1 and their previously-described qualities in terms of antimicrobial defence^17, 27, 45, 46^ and (ii) capable of becoming heavily loaded with fgTiO_2_. Indeed, the exquisite particle selectivity for this tissue region means that the concentration gradient for area-corrected reflectance signal was ∼14-fold higher than for the Peyer’s patch overall and, therefore, ∼1,245 fold higher than that of villous mucosa (**Supplementary Figure 16**). Given that the particulate-loaded phagocytic cells make up only a minority of the SED^17, 23^, then the cell concentration of fgTiO_2_ is *very* high (**Supplementary Video 1**).

We know of no other active particle-funnelling and specific cell-targeting strategy elsewhere in mammalian biology so it is very likely that SED LysoMacs and LysoDCs provide the first and only target to enable highly sensitive, single-cell risk assessment of fgTiO_2_ and other persistent oral particulates *in vivo*. By exemplification, we addressed the question of whether fgTiO_2_ could initiate or augment inflammatory signaling in the ileum / Peyer’s patches of wild type mice: a long standing question in the field. The clear answer was ‘no’, at least in wild-type genotypes, despite some *in vitro* evidence to the contrary^1^. To explain such differences it is worth noting that cell culture studies fail to replicate *in vivo* fidelity in many ways including rates of particle uptake, presence of functional lymphatics and vasculature, cellular cross-talk, compensatory mechanisms and cell migration prior to particle saturation (cell gorging). Whatever the reasons, after decades of uncertainty, other questions regarding the safety of persistent oral particles can now be precisely addressed using *in vivo* studies with human-relevant dosimetry and identified target cells. Indeed, understanding the consequences of LysoMac / LysoDC accumulation of persistent particles in terms of genotoxicity and any differential responses resultant from distinct host genotypes is now key to understanding human risk following oral exposure to nanoparticles and microparticles.

## METHODS

### Mouse Study 1

The first mouse fgTiO_2_ feeding study was approved by the Grasslands Animal Ethics Committee (Palmerston North, New Zealand) in accordance with the New Zealand Animal Welfare Act 1999. Food-grade, anatase TiO_2_ particles (fgTiO_2_) were purchased from Sensient Colors (St. Louis, MO, USA). Physico-chemical characterisation of these particles in our previous work shows a particle diameter distribution of ∼ 50 – 300 nm, with a median diameter of ∼ 130 nm^1^. As described and extensively characterised previously^15^, to enable successful oral delivery, the standard rodent diet American Institute of Nutrition (AIN)-76A was supplemented with 0.000625%, 0.00625% or 0.0625% (w/w) fgTiO_2_ (translating to approximately 1, 10 or 100 mg fgTiO_2_ / kg body-weight / day respectively^15^). These diets were prepared by Research Diets (New Brunswick, NJ, USA), with straight AIN-76A without fgTiO_2_ supplementation used as the ‘negative control diet’. Six-week-old mice (C57BL/6, 50:50 male:female) were housed conventionally with normal feed intake and weight-gain confirmed bi-weekly^15^. After eighteen weeks feeding, mice were euthanised by CO_2_ asphyxiation and cervical dislocation. The gastrointestinal tract was quickly removed and placed in cold phosphate-buffered saline (PBS). Ileal tissues containing the most distal Peyer’s patches (*i.e.,* closest to the ileal/caecal junction) and adjacent caecal patches were excised and frozen on a cold, stainless steel bar embedded in dry ice. Tissues were then transferred into cryomolds filled with pre-cooled optimal cutting temperature compound (OCT) and transferred to – 80°C for storage.

### Mouse Study 2

Animal experiments conducted at Cambridge were performed under the authority of Home Office personal license number PIL IEDB62633, and project license number PPL PF86EABB1. Female mice (C57BL/6) were purchased from Charles River Laboratories (UK). Upon arrival, mice were housed conventionally and were monitored for signs of normal health, food ingestion and body-weight gain throughout the study period. At the start of the study, thirty-six mice (six weeks old) were randomly divided into two feeding groups. As before, one group received the standard rodent diet American Institute of Nutrition (AIN)-76A supplemented with 0.0625% (w/w) fgTiO_2_ (translating to approximately 100 mg fgTiO_2_ / kg body-weight / day respectively^15^) whereas the other received straight AIN-76A without fgTiO_2_ supplementation (*i.e.,* the negative control diet). After sixteen weeks of feeding, all mice were switched to the negative control diet, and twelve mice from each diet-group were orally challenged with Δ*aroA-*deficient, *S*. Typhimurium strain SL3261^47^ (Δ*aroA*-*Salmonella)*. We marginally separated the administrations of bacteria and fgTiO_2_ to (i) prevent artefactual adsorption due to co-administration (ii) enable luminal interactions between gavaged bacteria and residual luminal particles (demonstrated by faecal ICP-MS studies, **Supplementary Figure 17**) should such interactions occur and (iii) ensure that some food also was in the lumen under these circumstances as would be expected with normal feeding. Mice were lightly anaesthetised with isofluorane then orally gavaged with 0.2 mL of inoculum prepared at 5.25×10^9^ CFU/mL. Mice were then euthanised by CO_2_ asphyxiation and cervical dislocation at two time-points, either +3 or +28 days after *Salmonella* inoculation (*i.e.,* 16 weeks +3 days or 16 weeks +28 days). In this way, four treatment groups (n = 6 control diet, n = 12 control diet + *Salmonella*, n = 6 fgTiO_2_ diet, n = 12 fgTiO_2_ diet + *Salmonella*) were established, with half of the animals analysed at each of the two time-points. Alongside, faecal samples were collected at three time-points (pre-study, 16 weeks +3 days and 16 weeks +28 days) to enable enzyme-linked immunosorbent assay (ELISA) studies for *Salmonella*-specific IgA and faecal Ti measurements by ICP-MS.

### Human tissue collection

Following informed consent, and after approval from the Regional Ethical Review Board, Linköping, Sweden, surgical specimens containing Peyer’s patches were collected from the neo-terminal ileum from six patients with Crohn’s disease or colonic cancer (3 males, 3 females; resection date-range 2017-2018; median age 48 years; age-range 21 – 77 years). In all instances the ileal tissue specimens formed the resection margins and were macroscopically normal. Peyer’s patches were microscopically identified as previously described^48^. The tissue specimens were fixed with 4% PBS-buffered (pH 7.4) paraformaldehyde for 12 h at 4°C before immersion in 30% sucrose and freezing in OCT (LAMB/OCT, Thermo Scientific) in cryomolds. For analysis, the OCT-embedded tissue samples were rapidly shipped overnight to Cambridge on dry ice. Three anonymised, formalin-fixed paraffin embedded (FFPE) tissue samples containing ileal Peyer’s patch lymphoid tissue were also analysed to confirm the presence of fgTiO_2_ in the FFPE specimen-type. Studies using human tissues at Cambridge were also approved by the UK NHS Health Research Authority, North West – Greater Manchester East Research Ethics Committee, REC reference 18/NW/0690.

### Tissue sectioning and immunofluorescence labeling

Frozen sections were cut at 20-micron thickness using a Leica CM 1900 cryostat, picked up on Superfrost Plus slides (ThermoFisher, J1800AMNT) and rested at room temperature for 2 h prior to labelling. FFPE sections were cut at 5-micron thickness, then dewaxed and rehydrated by baking at 60°C for 1 h, transferring through two changes of xylene, a reverse ethanol series (100%, 70%, 50%, 10%) followed by 1 minute rehydration in water. Sections were ringed with hydrophobic barrier pen (Vector Laboratories, H-4000) and cryostat sections fixed by exposure to fresh 4%, 0.1M PBS-buffered paraformaldehyde (pH 7.4) for 10 min. All subsequent steps were conducted under gentle agitation on a rotating shaker. To facilitate antibody penetration, sections were permeabilised for 90 min using 0.3 % (vol/vol) Triton X-100 in 0.1 M PBS (pH 7.4). Block buffer (10% goat serum, 2% bovine serum albumin, 25 mM glycine diluted in 25 mM Tris-buffered (pH 7.4) saline (TBS)) was then added to all sections for 1 hour. Primary antibodies or concentration-matched isotype controls were diluted in block-buffer and incubated at 4°C overnight. All subsequent steps took place at room temperature. Sections were washed with three changes of block-buffer. Secondary antibodies (where directly-conjugated primary antibodies were not used) were diluted 1:400 in block buffer and incubated for 2 h. All antibody concentrations, fluorophore conjugations and manufacturer information are provided in **Supplementary Table 2**. 500 nM phalloidin-AlexaFluor 647 (ThermoFisher, A22287) and 2 µg/mL Hoechst 33342 (ThermoFisher, H3570) were also included alongside secondary antibodies to simultaneously label cell outlines (*i.e.,* cytoskeletal f-actin) and cell nuclei (respectively). Sections were then washed once with TBS prior to mounting with #1.5 coverslips in Prolong Diamond (ThermoFisher, P36965).

### Confocal microscopy & imaging controls

Fluorescence and reflectance image-data (2048×2048 pixels per tile) were collected using a diode-controlled (retrofitted) Zeiss LSM780 confocal microscope equipped with plan-apochromat 63X/1.4 and 40X/1.3 oil-immersion objectives. Fluorescence and reflectance data were collected at the same time during the same imaging run by sequential scanning with reflected (*i.e.,* back-scattered) light from a 488 nm excitation laser collected in the range 485 – 491 nm to detect fgTiO_2_^15^. Where tilescanning was used, images were acquired with 10% edge overlap. Tiles were stitched using the Zen Black 2011 SP2 software (Carl-Zeiss, UK) using the nuclei and actin channels for registration and a ‘strict’ correlation threshold setting of 1.0. For all datasets under comparison, images were obtained in a single run using identical settings. For the reflectance imaging, tissues from negative control mice (*i.e.,* fed the AIN-76 diet only, so no fgTiO_2_ present) were used to determine the image acquisition settings and thresholding for fgTiO_2_ detection (strategy shown, **Supplementary Figure 1**). For immunofluorescence work, supporting secondary-only and concentration-matched isotype antibody controls in tissue-matched serial sections were included. For datasets collected as sets of single images from the villous mucosa (VM), sub-epithelial dome (SED), germinal center (GC) and TCZ (T-cell zone) regions of mouse Peyer’s patches, landmarks (*e.g.,* villi / follicle-associated epithelium) and T cell (CD3), B cell (B220) and mononuclear phagocyte (CD11c) immunofluorescence staining in adjacent sections were used to inform imaging locations. When tilescanning was performed, images were collected with 10% edge-overlap using focus mapping to facilitate accurate image stitching whilst minimising acquisition time. To define the autofluorescence signature of the fgTiO_2_-recipient cells in the mouse Peyer’s patches, lambda scans were performed using the LSM 780 microscope. Using unstained tissue sections, autofluorescence upon excitation from a 405 nm laser was detected in ∼ 10 nm intervals in the range 415-700 nm (spectra presented, **Supplementary Figure 8**).

### Deconvolution

Z-stacked image-data (pixel size (XYZ) 1024×1024×123; voxel size (XYZ) 26×26×100 nm) of fgTiO_2_-recipient cells was collected using a Zeiss LSM 780 confocal microscope equipped with a 63X/1.4 plan-apochromat oil immersion objective using murine tissue sections embedded in Prolong Glass (P36984, ThermoFisher) anti-fade reagent (refractive index ∼ 1.5). Cell autofluorescence was excited using the 405 nm laser. The image-stacks were registered to correct spatial drift using StackReg^49^. The Born and Wolf scalar-based diffraction model^50^ was then used to estimate a theoretical point spread function assuming a refractive index of 1.51 and a autofluorescence emission maximum of 495 nm (measured, **Supplementary Figure 8**). The registered Z-stack was then deconvolved using the Richardson-Lucy total variation algorithm (75 iterations, regularisation 1e-12) within the open-source DeconvolutionLab2^51^ software.

### fgTiO_2_ detection using reflected light and circle-marker placement

To enable fgTiO_2_ detection in biological tissue sections, reflected light and immunofluorescence information was collected at the same time using a Zeiss LSM780 confocal microscope. The reflectance images were thresholded to isolate the punctate, reflectant foci caused by the fgTiO_2_ particles from the background light scatter caused by the biological tissue. As the background level was similar in both mouse and human specimens (shown, **Supplementary Figure 4**) the same threshold was used for both mouse and human images, and was determined as the level required to remove all signal in the reflectance channel for images collected for tissue sections taken from the negative control mice (*i.e.,* fed the AIN-76 diet only, so no fgTiO_2_ present). This approach was then further validated using correlative scanning electron microscopy and energy dispersive X-ray analyses to confirm that the signal remaining after thresholding contained the expected X-ray signal for titanium (strategy shown, **Supplementary Figure 1**). Given the small size of the reflectant foci caused by fgTiO_2_, a translucent circle-marker was placed on each pixel above the threshold using the ‘insertShape’ function in MATLAB R2021a (MathWorks) to aid visualisation given the relatively small size of display in the Figure panels^15^. Examples of the raw signal and subsequent marker placements are shown for mouse and humans in **Supplementary Figure 5**. Example data and code demonstrating thresholding and circle marker placement are available at the BioStudies database under accession number S-BSST875.

### Correlative confocal-electron microscopy and energy dispersive X-ray analyses

Once fgTiO_2_ detection by confocal reflectance microscopy was completed, Prolong-mounted coverslips were removed from the tissue sections by overnight incubation in PBS at 37°C. Slides were air dried at room temperature for 24 h, cut down and mounted on aluminum SEM stubs using colloidal graphite and coated with a thin (∼ 15 nm) layer of amorphous carbon (evaporative coater). Tissue sections were then loaded into a Helios G4 CX Dual Beam high-resolution, monochromated, field emission gun, scanning electron microscope (SEM) equipped with a precise focused ion beam (FIB) and in-lens secondary electron and circular backscattered detectors (FEI/ThermoFisher). Secondary and backscattered electron images were collected at 3 – 5 kV as tilescans for the complete tissue surface, permitting manual overlay onto the previously-collected confocal tilescans. To enable correlative confocal-SEM imaging, the ‘outline’ of the Peyer’s patch – particularly the follicle associated epithelium and basal muscularis layer – were manually registered together using image overlay, enabling triangulation to target specific regions containing reflectance-positive cells. The correlative confocal-to-EDX analyses of the human tissues were then carried out directly in the SEM, using a 150 mm^2^ silicon drift detector and AZTEC software (Oxford Instruments). For the correlative scanning transmission electron microscopy (STEM) used to confirm single-particle detection by reflectance microscopy, thin lamella of the tissue sample were prepared in the SEM via the *in situ* lift-out method. Once sites-of-interest had been identified, 500 nm of electron beam platinum (Pt) was deposited (at 5 kV, 6.4 nA for the electron source) to the surface of the target area. This was followed by a second Pt layer (1 ∝m) using the FIB (at 30 kV, 80 pA for the liquid gallium (Ga) ion source). A bulk lamella was initially cut using the FIB (at 30 kV, 9 nA), before the final cut-out was performed (at 30 kV, 80 pA). The lamella was attached, using ion beam Pt, to a copper FIB lift-out grid (Omniprobe, USA) mounted within the SEM chamber (*i.e., in situ*). Final thinning and polishing of the lamella to electron translucency was performed with a gentle polish/clean using the ion beam (5 kV, 41 pA). A lamella was then transferred into a Titan^3^ Themis G2 300 kV S/TEM equipped with an S-TWIN objective lens, monochromator, HAADF detector and a Super-X 4-detector EDX system in a double-tilt specimen holder (FEI/ThermoFisher). Transmission electron microscopy (TEM) images and on-zone diffraction patterns were recorded using a OneView CMOS camera (Gatan) and STEM-EDX elemental maps were recorded and processed using the Velox software (FEI/ThermoFisher).

### Cell segmentation of intestinal tissue images

In previous work, we carefully developed and validated staining and image analysis methods enabling extraction of single-cell data from confocal microscopy images of gastrointestinal tissues^37^. Based on this framework, three cell segmentation pipelines were used to accommodate different tissue types and differing availabilities of training data. To confirm the reliability of our previously-published pipelines with the new image-data presented here, the accuracy of the outputs from all three cell segmentation pipelines were assessed relative to manually-segmented cells by Jaccard index (presented, **Supplementary Figure 2)**.

### Cell segmentation: mouse villous mucosa images

Villous image-fields were directly segmented into cell-objects using the freely-available CellProfiler software^52^. Using the Hoechst 33342 fluorescence information, nuclei were first segmented as primary objects. These were then used as seeds during a marker-controlled watershed process to segment each cell’s outline using fluorescence information from the actin channel. The process is schematically demonstrated in **Supplementary Figure 18** with the complete CellProfiler pipeline shown in **Supplementary Note 1**. Test images and the CellProfiler pipeline are also available for download at the BioStudies database under accession number S-BSST875.

### Cell segmentation: mouse lymphoid tissue images

Lymphoid tissues (*e.g.,* Peyer’s patches and caecal patches) exhibit extremely high cell densities leading to poor cell segmentation accuracies using the simple, marker-controlled watershed approach used for the villous mucosa images^37^. Pixel-classification machine learning can be used to alleviate this problem through the generation of probability images that more precisely delineate each cell’s boundary^37^. Here, in mouse lymphoid tissues, these probability images were provided using a 2-D U-Net fully convolutional neural network^53^ deployed in MATLAB R2021a (MathWorks, MA). The network has a 256×256×2 (x, y, channels) input layer and uses the nuclear and actin fluorescence information to predict the probability that each pixel belongs to either ‘cell outline’, ‘intracellular environment’ or ‘background’ classifications. The training data consisted of twelve lymphoid tissues images (each containing 2138 x 1900 annotated pixels covering ∼16,000 cells prepared by an experienced cell biologist). The image-data flows through a four-layer contracting convolutional path before complete up-convolutional expansion, yielding probability images that exactly match the input-image dimensions. The network was trained for 50 epochs using a batch-size of 12 for 1,200 iterations/epoch. Patches were shuffled every epoch and augmented by simple, random x / y reflection and rotation. Model training was optimised under stochastic gradient descent using cross-entropy loss. The initial learning rate was 0.05, dropping every 10 epochs by 0.1 under momentum 0.9 and L2 regularisation 1×10^-4^. The resultant ‘Cell Outline’ probability images were then loaded into the CellProfiler pipeline alongside immunofluorescence and reflectance information collected by the confocal microscope and were used to enable segmentation of individual cells via an ‘IdentifyPrimaryObjects module’. This process is schematically demonstrated in **Supplementary Figure 19**. The complete CellProfiler image analysis pipeline is presented in **Supplementary Note 2**. All U-Net code, the trained network with test-data and the complete CellProfiler pipeline are available for download from the BioStudies database under accession number S-BSST875.

### Cell / TLV segmentation: correlative mouse-human dosimetry

For the correlative analyses investigating the relative amounts of fgTiO_2_ delivered to individual SED cells in mice versus in normally-exposed humans, frozen ileal tissue sections containing Peyer’s patches were collected at random from six normally-exposed humans, and six mice fed the AIN-67 diet supplemented with 0.0625% (w/w) fgTiO_2_. In every instance, care was taken to collect tissue sections with clear follicle-associated epithelium (*i.e.,* without any overlying villi) to ensure maintenance of similar histological positioning near the ‘center of the dome’ of each Peyer’s patch. Reflectance confocal images of the murine and human tissues were then acquired in a single run under identical settings. After acquisition, the image-fields per specimen were manually laid out in order of lowest-to-highest fgTiO_2_ accumulation to visually present the spectrum of (wide) variation in fgTiO_2_ cellular loading observed across samples in both the mouse and human tissue specimens. This was followed by quantitative image analysis to determine the thresholded reflectance per unit tissue area; then the integrated intensity of the threshold reflectance information per-cell and per-vesicle objects (method exemplified, **Supplementary Figure 3**). To extract the single-cell and single-vesicle data, a CellProfiler pipeline was used. To tackle cell segmentation in lymphoid tissues with high cell densities we have shown in previous work^37^ that pixel-classification machine learning can be used to produce probability images that more precisely delineate each cell’s boundary improving segmentation accuracies when compared to using the raw fluorescence information. Unlike in mouse lymphoid tissues (method described above), because large amounts of manually-annotated training data for the human lymphoid images was not available, these probability images were created by sparse annotation pixel-classification machine learning in the ILASTIK software^54^. Two ILASTIK pixel-classification projects were created (one for the human images, one for the mouse). Pixel annotations representing ‘intracellular environment’, ‘cell outlines’ and ‘other’ classifications were made on the fluorescence channel describing the cell outlines (*i.e.,* the actin channel) (method shown schematically, **Supplementary Figure 20**). The ILASTIK software then generated probability images representing the probability of each pixel belonging to each classification. The resultant ‘Cell Outline’ probability images were loaded into the CellProfiler pipeline alongside the fluorescence and reflectance information captured by the confocal microscope and were used to segment individual cells via an ‘IdentifyPrimaryObjects’ module. As part of this pipeline, the reflectant foci representing fgTiO_2_ events were also directly-segmented from the thresholded reflectance images into ‘fgTiO_2_-loaded vesicle’ (TLV) objects using a second, ‘IdentifyPrimaryObjects’ module (method schematically demonstrated, **Supplementary Figure 21**). The complete image analysis pipeline is presented in **Supplementary Note 3**. Test images and the complete CellProfiler pipeline are available for download from the BioStudies database under accession number S-BSST875. The accuracy of the segmented-cell outputs were checked by Jaccard index relative to manually-segmented cells (presented, **Supplementary Figure 2)**. The accuracy of TLV segmentation from the raw reflectance information is explored in both mouse and human images in **Supplementary Figure 5**.

### Cell / TLV feature extraction

As described in previous work^37^, raw immunofluorescence data were pre-processed by manual thresholding at the level required to remove ≥ 95% of fluorescence in tissue-matched, secondary antibody-only control images. Alongside, the reflectance data were carefully thresholded at the level required to remove all signal from the images collected from the negative control group mice (*e.g.,* fed the AIN-67 diet alone; strategy shown, **Supplementary Figure 1**). Thresholded reflectance and fluorescence intensity information per cell or TLV object, alongside size and shape features were then measured for both the immunofluorescence and reflectance channels using the ‘MeasureObjectSizeShape’ and ‘MeasureObjectIntensity’ modules in CellProfiler. The number of TLV objects per cell was measured using a ‘RelateObjects’ module. Data were preprocessed by discarding objects outside of the 5^th^ and 95^th^ percentiles by size prior to analysis as is recommended best-practice^55^.

### Cell segmentation: block processing tilescans

To avoid memory limitations, when necessary, tilescanned images were cut into smaller pieces, run through one of the CellProfiler pipelines before reassembling the object segmentation masks and assigning global cell position coordinates to the extracted object features. This was achieved using two MATLAB functions developed in our previous work^37^. The first ‘TilescanToCellProfiler’, reads stitched images in most microscopy formats and cuts them into manageable tiles with edge-overlap for processing. The second, ‘CellProfilerToTilescan’ reassembles the object masks, removes ‘double hits’ on overlap edges and assigns global position coordinates values to the extracted features. Example images and MATLAB code demonstrating these functions is available for download at the BioStudies database under accession number S-BSST875.

### Cell segmentation: accuracy assessment by Jaccard index

The Jaccard index (intersection over union) approach was used to check the accuracy of the cell segmentation results used in this work. This was achieved by comparing pixel positions inside the automatically-segmented cell objects against those obtained by manual annotations performed by an experience cell biologist. The Jaccard index was calculated as:

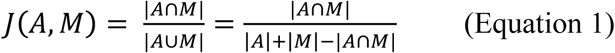

Where J is the Jaccard distance for two sets of pixel positions for the automated (A) and manual (M) segmentations, respectively. Scores of zero represent no overlap (false negatives) whereas scores of 1 represent exact pixel-for-pixel intersection. In this way, it is acknowledged that this approach is a relatively harsh success measure, and that scores of ∼ 0.7 indicate a good segmentation result^53^. This is partially due to the inevitable inaccuracies that are present – even in the manually-annotated data (*e.g.,* due to outline smoothing, ambiguity in determining the precise position of each cell’s boundary from the fluorescent staining information, and the available image resolution *etc*.).

### Cell segmentation: cell mapping visualisations

The cell map visualisations displaying the delivered cellular doses of fgTiO_2_ or TLV counts per-cell were produced using cell feature information extracted by the CellProfiler ‘MeasureObjectIntensities’ and ‘RelateObjects’ modules alongside the object segmentation masks outputted by each pipeline. Using MATLAB scripts, object feature data were binned into categories (*e.g.,* ‘low, ‘intermediate’ and ‘high’ fgTiO_2_-load based on the integrated intensity of the thresholded reflectance signal per-cell). Using the segmentation masks, individual cell-objects were then colour-coded according to these categories to provide a ‘tissue map’ visualisation. Example data and MATLAB code demonstrating the creation of these visualisations are available for download from the BioStudies database under accession number S-BSST875.

### Statistical analysis of image-based data

Integrated intensity per object (*i.e.,* per cell or NLV) distributions were compared using two-sided Wilcoxon rank-sum analysis. This non-parametric approach was chosen for compatibility with unpaired samples with unequal group sizes^37^. The approach tests the null hypothesis that the distributions under comparison might reasonably be drawn from one continuous distribution with equal medians (P > 0.05) versus the alternate hypothesis that the distributions are distinct (P ≤ 0.05). Pairwise comparisons of cell counts were first checked for distribution normality and variance homogeneity using Shapiro-Wilk and Barlett tests, respectively. Where these tests were passed (P >0.05), groups were compared using a two-sided, unpaired samples T-test assuming equal variance.

### Olink^®^ proximity extension assay

Frozen mouse ileal samples were weighed and stored in Kimble pellet pestle tubes (DWK Life Sciences, K-749520-0000) at -70°C until processing. The Peyer’s patch-enriched samples were first trimmed tightly to either side of Peyer’s patch using a cold safety razorblade such that each sample primarily consisted of only the patch itself. Tissues were lysed using the BioPlex Cell Lysis Kit (BioRad, 171-304011). 500 µL of Cell Wash Buffer per sample was used for rinsing the tissues. Cell Lysis Buffer was prepared by adding Factors 1 and 2 as per the manufacturer’s instructions, as well as Halt Protease Inhibitor Cocktail (ThermoFisher, 78437; 10 µL of cocktail per 1 mL of buffer). Following a rinse, tissues were homogenised by adding 500 µL of ice-cold Cell Lysis Buffer and hand grinding on ice using disposable pestles (DWK Life Sciences, 749520-0000; 20 strokes).

Homogenised samples were frozen at -70°C for at least 4 hours, thawed on ice, sonicated in an Ultrawave U300 ultrasonic bath (44 kHz, 35 W) on ice for 40 seconds and centrifuged for 5 min at 6,000 x g. Supernatant was collected into 2 mL cryogenic vials (Corning, 430659) and frozen at - 70°C until quantification. Quantification was performed using the DC Protein Quantification Kit I (BioRad, 5000111) as per the manufacturer’s instructions (Microplate Assay Protocol). Plates (Greiner Bio-One, 650185) were read using FLUOstar Omega microplate reader at 750 nm and protein concentrations were calculated by the Omega’s MARS Data Analysis software using linear regression fit. All samples were normalised to 0.77 mg/mL (+/- 0.12) by dilution with Cell Lysis Buffer and re-quantified to confirm protein concentration. A microplate was prepared for the proximity extension assay by transferring 40 µL aliquots of all samples, negative controls (Cell Lysis Buffer), duplicates of 6 samples, and dilutions (1:4, 1:8 and 1:16 v/v) of two samples into a 96-well skirted plate (ThermoFisher, AB0800) in a randomised manner (controls presented, **Supplementary Figure 11**). The plate was sealed (ThermoFisher, 4306311) and stored at -70°C until shipment on dry ice to Olink (Uppsala, Sweden) for analysis using the Target 96 Mouse Exploratory panel (www.olink.com/mouse-exploratory). All protein names and abbreviations are listed in **Supplementary Table 1**. Statistical analysis was performed using the Olink Analyze 3.6.0 R-package (https://CRAN.R-project.org/package=OlinkAnalyze). Treatment groups were compared using a two-sample Mann-Whitney U test with correction for multiple testing using the Benjamini-Hochberg method. In the main text, results for proteins that could be involved in immuno-inflammatory signaling are shown (**Figure 5**) whereas in the Supplementary Information results of the entire protein set are shown alongside repeat-sample, serial dilution and blank sample proximity extension assay controls (**Supplementary Figures 11-13**). The proximity extension assay data are available for download at the BioStudies database under accession number S-BSST875.

### Enzyme-linked immunosorbent assay for *Salmonella*-specific IgA

During Mouse Study 2, faecal pellets were collected pre-study and at the 16-week +3 day and 16-week +28 day time-points. Samples were weighed and diluted 1:10 (w/v) in 10% PBS containing 1 mM EDTA. The resulting suspension was vortexed for 15 min to facilitate liquefaction then centrifuged at 400 x g at 4°C for 5 min. The supernatant was aspirated and centrifuged for a second time at 12,000 x g for 10 min at 4°C. *Salmonella*-specific IgA quantification was then carried by sandwich ELISA method. Plates were coated with 50 µL lipopolysaccharide (LPS), (Sigma, L2262) diluted in Reggiardo’s buffer (0.05 M glycine, 0.1 M NaCl, 1 mM EDTA, 0.05 M NaF and 0.1% sodium deoxycholate). Plates were then sealed and incubated overnight at 37°C. Next day, the plates were washed 3 times with PBS-tween and blotted dry. Blocking buffer from the whole IgA kit (Affymetrix, 88-50450) was added to each well for 1 hr at 37°C. The faecal supernatant was then diluted in kit buffer A (Affymetrix, 88-50450) and 50 µL of diluted sample was added to each well. After three hours of incubation, the plates were washed three times with PBS-tween before addition of 50 µL horseradish peroxidase-conjugated IgA. Three more washes were carried out before the addition of 50 µL tetramethylbenzidine substrate for 10 min. This was followed by 50 µL of stop solution prior to absorption reading at 492 nm on a plate reader. All outputs were calibrated using a standard curve and corrected for dilution.

### Faecal ICP-MS analysis

Faecal samples were collected and stored at -80°C. Samples were weighed and a digestion solution of 1:1 (vol/vol) nitric acid (Fisher, 7697-37-2) and hydrogen peroxide (Sigma, 7722-84-1) was added to the samples (20-70 mg faecal weight) at a 1:10 (wt/vol) ratio. Samples were digested at room temperature for 72 h before loosely-capped sample containers were transferred to a water bath (40°C) for 5-6 h to release the peroxide. Samples were then transferred to a PTFE vial and sulphuric acid was added (1:1 vol acid/sample weight) prior to final digestion using a Milestone UltraWave microwave. The resultant liquid was decanted and diluted 1:10 (vol/vol) using ultra-high purity water. Quantification of elemental Ti was performed using an 8900 triple quadruple inductively-coupled plasma mass spectrometer (Agilent, USA). Scans were performed in MS/MS mode with the reaction cell adjusted for analysis in H_2_/O_2_ gas. The instrument was set up to identify the mass pairs 48 and 64 for isotopic Ti-48 and TiO_2_-64. Calibration standards were prepared in 2% nitric acid and spiked with elemental Ti (Sigma, 12237) to final concentrations between 0 and 1 ppm. All samples were spiked with an internal standard solution containing gallium (Sigma, 16639), germanium (Sigma, 05419), yttrium (Sigma, 01357) & europium (Sigma, 05779) at final concentrations of 2 ppb. Counts per-second (cps) values for yttrium were collected to reference signal stability. A 2% solution of nitric acid was used as sample carrier and rinse.

## ONLINE CONTENT

Data and code are available at the BioStudies database (https://www.ebi.ac.uk/biostudies/) under accession number S-BSST875.

## DATA AVAILABILITY

Data supporting the presented analyses are available for download from the BioStudies database (https://www.ebi.ac.uk/biostudies/) under accession number S-BSST875. Source data are provided with this submission.

## CODE AVAILABILITY

MATLAB code (using Deep Learning, Image Processing and Computer Vision toolboxes) and CellProfiler pipelines enabling the presented image analysis workflows are downloadable from the BioStudies database (http://www.ebi.ac.uk/biostudies) under accession number S-BSST875.

## ACKNOWLEDGEMENTS

The authors acknowledge the UK Medical Research Council (grant number MR/R005699/1), the UK Engineering and Physical Sciences Research Council (grant EP/N013506/1) and the UK Biotechnology and Biological Sciences Research Council (grant number BB/P026818/1) for supporting the work. J.W.W. is grateful to Girton College and the University of Cambridge Herchel-Smith Fund for supporting him with Fellowships. J.W.W. thanks Debbie Edwards for her assistance with the microscopy studies and Guy Dew for his comments on the manuscript. The authors thank Prof. Neil Mabbot and Prof. Simon Milling for their helpful comments. J.D.S. and Å.V.K. were supported by Swedish Research Council Grant No. 2021-02566. Mouse Study 1 was funded by the Riddet Institute through its Centre of Research Excellence funding which was awarded by the New Zealand government. Additional funding was provided by the Ministry for Science and Innovation contract C11 × 1009 through Nutrigenomics New Zealand, a collaboration between AgResearch, Plant and Food Research, and The University of Auckland. S.R. was supported by doctoral scholarships from Massey University. The authors are grateful to Dr. Andrew Grant for his input into the study design for Mouse Study 2.

## AUTHOR CONTRIBUTIONS

JWW and JJP conceived the concept and designed the analyses. NR obtained funding for SR Ph.D. studies including Mouse Study 1. SR, NR and DO carried out Mouse Study 1. RJ, ABDS, MM, CB, JW, PM and JJP carried out Mouse Study 2. AVK and JDS collected the human surgical specimens. JWW, JR, MM, AF and REH conducted the light microscopical studies. AD, MM, JWW and JJP performed the protein expression analyses. APB, SM and JWW carried out the electron microanalytical work. JWW, PR, HDS and JJP performed the data analysis. JWW and JJP wrote the manuscript in close collaboration with all authors.

