## Supplementary Information for "Immunocompetent Cell Targeting by Food-Additive Titanium Dioxide"

**List of Supplementary Information –**

**Supplementary Figure 1** – Quantifying fg$TiO_2$ in biological tissues using confocal reflectance microscopy. This explains the methodology used to enable robust quantification of fg$TiO_2$ in mouse and human tissues.

**Supplementary Figure 2** – Assessing cell segmentation accuracies. This presents Jaccard index results for each of the cell segmentation strategies used in the study relative to results obtained by hand-drawing.

**Supplementary Figure 3** – Image-analysis strategies enabling fg$TiO_2$ quantification. This explains the different image analysis approaches used to measure fg$TiO_2$ at the tissue-region, cell and fg$TiO_2$-loaded vesicle levels.

**Supplementary Figure 4** – Background reflectance comparison in human and mouse tissue sections. This demonstrates that the background reflectance signal is highly similar in both mouse and human Peyer's patch tissue enabling reflectant foci to be established using a single threshold.

**Supplementary Figure 5** – Circle-marker placement and TLV segmentation examples. This shows examples of the raw reflectance signal and resulting circle-marker and fg$TiO_2$-loaded vesicle segmentations after image processing for a series of mouse and human examples.

**Supplementary Figure 6** – Statistical comparisons of the delivered cellular dose of fg$TiO_2$ in mice and humans. Statistical comparisons are presented for the measured dose of fg$TiO_2$ delivered to subepithelial dome cells at both the cell and endosome level.

**Supplementary Figure 7** – fg$TiO_2$ dose delivery using different diets. Further confocal reflectance microscopy data is presented comparing fg$TiO_2$ loading into mouse SED cells using diets supplemented with 0.000625, 0.00625 or 0.0625% (w/w) fg$TiO_2$.

**Supplementary Figure 8** – Autofluorescence properties of LysoMac/LysoDC cells. This Figure presents measured emission spectra for the LysoMac/LysoDC autofluorescence signal.

**Supplementary Figure 9** – Association of fg$TiO_2$ with LysoMac/LysoDC cells. This demonstrates the selectivity and specificity of fg$TiO_2$ for the autofluorescent, LysoMac/LysoDC cell population of the murine subepithelial dome.

**Supplementary Figure 10** – Immunofluorescence controls for PD-L1. This provides secondary-only and isotype control image-data in support of the PD-L1 immunofluorescence studies.

**Supplementary Figure 11** – Olink proximity extension assay controls. This shows the results of the random duplication, sample dilution and blank-buffer controls analysed by proximity extension assay.

**Supplementary Figure 12 –** Full protein expression analyses of ileal tissue digests by proximity extension assay showing all targets covered by the Olink mouse exploratory panel.

**Supplementary Figure 13 –** Protein expression analyses of Peyer's patch-enriched tissue digests by proximity extension assay showing all targets covered by the Olink mouse exploratory panel.

**Supplementary Figure 14 –** Confocal reflectance tilescans showing autofluorescence, CD4 and MHCII expression in fg$TiO_2$-recipient cells of the Peyer's patch at the +28 day timepoint.

**Supplementary Figure 15 –** Single-cell image analysis showing the ileal tissue distribution of fg$TiO_2$ following *Salmonella* challenge.

**Supplementary Figure 16** – Comparison of fg$TiO_2$ loading into different ileal tissue compartments. This presents a quantitative comparison of the relative amounts of fg$TiO_2$ delivered to different ileal tissue regions in the murine model.

**Supplementary Figure 17 –** Faecal titanium analyses by ICP-MS: This indicates the amount of fg$TiO_2$ passed in the faeces at the 16 week, 16 week +3-day and 16 week +28-day timepoints.

**Supplementary Figure 18** – Mouse villous mucosa cell segmentation. Schematic explanation of the marker-controlled watershed approach used to enable cell segmentation of mouse villous mucosa images.

**Supplementary Figure 19** – Cell segmentation of mouse lymphoid tissues using UNET pixel classification. This schematically explains how cell segmentations were achieved in mouse lymphoid tissues using a UNET pixel-classification model.

**Supplementary Figure 20** – Mouse-human correlative dosimetry: Peyer's patch cell segmentation. This schematically demonstrates the process used to enable cell segmentation of mouse and human Peyer's patch tissue using sparse-annotation pixel classification machine learning in the Ilastik software.

**Supplementary Figure 21** – Mouse-human correlative dosimetry: fg$TiO_2$-loaded vesicle segmentation. This schematically demonstrates the TLV segmentation approach from the reflectance image data.

**Supplementary Table 1 –** Olink proximity extension assay full protein names. This Table summarises the full list of proteins assessed by the Olink proximity extension assay alongside their full and abbreviated names.

**Supplementary Table 2** – Antibody information Table. This provides the specifics of all antibodies used in the study, including manufacturer information, staining concentrations and the secondary antibodies and fluorophores used for detection.

**Supplementary Video 1** – fg$TiO_2$ targets autofluorescent LysoMac/LysoDC cells of the murine subepithelial dome. This video presents a 3-D render of the murine subepithelial dome demonstrating the selectivity and specificity of fg$TiO_2$ targeting for the autofluorescent LysoMac/LysoDC cell population.

**Supplementary Note 1** – Image analysis pipeline enabling cell segmentation of mouse villous mucosa image fields. This section presents the CellProfiler pipeline used to segment and extract cell features from the mouse villous mucosa images. The CellProfiler project and sample image data are available for download from the BioStudies archive accompanying this paper.

**Supplementary Note 2 –** Image analysis pipeline for fg$TiO_2$ and immunofluorescence marker quantification in mouse lymphoid tissues. This section presents the CellProfiler pipeline used to measure immunofluorescence marker and fg$TiO_2$ features in segmented cells. The CellProfiler project and sample image-data are available for download from the BioStudies archive accompanying this paper.

**Supplementary Note 3 –** Image analysis pipeline enabling the correlative mouse-human dosimetry studies. This section presents the CellProfiler pipeline used to measure cell and fg$TiO_2$ loaded vesicle (TLV) properties in the mouse and human tissue sections. The CellProfiler project and sample image data are available for download from the BioStudies archive accompanying this paper.

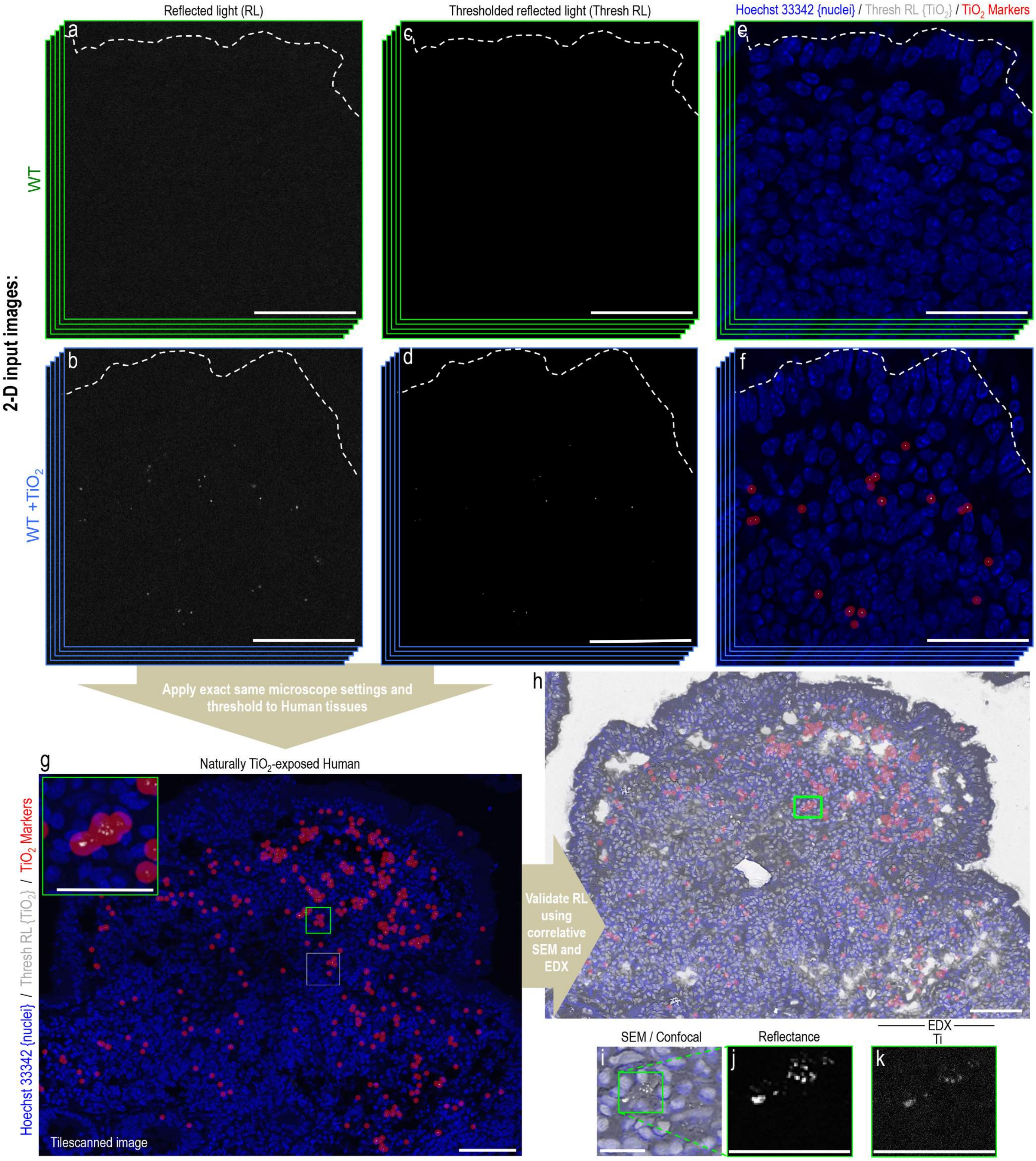


**Supplementary Figure 1 – Detecting and quantifying fg$TiO_2$ in biological tissues using reflectance confocal microscopy. a/b**, Using murine Peyer's patch tissue sections, known to be positive or negative for fg$TiO_2$ (*i.e.,* fed a diet with fg$TiO_2$ (WT +$TiO_2$) or without (WT) fg$TiO_2$), laser excitation and detector gain settings were optimised such that the signal from the brightest reflectant foci caused by fg$TiO_2$ approached saturation at the detector. At the same time, the confocal pinhole was used to obtain an optical section of ~1 micron in the Z-dimension enabling robust data collection from the centre of 20 micron-thickness tissue sections avoiding any surface-based signal. These microscope settings were universally applied during data acquisition using a Zeiss LSM780 laser scanning confocal microscope diode-retrofitted to maintain consistent laser excitation power. **c/d**, During the image analysis process, the murine tissues from the control-diet animals (*i.e.,* without fg$TiO_2$ supplementation) were used to inform a threshold for the reflectance channel ensuring that only the (**d**) signal from the highly reflectant foci caused by fg$TiO_2$ remained. **e/f**, To enable visualisation of these small reflectant foci, a translucent red circle marker was placed on each pixel above the threshold during the image analysis process. To orientate the images, dashed lines indicate the follicle-associated epithelium overlying the Peyer's patches. **g/h**, Once microscopy settings and image analysis thresholds were established using the murine tissues, the exact same settings were taken forward to enable the detection of fg$TiO_2$ in human tissues. **i-k**, To further validate the method in human tissues, correlative SEM-EDX analyses were carried out in the same tissue sections to confirm that the (**j**) reflectant foci detected by reflectance microscopy were an exact match for the (**k**) physical X-ray signal for Ti (further data shown, **Figure 1**). *Scale bars:* ***a-h*** *= 50 µm;* ***h****-inset and* ***i-k*** *= 10 µm.*

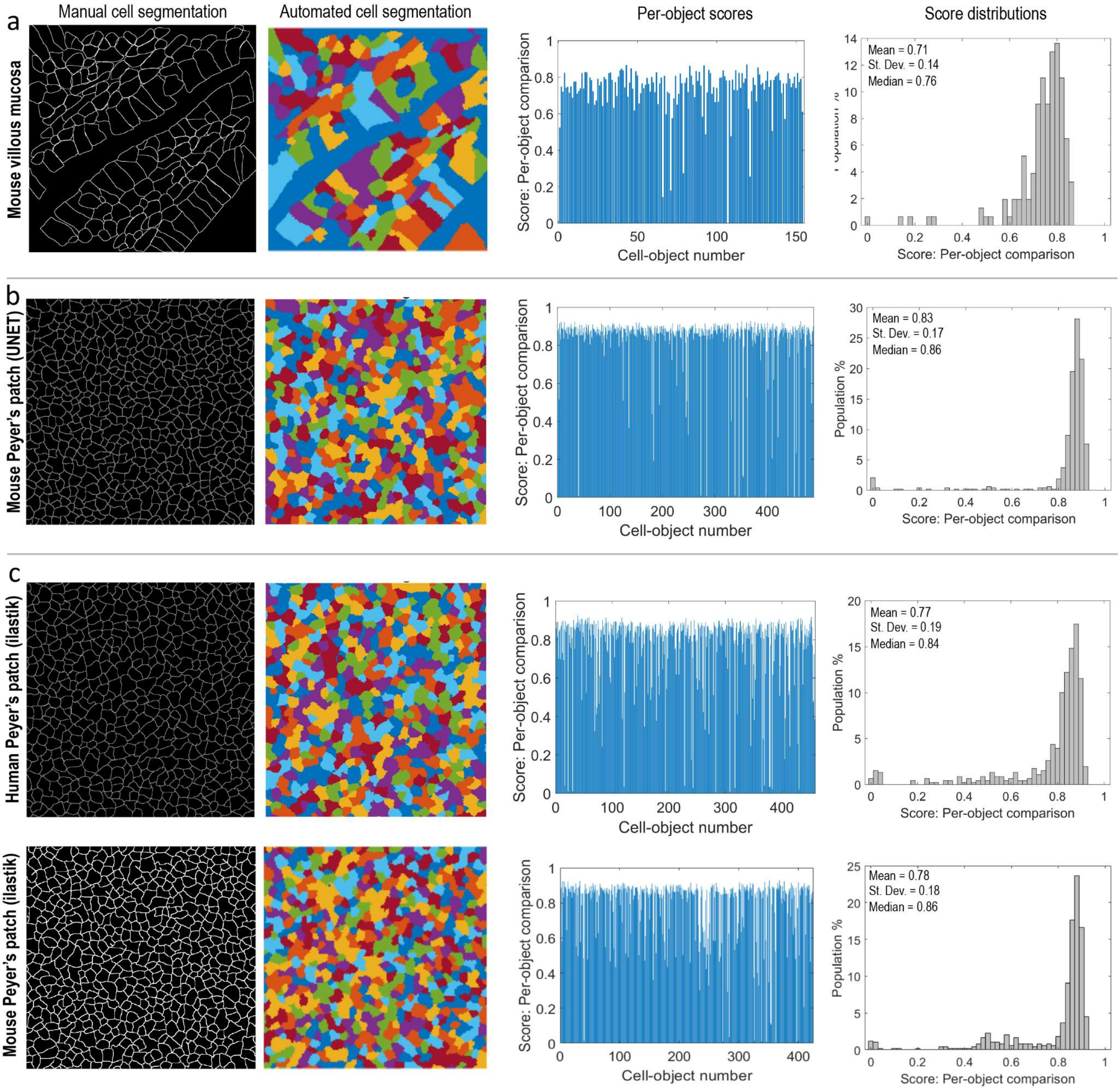


**Supplementary Figure 2 – Testing the accuracy of the automated cell segmentations against manually-drawn examples. a-c**, To check the accuracy of the three cell segmentation pipelines used, hand-drawn cell segmentations were performed by an experienced biologist. The results from the automated procedures were then compared – cell-object-by-cell-object – against the manually-drawn examples using the commonly employed intersection-over-union approach (Jaccard index). Per-object scores and distributions are shown for each analysis. A score of 1 represents a perfect pixel-area overlap between manual and automated cell segmentations. Due to the inherent inaccuracies present (*e.g.,* due to line thickness and outline smoothing *etc.,*) scores ≥ 0.7 typically represent a good segmentation result.

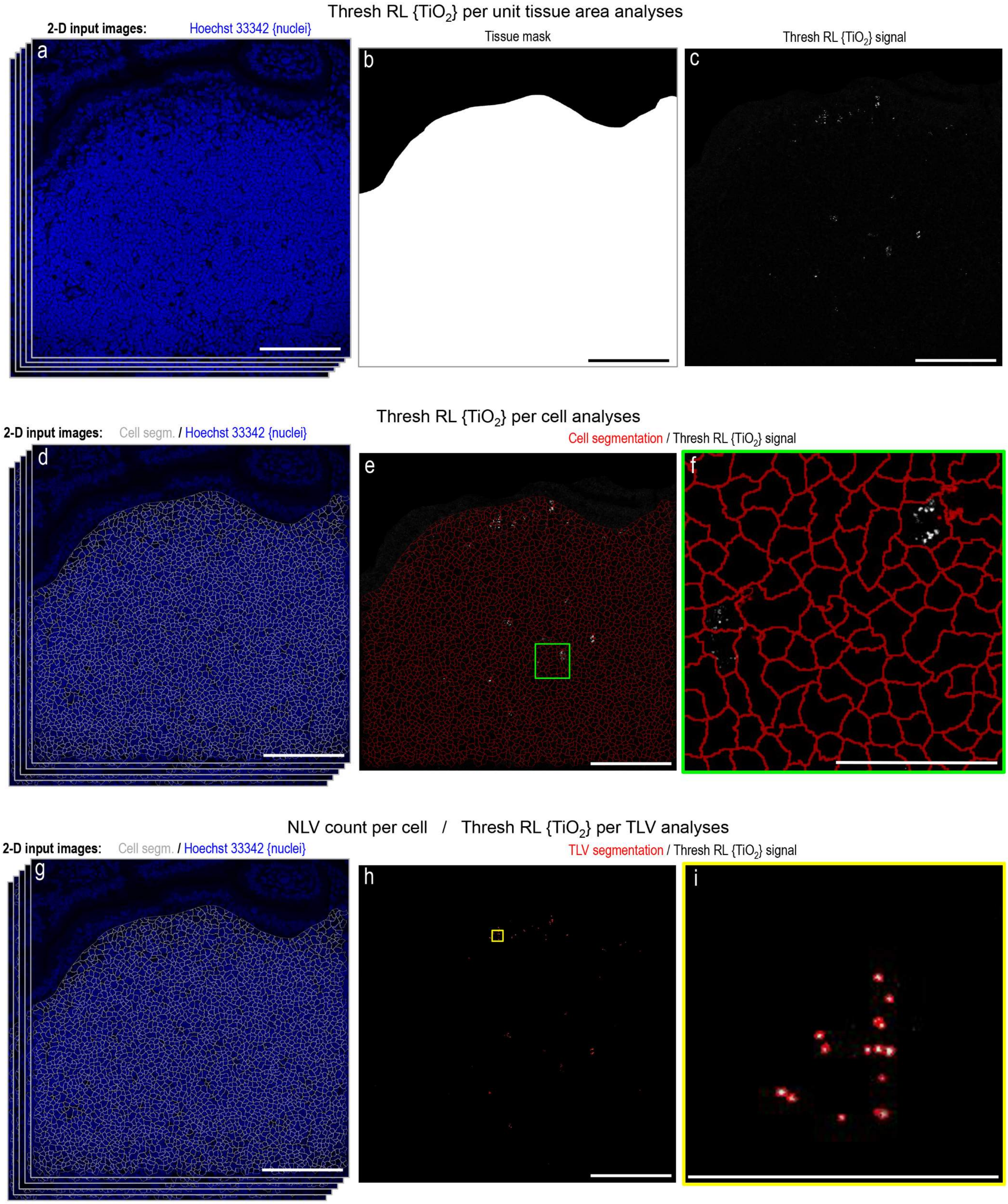


**Supplementary Figure 3 – Image analysis of fg$TiO_2$ at the tissue-region, single-cell and fg$TiO_2$-loaded vesicle levels. a-c**, To determine the abundance of fg$TiO_2$ per unit tissue area, masks of the tissue region-of-interest were (**b**) drawn manually. **c**, The thresholded reflectance signal attributable to fg$TiO_2$ (explained, **Supplementary Figure 1**) was then measured within these masked areas. **d-f**, The dose of fg$TiO_2$ delivered to individual cells was obtained by first segmenting all cells in the tissue region-of-interest and then (**f**) integrating the thresholded reflectance signal in each cell-object. **g-i**, Similarly, the count of fg$TiO_2$-loaded vesicles (TLV) per cell and the dose of fg$TiO_2$ per fg$TiO_2$-loaded vesicle were obtained by segmenting each individual, reflectant foci and counting these per (**f**) cell-object, or, by integrating the thresholded reflectance signal in each segmented TLV-object. *Scale bars:* ***a-h*** *= 50 µm,* ***i*** *= 10 µm.*

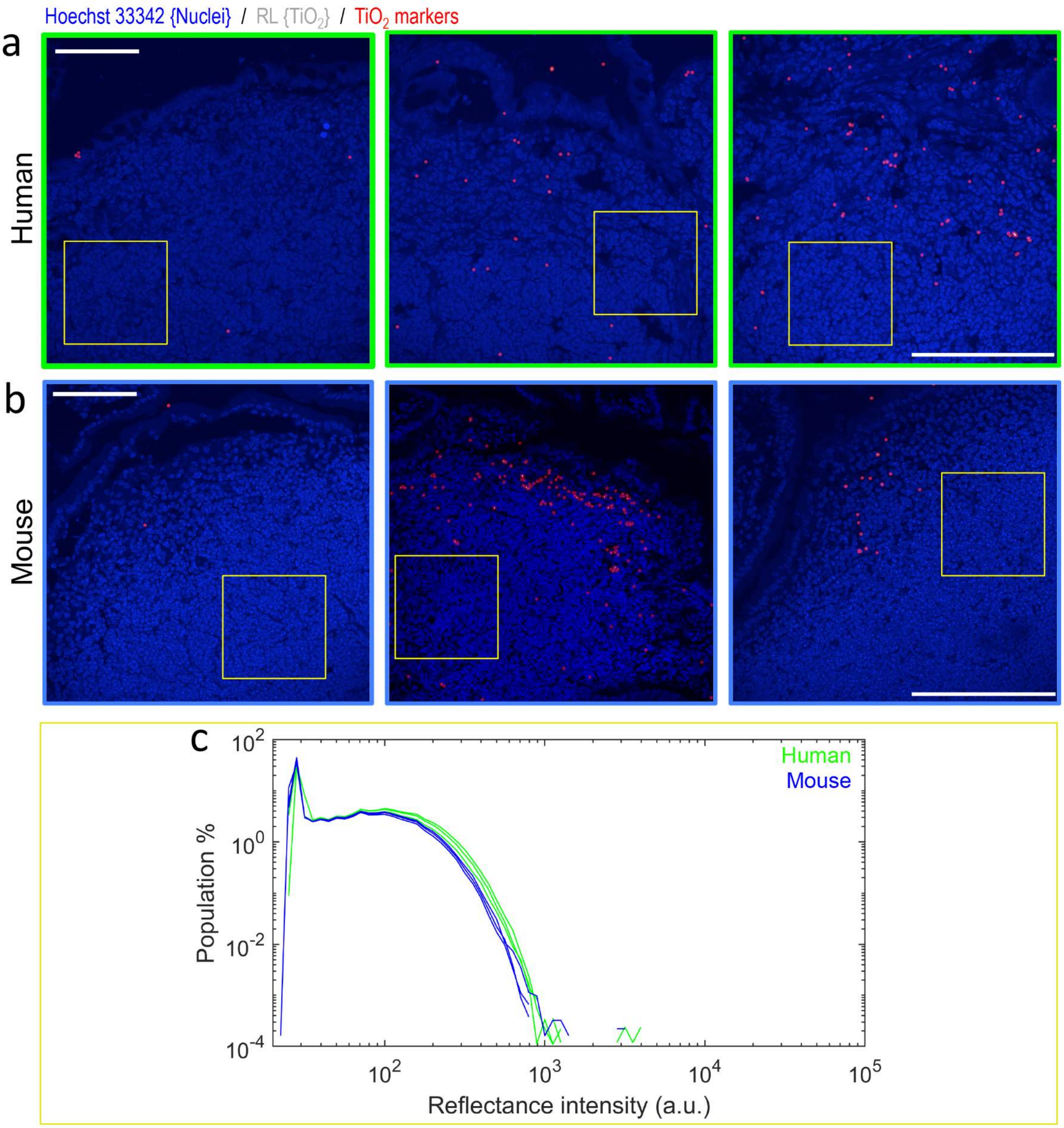


**Supplementary Figure 4 – Comparison of the background reflectance intensities in human and mouse tissue sections**. **a**/**b**, Image-regions containing subepithelial dome tissue but no reflectant foci were identified (yellow boxes) in (**a**) human or (**b**) mouse Peyer's patch datasets. **c**, The reflectance intensity information (per pixel) in these image-regions was overlaid as normalised histograms. The distributions are extremely similar: the optical sectioning imposed by the confocal microscopy technique means that the reflectance data is collected in a Z-depth of about 1 micron from the center of each 20-micron tissue section. The reflectance signal in this volume is largely driven by the cellular make-up of the tissue. In both mouse and humans, this tissue region predominantly consists of B-lymphocytes likely explaining the similar distributions observed. *Scale bars = 250 μm.*

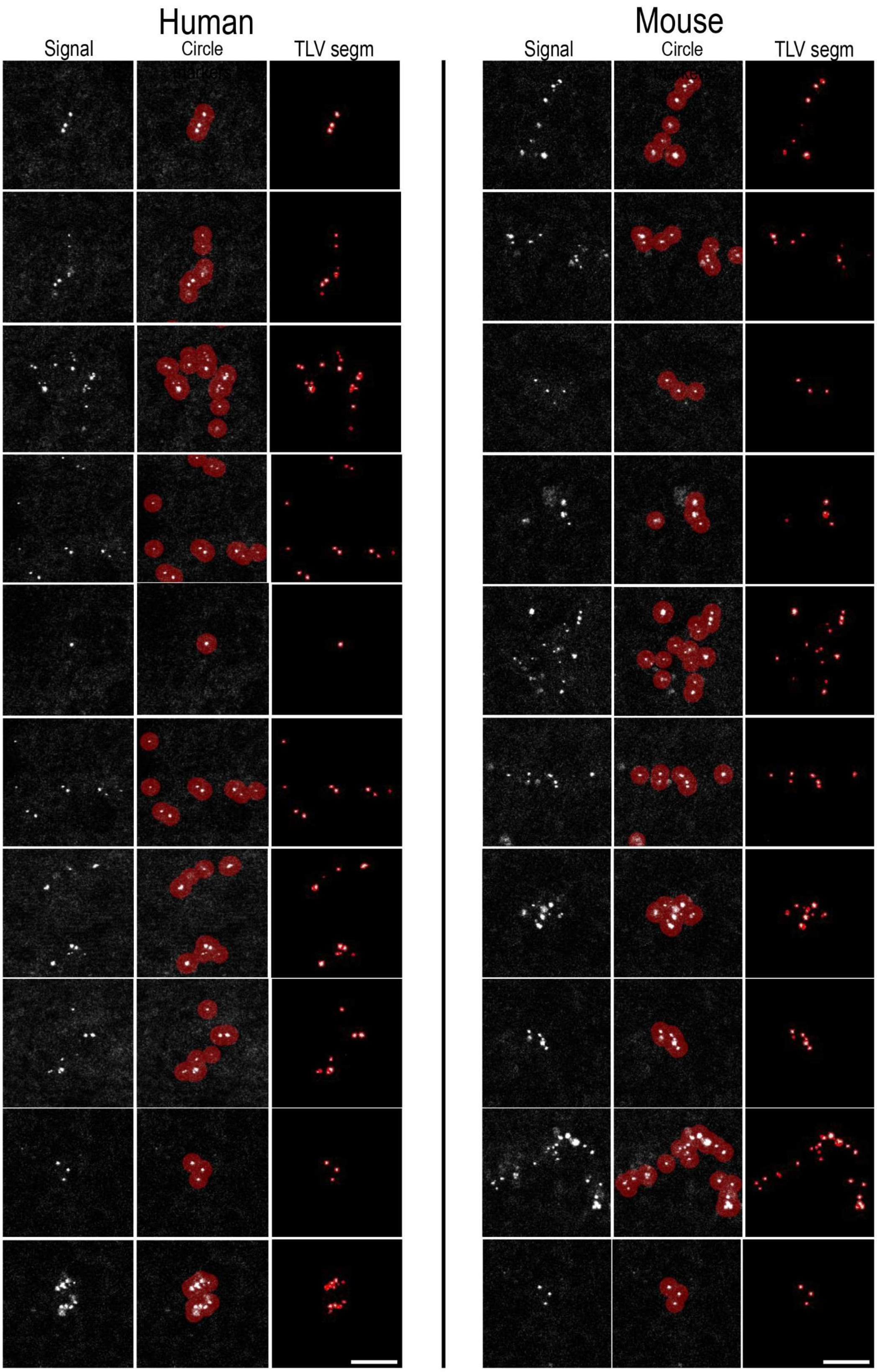


**Supplementary Figure 5 – Examples of circle-marker placement and fg$TiO_2$-loaded vesicle (TLV) segmentation using mouse or human reflectance data.** In the Left or Right panels, the first column exemplifies the raw reflectance signal collected by the confocal microscope. The second column shows the resultant circle-marker placements on thresholded reflectant foci (*i.e.,* representing fg$TiO_2$ events). The third column exemplifies reflectant foci segmentations into 'fg$TiO_2$-loaded vesicle' (TLV) objects. *Scale bars = 5 µm.*

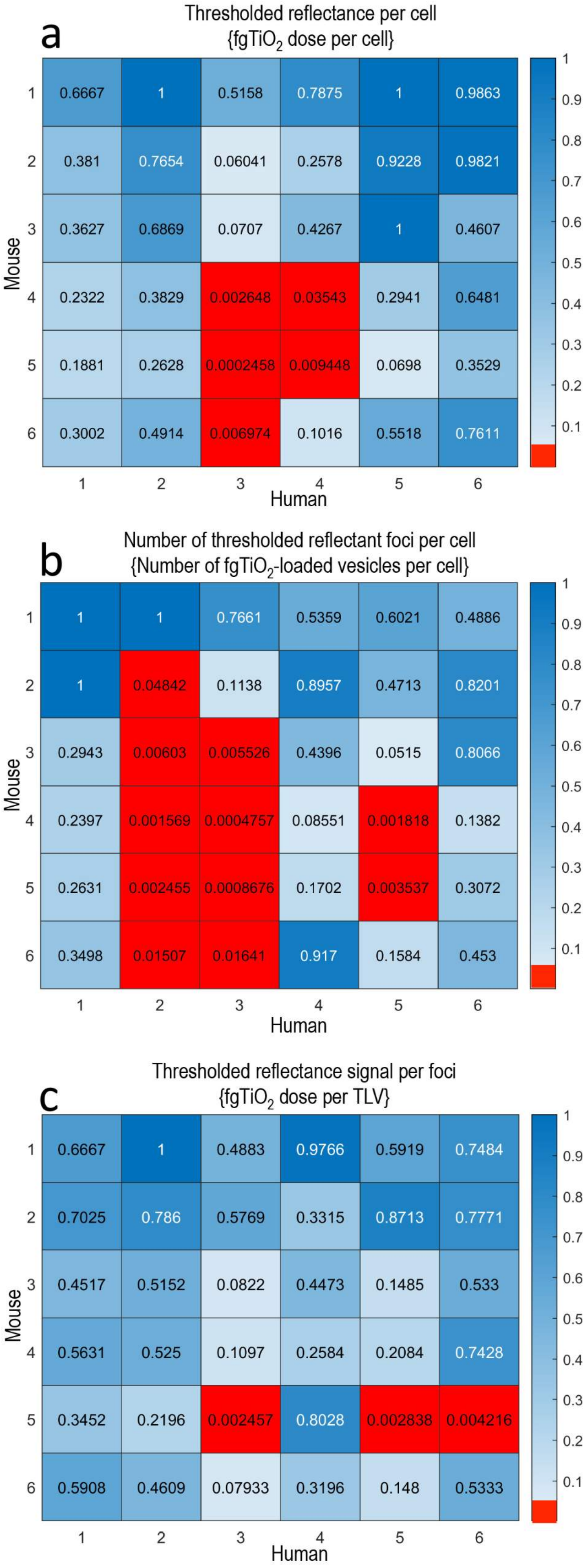


**Supplementary Figure 6 - Statistical comparison of fgTiO$_2$ loading into Peyer's patch subepithelial dome cells in mice and humans.** In each analysis, each of the six mouse and six human distributions presented in **Figure 3** were statistically compared by two-sided Wilcoxon rank-sum analysis. This tests the null hypothesis that each pairwise comparison could reasonably have been drawn from continuous distributions with equal medians ($P > 0.05$) against the alternate hypothesis that the distributions are distinct ($P < 0.05$). **a**, Using the measure of thresholded reflectance per cell (*i.e.* fgTiO$_2$ dose per cell) 31 out of 36 of the mouse-human comparisons reject the alternate hypothesis and thus might reasonably have been drawn from the same distributions. **b**, By the measure of the count of thresholded reflectant foci per cell (*i.e.,* the number of fgTiO$_2$-loaded vesicles (TLV) per cell) 25 out of 36 of the mouse-human comparisons were determined similar. **c**, Using the measure of thresholded reflectance signal per foci (*i.e.,* fgTiO$_2$ dose per TLV) 33 out of 36 of the mouse-human distributions were statistically similar. By each measure of the delivered cellular dose, feeding mice a diet supplemented with 0.0625% fgTiO$_2$ for eighteen weeks is seen to provide significant overlap with measured, real-world human exposures to fgTiO$_2$ in this tissue region.

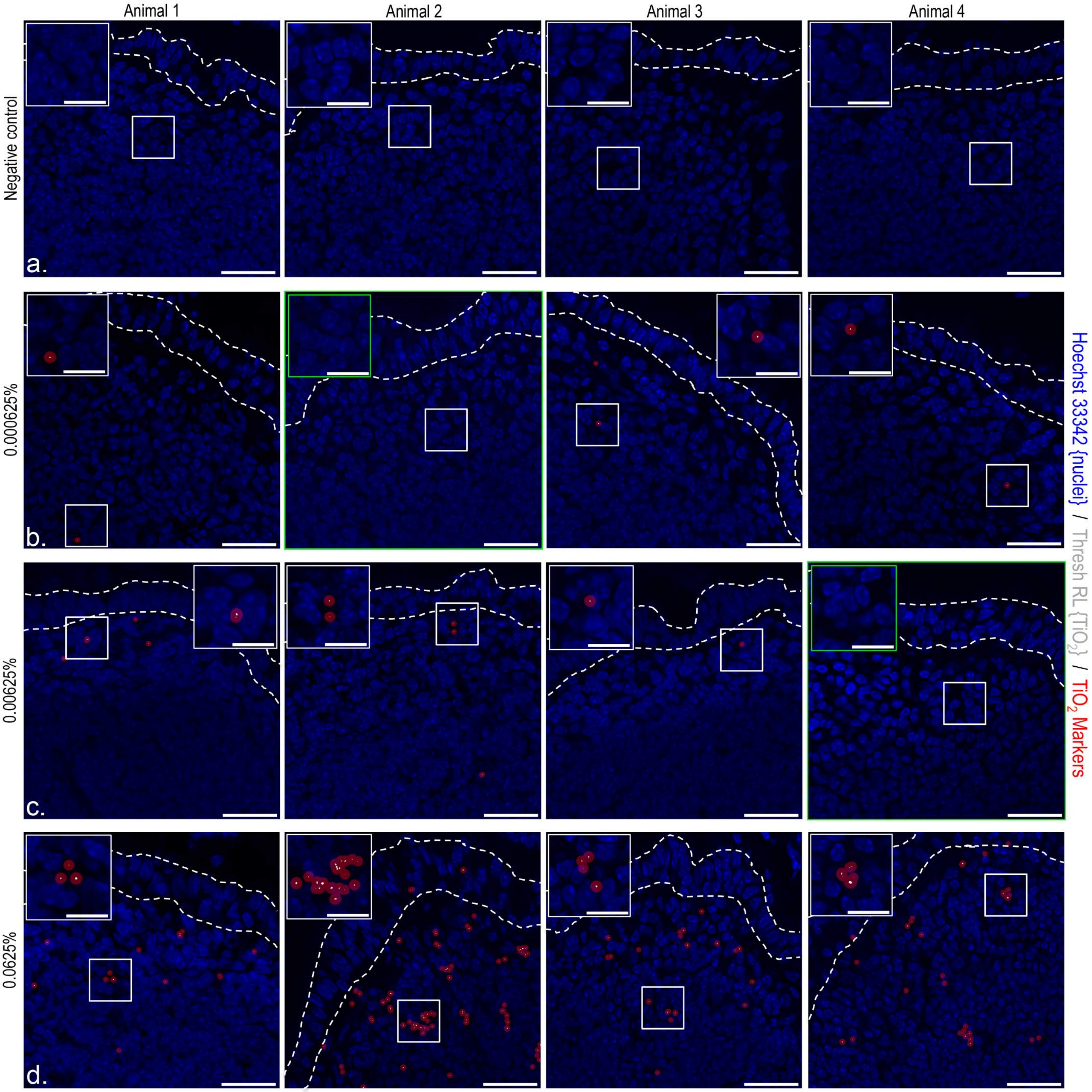


**Supplementary Figure 7 – Comparison of fg$TiO_2$ delivery into the murine Peyer's patch subepithelial dome using diets supplemented with differing levels of fg$TiO_2$.** American Institute of Nutrition (AIN)-76A diets containing 0.000625%, 0.00625% or 0.0625% fg$TiO_2$ (approximately 1.0, 10.0 or 100 mg fg$TiO_2$ / kg body-weight / day respectively) were prepared and fed alongside a negative control diet (AIN-76A alone) for eighteen weeks. **a-d**, After necropsy, the confocal reflectance microscopy approach (explained, **Supplementary Figure 1**) was able to detect delivery of fg$TiO_2$ into the SEDs for all three fg$TiO_2$-supplemented diets. Image analysis revealed that only the highest diet (0.0625% or 100 mg fg$TiO_2$ / kg body-weight / day) was able to recreate a similar cellular loading in the murine Peyer's patches to that observed in humans (presented, **Figure 3** with statistical comparison in **Supplementary Figure 6**). Dashed lines indicate the follicle-associated epithelium overlying the subepithelial dome tissue-region. *Scale bars: **a-d** = 25 µm; insets = 10 µm.*

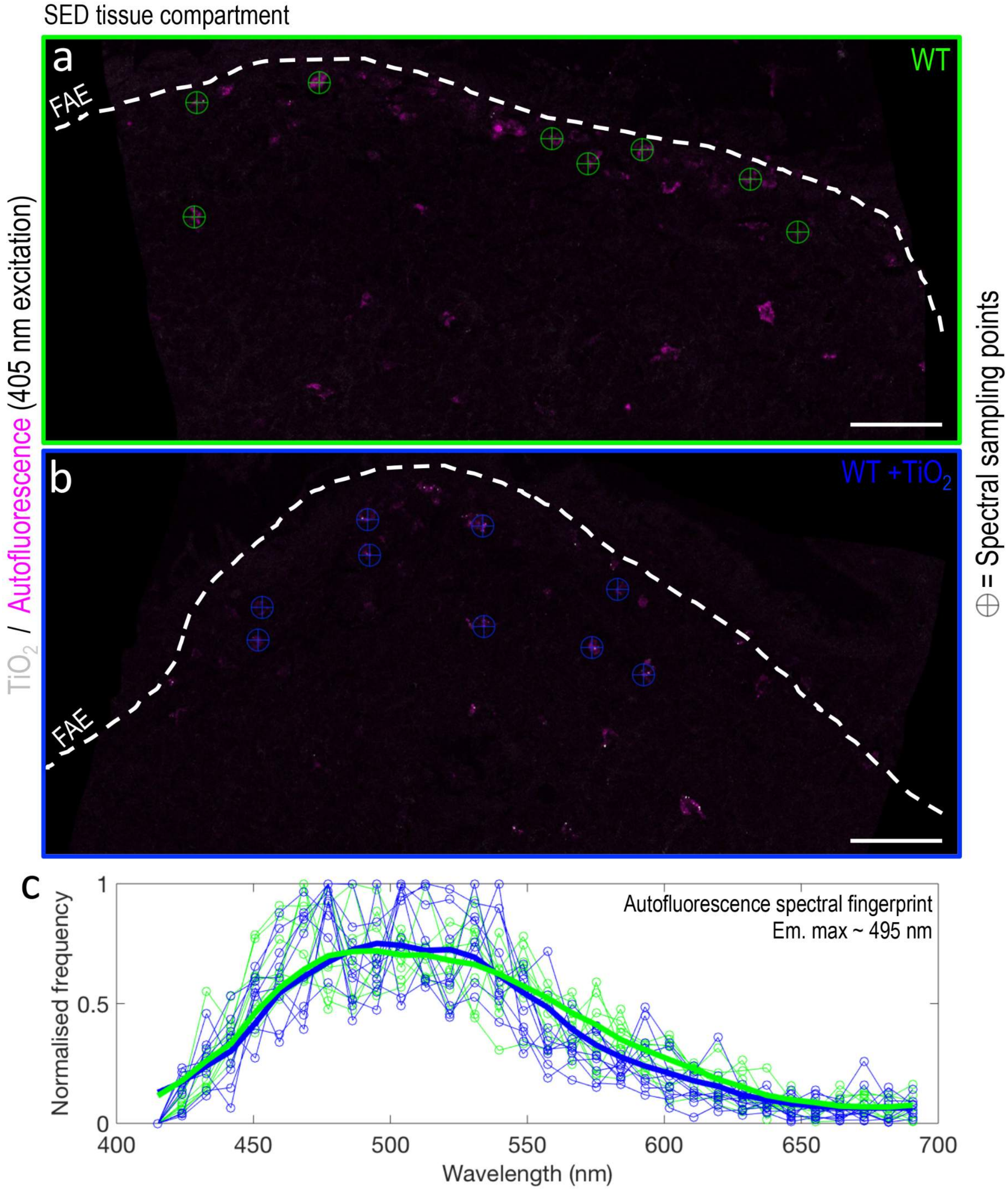


**Supplementary Figure 8 – Determining the autofluorescence emission spectrum of murine LysoMac/LysoDC cells. a/b,** Under 405 nm laser excitation, reflectance data and lambda stacks for unstained murine Peyer's patch tissue sections were collected from animals fed either the (**a**) negative control diet (WT) or the (**b**) control diet supplemented with 0.0625% fg$TiO_2$ (WT +$TiO_2$). In both groups, a population of highly autofluorescent cells were observed in the Peyer's patch subepithelial dome (magenta) that (**b**) co-located with fg$TiO_2$ in the WT+$TiO_2$ samples. **c**, From each lambda stack, spectra were collected from eight locations (indicated by the cross-hair markers in **a/b**) and then averaged (**c**, thick lines) to determine the spectral fingerprint of the autofluorescence signal. Under 405 nm excitation, the emission max was ~ 495 nm. The presence of fg$TiO_2$ did not appear to affect the emission characteristics of the autofluorescence signal. *Scale bars: **a/b** = 50 μm.*

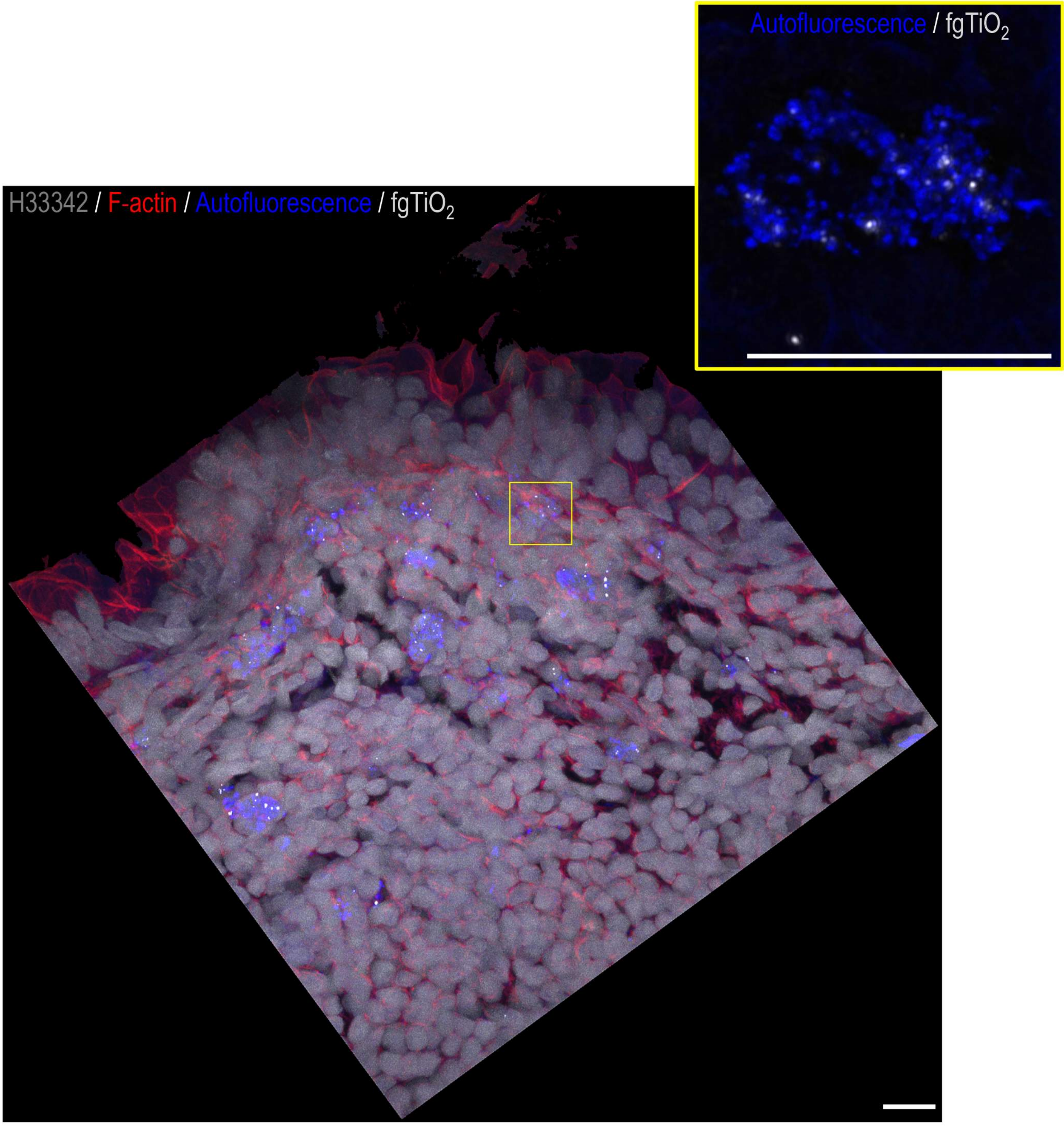


**Supplementary Figure 9 – Selectivity and specificity of fg$TiO_2$ for autofluorescent LysoMac/LysoDC cells in the murine subepithelial dome.** (Main) Max projection of image data collected as a Z-stack using confocal reflectance microscopy. Cell nuclei were fluorescently labelled using Hoechst 33342 (grey). Cytoskeletal actin was labelled with phalloidin-AlexaFluor 647 (red). fg$TiO_2$ (white foci) was seen to selectively and specifically target the highly autofluorescent (blue) mononuclear phagocytic cells. The inset shows a high magnification max projection image of a single autofluorescent cell taken from the area indicated by the yellow box in the main image. The autofluorescence signal was deconvolved (Born and Wolf theoretical point-spread function, Richardson-Lucy deconvolution for 75 iterations via DeconvolutionLab2 software). The output demonstrates the punctate, vesicular nature of the autofluorescence signal and its close association with fg$TiO_2$ at the subcellular level. *Scale bars = 10 µm.*

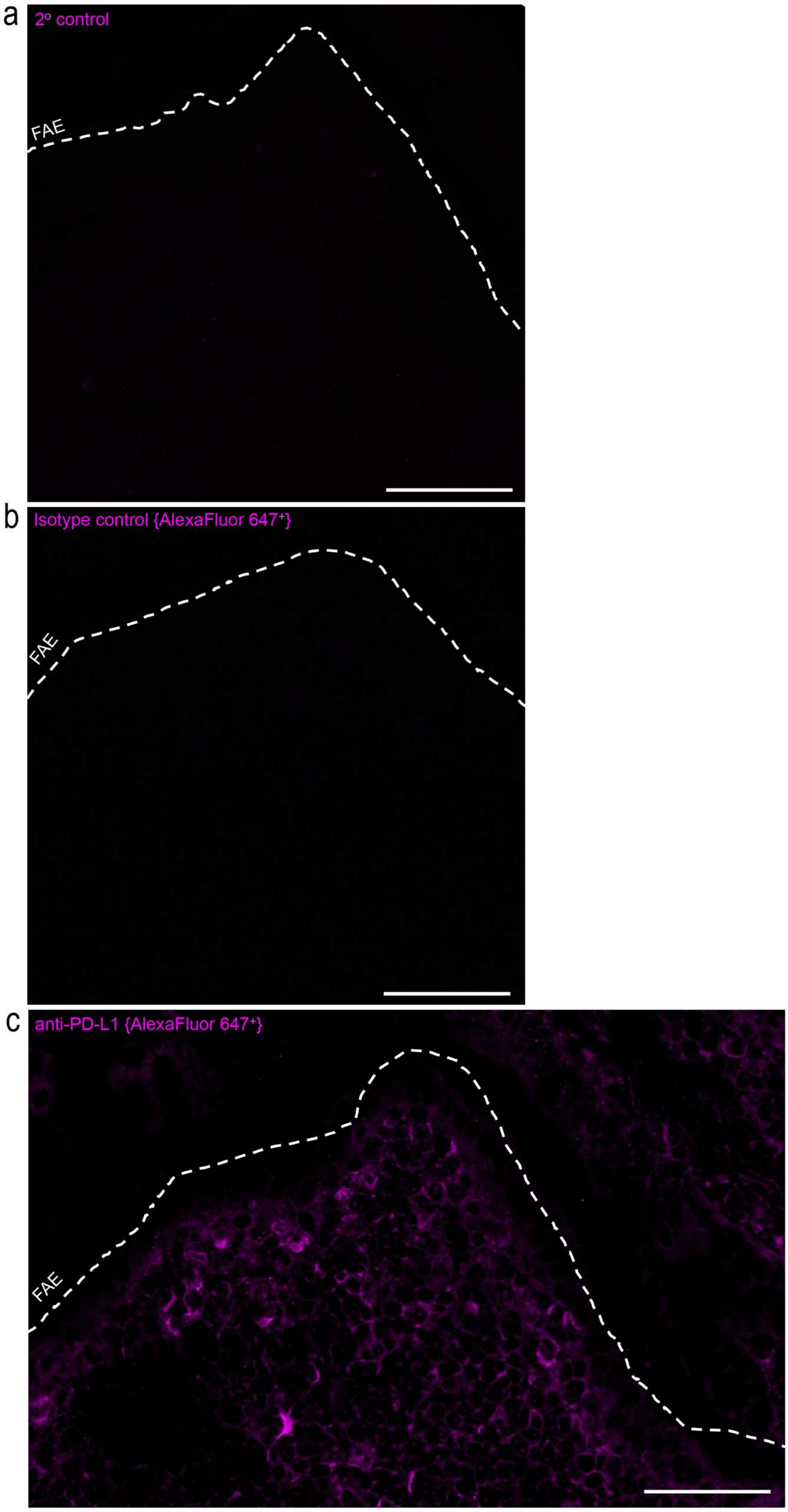


**Supplementary Figure 10 – immunofluorescence control data supporting the anti-PD-L1 immunofluorescence labelling.** Image-data were collected by laser scanning confocal microscopy. **a**, Secondary-only control data (*i.e.,* AlexaFluor 647+ secondary antibody staining in absence of the anti-PD-L1 primary antibody). This shows a clean image with minimal signal in the AlexaFluor 647 channel ruling out non-specific secondary antibody binding and autofluorescence contributions to the measured PD-L1 immunofluorescence data. **b**, Clean images were also obtained when the anti-PD-L1 primary antibody was switched for a concentration-matched 'isotype control' (*i.e.,* an irrelevant antibody of the same isoclass as the anti-PD-L1 primary antibody). This suggests that the anti-PD-L1 primary antibody is not bound non-specifically in this tissue region. **c**, In contrast, strong immunofluorescence labelling was detected in the subepithelial dome using the anti-PD-L1 primary antibody sequentially detected by an AlexaFluor 647-plus secondary antibody. All image data were collected using identical microscope settings. The apical aspect of the follicle-associated epithelium is indicated by the dashed line. *Scale bars = 50 µm.*

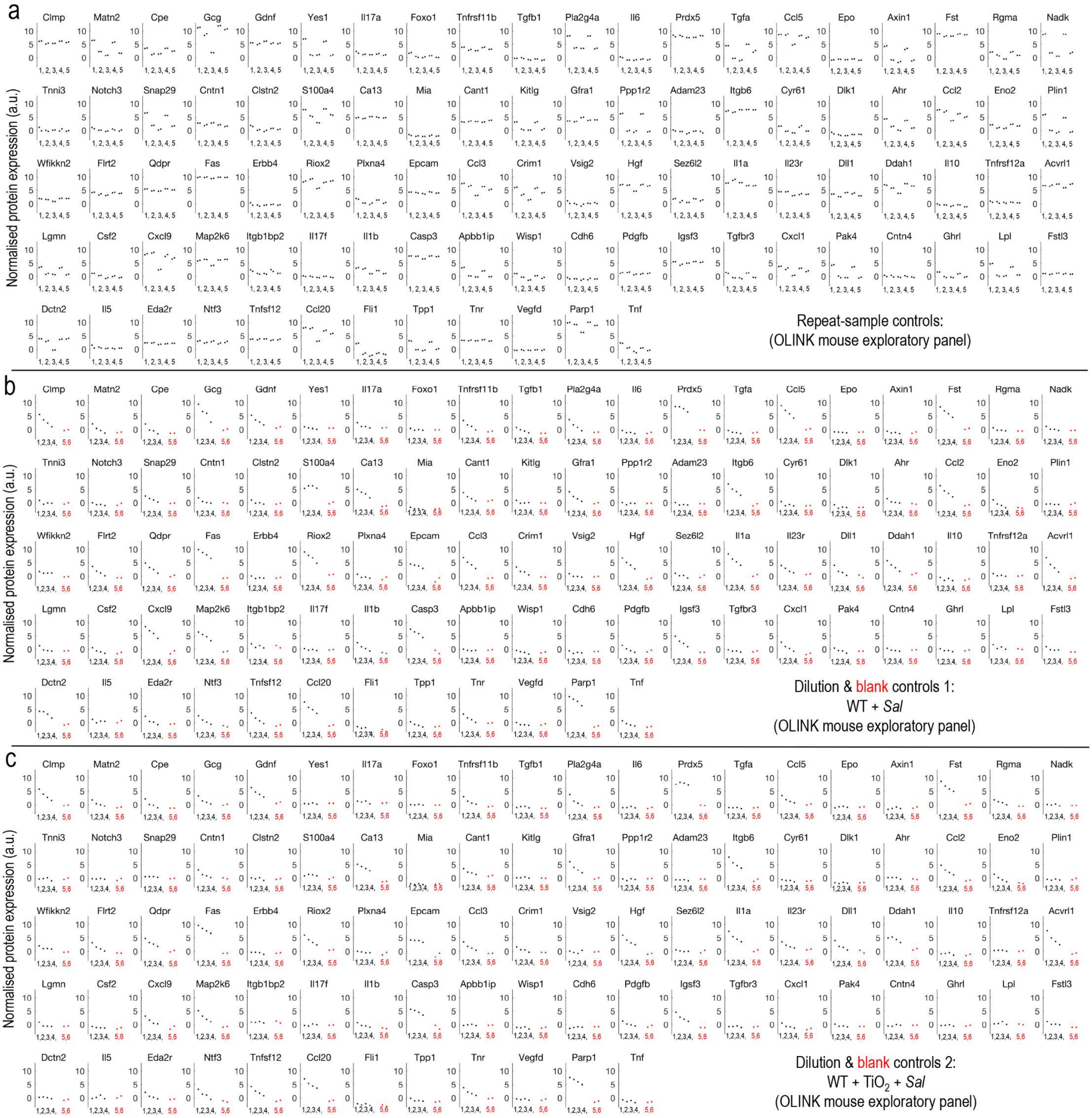


**Supplementary Figure 11 – Proximity extension assay controls. a**, Five random samples across treatment groups were duplicated at random positions on the 96-well plate to assess measurement reproducibility (1=WT+Sal, 2=WT+$TiO_2$+Sal, 3=WT, 4=WT+Sal, 5=WT+$TiO_2$+Sal). **b**/**c**, Serial dilutions of one (**b**) WT + Sal sample and one (**c**) WT +$TiO_2$ +Sal sample were prepared and placed at random positions on the 96-well plate to test for measurement linearity across a wide concentration range. Alongside, two blank lysis buffer-only controls were also included to assess background and indicate limits of detection (1=1:1, 2=1:4, 3=1:8, 4=1:16 dilutions; 5=blank1, 6=blank2).

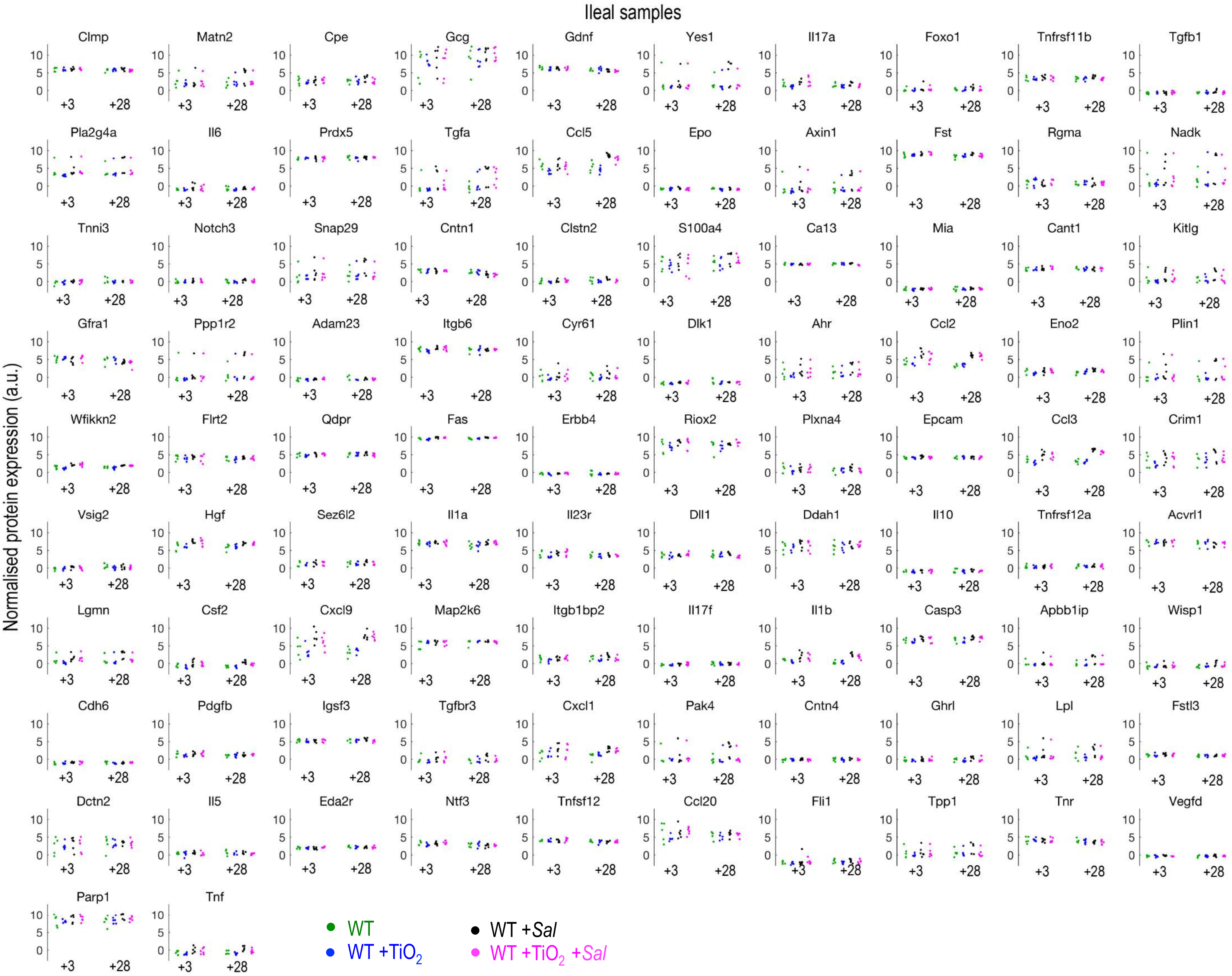


**Supplementary Figure 12 –** Protein expression analyses of ileal tissue digests by proximity extension assay (Olink mouse exploratory panel). In "Mouse Study 2", after the 16-week fg$TiO_2$ feeding period (+$TiO_2$), wild-type animals (WT) were switched to a normal laboratory chow diet (*i.e.,* without fg$TiO_2$-supplementation) then half were orally inoculated with attenuated, Δ*aroA*-*Salmonella* (+*Sal*). Tissues were harvested +3 or +28 days after infection. No significant differences in protein expression levels were observed between the WT and WT +$TiO_2$ ($P \geq 0.53$, two-sided Wilcoxon rank-sum test, n = 3 animals per group) or WT + *Sal* and WT +$TiO_2$ +*Sal* groups ($P \geq 0.79$, two-sided Wilcoxon rank-sum test, n = 6 animals per group) for the 92 protein targets covered by the Olink panel. Protein abbreviations are defined in **Supplementary Table 1**.

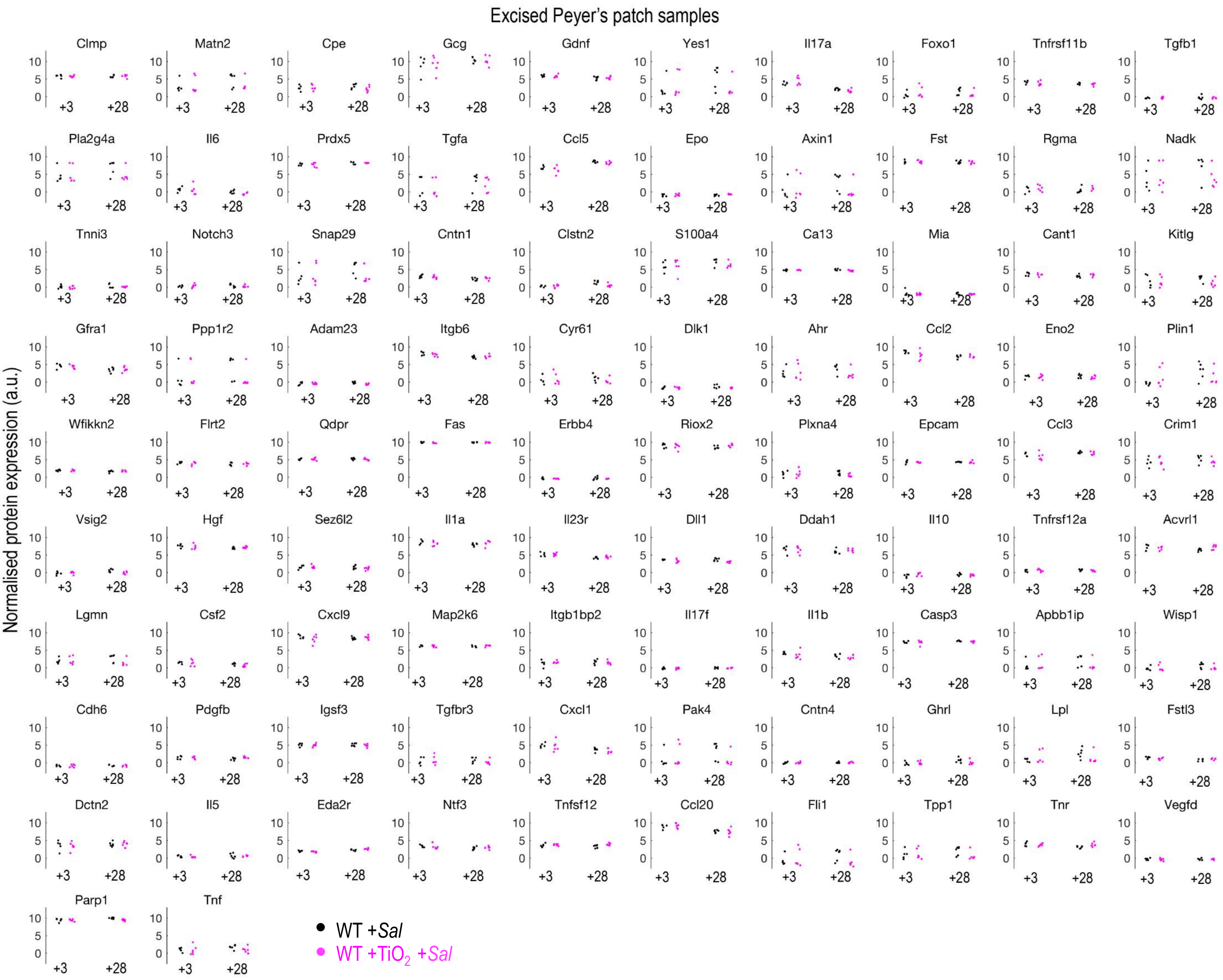


**Supplementary Figure 13 –** Protein expression analyses of Peyer's patch tissue digests by proximity extension assay (Olink mouse exploratory panel). Peyer's patch-focussed analyses were carried out using carefully-excised patch digests from wild-type mice (WT) treated with either Δ*aroA-Salmonella* alone (WT +*Sal*) or with fg$TiO_2$ and Δ*aroA-Salmonella* (WT +$TiO_2$ +*Sal*) at +3 or +28 day timepoints. No significant differences were observed between the two groups ($P \geq 0.2$, two-sided Wilcoxon rank-sum test, n = 6 animals per group) at either timepoint demonstrating no measurable interaction of fg$TiO_2$ and Δ*aroA-Salmonella* for the 92 protein targets covered by the Olink panel. Protein abbreviations are defined in **Supplementary Table 1**.

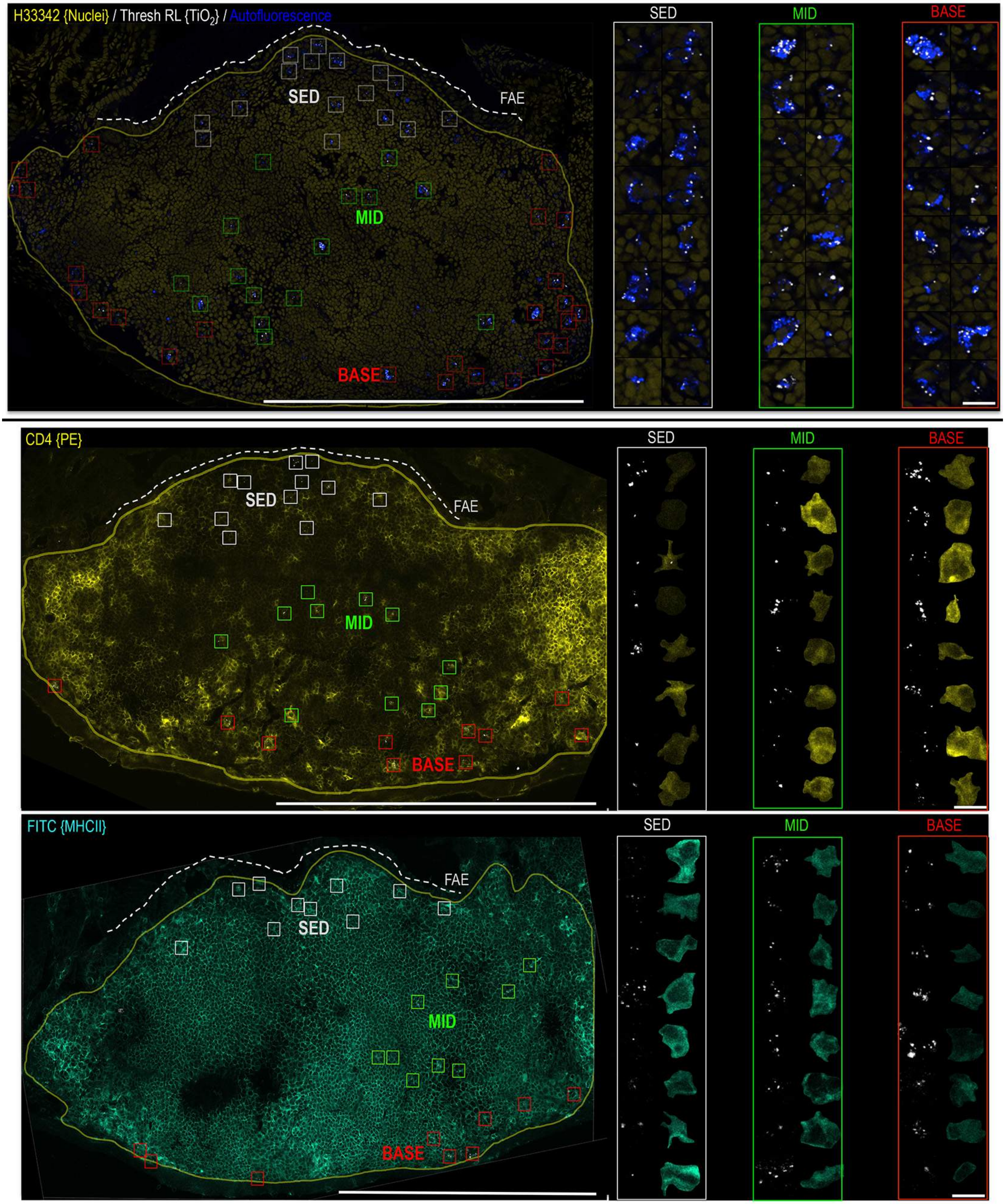


**Supplementary Figure 14 – Autofluorescence, CD4 and MHCII expression in fg$TiO_2$-recipient cells of the Peyer's patch at the +28 day timepoint.** fg$TiO_2$-recipient cells were selected at random from the subepithelial dome (SED), mid and base regions of the patch and montaged to display their fg$TiO_2$ load and concomitant cell expression. Cells maintained the same autofluorescent profile and mixed MHCII and CD4 expression as described in "Mouse Study 1" suggesting a new population of long-lived phagocytes was not formed over time. Visually however, with increasing distance from the SED, an increase in CD4 expression in conjunction with a decrease in MHCII expression was observed, suggesting an shift to sequestration in cells with the longer-lived, LysoMac phenotype. *Scale bars: left = 250 μm; right = 10 μm.*

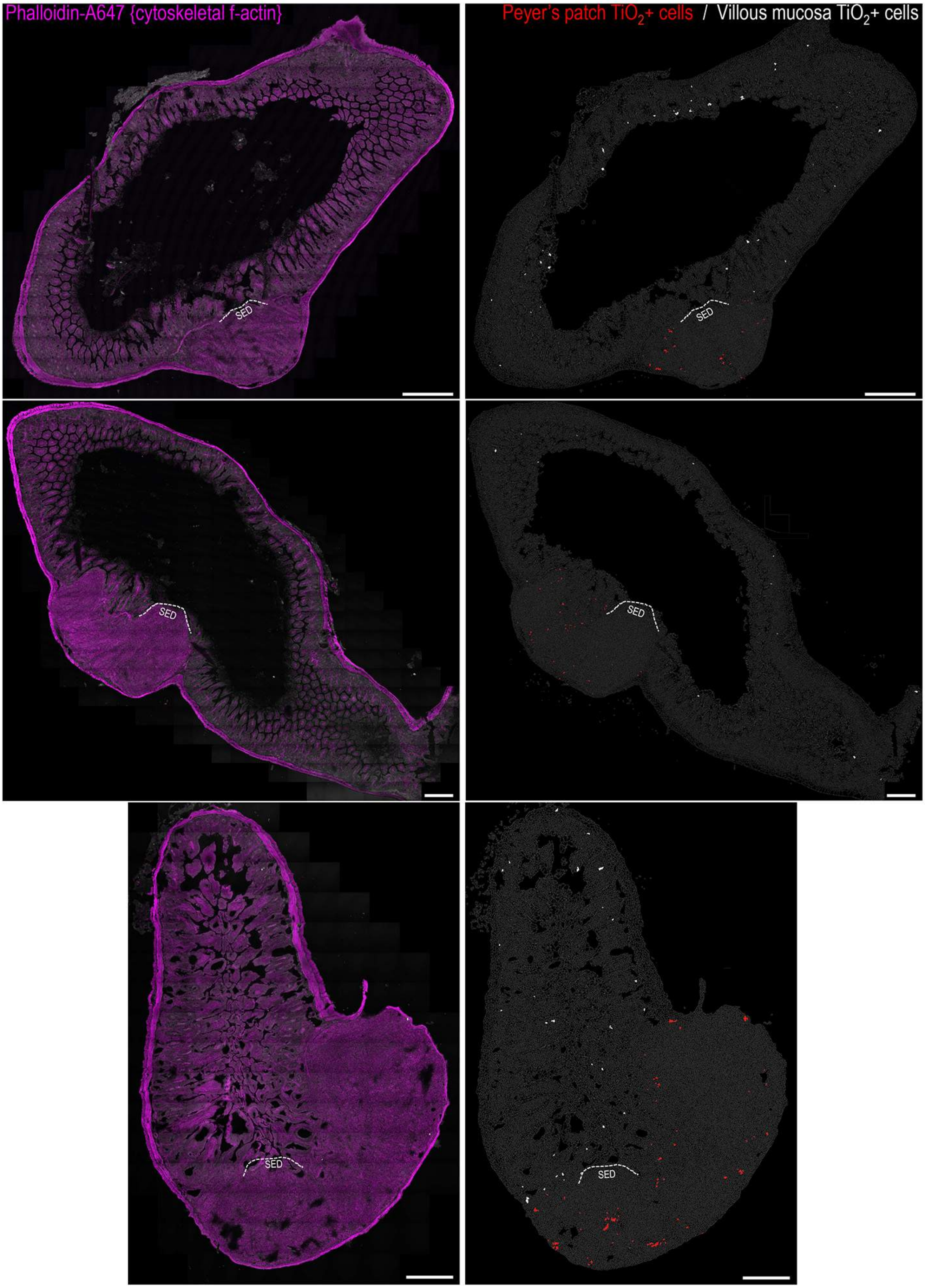


**Supplementary Figure 15 – Ileal distribution of fg$TiO_2$ following *Salmonella* challenge.** (**Right**) Single-cell image analysis of tilescanned (**left**) confocal reflectance images collected from transverse sections of mouse ileal tissue from the fg$TiO_2$-exposed, Δ*aroA*-*Salmonella*-infected treatment group (+28-day timepoint, n = 3 animals). fg$TiO_2$-positive cells in the Peyer's patches are displayed in red. As in **Figure 6**, the subepithelial dome (SED) regions were devoid of fg$TiO_2$ with the majority of the positive cells located basally around the follicle margins. In contrast to all previous data collected without Δ*aroA*-*Salmonella* exposure, fg$TiO_2$-positive cells (displayed in white) were now also present in the villous mucosa. *Scale bars = 250 µm.*

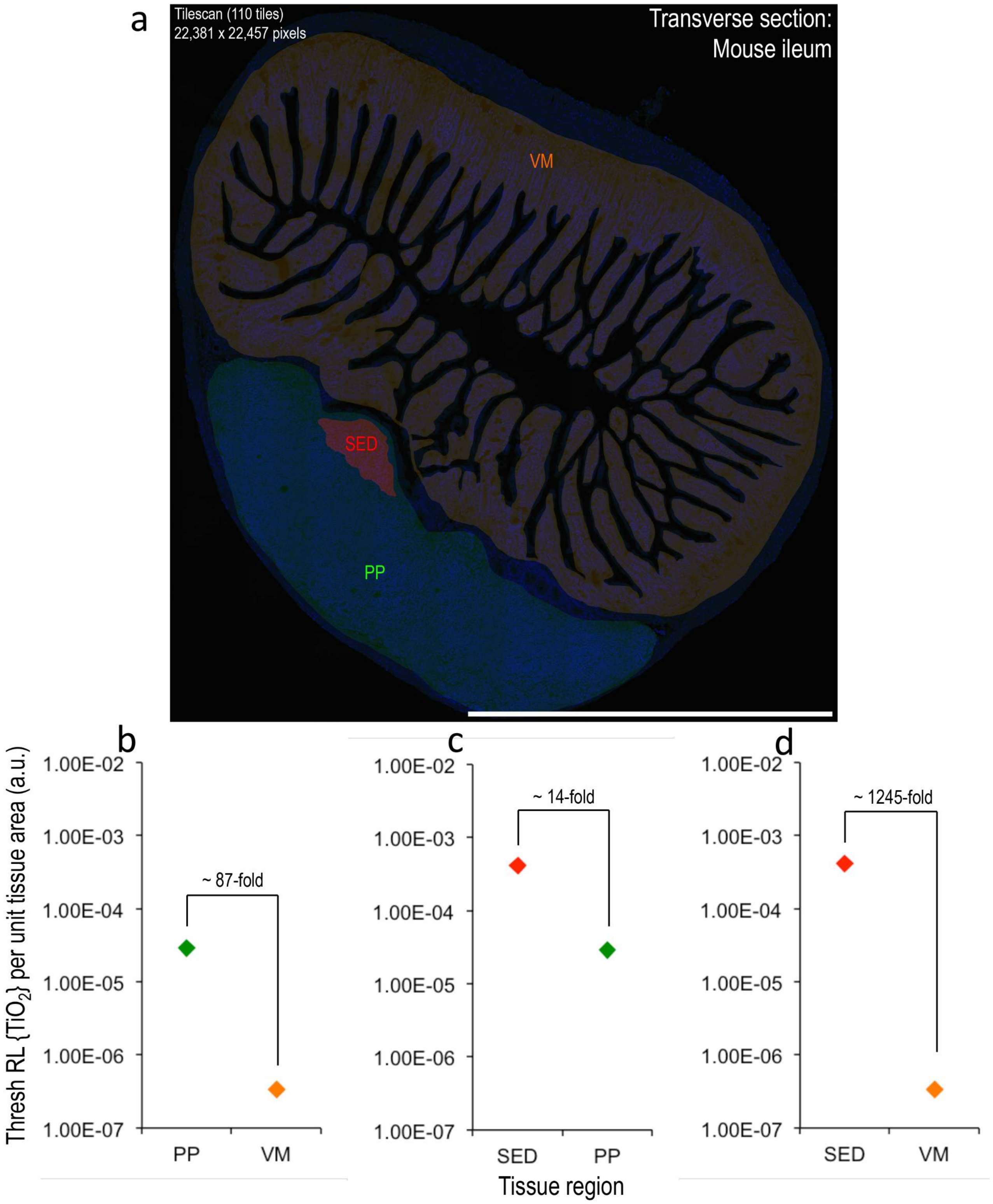


**Supplementary Figure 16 – Quantitative comparison of fg$TiO_2$ loading into different tissue compartments of the murine ileum. a**, Tilescanned reflectance confocal microscopy data (110 stitched images) covering a complete, ileal transverse tissue section containing a Peyer's patch. Manually-drawn region-of-interest masks for the villous mucosa (VM), Peyer's patch (PP plus red SED region) and subepithelial dome (SED) tissue regions are overlaid. **b-d**, The thresholded reflectance signal attributable to fg$TiO_2$ (methodology explained, **Supplementary Figure 1**) was then measured per unit tissue area within these different regions. Despite its large surface area, the normally-absorptive villous mucosa is practically impermeable to fg$TiO_2$. In contrast – facilitated by a overlying epithelium rich in M-cells – the Peyer's patches show considerable uptake. The selectivity for this site means that the concentration gradient for area-corrected reflectance signal was (**b**) ~87-fold higher in the Peyer's patch when directly compared against the surrounding villous mucosa. Moreover, most of this Peyer's patch signal was localised within the immunocompetant SED region where the area-corrected reflectance was (**c**) ~14-fold higher than for the patch overall and, therefore, (**d**) ~1,245 fold higher than that of villous mucosa. *Scale bar = 1 mm.*

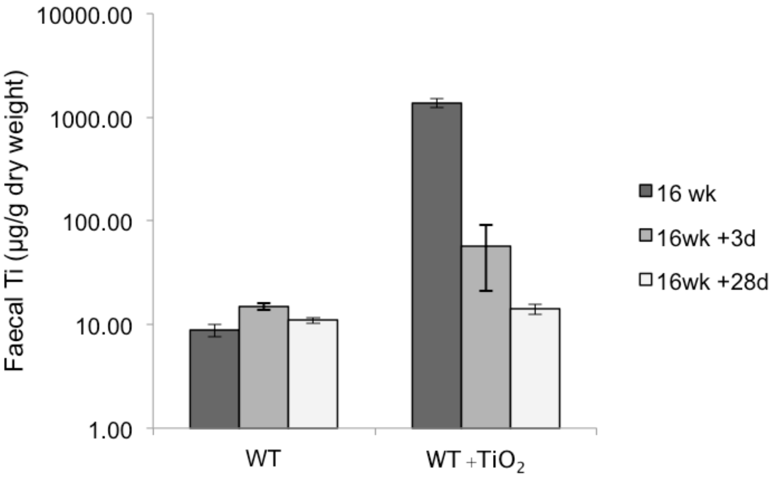


**Supplementary Figure 17 – Analyses of faecal titanium using inductively-coupled plasma mass spectrometry (ICP-MS).** In 'Mouse Study 2', diets containing 0% or 0.0625% fg$TiO_2$ (labelled 'WT' or 'WT+$TiO_2$', respectively) were fed for sixteen weeks. All animals were then switched to a normal diet (*i.e.,* without supplemental fg$TiO_2$) for the remainder of the study. ICP-MS analyses for titanium were conducted from two faecal samples collected from three animals per treatment group at the 16 week, 16 week +3-day or 16 week +28-day timepoints. Elevated Ti (~ 5-fold higher than control-diet animals) was measureable at the 16 week +3-day timepoint but returned to WT control levels by 16 weeks +28 days. The instrument was set up to identify the mass pairs 48 and 64 for isotopic Ti-48 and $TiO_2$-64. The graph represents the average and standard deviation per treatment group and timepoint (n = 3 animals).

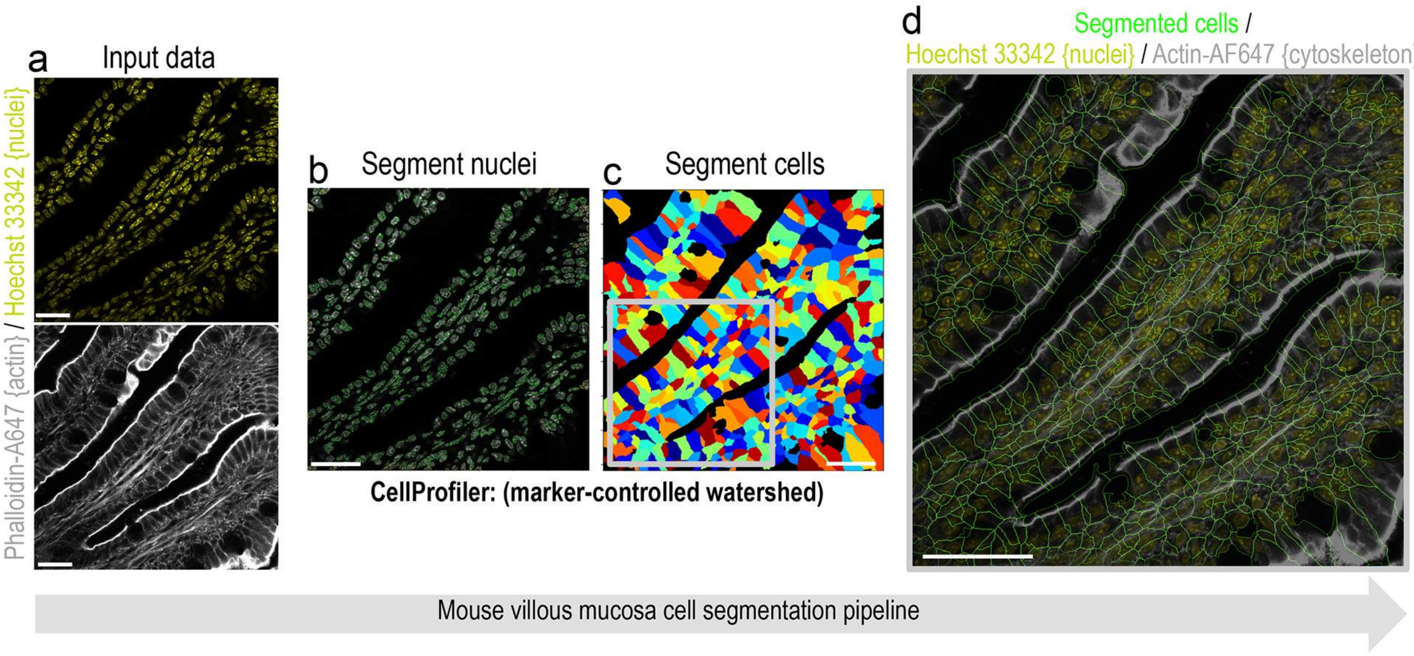


**Supplementary Figure 18 – Cell segmentation of mouse villous mucosa images using a marker-controlled watershed algorithm.** **a,** Nuclei and actin fluorescence images were loaded into a CellProfiler pipeline. **b**, First, nuclei were segmented via an 'IdentifyPrimaryObjects' module using information from the Hoechst 33342 channel. **c**, The resultant nuclei-objects were then used as seeds during deployment of a marker-controlled watershed approach to find each cell's boundary using fluorescence information from the actin channel. **d**, Cell segmentation boundaries overlaid on the nuclei and actin fluorescence information. *Scale bars = 25 µm.*

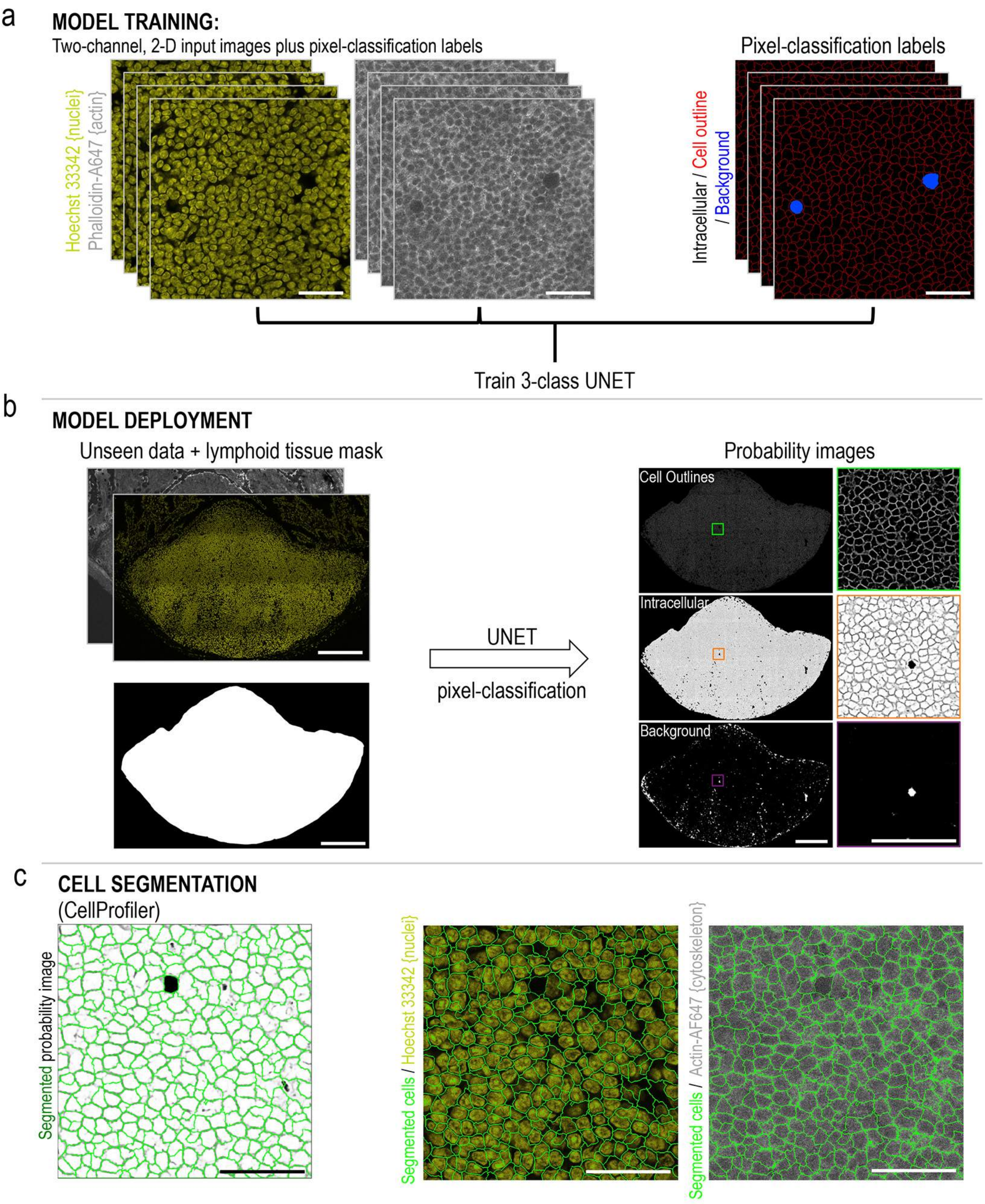


**Supplementary Figure 19 – Cell segmentation of mouse lymphoid tissues using UNET pixel-classification**. **a**, Using the nuclei and actin fluorescence channels alongside paired pixel-classification labels as input, the network was trained to predict the probability that pixels in the input images belonged to one of three classes ('intracellular', 'cell outline' or 'background'). **b**, The trained model was then used to generate probability images for unseen lymphoid tissue images. These probability images represent the likelihood for every pixel in the image belonging to the indicated class. **c**, To segment cell-objects and extract features (*e.g.,* per-cell intensity and size/shape information), the 'intracellular' probability images were loaded into CellProfiler alongside the fluorescence and reflectance data and cell segmentation was achieved from the 'intracellular' probability image using an 'IdentifyPrimaryObjects' module. *Scale bars:* ***a*** *= 25 μm,* ***b*** *= 200 μm (with insets 50 μm),* ***c*** *= 50 μm.*

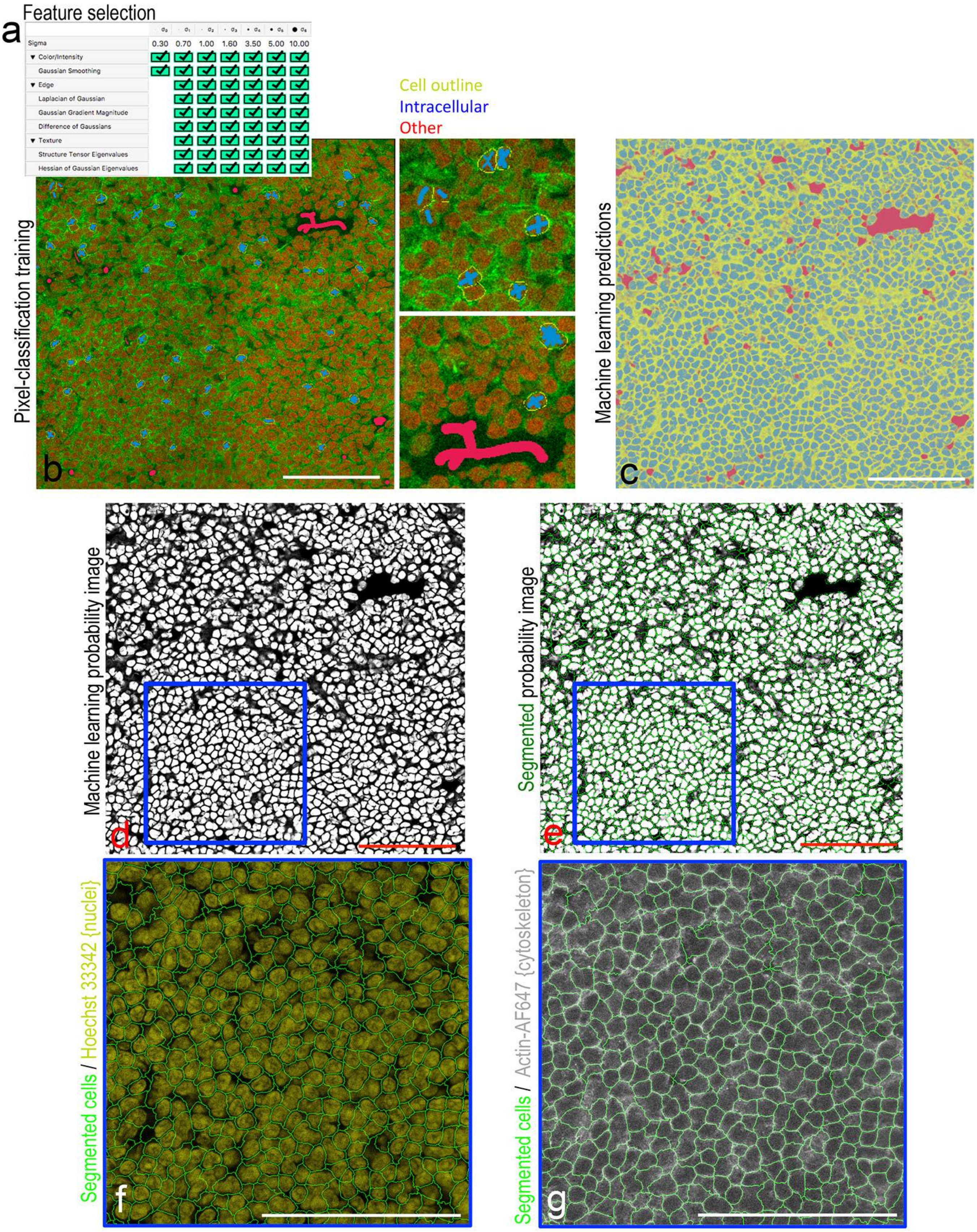


**Supplementary Figure 20 – Cell segmentation of human or mouse Peyer's patch images using pixel-classification machine learning in Ilastik**. (**a**) Feature selection in the Ilastik software. (**b**) A small number of pixel annotation indicating the different classes desired are made by drawing onto the nuclei and actin image data. (**c**) The Ilastik software then uses machine learning to attempt classification of all pixels in the image into each category. The images represent the likelihood for every pixel in the image belonging to the indicated class. (**d-g**), The probability map information can then be segmented into cell objects via an 'IdentifyPrimaryObjects' module in Cell Profiler. *Scale bars = 100 µm.*

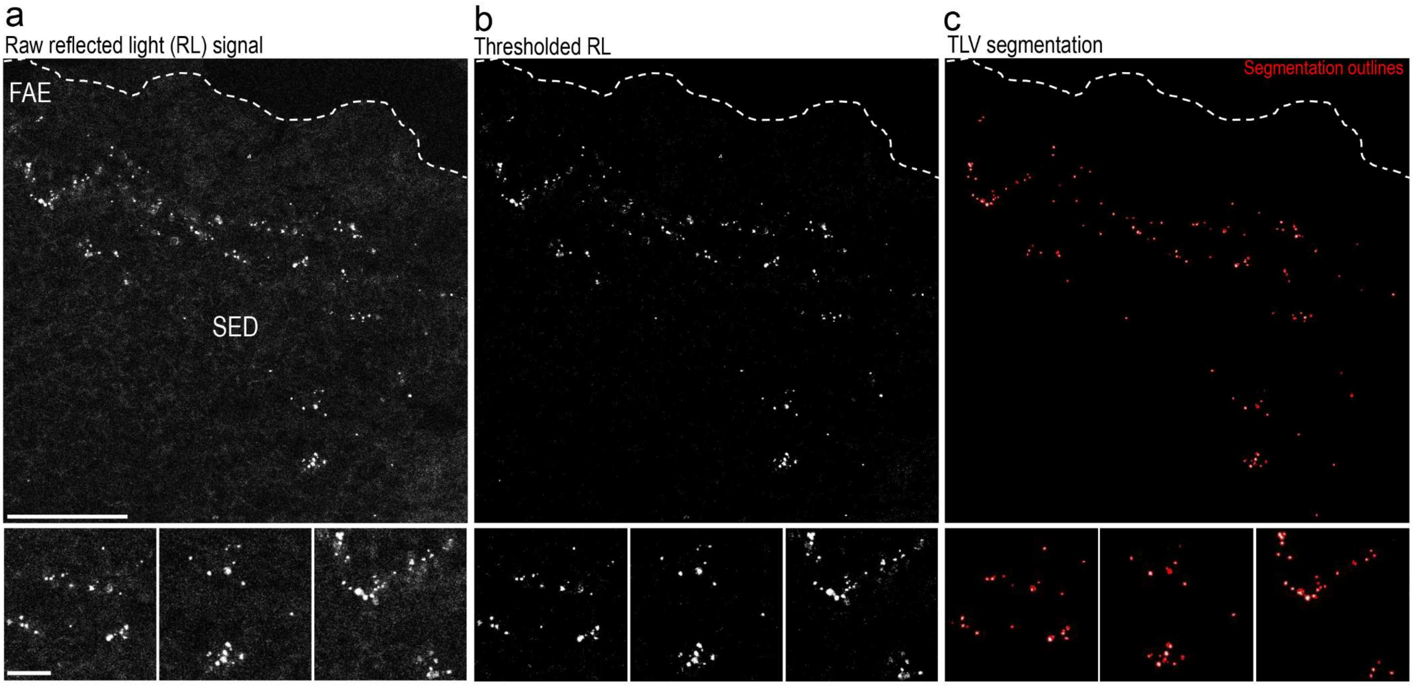


**Supplementary Figure 21 – fg$TiO_2$-loaded vesicle (TLV) object segmentation from reflectance images.** Using a CellProfiler pipeline, the (**a**) raw reflectance information was (**b**) thresholded to remove background signal caused by the biological tissue whilst leaving the punctate, reflectant foci representing fg$TiO_2$. **c**, These thresholded reflectant foci were then segmented into 'fg$TiO_2$-loaded vesicle' objects using an 'IdentifyPrimaryObjects' module in CellProfiler. Red represents the segmentation boundaries for each TLV object. *Scale bars = 100 µm with 10 µm in insets.*

**Supplementary Table 1– Full list of proteins and abbreviated names assessed by the proximity extension assay** (Olink Mouse Exploratory Panel)

| | | | |
|---|---|---|---|
| Amyloid beta A4 precursor protein-binding family B member 1-interacting protein (Apbb1ip ) | Q8R5A3 | Epithelial cell adhesion molecule (Epcam ) | Q99JW5 |
| Appetite-regulating hormone (Ghrl ) | Q9EQX0 | Erythropoietin (Epo ) | P07321 |
| Aryl hydrocarbon receptor (Ahr ) | P30561 | Follistatin (Fst ) | P47931 |
| Axin-1 (Axin1 ) | O35625 | Follistatin-related protein 3 (Fstl3 ) | Q9EQC7 |
| C-C motif chemokine 2 (Ccl2 ) | P10148 | Forkhead box protein O1 (Foxo1 ) | Q9R1E0 |
| C-C motif chemokine 20 (Ccl20 ) | O89093 | Friend leukemia integration 1 transcription factor (Fli1 ) | P26323 |
| C-C motif chemokine 3 (Ccl3 ) | P10855 | Gamma-enolase (Eno2 ) | P17183 |
| C-C motif chemokine 5 (Ccl5 ) | P30882 | GDNF family receptor alpha-1 (Gfra1 ) | P97785 |
| C-X-C motif chemokine 9 (Cxcl9 ) | P18340 | Glial cell line-derived neurotrophic factor (Gdnf ) | P48540 |
| Cadherin-6 (Cdh6 ) | P97326 | Glucagon (Gcg ) | P55095 |
| Calsyntenin-2 (Clstn2 ) | Q9ER65 | Granulocyte-macrophage colony-stimulating factor (Csf2 ) | P01587 |
| Carbonic anhydrase 13 (Ca13 ) | Q9D6N1 | Growth-regulated alpha protein (Cxcl1 ) | P12850 |
| Carboxypeptidase E (Cpe ) | Q00493 | Hepatocyte growth factor (Hgf ) | Q08048 |
| Caspase-3 (Casp3 ) | P70677 | Immunoglobulin superfamily member 3 (Igsf3 ) | Q6ZQA6 |
| Contactin-1 (Cntn1 ) | P12960 | Integrin beta-1-binding protein 2 (Itgb1bp2 ) | Q9R000 |
| Contactin-4 (Cntn4 ) | Q69Z26 | Integrin beta-6 (Itgb6 ) | Q9Z0T9 |
| CXADR-like membrane protein (Clmp ) | Q8R373 | Interleukin-1 alpha (Il1a ) | P01582 |
| Cysteine-rich motor neuron 1 protein (Crim1 ) | Q9JLL0 | Interleukin-1 beta (Il1b ) | P10749 |
| Cytosolic phospholipase A2 (Pla2g4a ) | P47713 | Interleukin-10 (Il10 ) | P18893 |
| Delta-like protein 1 (Dll1 ) | Q61483 | Interleukin-17A (Il17a ) | Q62386 |
| Dihydropteridine reductase (Qdpr ) | Q8BVI4 | Interleukin-17F (Il17f ) | Q7TNI7 |
| Disintegrin and metalloproteinase domain-containing protein 23 (Adam23 ) | Q9R1V7 | Interleukin-23 receptor (Il23r ) | Q8K4B4 |
| | | Interleukin-5 (Il5 ) | P04401 |
| Dual specificity mitogen-activated protein kinase kinase 6 (Map2k6 ) | P70236 | Interleukin-6 (Il6 ) | P08505 |
| Dynactin subunit 2 (Dctn2 ) | Q99KJ8 | Kit ligand (Kitlg ) | P20826 |
| Legumain (Lgmn ) | O89017 | Seizure 6-like protein 2 (Sez6l2 ) | Q4V9Z5 |
| Leucine-rich repeat transmembrane protein FLRT2 (Flrt2 ) | Q8BLU0 | Serine/threonine-protein kinase PAK 4 (Pak4 ) | Q8BTW9 |
| Lipoprotein lipase (Lpl ) | P11152 | Serine/threonine-protein kinase receptor R3 (Acvrl1 ) | Q61288 |
| Matrilin-2 (Matn2 ) | O08746 | Soluble calcium-activated nucleotidase 1 (Cant1 ) | Q8VCF1 |
| Melanoma-derived growth regulatory protein (Mia ) | Q61865 | Synaptosomal-associated protein 29 (Snap29 ) | Q9ERB0 |
| N(G),N(G)-dimethylarginine dimethylaminohydrolase 1 (Ddah1 ) | Q9CWS0 | Tenascin-R (Tnr ) | Q8BYI9 |
| NAD kinase (Nadk ) | P58058 | Transforming growth factor beta receptor type 3 (Tgfbr3 ) | O88393 |
| Neurogenic locus notch homolog protein 3 (Notch3 ) | Q61982 | Latency-associated peptide transforming growth factor beta-1 (Tgfb1 ) | P04202 |
| Neurotrophin-3 (Ntf3 ) | P20181 | Tripeptidyl-peptidase 1 (Tpp1 ) | O89023 |
| Perilipin-1 (Plin1 ) | Q8CGN5 | Troponin I, cardiac muscle (Tnni3 ) | P48787 |
| Peroxiredoxin-5, mitochondrial (Prdx5 ) | P99029 | Tumor necrosis factor (Tnf ) | P06804 |
| Platelet-derived growth factor subunit B (Pdgfb ) | P31240 | Tumor necrosis factor ligand superfamily member 12 (Tnfsf12 ) | O54907 |
| Plexin-A4 (Plxna4 ) | Q80UG2 | Tumor necrosis factor receptor superfamily member 11B (Tnfrsf11b ) | O08712 |
| Poly [ADP-ribose] polymerase 1 (Parp1 ) | P11103 | Tumor necrosis factor receptor superfamily member 12A (Tnfrsf12a ) | Q9CR75 |
| Protein CYR61 (Cyr61 ) | P18406 | Tumor necrosis factor receptor superfamily member 27 (Eda2r ) | Q8BX35 |
| Protein delta homolog 1 (Dlk1 ) | Q09163 | Tumor necrosis factor receptor superfamily member 6 (Fas ) | P25446 |
| Protein phosphatase inhibitor 2 (Ppp1r2 ) | Q9DCL8 | Tyrosine-protein kinase Yes (Yes1 ) | Q04736 |
| Protein S100-A4 (S100a4 ) | P07091 | V-set and immunoglobulin domain-containing protein 2 (Vsig2 ) | Q9Z109 |
| Protransforming growth factor alpha (Tgfa ) | P48030 | Vascular endothelial growth factor D (Vegfd ) | P97946 |
| Receptor tyrosine-protein kinase erbB-4 (Erbb4 ) | Q61527 | WAP, Kazal, immunoglobulin, Kunitz and NTR domain-containing protein 2 (Wfikkn2 ) | Q7TQN3 |
| Repulsive guidance molecule A (Rgma ) | Q6PCX7 | | |
| Ribosomal oxygenase 2 (Riox2 ) | Q8CD15 | WNT1-inducible-signaling pathway protein 1 (Wisp1 ) | O54775 |

**Supplementary Table 2 – Antibody information Table**

| PRIMARY ANTIBODIES | Product no | Supplier | Dilution primary | Stock concentration | Host | Secondary | Detection | Figure |
|---|---|---|---|---|---|---|---|---|
| Anti-mouse GP2 | D278-3 | MBL | | 1 mg/mL | Rat | aRat AF647+ | AlexaFluor 647+ | 2 |
| Anti-mouse CD3 | AB5690 | Abcam | 1:150 | 0.2 mg/mL | Rabbit | aRb AF568 | AlexaFluor 568 | 2 |
| Anti-mouse CD11c | AB33483 | Abcam | 1:400 | 0.5 mg/mL | Armenian Hamster | aHam A488 | AlexaFluor 488 | 2 |
| Anti-mouse B220 | 50045280 | ThermoFisher | 1:50 | 0.2 mg/mL | Rat | - | EFluor 660 | 4 |
| Anti-mouse CD11c | 53011482 | ThermoFisher | 1:25 | 0.2 mg/mL | Rat | - | AlexaFluor 488 | 4 |
| Anti-mouse CD3 | 17003282 | ThermoFisher | 1:50 | 0.2 mg/mL | Rat | - | EFluor 570 | 4 |
| Anti-mouse PD-L1 | AB213480 | Abcam | 1:100 | 0.5 mg/mL | Rabbit | aRb A647+ | AlexaFluor A647+ | 4 |
| Anti-mouse MHCII | PE-65122-100UG | ThermoFisher | 1:25 | 0.2 mg/mL | Rat | - | PE | 4 |
| Anti-mouse CD4 | 12-0041-82 | ThermoFisher | 1:25 | 0.2 mg/mL | Rat | - | PE | 4 |
| Anti-flagellin | AB93713 | Abcam | 1:100 | 1 mg/mL | Rabbit | aRb A568 | AlexaFluor A568 | 5 |
| Anti-CD80 | 17-0801-81 | ThermoFisher | 1:25 | 0.2 mg/mL | Rat | - | APC | Supplementary |
| Anti-CD86 | 12-0862-81 | ThermoFisher | 1:25 | 0.2 mg/mL | Rat | - | PE | Supplementary |
| Anti-MHCII | 11-5321-81 | ThermoFisher | 1:50 | 0.5 mg/mL | Rat | - | FITC | 5 / Supplementary |

| SECONDARY ANTIBODIES | Product no | Supplier | Stock concentration | Host | Secondary dilution | Fluorophore |
|---|---|---|---|---|---|---|
| Goat anti-Rabbit IgG (H+L) | A-11011 | Thermo-Fisher | 2 mg/mL | Goat | 1:400 | AlexaFluor568 |
| Goat anti-Hamster IgG (H+L) | A-11008 | Thermo-Fisher | 2 mg/mL | Goat | 1:400 | AlexaFluor488 |
| Goat anti-Rabbit IgG (H+L) | A32733TR | Thermo-Fisher | 2 mg/mL | Goat | 1:400 | AlexFluor647 plus |
| Goat anti-Rat IgG (H+L) | A48265 | Thermo-Fisher | 2 mg/mL | Goat | 1:400 | AlexFluor647 plus |

**Supplementary Video 1 – 3-D animation demonstrating the selectivity and specificity of fg$TiO_2$ for autofluorescent LysoMac / LysoDC immune cells.** The video shows a 3-D render of image-data collected as a Z-stack from the murine subepithelial dome tissue compartment using confocal reflectance microscopy. Cell nuclei were fluorescently labelled using Hoechst 33342 (grey). Cytoskeletal actin was labelled with phalloidin-AlexaFluor 647 (red). fg$TiO_2$ (white foci) are seen to selectively and specifically target the highly autofluorescent (blue) mononuclear phagocytic cells.

**Supplementary Note 1 – Mouse villous mucosa image analysis pipeline.** This section presents screenshots of the CellProfiler image analysis pipeline used to measure per-cell intensity and shape/size features enabling the detection of fg$TiO_2$ in the mouse villous mucosa images. The pipeline and sample image data are available for download from the BioStudies archive accompanying the paper.

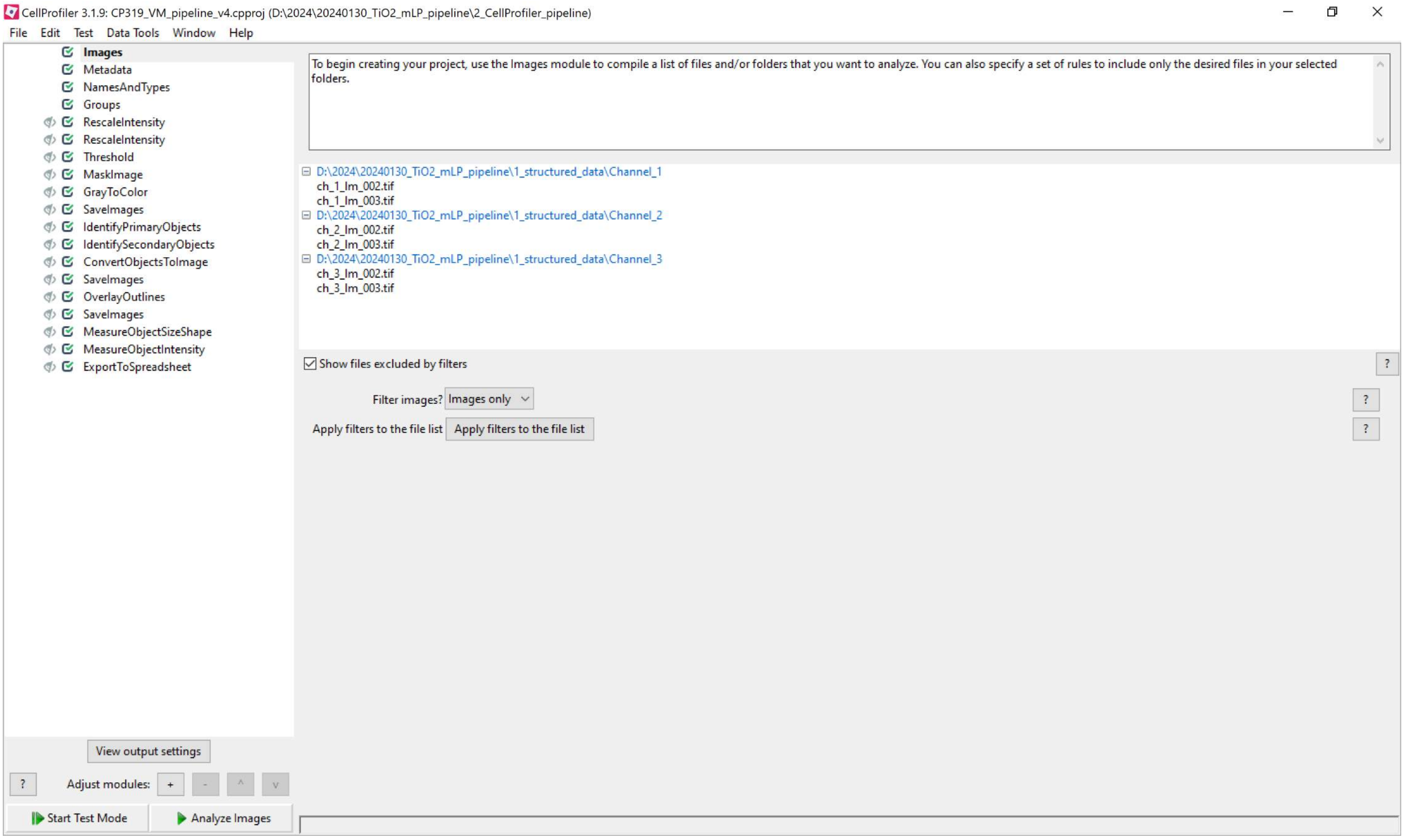

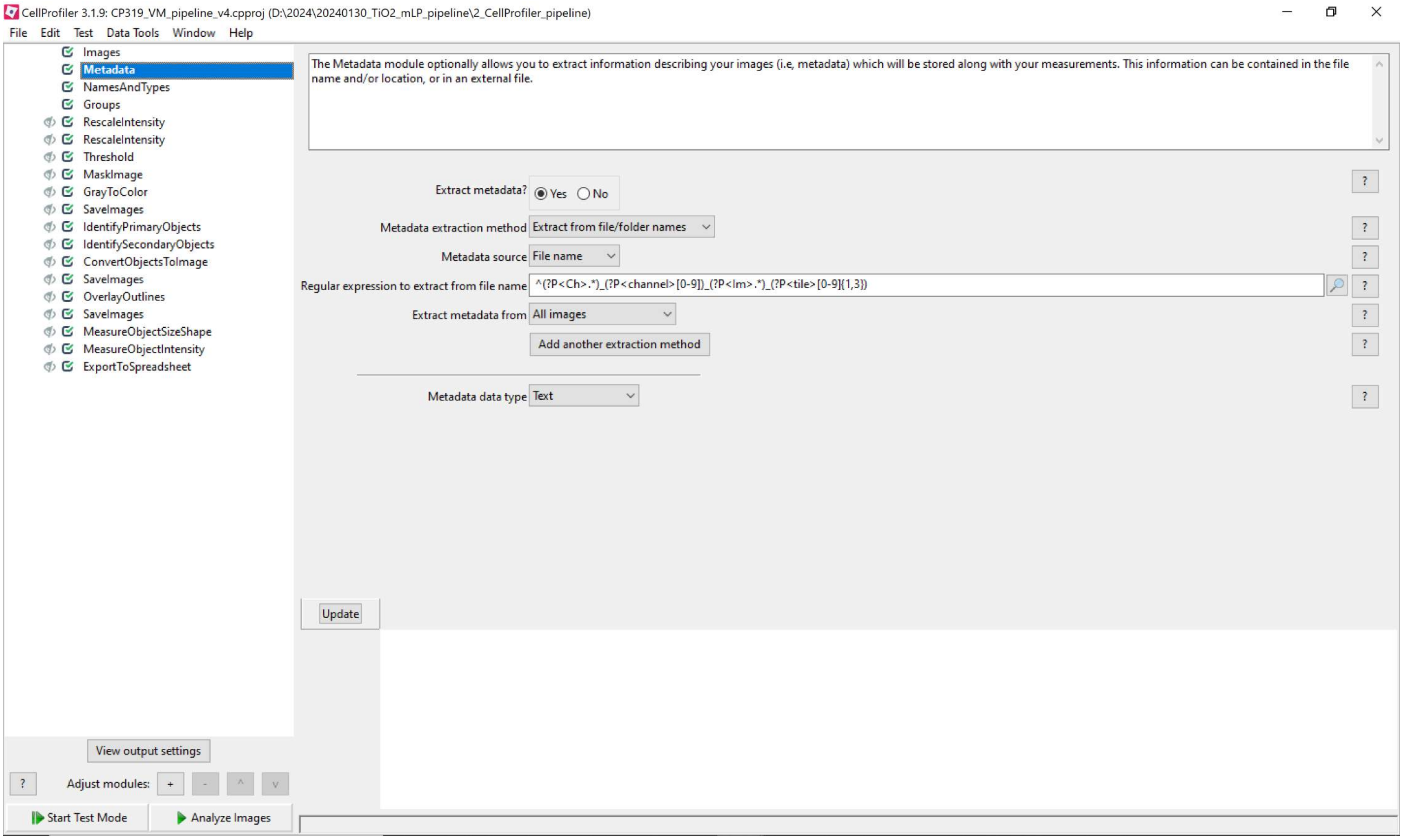
CellProfiler 3.1.9: CP319_VM_pipeline_v4.cpproj (D:\2024\20240130_TiO2_mLP_pipeline\2_CellProfiler_pipeline)
File Edit Test Data Tools Window Help
Images
Metadata
NamesAndTypes
Groups
RescaleIntensity
RescaleIntensity
Threshold
MaskImage
GrayToColor
SaveImages
IdentifyPrimaryObjects
IdentifySecondaryObjects
ConvertObjectsToImage
SaveImages
OverlayOutlines
SaveImages
MeasureObjectSizeShape
MeasureObjectIntensity
ExportToSpreadsheet
The Metadata module optionally allows you to extract information describing your images (i.e, metadata) which will be stored along with your measurements. This information can be contained in the file name and/or location, or in an external file.
Extract metadata? Yes No
Metadata extraction method Extract from file/folder names
Metadata source File name
Regular expression to extract from file name ^(?P<Ch>.*)_(?P<channel>[0-9])_(?P<Im>.*)_(?P<tile>[0-9]{1,3})
Extract metadata from All images
Add another extraction method
Metadata data type Text
Update
View output settings
Adjust modules:
Start Test Mode
Analyze Images

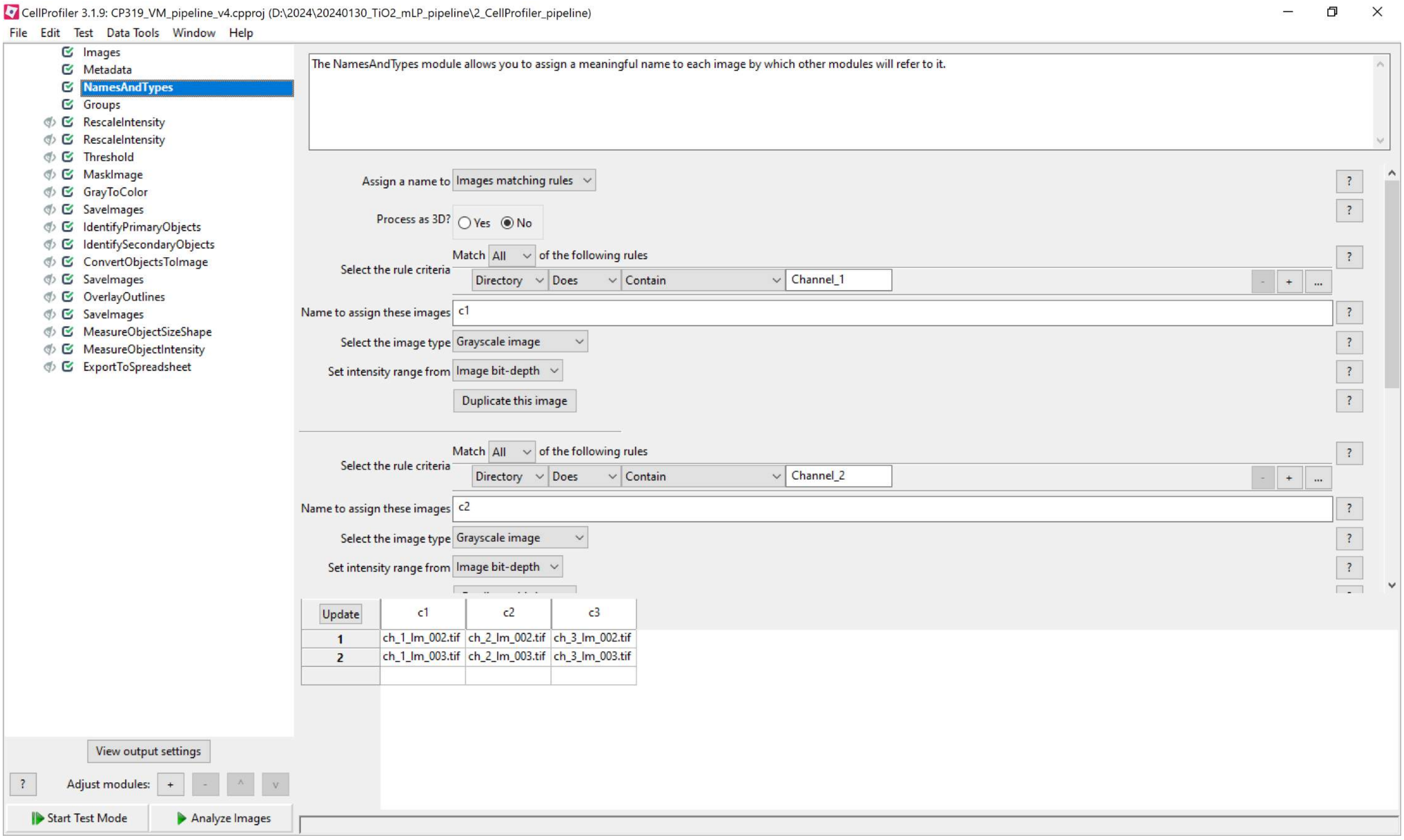
CellProfiler 3.1.9: CP319_VM_pipeline_v4.cpproj (D:\2024\20240130_TiO2_mLP_pipeline\2_CellProfiler_pipeline)
File Edit Test Data Tools Window Help
Images
Metadata
NamesAndTypes
Groups
RescaleIntensity
RescaleIntensity
Threshold
MaskImage
GrayToColor
SaveImages
IdentifyPrimaryObjects
IdentifySecondaryObjects
ConvertObjectsToImage
SaveImages
OverlayOutlines
SaveImages
MeasureObjectSizeShape
MeasureObjectIntensity
ExportToSpreadsheet
The NamesAndTypes module allows you to assign a meaningful name to each image by which other modules will refer to it.
Assign a name to Images matching rules
Process as 3D? Yes No
Match All of the following rules
Select the rule criteria
Directory Does Contain Channel_1
Name to assign these images c1
Select the image type Grayscale image
Set intensity range from Image bit-depth
Duplicate this image
Match All of the following rules
Select the rule criteria
Directory Does Contain Channel_2
Name to assign these images c2
Select the image type Grayscale image
Set intensity range from Image bit-depth
Update
c1
c2
c3
1 ch_1_Im_002.tif ch_2_Im_002.tif ch_3_Im_002.tif
2 ch_1_Im_003.tif ch_2_Im_003.tif ch_3_Im_003.tif
View output settings
Adjust modules:
Start Test Mode
Analyze Images

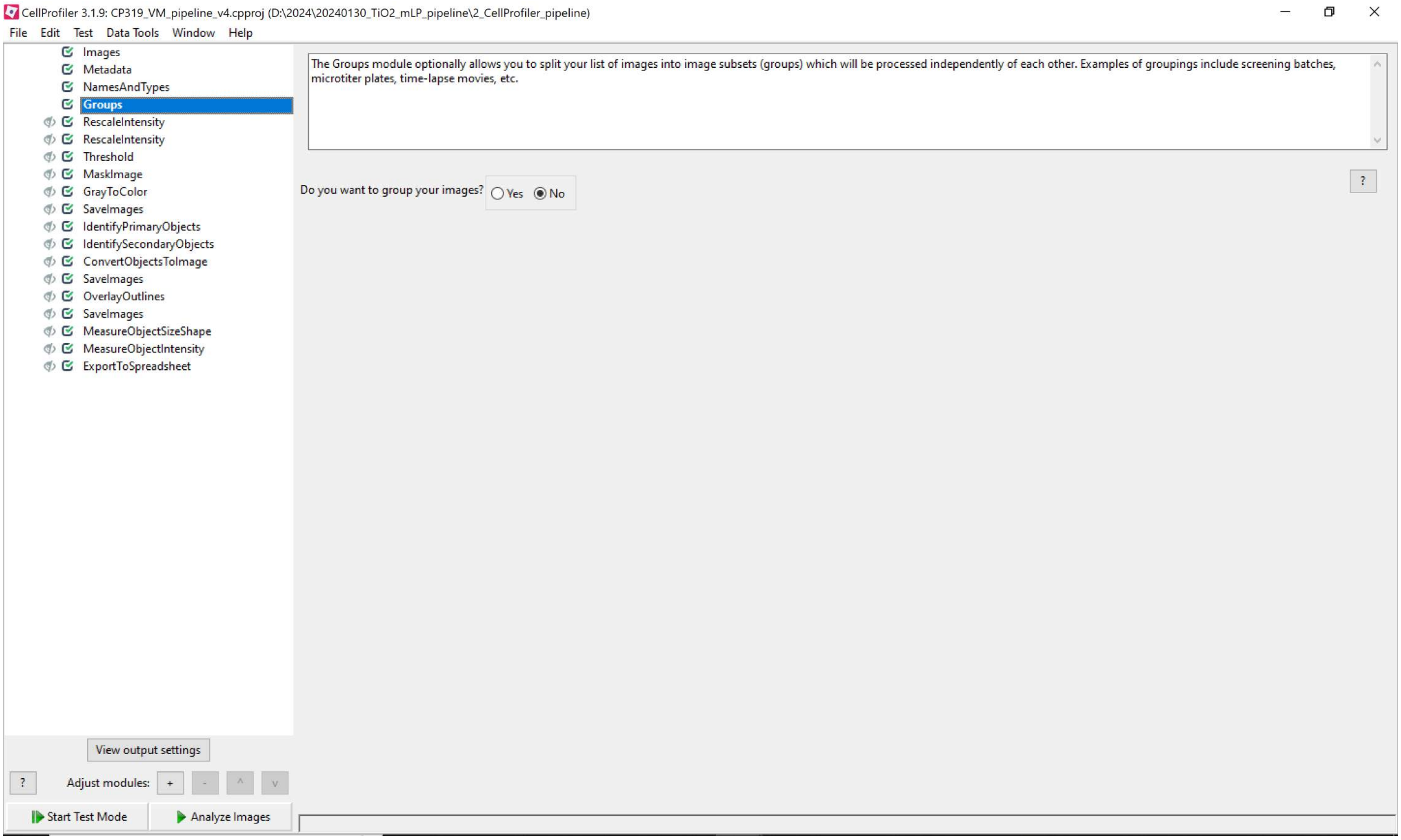
CellProfiler 3.1.9: CP319_VM_pipeline_v4.cpproj (D:\2024\20240130_TiO2_mLP_pipeline\2_CellProfiler_pipeline)
File Edit Test Data Tools Window Help
Images
Metadata
NamesAndTypes
Groups
RescaleIntensity
RescaleIntensity
Threshold
MaskImage
GrayToColor
SaveImages
IdentifyPrimaryObjects
IdentifySecondaryObjects
ConvertObjectsToImage
SaveImages
OverlayOutlines
SaveImages
MeasureObjectSizeShape
MeasureObjectIntensity
ExportToSpreadsheet
The Groups module optionally allows you to split your list of images into image subsets (groups) which will be processed independently of each other. Examples of groupings include screening batches, microtiter plates, time-lapse movies, etc.
Do you want to group your images?
Yes
No
View output settings
Adjust modules:
Start Test Mode
Analyze Images

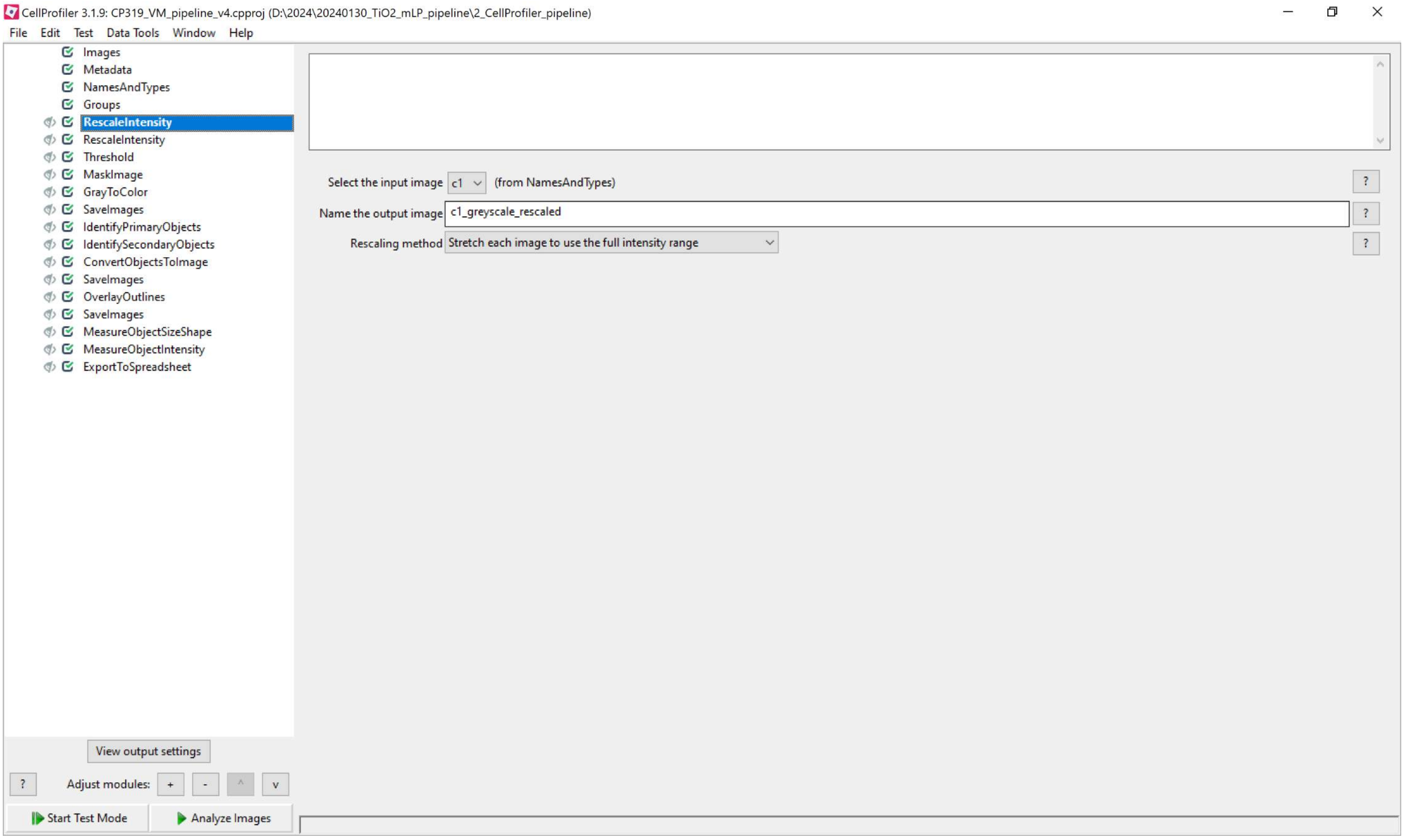
CellProfiler 3.1.9: CP319_VM_pipeline_v4.cpproj (D:\2024\20240130_TiO2_mLP_pipeline\2_CellProfiler_pipeline)
File Edit Test Data Tools Window Help
Images
Metadata
NamesAndTypes
Groups
RescaleIntensity
RescaleIntensity
Threshold
MaskImage
GrayToColor
SaveImages
IdentifyPrimaryObjects
IdentifySecondaryObjects
ConvertObjectsToImage
SaveImages
OverlayOutlines
SaveImages
MeasureObjectSizeShape
MeasureObjectIntensity
ExportToSpreadsheet
Select the input image c1 (from NamesAndTypes)
Name the output image c1_greyscale_rescaled
Rescaling method Stretch each image to use the full intensity range
View output settings
Adjust modules: + - ^ v
Start Test Mode
Analyze Images

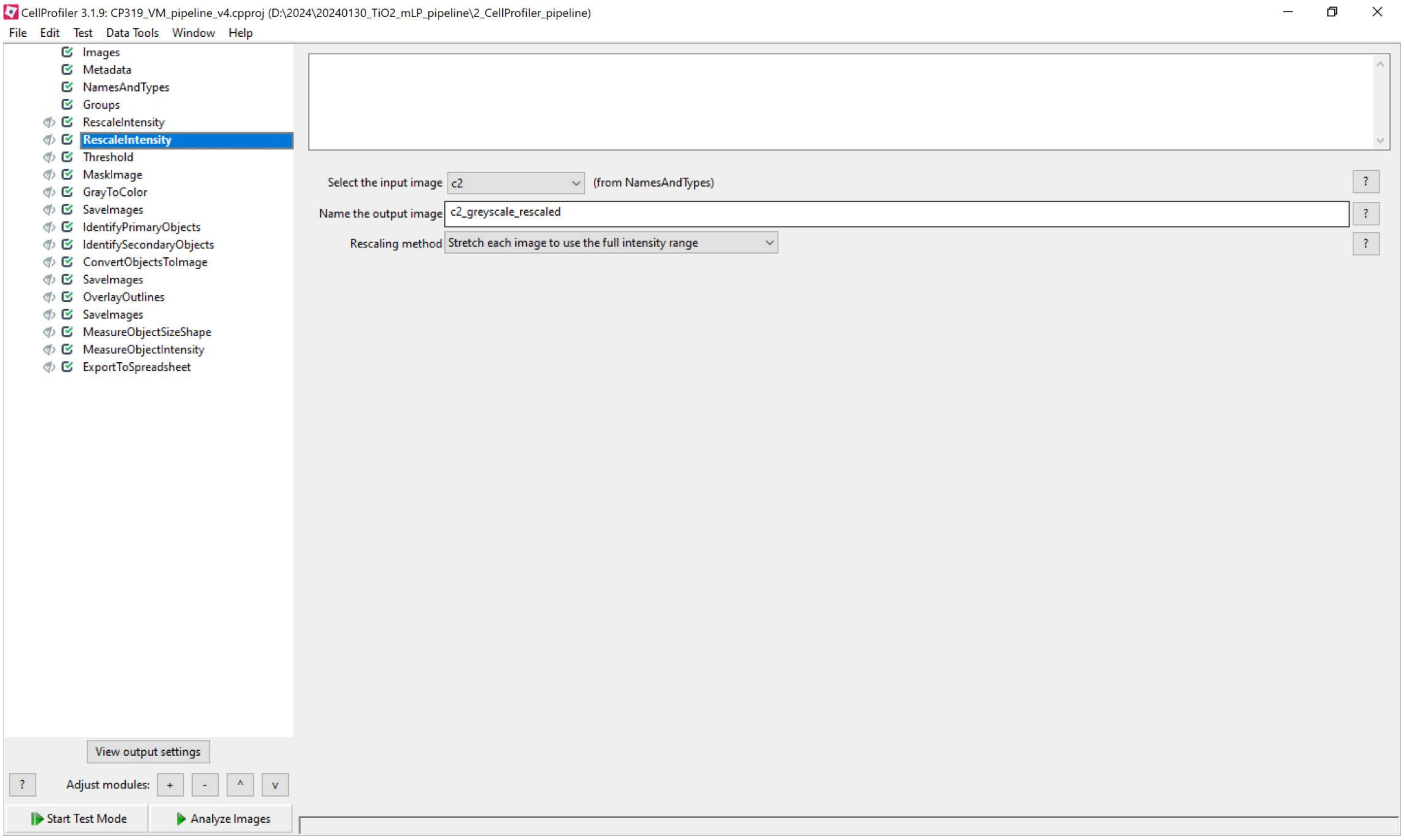
CellProfiler 3.1.9: CP319_VM_pipeline_v4.cpproj (D:\2024\20240130_TiO2_mLP_pipeline\2_CellProfiler_pipeline)
File Edit Test Data Tools Window Help
Images
Metadata
NamesAndTypes
Groups
RescaleIntensity
RescaleIntensity
Threshold
MaskImage
GrayToColor
SaveImages
IdentifyPrimaryObjects
IdentifySecondaryObjects
ConvertObjectsToImage
SaveImages
OverlayOutlines
SaveImages
MeasureObjectSizeShape
MeasureObjectIntensity
ExportToSpreadsheet
Select the input image
c2
(from NamesAndTypes)
Name the output image
c2_greyscale_rescaled
Rescaling method
Stretch each image to use the full intensity range
View output settings
Adjust modules:
Start Test Mode
Analyze Images

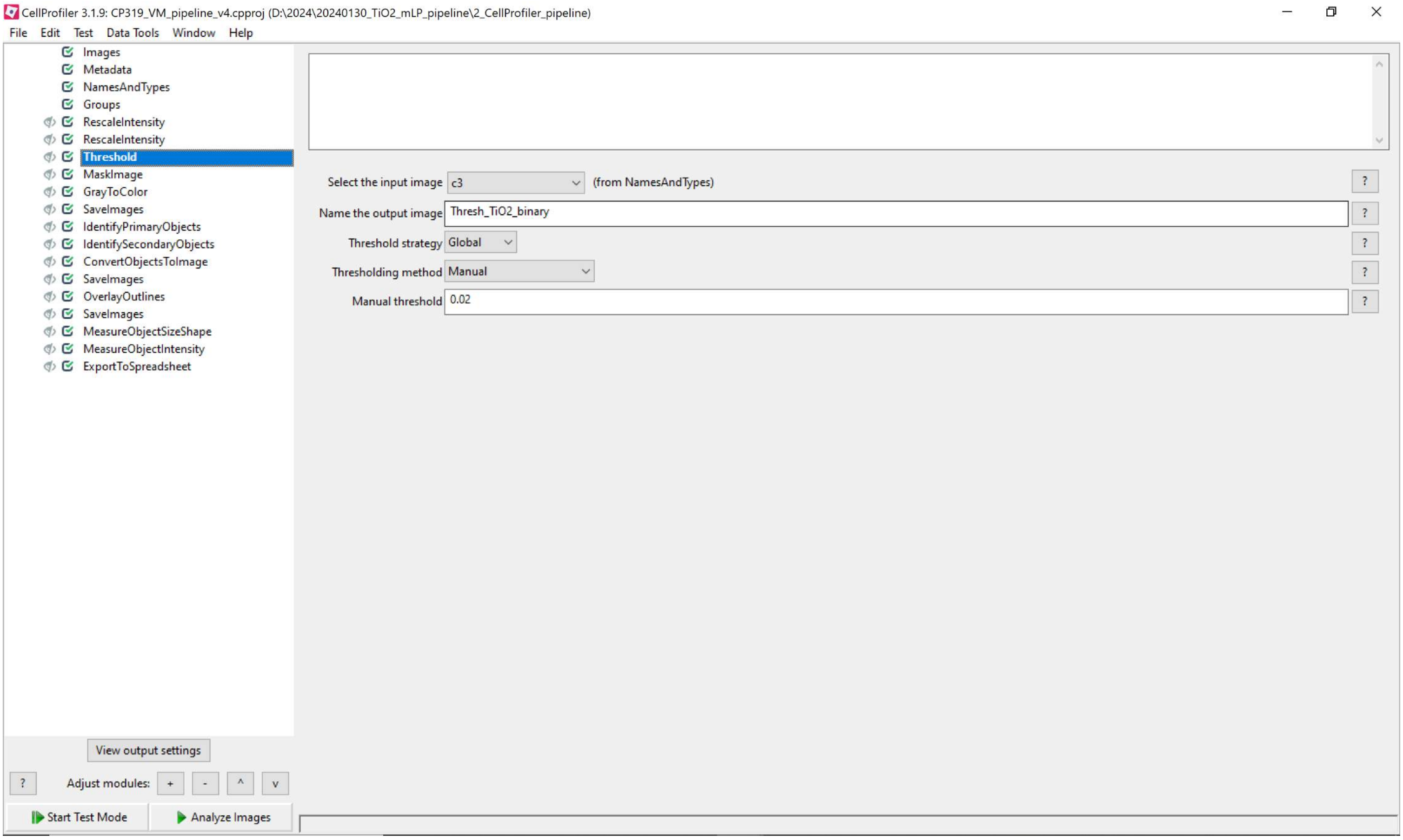
CellProfiler 3.1.9: CP319_VM_pipeline_v4.cpproj (D:\2024\20240130_TiO2_mLP_pipeline\2_CellProfiler_pipeline)
File Edit Test Data Tools Window Help
Images
Metadata
NamesAndTypes
Groups
RescaleIntensity
RescaleIntensity
Threshold
MaskImage
GrayToColor
SaveImages
IdentifyPrimaryObjects
IdentifySecondaryObjects
ConvertObjectsToImage
SaveImages
OverlayOutlines
SaveImages
MeasureObjectSizeShape
MeasureObjectIntensity
ExportToSpreadsheet
Select the input image c3 (from NamesAndTypes)
Name the output image Thresh_TiO2_binary
Threshold strategy Global
Thresholding method Manual
Manual threshold 0.02
View output settings
Adjust modules: + - ^ v
Start Test Mode
Analyze Images

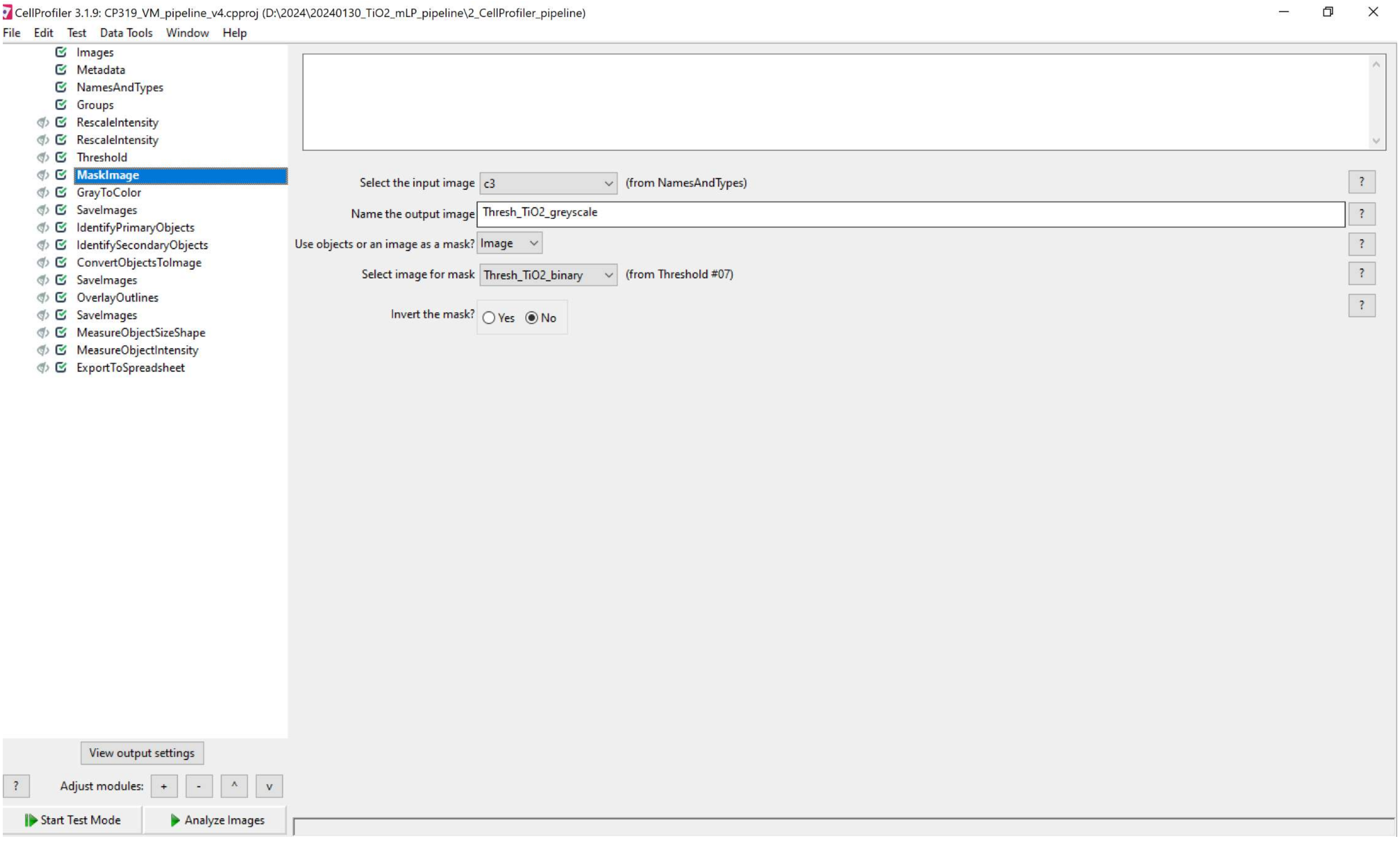
CellProfiler 3.1.9: CP319_VM_pipeline_v4.cpproj (D:\2024\20240130_TiO2_mLP_pipeline\2_CellProfiler_pipeline)
File Edit Test Data Tools Window Help
Images
Metadata
NamesAndTypes
Groups
RescaleIntensity
RescaleIntensity
Threshold
MaskImage
GrayToColor
SaveImages
IdentifyPrimaryObjects
IdentifySecondaryObjects
ConvertObjectsToImage
SaveImages
OverlayOutlines
SaveImages
MeasureObjectSizeShape
MeasureObjectIntensity
ExportToSpreadsheet
Select the input image c3 (from NamesAndTypes)
Name the output image Thresh_TiO2_greyscale
Use objects or an image as a mask? Image
Select image for mask Thresh_TiO2_binary (from Threshold #07)
Invert the mask? Yes No
View output settings
Adjust modules: + - ^ v
Start Test Mode
Analyze Images

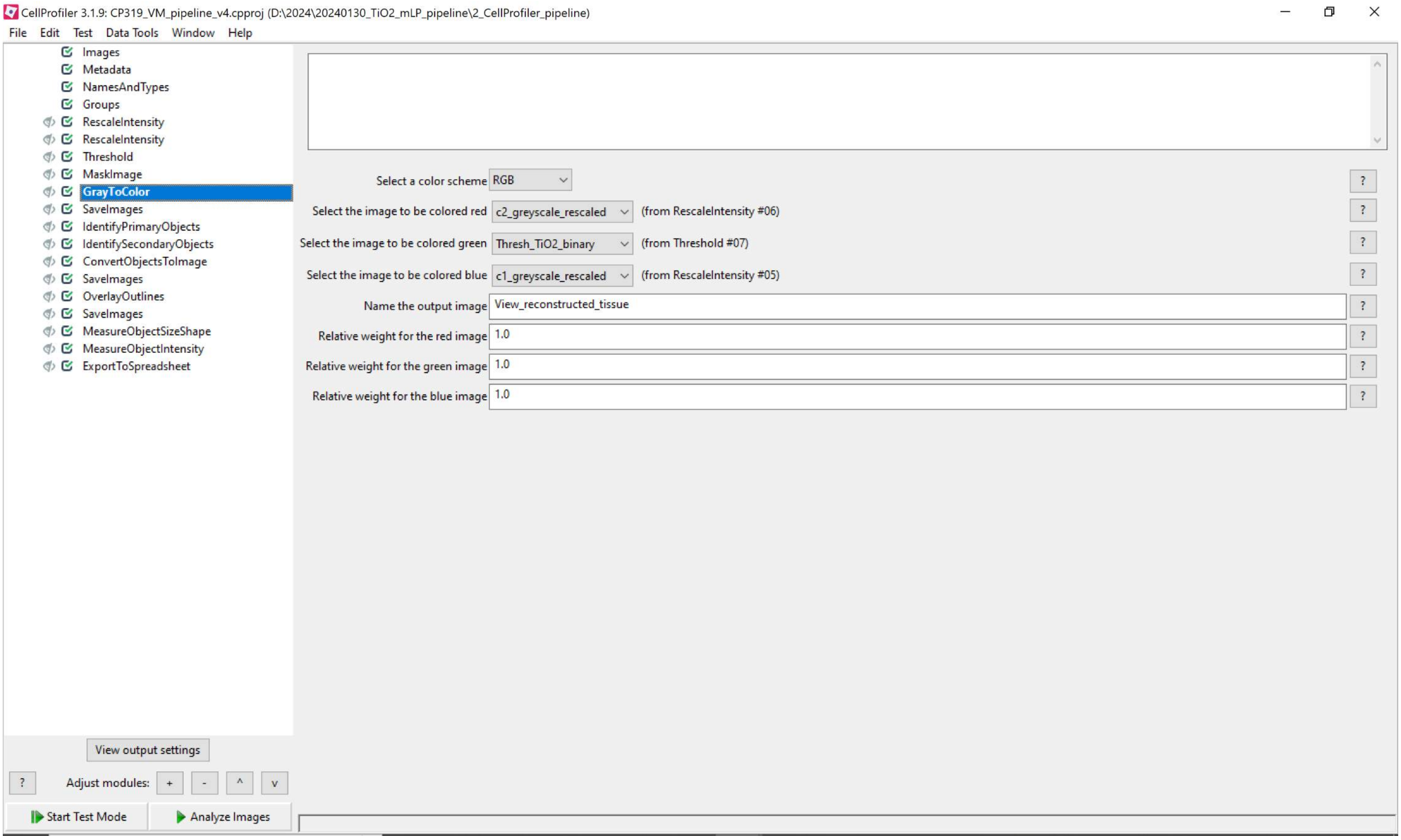
CellProfiler 3.1.9: CP319_VM_pipeline_v4.cpproj (D:\2024\20240130_TiO2_mLP_pipeline\2_CellProfiler_pipeline)
File Edit Test Data Tools Window Help
Images
Metadata
NamesAndTypes
Groups
RescaleIntensity
RescaleIntensity
Threshold
MaskImage
GrayToColor
SaveImages
IdentifyPrimaryObjects
IdentifySecondaryObjects
ConvertObjectsToImage
SaveImages
OverlayOutlines
SaveImages
MeasureObjectSizeShape
MeasureObjectIntensity
ExportToSpreadsheet
Select a color scheme RGB
Select the image to be colored red c2_greyscale_rescaled (from RescaleIntensity #06)
Select the image to be colored green Thresh_TiO2_binary (from Threshold #07)
Select the image to be colored blue c1_greyscale_rescaled (from RescaleIntensity #05)
Name the output image View_reconstructed_tissue
Relative weight for the red image 1.0
Relative weight for the green image 1.0
Relative weight for the blue image 1.0
View output settings
Adjust modules:
Start Test Mode
Analyze Images

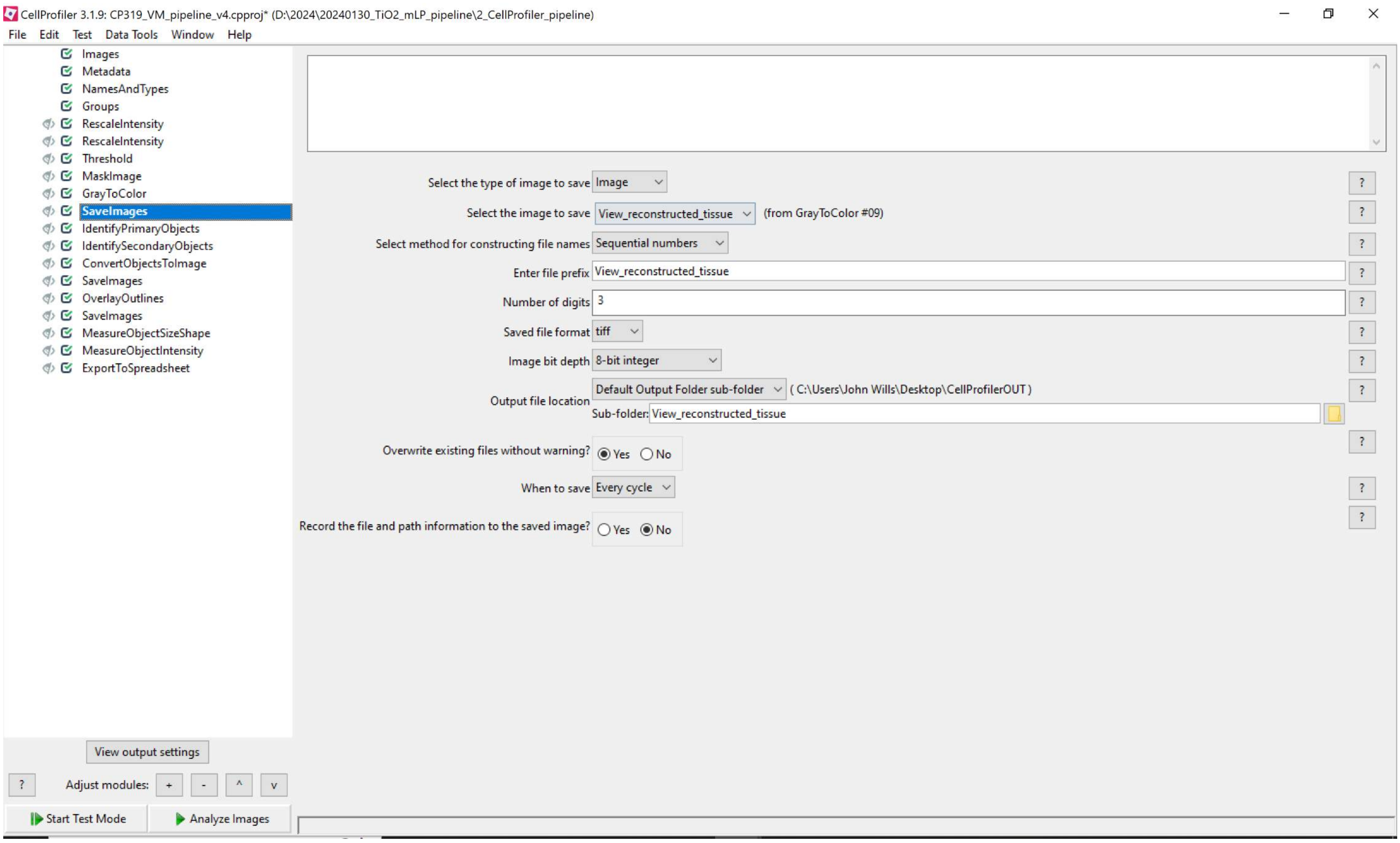
CellProfiler 3.1.9: CP319_VM_pipeline_v4.cpproj* (D:\2024\20240130_TiO2_mLP_pipeline\2_CellProfiler_pipeline)
File Edit Test Data Tools Window Help
Images
Metadata
NamesAndTypes
Groups
RescaleIntensity
RescaleIntensity
Threshold
MaskImage
GrayToColor
SaveImages
IdentifyPrimaryObjects
IdentifySecondaryObjects
ConvertObjectsToImage
SaveImages
OverlayOutlines
SaveImages
MeasureObjectSizeShape
MeasureObjectIntensity
ExportToSpreadsheet
Select the type of image to save
Image
Select the image to save
View_reconstructed_tissue
(from GrayToColor #09)
Select method for constructing file names
Sequential numbers
Enter file prefix
View_reconstructed_tissue
Number of digits
3
Saved file format
tiff
Image bit depth
8-bit integer
Output file location
Default Output Folder sub-folder
( C:\Users\John Wills\Desktop\CellProfilerOUT )
Sub-folder:
View_reconstructed_tissue
Overwrite existing files without warning?
Yes
No
When to save
Every cycle
Record the file and path information to the saved image?
Yes
No
View output settings
Adjust modules:
Start Test Mode
Analyze Images

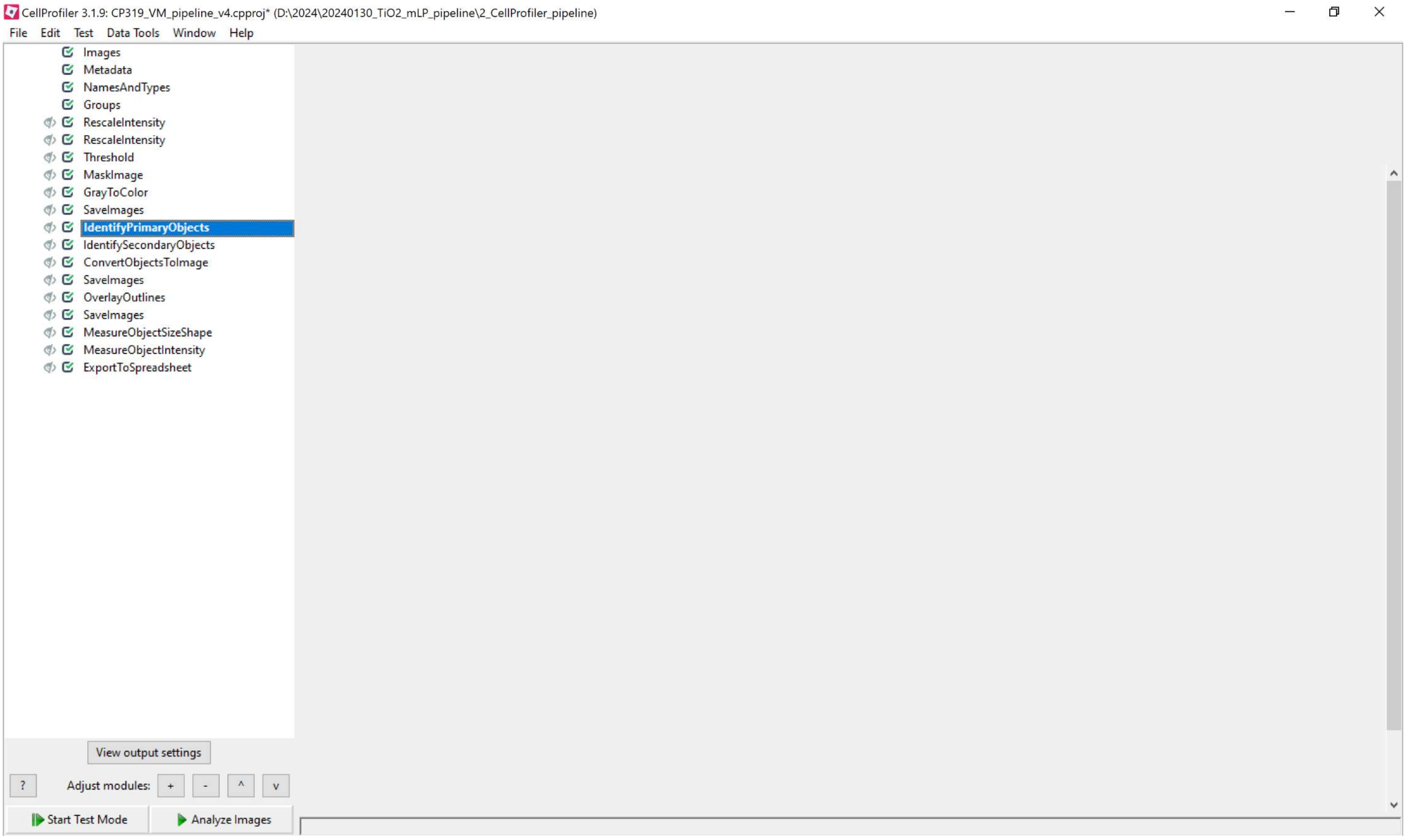
CellProfiler 3.1.9: CP319_VM_pipeline_v4.cpproj* (D:\2024\20240130_TiO2_mLP_pipeline\2_CellProfiler_pipeline)
File Edit Test Data Tools Window Help
Images
Metadata
NamesAndTypes
Groups
RescaleIntensity
RescaleIntensity
Threshold
MaskImage
GrayToColor
SaveImages
IdentifyPrimaryObjects
IdentifySecondaryObjects
ConvertObjectsToImage
SaveImages
OverlayOutlines
SaveImages
MeasureObjectSizeShape
MeasureObjectIntensity
ExportToSpreadsheet
View output settings
?
Adjust modules:
Start Test Mode
Analyze Images

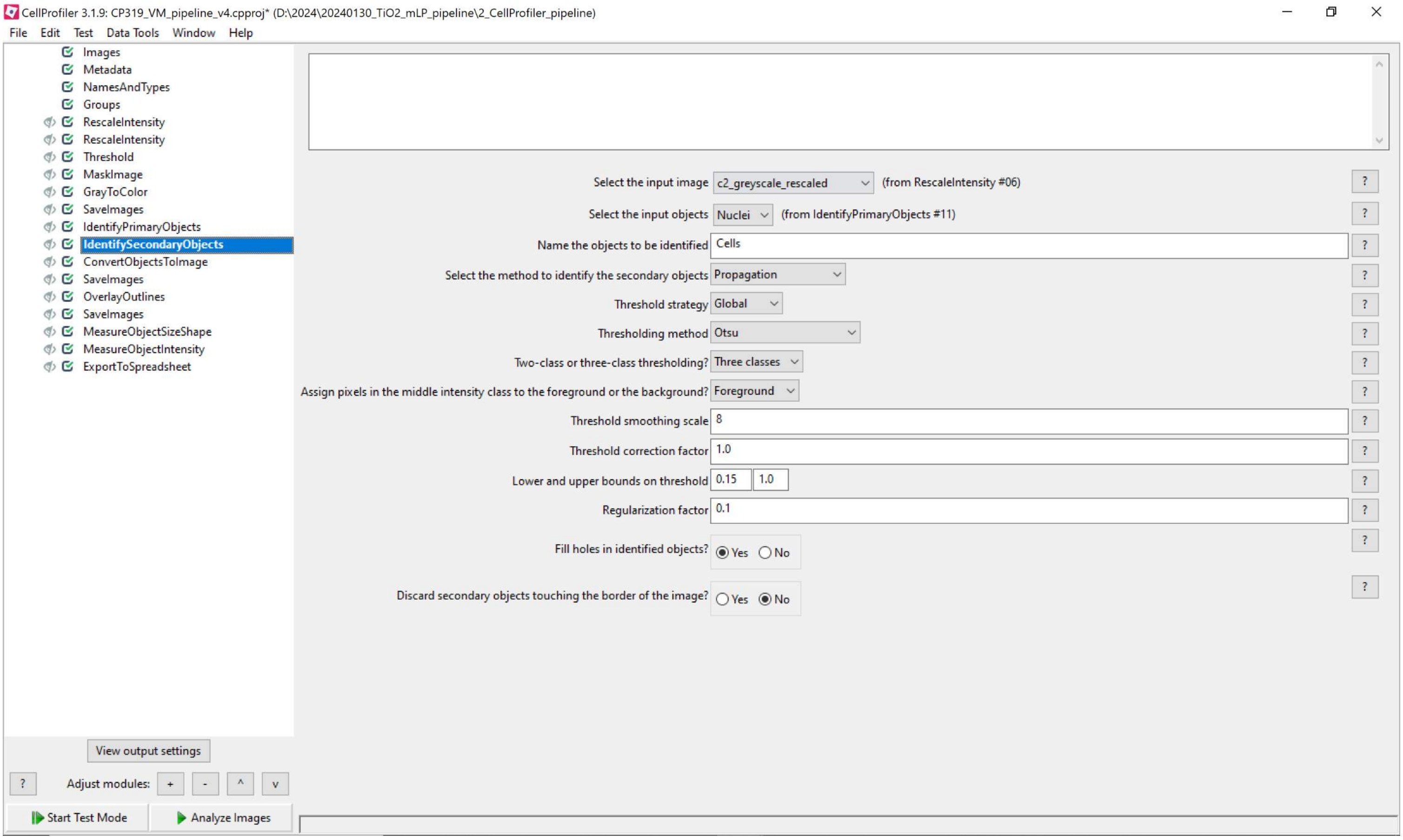
CellProfiler 3.1.9: CP319_VM_pipeline_v4.cpproj* (D:\2024\20240130_TiO2_mLP_pipeline\2_CellProfiler_pipeline)
File Edit Test Data Tools Window Help
Images
Metadata
NamesAndTypes
Groups
RescaleIntensity
RescaleIntensity
Threshold
MaskImage
GrayToColor
SaveImages
IdentifyPrimaryObjects
IdentifySecondaryObjects
ConvertObjectsToImage
SaveImages
OverlayOutlines
SaveImages
MeasureObjectSizeShape
MeasureObjectIntensity
ExportToSpreadsheet
Select the input image c2_greyscale_rescaled (from RescaleIntensity #06)
Select the input objects Nuclei (from IdentifyPrimaryObjects #11)
Name the objects to be identified Cells
Select the method to identify the secondary objects Propagation
Threshold strategy Global
Thresholding method Otsu
Two-class or three-class thresholding? Three classes
Assign pixels in the middle intensity class to the foreground or the background? Foreground
Threshold smoothing scale 8
Threshold correction factor 1.0
Lower and upper bounds on threshold 0.15 1.0
Regularization factor 0.1
Fill holes in identified objects? Yes No
Discard secondary objects touching the border of the image? Yes No
View output settings
Adjust modules: + - ^ v
Start Test Mode
Analyze Images

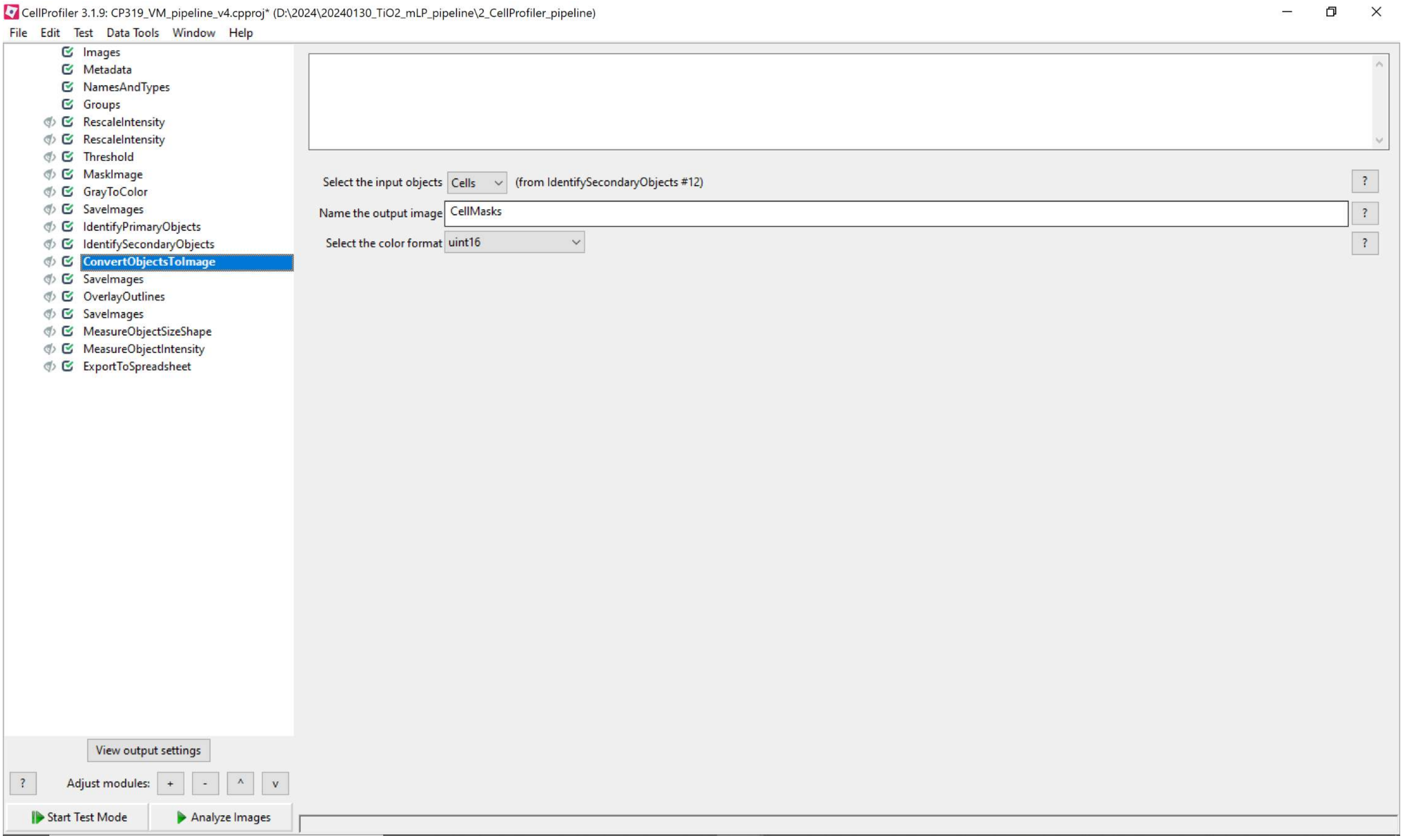
CellProfiler 3.1.9: CP319_VM_pipeline_v4.cpproj* (D:\2024\20240130_TiO2_mLP_pipeline\2_CellProfiler_pipeline)
File Edit Test Data Tools Window Help
Images
Metadata
NamesAndTypes
Groups
RescaleIntensity
RescaleIntensity
Threshold
MaskImage
GrayToColor
SaveImages
IdentifyPrimaryObjects
IdentifySecondaryObjects
ConvertObjectsToImage
SaveImages
OverlayOutlines
SaveImages
MeasureObjectSizeShape
MeasureObjectIntensity
ExportToSpreadsheet
Select the input objects
Cells
(from IdentifySecondaryObjects #12)
Name the output image
CellMasks
Select the color format
uint16
View output settings
Adjust modules:
Start Test Mode
Analyze Images

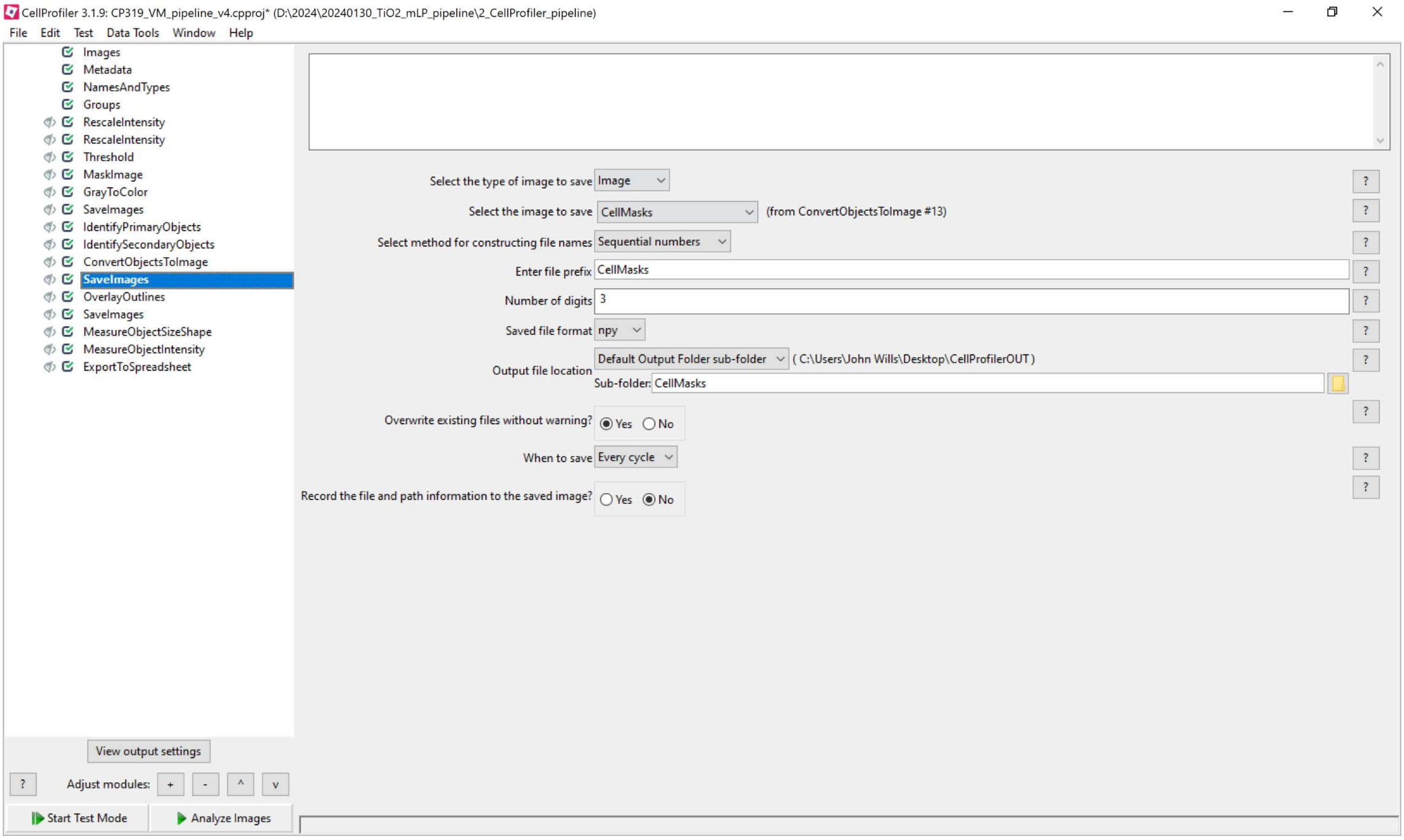
CellProfiler 3.1.9: CP319_VM_pipeline_v4.cpproj* (D:\2024\20240130_TiO2_mLP_pipeline\2_CellProfiler_pipeline)
File Edit Test Data Tools Window Help
Images
Metadata
NamesAndTypes
Groups
RescaleIntensity
RescaleIntensity
Threshold
MaskImage
GrayToColor
SaveImages
IdentifyPrimaryObjects
IdentifySecondaryObjects
ConvertObjectsToImage
SaveImages
OverlayOutlines
SaveImages
MeasureObjectSizeShape
MeasureObjectIntensity
ExportToSpreadsheet
Select the type of image to save Image
Select the image to save CellMasks (from ConvertObjectsToImage #13)
Select method for constructing file names Sequential numbers
Enter file prefix CellMasks
Number of digits 3
Saved file format npy
Output file location Default Output Folder sub-folder ( C:\Users\John Wills\Desktop\CellProfilerOUT )
Sub-folder: CellMasks
Overwrite existing files without warning? Yes No
When to save Every cycle
Record the file and path information to the saved image? Yes No
View output settings
Adjust modules: + - ^ v
Start Test Mode
Analyze Images

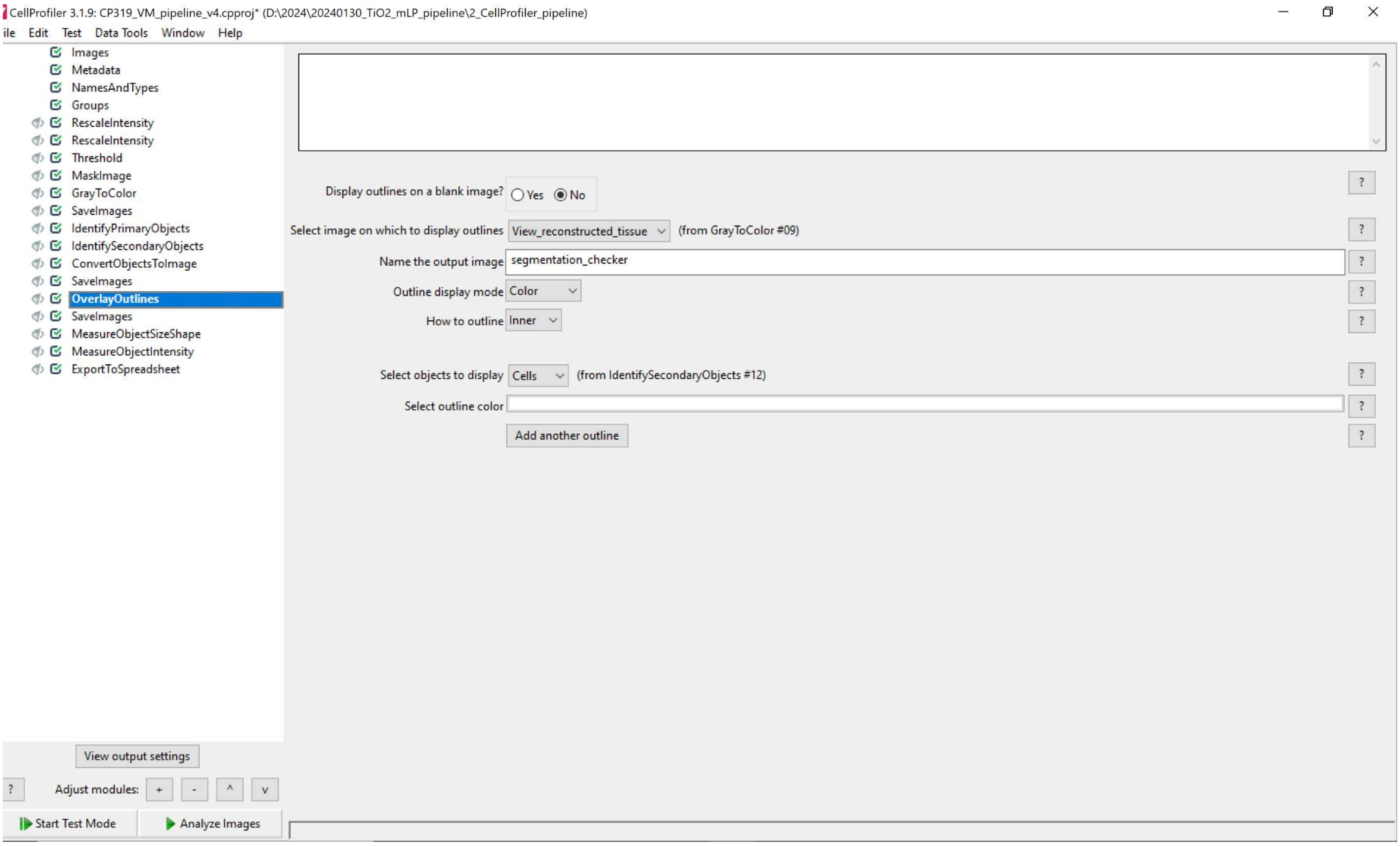
CellProfiler 3.1.9: CP319_VM_pipeline_v4.cpproj* (D:\2024\20240130_TiO2_mLP_pipeline\2_CellProfiler_pipeline)
ile Edit Test Data Tools Window Help
Images
Metadata
NamesAndTypes
Groups
RescaleIntensity
RescaleIntensity
Threshold
MaskImage
GrayToColor
SaveImages
IdentifyPrimaryObjects
IdentifySecondaryObjects
ConvertObjectsToImage
SaveImages
OverlayOutlines
SaveImages
MeasureObjectSizeShape
MeasureObjectIntensity
ExportToSpreadsheet
Display outlines on a blank image?
Yes
No
Select image on which to display outlines
View_reconstructed_tissue
(from GrayToColor #09)
Name the output image
segmentation_checker
Outline display mode
Color
How to outline
Inner
Select objects to display
Cells
(from IdentifySecondaryObjects #12)
Select outline color
Add another outline
View output settings
Adjust modules:
Start Test Mode
Analyze Images

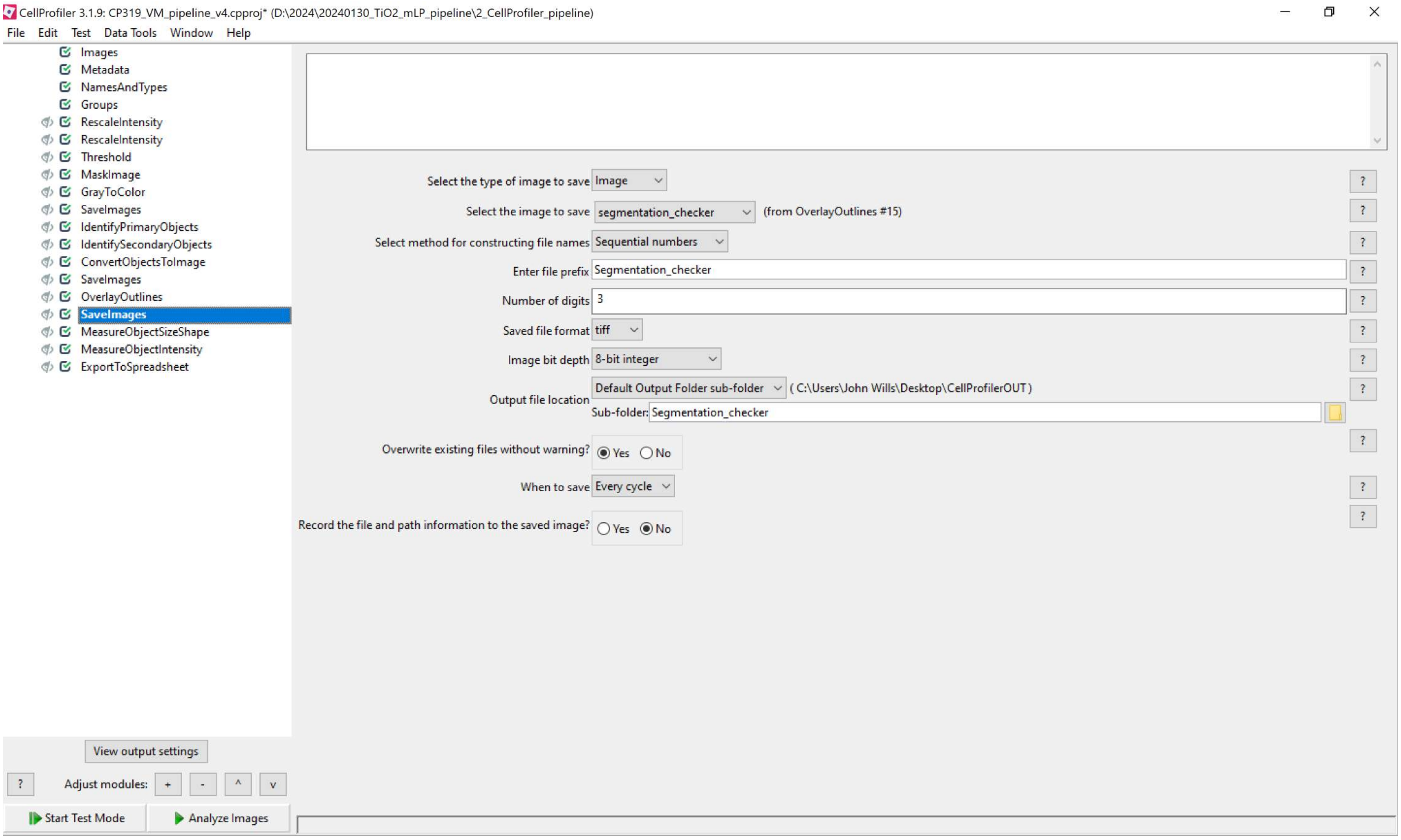
CellProfiler 3.1.9: CP319_VM_pipeline_v4.cpproj* (D:\2024\20240130_TiO2_mLP_pipeline\2_CellProfiler_pipeline)
File Edit Test Data Tools Window Help
Images
Metadata
NamesAndTypes
Groups
RescaleIntensity
RescaleIntensity
Threshold
MaskImage
GrayToColor
SaveImages
IdentifyPrimaryObjects
IdentifySecondaryObjects
ConvertObjectsToImage
SaveImages
OverlayOutlines
SaveImages
MeasureObjectSizeShape
MeasureObjectIntensity
ExportToSpreadsheet
Select the type of image to save Image
Select the image to save segmentation_checker (from OverlayOutlines #15)
Select method for constructing file names Sequential numbers
Enter file prefix Segmentation_checker
Number of digits 3
Saved file format tiff
Image bit depth 8-bit integer
Output file location Default Output Folder sub-folder ( C:\Users\John Wills\Desktop\CellProfilerOUT )
Sub-folder: Segmentation_checker
Overwrite existing files without warning? Yes No
When to save Every cycle
Record the file and path information to the saved image? Yes No
View output settings
Adjust modules: + - ^ v
Start Test Mode
Analyze Images

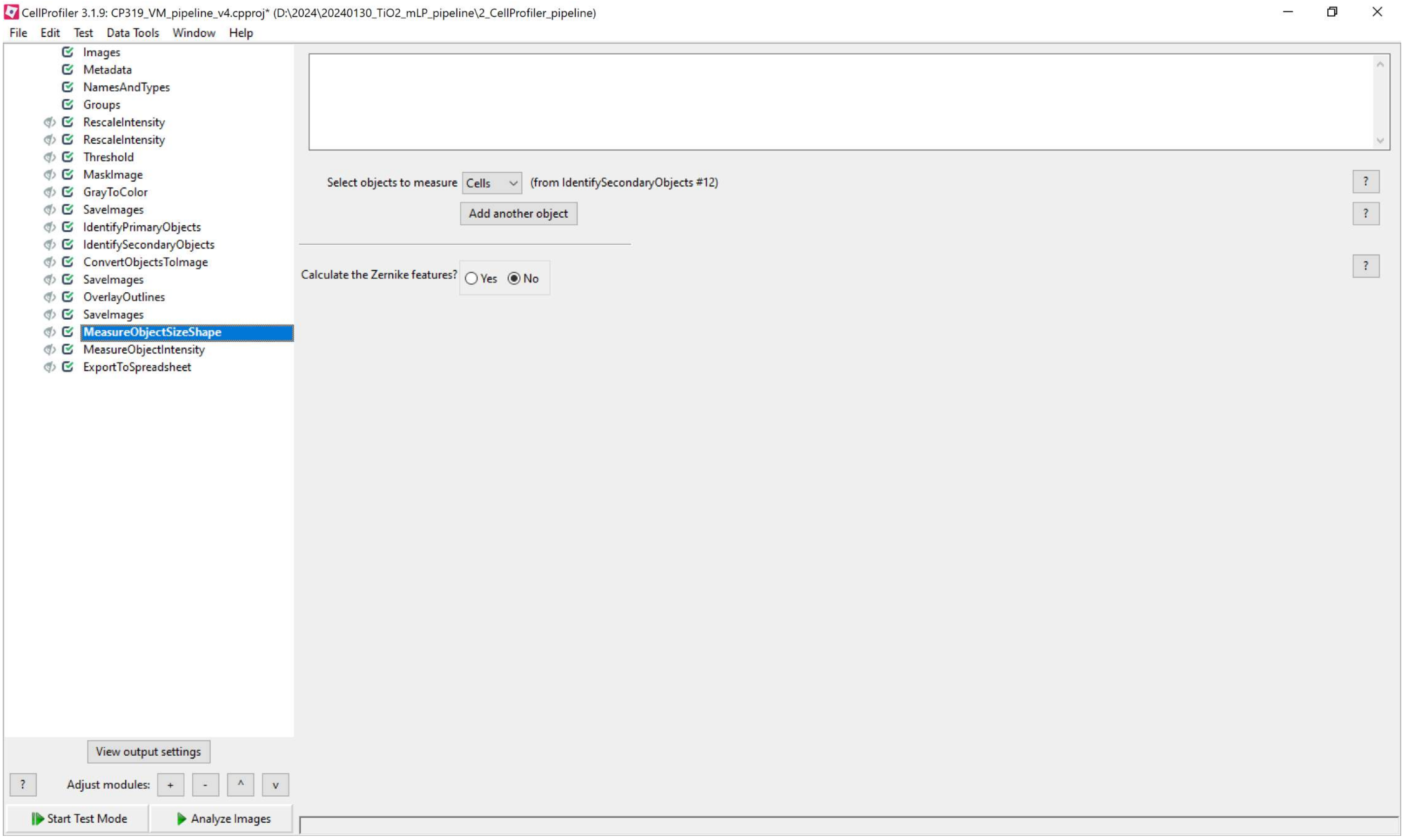
CellProfiler 3.1.9: CP319_VM_pipeline_v4.cpproj* (D:\2024\20240130_TiO2_mLP_pipeline\2_CellProfiler_pipeline)
File Edit Test Data Tools Window Help
Images
Metadata
NamesAndTypes
Groups
RescaleIntensity
RescaleIntensity
Threshold
MaskImage
GrayToColor
SaveImages
IdentifyPrimaryObjects
IdentifySecondaryObjects
ConvertObjectsToImage
SaveImages
OverlayOutlines
SaveImages
MeasureObjectSizeShape
MeasureObjectIntensity
ExportToSpreadsheet
Select objects to measure
Cells
(from IdentifySecondaryObjects #12)
Add another object
Calculate the Zernike features?
Yes
No
View output settings
Adjust modules:
Start Test Mode
Analyze Images

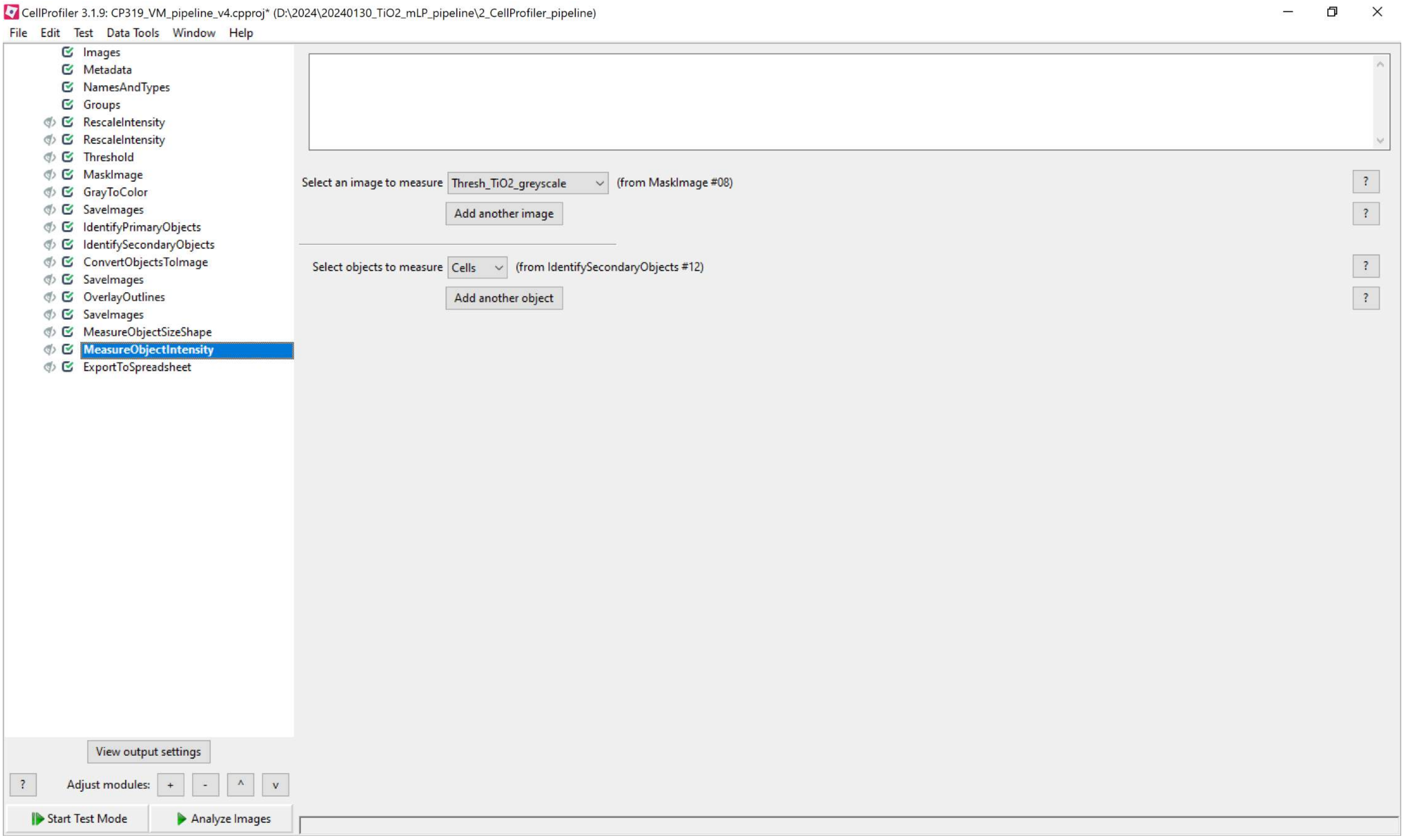
CellProfiler 3.1.9: CP319_VM_pipeline_v4.cpproj* (D:\2024\20240130_TiO2_mLP_pipeline\2_CellProfiler_pipeline)
File Edit Test Data Tools Window Help
Images
Metadata
NamesAndTypes
Groups
RescaleIntensity
RescaleIntensity
Threshold
MaskImage
GrayToColor
SaveImages
IdentifyPrimaryObjects
IdentifySecondaryObjects
ConvertObjectsToImage
SaveImages
OverlayOutlines
SaveImages
MeasureObjectSizeShape
MeasureObjectIntensity
ExportToSpreadsheet
Select an image to measure
Thresh_TiO2_greyscale
(from MaskImage #08)
Add another image
Select objects to measure
Cells
(from IdentifySecondaryObjects #12)
Add another object
View output settings
Adjust modules:
Start Test Mode
Analyze Images

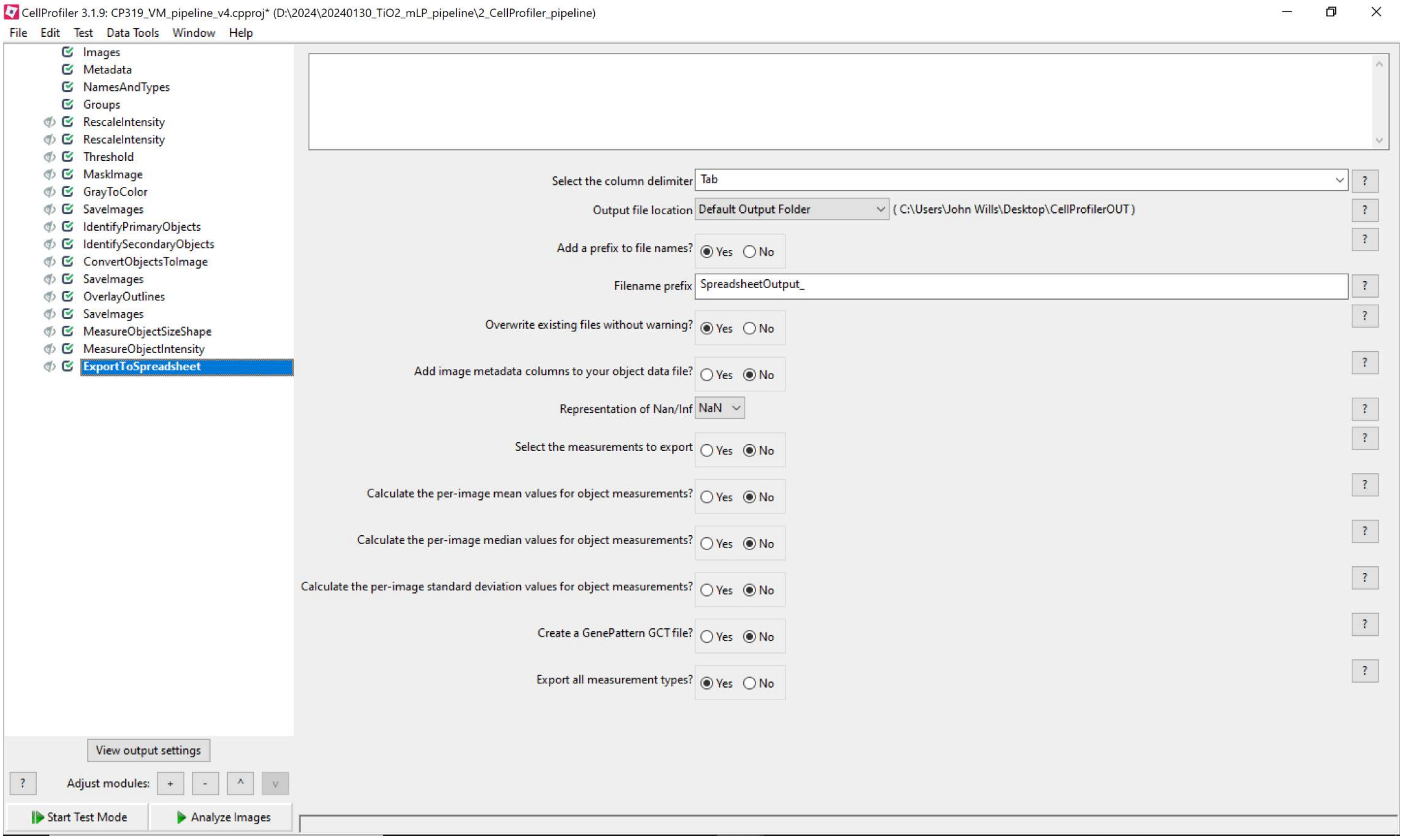
CellProfiler 3.1.9: CP319_VM_pipeline_v4.cpproj* (D:\2024\20240130_TiO2_mLP_pipeline\2_CellProfiler_pipeline)
File Edit Test Data Tools Window Help
Images
Metadata
NamesAndTypes
Groups
RescaleIntensity
RescaleIntensity
Threshold
MaskImage
GrayToColor
SaveImages
IdentifyPrimaryObjects
IdentifySecondaryObjects
ConvertObjectsToImage
SaveImages
OverlayOutlines
SaveImages
MeasureObjectSizeShape
MeasureObjectIntensity
ExportToSpreadsheet
Select the column delimiter Tab
Output file location Default Output Folder ( C:\Users\John Wills\Desktop\CellProfilerOUT )
Add a prefix to file names? Yes No
Filename prefix SpreadsheetOutput_
Overwrite existing files without warning? Yes No
Add image metadata columns to your object data file? Yes No
Representation of Nan/Inf NaN
Select the measurements to export Yes No
Calculate the per-image mean values for object measurements? Yes No
Calculate the per-image median values for object measurements? Yes No
Calculate the per-image standard deviation values for object measurements? Yes No
Create a GenePattern GCT file? Yes No
Export all measurement types? Yes No
View output settings
Adjust modules: + - ^ v
Start Test Mode
Analyze Images

**Supplementary Note 2 – Mouse lymphoid tissue image analysis pipeline using UNET-generated probability maps:** This section presents screenshots of the CellProfiler image analysis pipeline used to measure per-cell reflectance (fgTi$O_2$) and immunofluorescence intensities (here, PD-L1 quantification from **Figure 4** is shown) and cell shape/size features. The pipeline and sample image data are available for download from the BioStudies archive accompanying the paper.

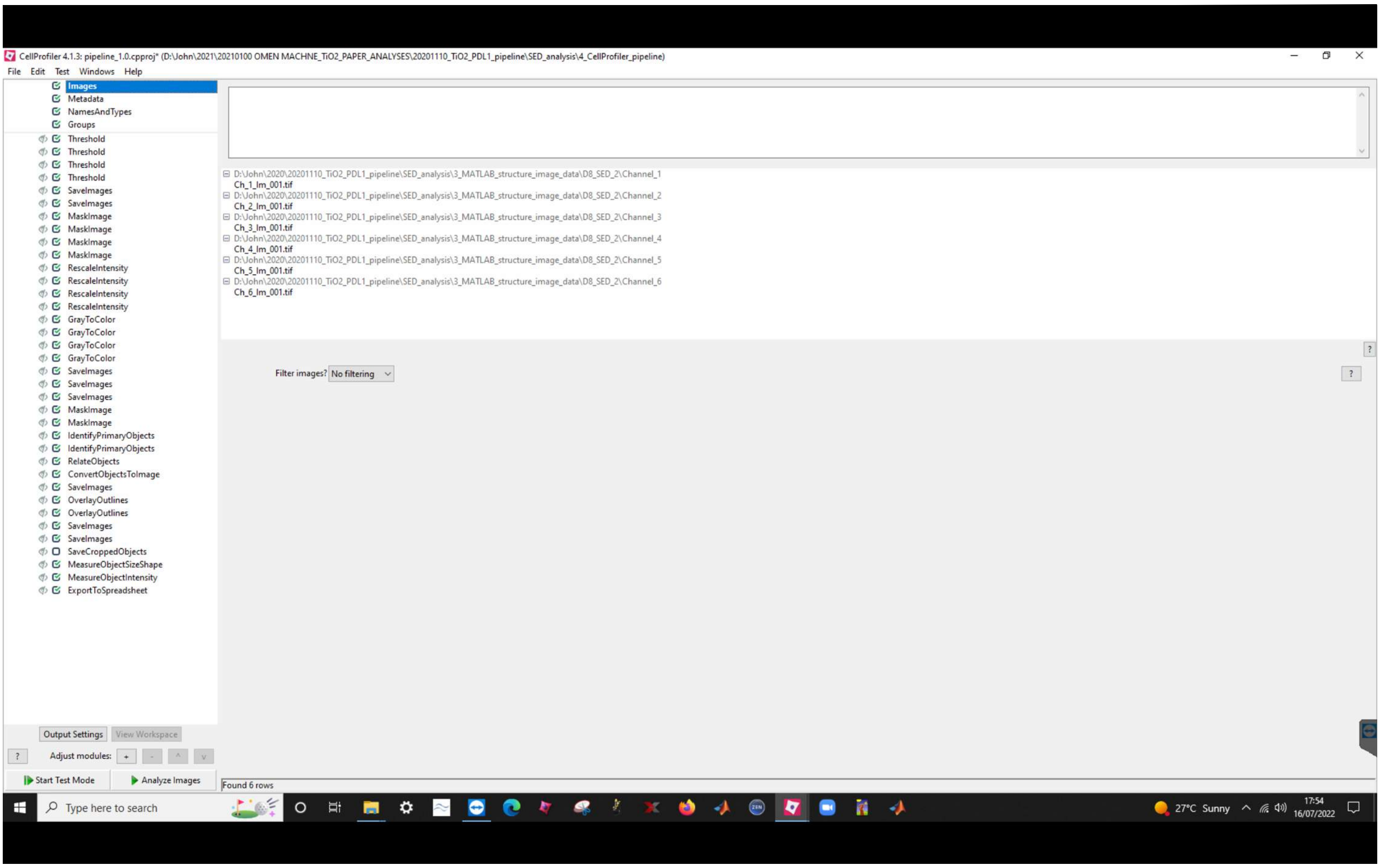

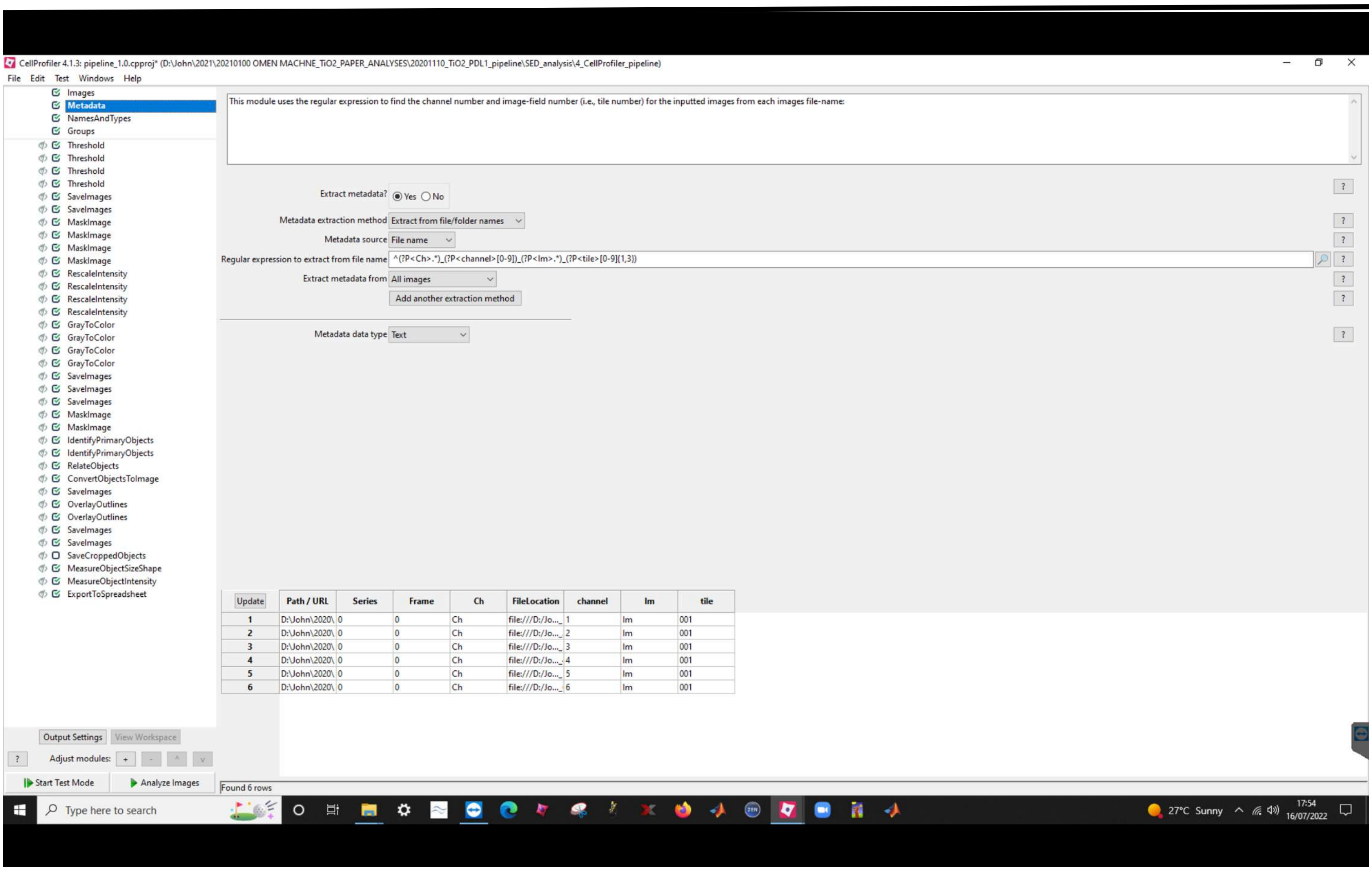


| | Path / URL | Series | Frame | Ch | FileLocation | channel | Im | tile |
|---|---|---|---|---|---|---|---|---|
| 1 | D:\John\2020\ | 0 | 0 | Ch | file:///D:/Jo... | 1 | Im | 001 |
| 2 | D:\John\2020\ | 0 | 0 | Ch | file:///D:/Jo... | 2 | Im | 001 |
| 3 | D:\John\2020\ | 0 | 0 | Ch | file:///D:/Jo... | 3 | Im | 001 |
| 4 | D:\John\2020\ | 0 | 0 | Ch | file:///D:/Jo... | 4 | Im | 001 |
| 5 | D:\John\2020\ | 0 | 0 | Ch | file:///D:/Jo... | 5 | Im | 001 |
| 6 | D:\John\2020\ | 0 | 0 | Ch | file:///D:/Jo... | 6 | Im | 001 |

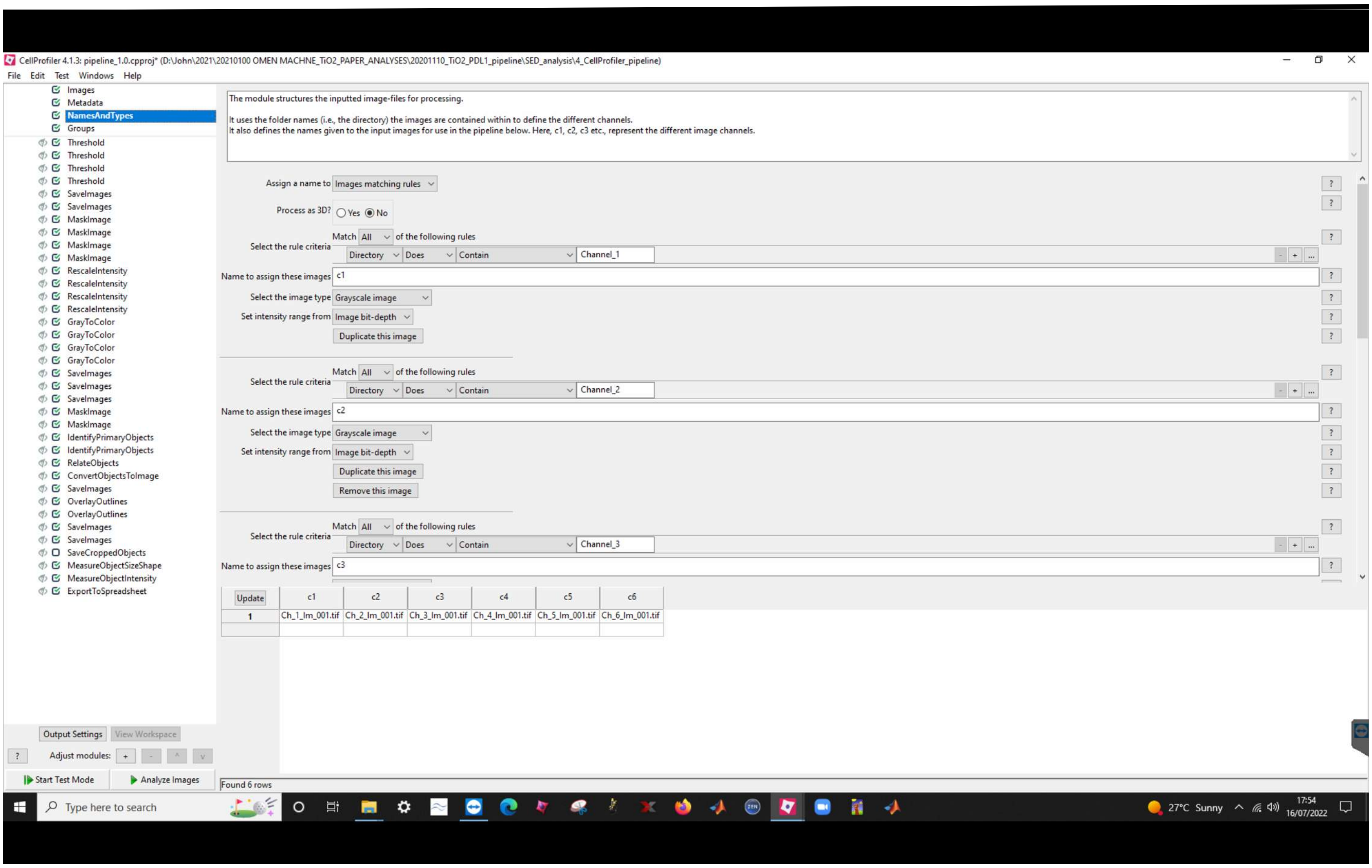
CellProfiler 4.1.3: pipeline_1.0.cpproj* (D:\John\2021\20210100 OMEN MACHNE_TiO2_PAPER_ANALYSES\20201110_TiO2_PDL1_pipeline\SED_analysis\4_CellProfiler_pipeline)
File Edit Test Windows Help
Images
Metadata
NamesAndTypes
Groups
Threshold
SaveImages
MaskImage
RescaleIntensity
GrayToColor
IdentifyPrimaryObjects
RelateObjects
ConvertObjectsToImage
OverlayOutlines
SaveCroppedObjects
MeasureObjectSizeShape
MeasureObjectIntensity
ExportToSpreadsheet
The module structures the inputted image-files for processing.
It uses the folder names (i.e., the directory) the images are contained within to define the different channels.
It also defines the names given to the input images for use in the pipeline below. Here, c1, c2, c3 etc., represent the different image channels.
Assign a name to Images matching rules
Process as 3D? Yes No
Match All of the following rules
Select the rule criteria Directory Does Contain Channel_1
Name to assign these images c1
Select the image type Grayscale image
Set intensity range from Image bit-depth
Duplicate this image
Select the rule criteria Directory Does Contain Channel_2
Name to assign these images c2
Remove this image
Select the rule criteria Directory Does Contain Channel_3
Name to assign these images c3
Update c1 c2 c3 c4 c5 c6
1 Ch_1_Im_001.tif Ch_2_Im_001.tif Ch_3_Im_001.tif Ch_4_Im_001.tif Ch_5_Im_001.tif Ch_6_Im_001.tif
Output Settings View Workspace
Adjust modules:
Start Test Mode
Analyze Images
Found 6 rows
Type here to search
27°C Sunny
17:54
16/07/2022

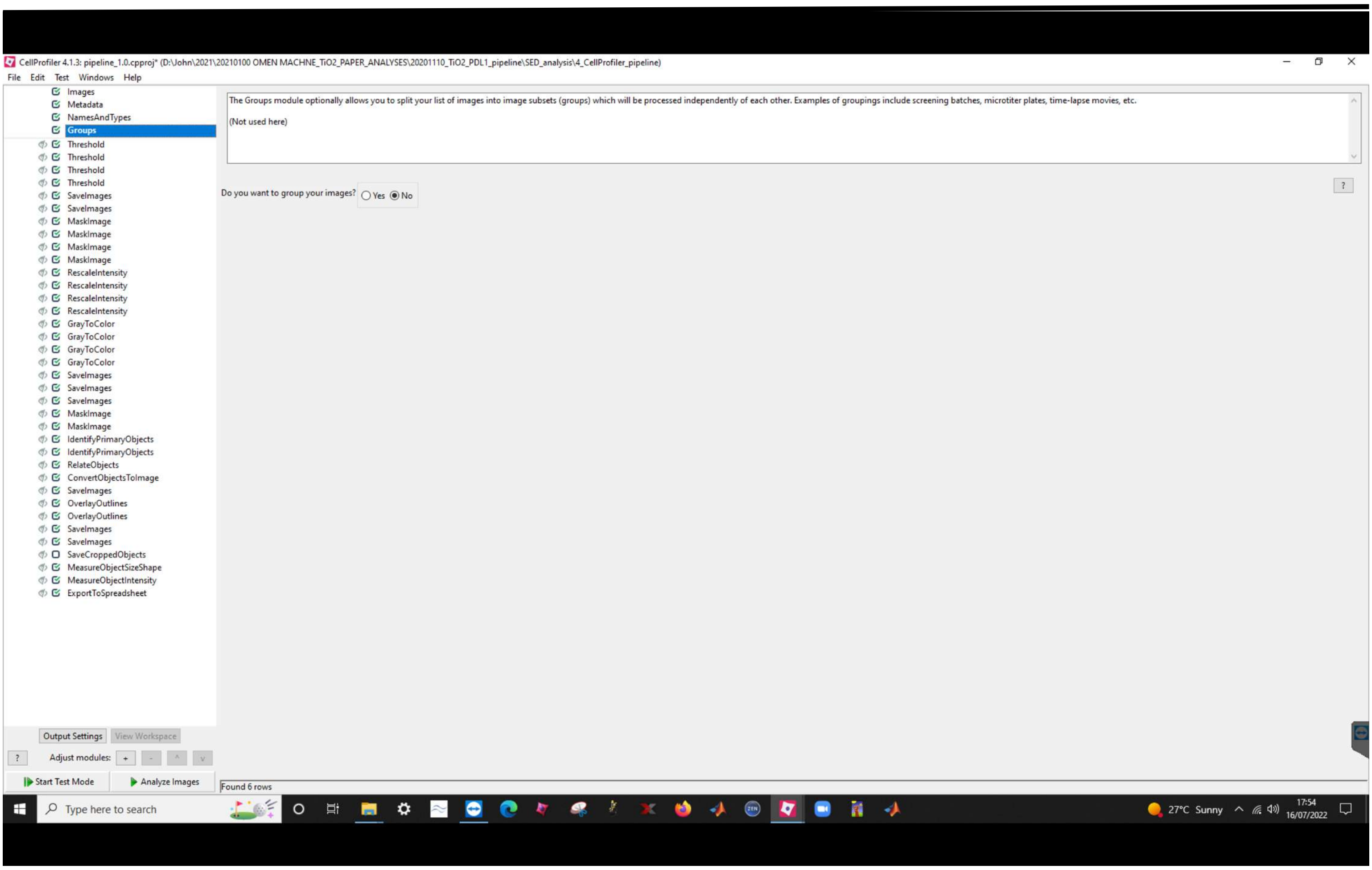
CellProfiler 4.1.3: pipeline_1.0.cpproj* (D:\John\2021\20210100 OMEN MACHNE_TiO2_PAPER_ANALYSES\20201110_TiO2_PDL1_pipeline\SED_analysis\4_CellProfiler_pipeline)
File Edit Test Windows Help
Images
Metadata
NamesAndTypes
Groups
Threshold
SaveImages
MaskImage
RescaleIntensity
GrayToColor
IdentifyPrimaryObjects
RelateObjects
ConvertObjectsToImage
OverlayOutlines
SaveCroppedObjects
MeasureObjectSizeShape
MeasureObjectIntensity
ExportToSpreadsheet
The Groups module optionally allows you to split your list of images into image subsets (groups) which will be processed independently of each other. Examples of groupings include screening batches, microtiter plates, time-lapse movies, etc.
(Not used here)
Do you want to group your images? Yes No
Output Settings
View Workspace
Adjust modules:
Start Test Mode
Analyze Images
Found 6 rows
Type here to search
27°C Sunny
17:54
16/07/2022

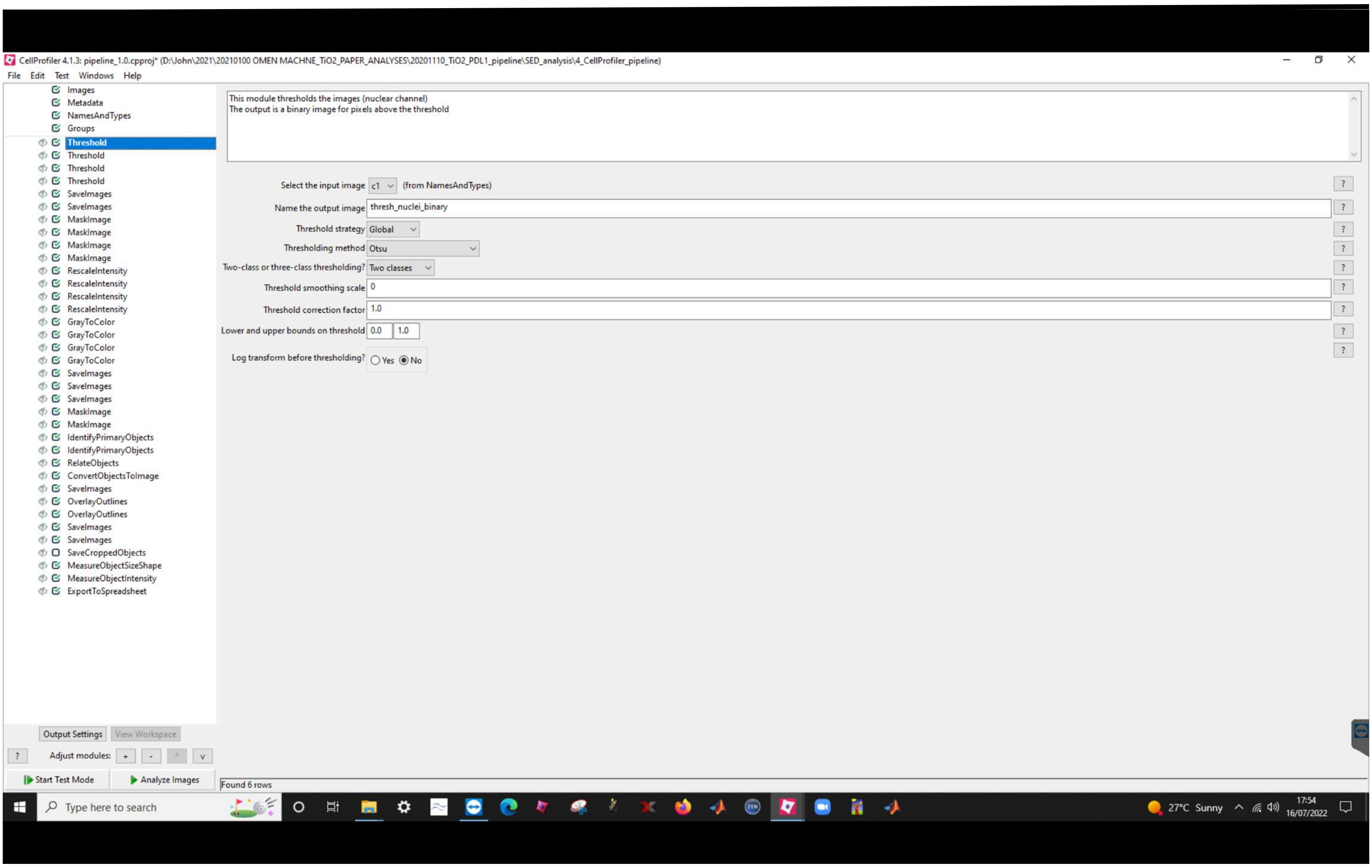
CellProfiler 4.1.3: pipeline_1.0.cpproj* (D:\John\2021\20210100 OMEN MACHNE_TiO2_PAPER_ANALYSES\20201110_TiO2_PDL1_pipeline\SED_analysis\4_CellProfiler_pipeline)
File Edit Test Windows Help
Images
Metadata
NamesAndTypes
Groups
Threshold
Threshold
Threshold
Threshold
SaveImages
SaveImages
MaskImage
MaskImage
MaskImage
MaskImage
RescaleIntensity
RescaleIntensity
RescaleIntensity
RescaleIntensity
GrayToColor
GrayToColor
GrayToColor
GrayToColor
SaveImages
SaveImages
SaveImages
MaskImage
MaskImage
IdentifyPrimaryObjects
IdentifyPrimaryObjects
RelateObjects
ConvertObjectsToImage
SaveImages
OverlayOutlines
OverlayOutlines
SaveImages
SaveImages
SaveCroppedObjects
MeasureObjectSizeShape
MeasureObjectIntensity
ExportToSpreadsheet
This module thresholds the images (nuclear channel)
The output is a binary image for pixels above the threshold
Select the input image c1 (from NamesAndTypes)
Name the output image thresh_nuclei_binary
Threshold strategy Global
Thresholding method Otsu
Two-class or three-class thresholding? Two classes
Threshold smoothing scale 0
Threshold correction factor 1.0
Lower and upper bounds on threshold 0.0 1.0
Log transform before thresholding? Yes No
Output Settings
View Workspace
Adjust modules: + - ^ v
Start Test Mode
Analyze Images
Found 6 rows
Type here to search
27°C Sunny
17:54
16/07/2022

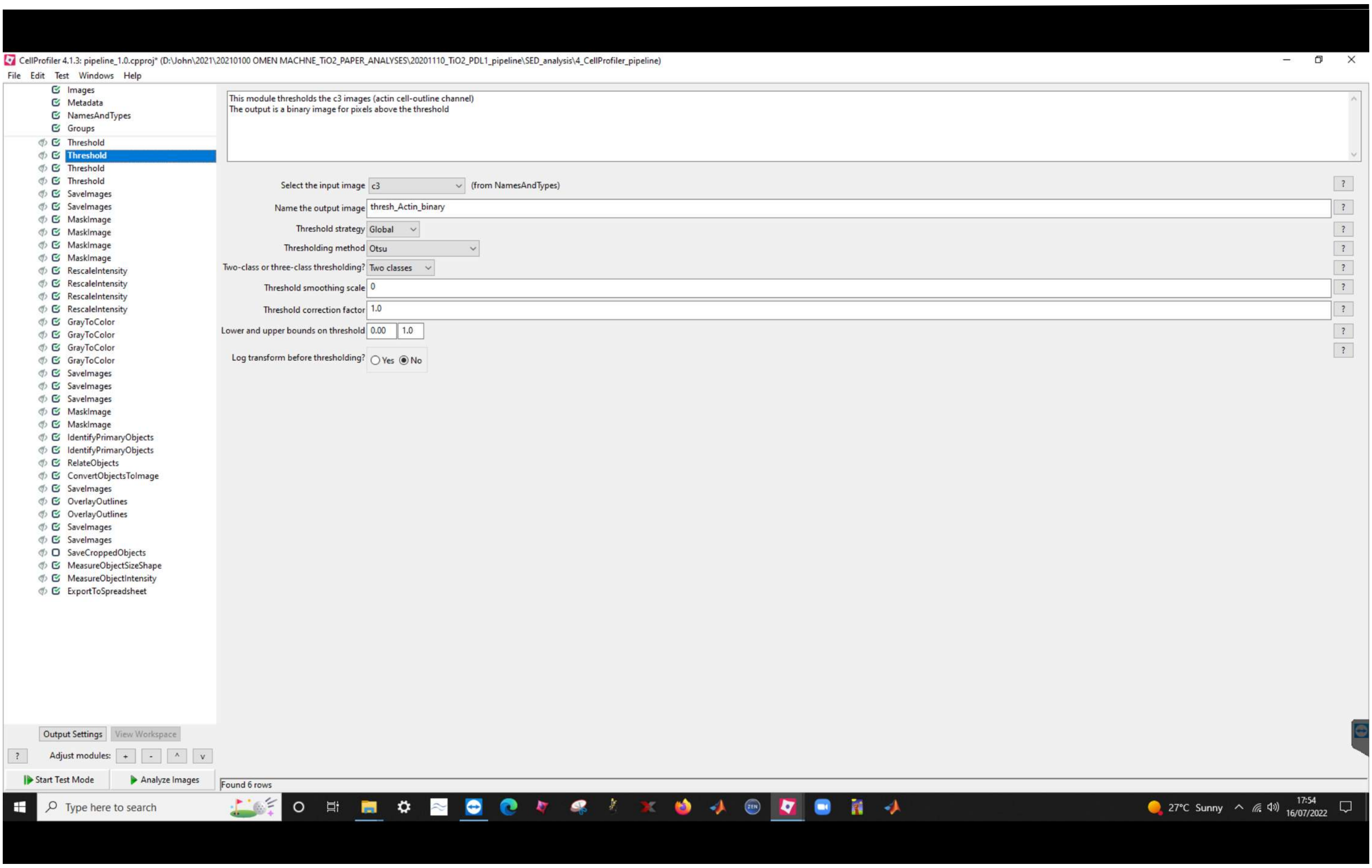
CellProfiler 4.1.3: pipeline_1.0.cpproj* (D:\John\2021\20210100 OMEN MACHNE_TiO2_PAPER_ANALYSES\20201110_TiO2_PDL1_pipeline\SED_analysis\4_CellProfiler_pipeline)
File Edit Test Windows Help
Images
Metadata
NamesAndTypes
Groups
Threshold
Threshold
Threshold
Threshold
SaveImages
SaveImages
MaskImage
MaskImage
MaskImage
MaskImage
RescaleIntensity
RescaleIntensity
RescaleIntensity
RescaleIntensity
GrayToColor
GrayToColor
GrayToColor
GrayToColor
SaveImages
SaveImages
SaveImages
MaskImage
MaskImage
IdentifyPrimaryObjects
IdentifyPrimaryObjects
RelateObjects
ConvertObjectsToImage
SaveImages
OverlayOutlines
OverlayOutlines
SaveImages
SaveImages
SaveCroppedObjects
MeasureObjectSizeShape
MeasureObjectIntensity
ExportToSpreadsheet
This module thresholds the c3 images (actin cell-outline channel)
The output is a binary image for pixels above the threshold
Select the input image c3 (from NamesAndTypes)
Name the output image thresh_Actin_binary
Threshold strategy Global
Thresholding method Otsu
Two-class or three-class thresholding? Two classes
Threshold smoothing scale 0
Threshold correction factor 1.0
Lower and upper bounds on threshold 0.00 1.0
Log transform before thresholding? Yes No
Output Settings
View Workspace
Adjust modules: + - ^ v
Start Test Mode
Analyze Images
Found 6 rows
Type here to search
27°C Sunny
17:54
16/07/2022

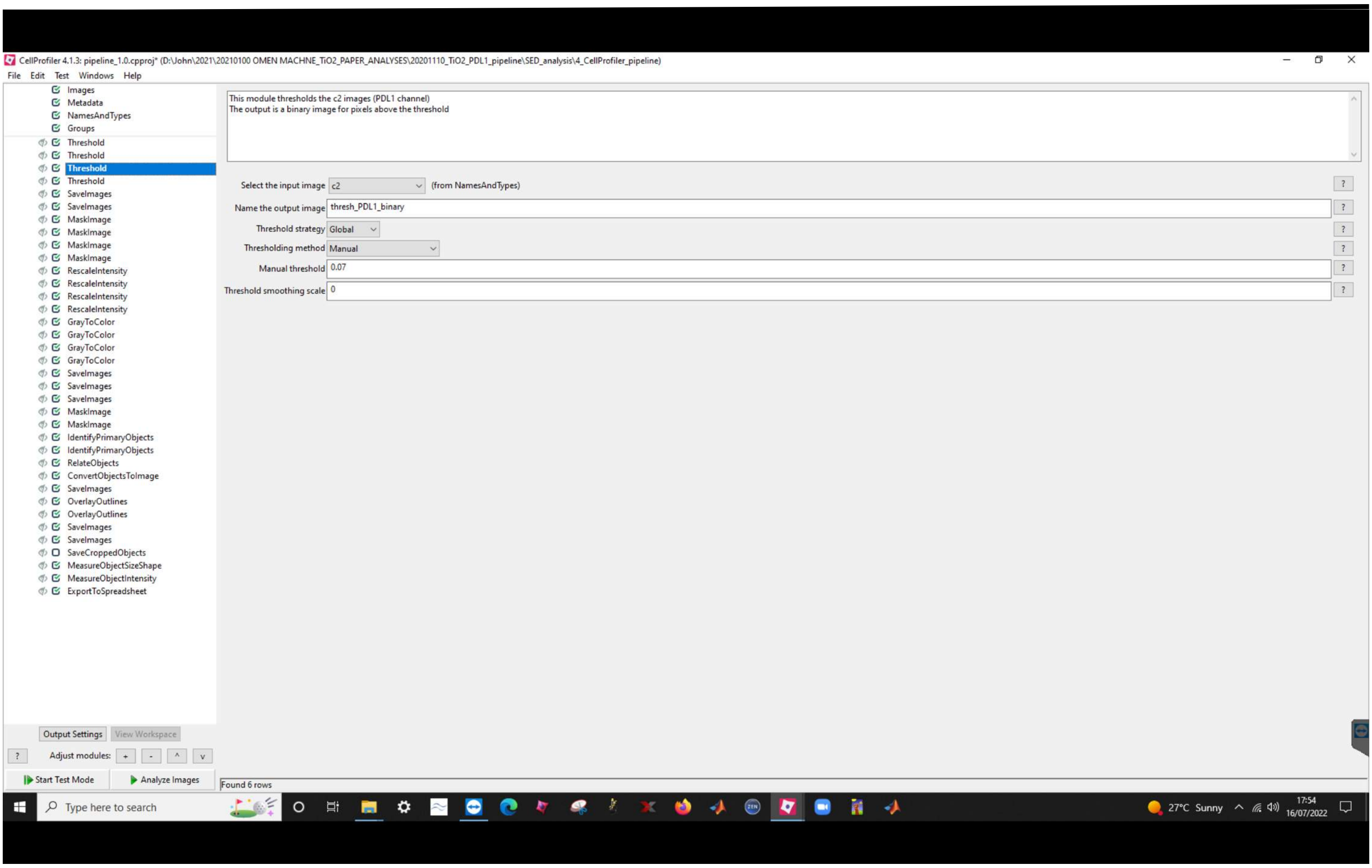
CellProfiler 4.1.3: pipeline_1.0.cpproj* (D:\John\2021\20210100 OMEN MACHNE_TiO2_PAPER_ANALYSES\20201110_TiO2_PDL1_pipeline\SED_analysis\4_CellProfiler_pipeline)
File Edit Test Windows Help
Images
Metadata
NamesAndTypes
Groups
Threshold
Threshold
Threshold
Threshold
SaveImages
SaveImages
MaskImage
MaskImage
MaskImage
MaskImage
RescaleIntensity
RescaleIntensity
RescaleIntensity
RescaleIntensity
GrayToColor
GrayToColor
GrayToColor
GrayToColor
SaveImages
SaveImages
SaveImages
MaskImage
MaskImage
IdentifyPrimaryObjects
IdentifyPrimaryObjects
RelateObjects
ConvertObjectsToImage
SaveImages
OverlayOutlines
OverlayOutlines
SaveImages
SaveImages
SaveCroppedObjects
MeasureObjectSizeShape
MeasureObjectIntensity
ExportToSpreadsheet
This module thresholds the c2 images (PDL1 channel)
The output is a binary image for pixels above the threshold
Select the input image c2 (from NamesAndTypes)
Name the output image thresh_PDL1_binary
Threshold strategy Global
Thresholding method Manual
Manual threshold 0.07
Threshold smoothing scale 0
Output Settings
View Workspace
Adjust modules: + - ^ v
Start Test Mode
Analyze Images
Found 6 rows
Type here to search
27°C Sunny
17:54
16/07/2022

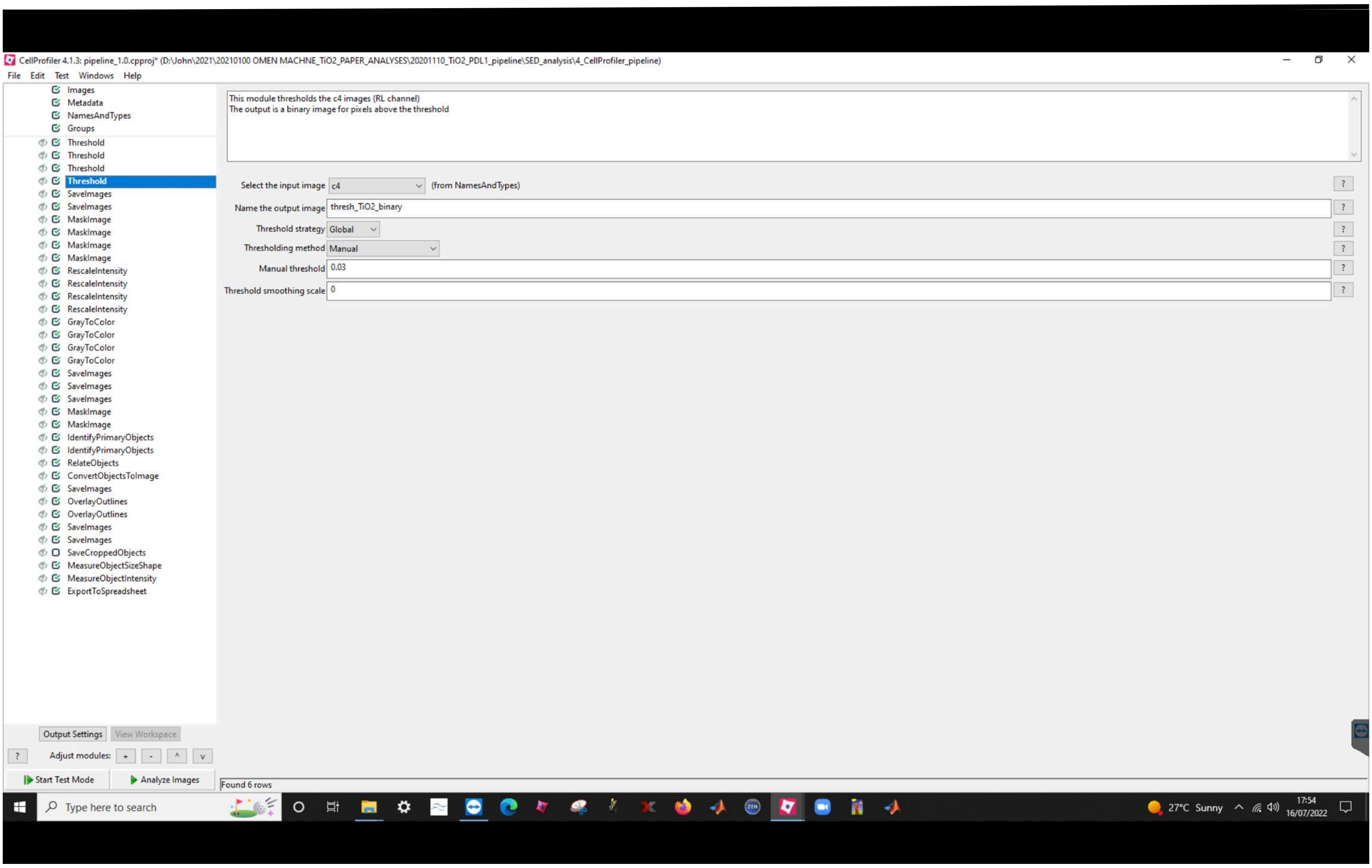
CellProfiler 4.1.3: pipeline_1.0.cpproj* (D:\John\2021\20210100 OMEN MACHNE_TiO2_PAPER_ANALYSES\20201110_TiO2_PDL1_pipeline\SED_analysis\4_CellProfiler_pipeline)
File Edit Test Windows Help
Images
Metadata
NamesAndTypes
Groups
Threshold
Threshold
Threshold
Threshold
SaveImages
SaveImages
MaskImage
MaskImage
MaskImage
MaskImage
RescaleIntensity
RescaleIntensity
RescaleIntensity
RescaleIntensity
GrayToColor
GrayToColor
GrayToColor
GrayToColor
SaveImages
SaveImages
SaveImages
MaskImage
MaskImage
IdentifyPrimaryObjects
IdentifyPrimaryObjects
RelateObjects
ConvertObjectsToImage
SaveImages
OverlayOutlines
OverlayOutlines
SaveImages
SaveImages
SaveCroppedObjects
MeasureObjectSizeShape
MeasureObjectIntensity
ExportToSpreadsheet
This module thresholds the c4 images (RL channel)
The output is a binary image for pixels above the threshold
Select the input image c4 (from NamesAndTypes)
Name the output image thresh_TiO2_binary
Threshold strategy Global
Thresholding method Manual
Manual threshold 0.03
Threshold smoothing scale 0
Output Settings
View Workspace
Adjust modules:
Start Test Mode
Analyze Images
Found 6 rows
Type here to search
27°C Sunny
17:54
16/07/2022

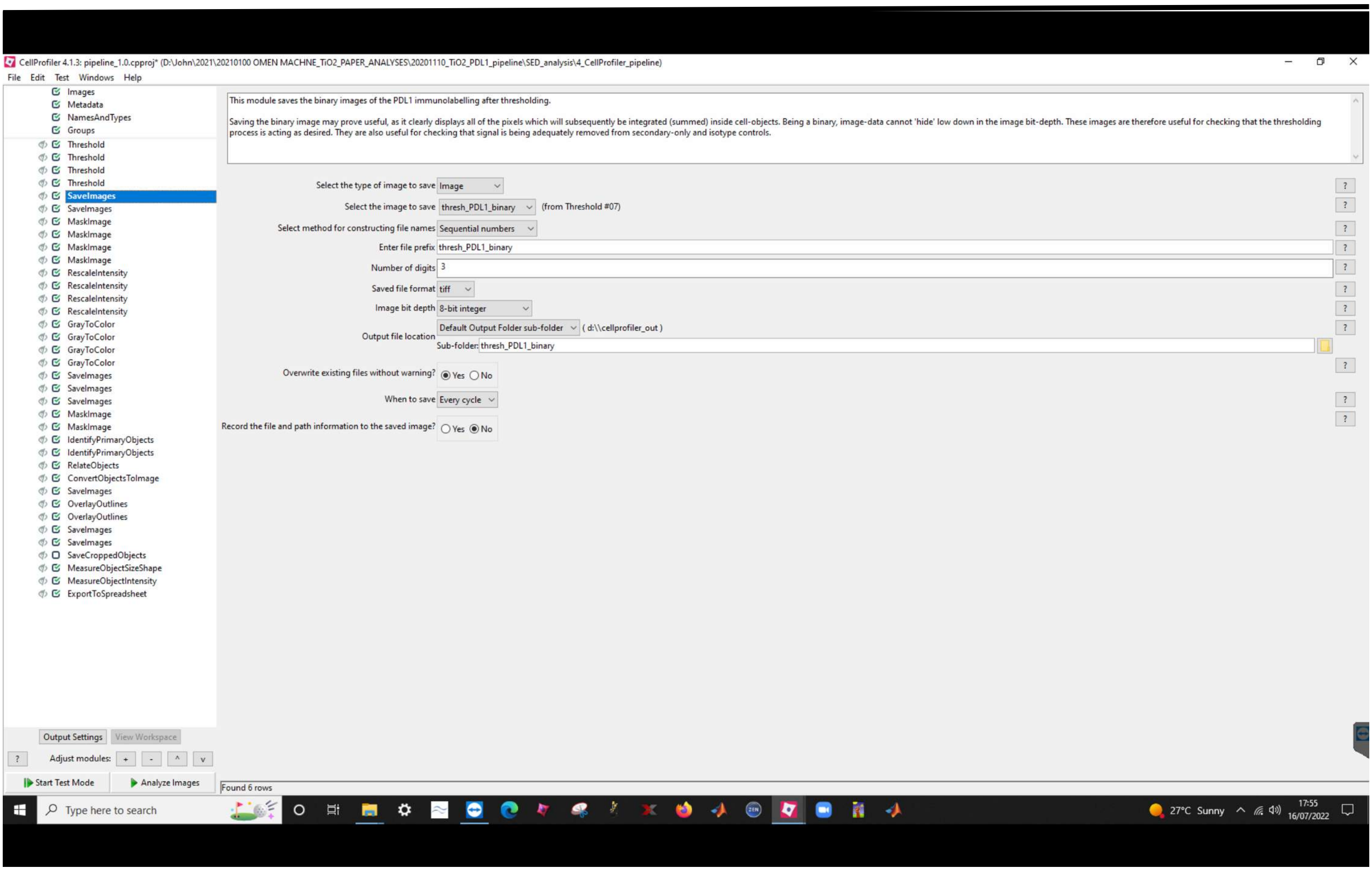
CellProfiler 4.1.3: pipeline_1.0.cpproj* (D:\John\2021\20210100 OMEN MACHNE_TiO2_PAPER_ANALYSES\20201110_TiO2_PDL1_pipeline\SED_analysis\4_CellProfiler_pipeline)
File Edit Test Windows Help
Images
Metadata
NamesAndTypes
Groups
Threshold
Threshold
Threshold
Threshold
SaveImages
SaveImages
MaskImage
MaskImage
MaskImage
MaskImage
RescaleIntensity
RescaleIntensity
RescaleIntensity
RescaleIntensity
GrayToColor
GrayToColor
GrayToColor
GrayToColor
SaveImages
SaveImages
SaveImages
MaskImage
MaskImage
IdentifyPrimaryObjects
IdentifyPrimaryObjects
RelateObjects
ConvertObjectsToImage
SaveImages
OverlayOutlines
OverlayOutlines
SaveImages
SaveImages
SaveCroppedObjects
MeasureObjectSizeShape
MeasureObjectIntensity
ExportToSpreadsheet
This module saves the binary images of the PDL1 immunolabelling after thresholding.
Saving the binary image may prove useful, as it clearly displays all of the pixels which will subsequently be integrated (summed) inside cell-objects. Being a binary, image-data cannot 'hide' low down in the image bit-depth. These images are therefore useful for checking that the thresholding process is acting as desired. They are also useful for checking that signal is being adequately removed from secondary-only and isotype controls.
Select the type of image to save Image
Select the image to save thresh_PDL1_binary (from Threshold #07)
Select method for constructing file names Sequential numbers
Enter file prefix thresh_PDL1_binary
Number of digits 3
Saved file format tiff
Image bit depth 8-bit integer
Output file location Default Output Folder sub-folder ( d:\\cellprofiler_out )
Sub-folder: thresh_PDL1_binary
Overwrite existing files without warning? Yes No
When to save Every cycle
Record the file and path information to the saved image? Yes No
Output Settings View Workspace
Adjust modules: + - ^ v
Start Test Mode
Analyze Images
Found 6 rows
Type here to search
27°C Sunny
17:55
16/07/2022

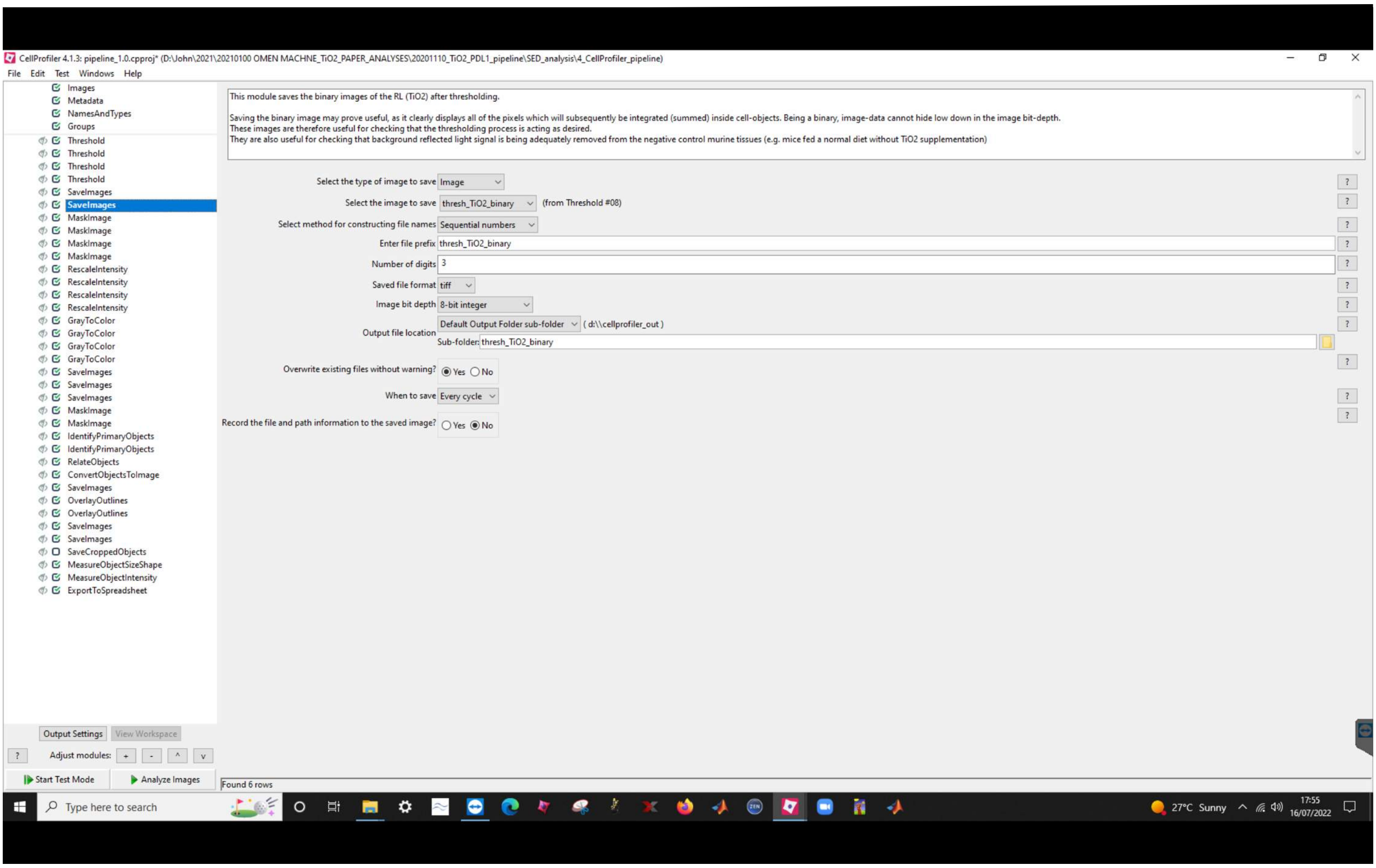
CellProfiler 4.1.3: pipeline_1.0.cpproj* (D:\John\2021\20210100 OMEN MACHNE_TiO2_PAPER_ANALYSES\20201110_TiO2_PDL1_pipeline\SED_analysis\4_CellProfiler_pipeline)
File Edit Test Windows Help
Images
Metadata
NamesAndTypes
Groups
Threshold
Threshold
Threshold
Threshold
SaveImages
SaveImages
MaskImage
MaskImage
MaskImage
MaskImage
RescaleIntensity
RescaleIntensity
RescaleIntensity
RescaleIntensity
GrayToColor
GrayToColor
GrayToColor
GrayToColor
SaveImages
SaveImages
SaveImages
MaskImage
MaskImage
IdentifyPrimaryObjects
IdentifyPrimaryObjects
RelateObjects
ConvertObjectsToImage
SaveImages
OverlayOutlines
OverlayOutlines
SaveImages
SaveImages
SaveCroppedObjects
MeasureObjectSizeShape
MeasureObjectIntensity
ExportToSpreadsheet
This module saves the binary images of the RL (TiO2) after thresholding.
Saving the binary image may prove useful, as it clearly displays all of the pixels which will subsequently be integrated (summed) inside cell-objects. Being a binary, image-data cannot hide low down in the image bit-depth.
These images are therefore useful for checking that the thresholding process is acting as desired.
They are also useful for checking that background reflected light signal is being adequately removed from the negative control murine tissues (e.g. mice fed a normal diet without TiO2 supplementation)
Select the type of image to save Image
Select the image to save thresh_TiO2_binary (from Threshold #08)
Select method for constructing file names Sequential numbers
Enter file prefix thresh_TiO2_binary
Number of digits 3
Saved file format tiff
Image bit depth 8-bit integer
Output file location Default Output Folder sub-folder ( d:\\cellprofiler_out )
Sub-folder: thresh_TiO2_binary
Overwrite existing files without warning? Yes No
When to save Every cycle
Record the file and path information to the saved image? Yes No
Output Settings View Workspace
Adjust modules: + - ^ v
Start Test Mode
Analyze Images
Found 6 rows
Type here to search
27°C Sunny
17:55
16/07/2022

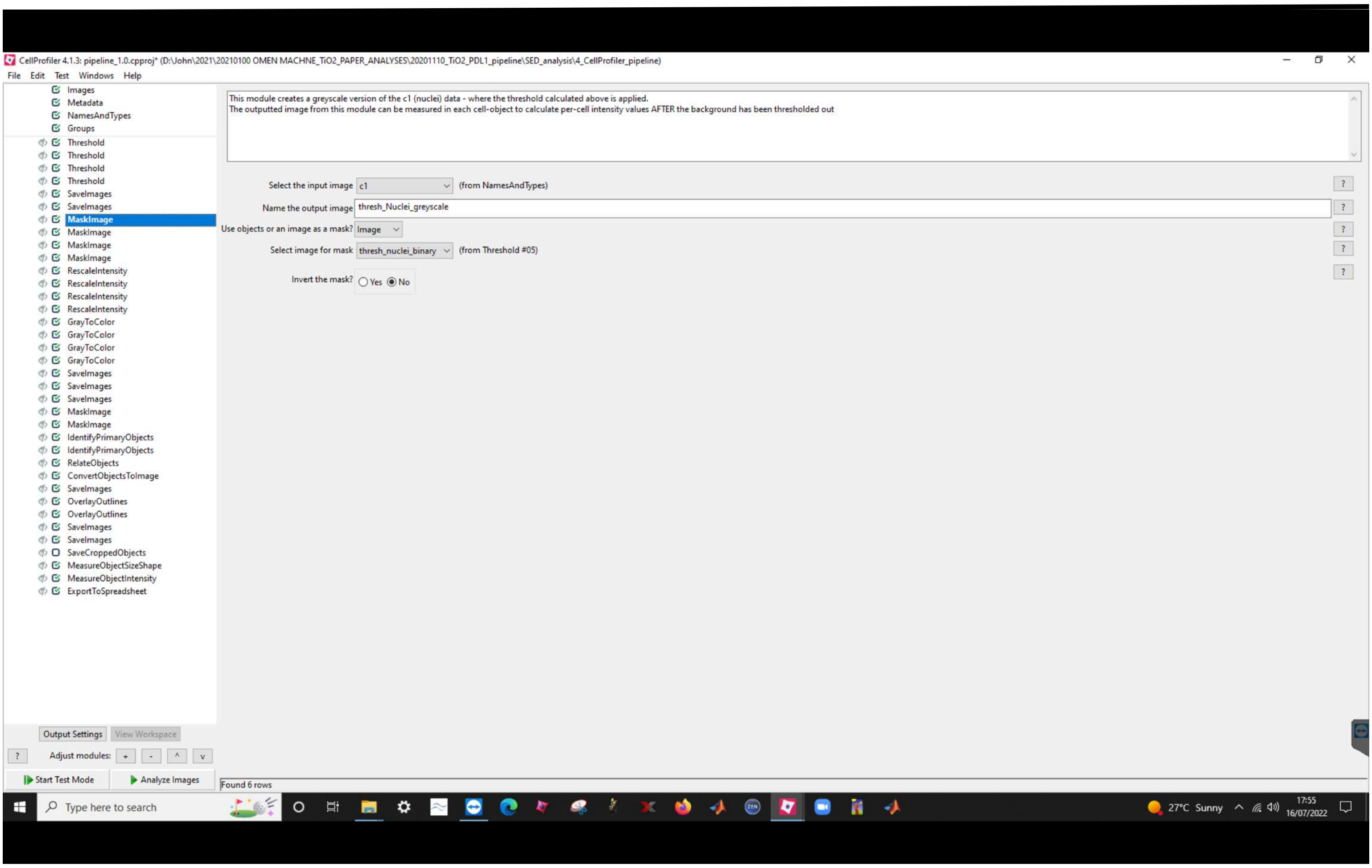
CellProfiler 4.1.3: pipeline_1.0.cpproj* (D:\John\2021\20210100 OMEN MACHNE_TiO2_PAPER_ANALYSES\20201110_TiO2_PDL1_pipeline\SED_analysis\4_CellProfiler_pipeline)
File Edit Test Windows Help
Images
Metadata
NamesAndTypes
Groups
Threshold
Threshold
Threshold
Threshold
SaveImages
SaveImages
MaskImage
MaskImage
MaskImage
MaskImage
RescaleIntensity
RescaleIntensity
RescaleIntensity
RescaleIntensity
GrayToColor
GrayToColor
GrayToColor
GrayToColor
SaveImages
SaveImages
SaveImages
MaskImage
MaskImage
IdentifyPrimaryObjects
IdentifyPrimaryObjects
RelateObjects
ConvertObjectsToImage
SaveImages
OverlayOutlines
OverlayOutlines
SaveImages
SaveImages
SaveCroppedObjects
MeasureObjectSizeShape
MeasureObjectIntensity
ExportToSpreadsheet
This module creates a greyscale version of the c1 (nuclei) data - where the threshold calculated above is applied.
The outputted image from this module can be measured in each cell-object to calculate per-cell intensity values AFTER the background has been thresholded out
Select the input image c1 (from NamesAndTypes)
Name the output image thresh_Nuclei_greyscale
Use objects or an image as a mask? Image
Select image for mask thresh_nuclei_binary (from Threshold #05)
Invert the mask? Yes No
Output Settings
View Workspace
Adjust modules: + - ^ v
Start Test Mode
Analyze Images
Found 6 rows
Type here to search
27°C Sunny
17:55
16/07/2022

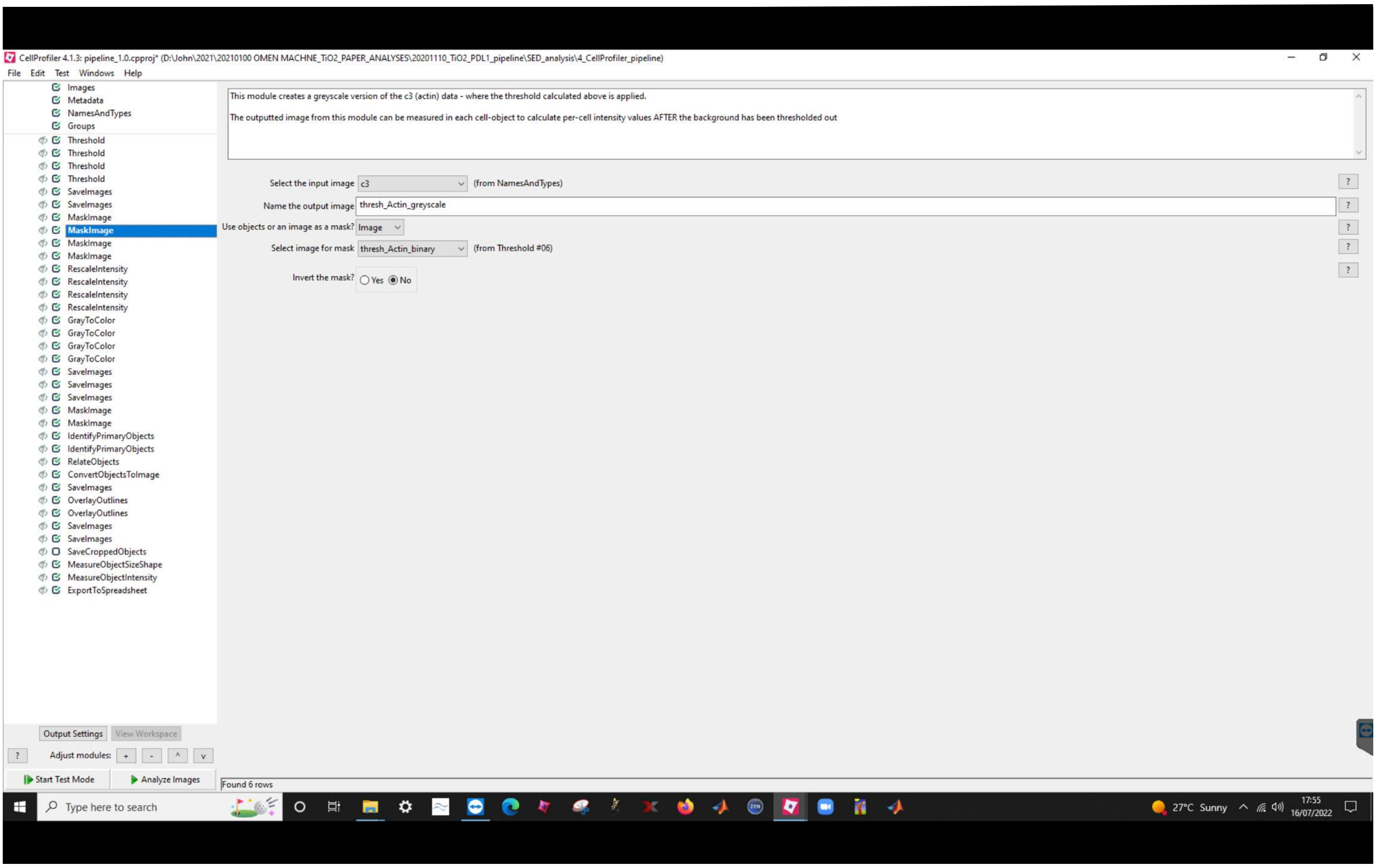
CellProfiler 4.1.3: pipeline_1.0.cpproj* (D:\John\2021\20210100 OMEN MACHNE_TiO2_PAPER_ANALYSES\20201110_TiO2_PDL1_pipeline\SED_analysis\4_CellProfiler_pipeline)
File Edit Test Windows Help
Images
Metadata
NamesAndTypes
Groups
Threshold
Threshold
Threshold
Threshold
SaveImages
SaveImages
MaskImage
MaskImage
MaskImage
MaskImage
RescaleIntensity
RescaleIntensity
RescaleIntensity
RescaleIntensity
GrayToColor
GrayToColor
GrayToColor
GrayToColor
SaveImages
SaveImages
SaveImages
MaskImage
MaskImage
IdentifyPrimaryObjects
IdentifyPrimaryObjects
RelateObjects
ConvertObjectsToImage
SaveImages
OverlayOutlines
OverlayOutlines
SaveImages
SaveImages
SaveCroppedObjects
MeasureObjectSizeShape
MeasureObjectIntensity
ExportToSpreadsheet
This module creates a greyscale version of the c3 (actin) data - where the threshold calculated above is applied.
The outputted image from this module can be measured in each cell-object to calculate per-cell intensity values AFTER the background has been thresholded out
Select the input image c3 (from NamesAndTypes)
Name the output image thresh_Actin_greyscale
Use objects or an image as a mask? Image
Select image for mask thresh_Actin_binary (from Threshold #06)
Invert the mask? Yes No
Output Settings View Workspace
Adjust modules:
Start Test Mode
Analyze Images
Found 6 rows
Type here to search
27°C Sunny
17:55
16/07/2022

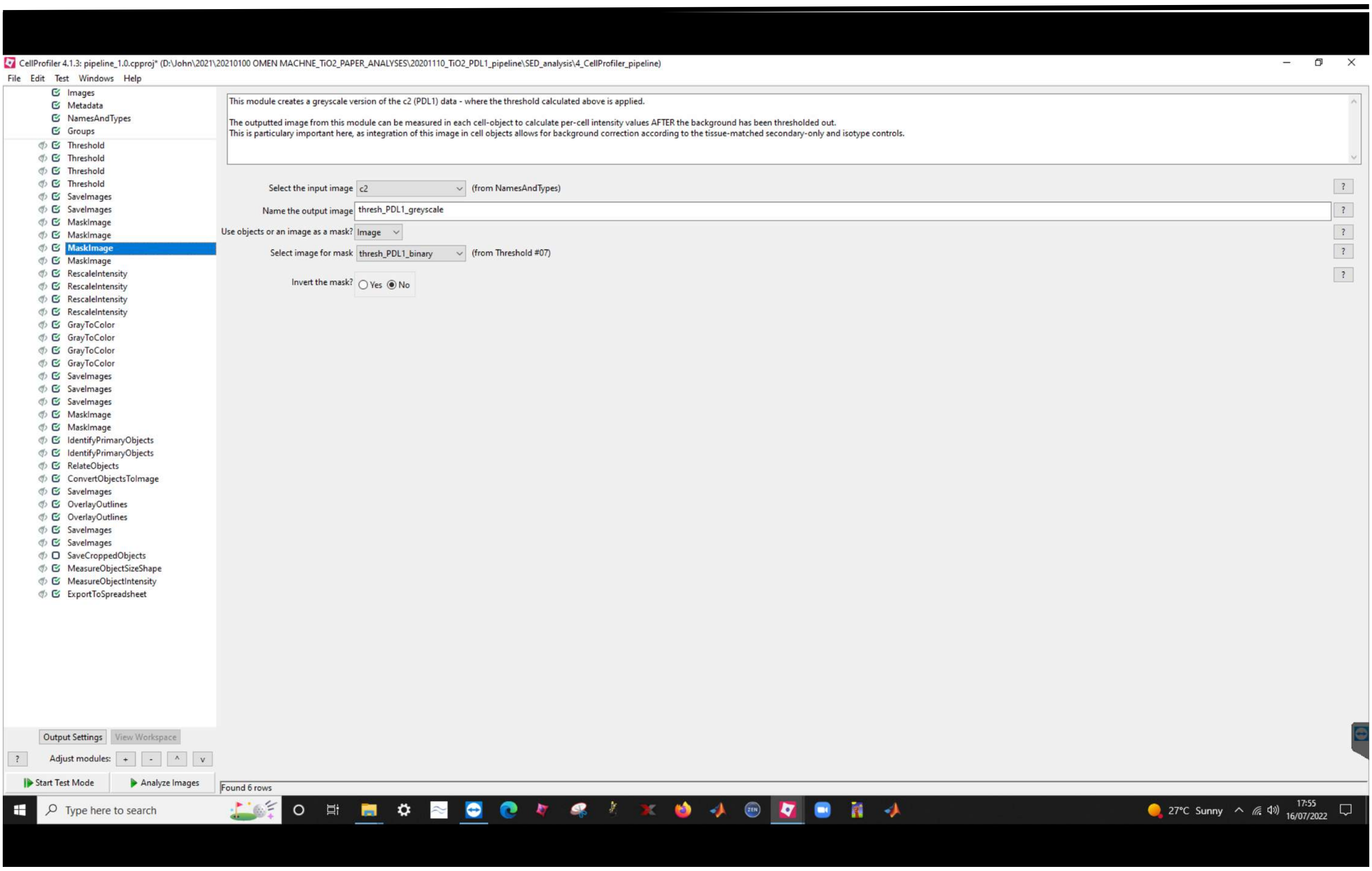
CellProfiler 4.1.3: pipeline_1.0.cpproj* (D:\John\2021\20210100 OMEN MACHNE_TiO2_PAPER_ANALYSES\20201110_TiO2_PDL1_pipeline\SED_analysis\4_CellProfiler_pipeline)
File Edit Test Windows Help
Images
Metadata
NamesAndTypes
Groups
Threshold
Threshold
Threshold
Threshold
SaveImages
SaveImages
MaskImage
MaskImage
MaskImage
MaskImage
RescaleIntensity
RescaleIntensity
RescaleIntensity
RescaleIntensity
GrayToColor
GrayToColor
GrayToColor
GrayToColor
SaveImages
SaveImages
SaveImages
MaskImage
MaskImage
IdentifyPrimaryObjects
IdentifyPrimaryObjects
RelateObjects
ConvertObjectsToImage
SaveImages
OverlayOutlines
OverlayOutlines
SaveImages
SaveImages
SaveCroppedObjects
MeasureObjectSizeShape
MeasureObjectIntensity
ExportToSpreadsheet
This module creates a greyscale version of the c2 (PDL1) data - where the threshold calculated above is applied.
The outputted image from this module can be measured in each cell-object to calculate per-cell intensity values AFTER the background has been thresholded out.
This is particulary important here, as integration of this image in cell objects allows for background correction according to the tissue-matched secondary-only and isotype controls.
Select the input image c2 (from NamesAndTypes)
Name the output image thresh_PDL1_greyscale
Use objects or an image as a mask? Image
Select image for mask thresh_PDL1_binary (from Threshold #07)
Invert the mask? Yes No
Output Settings
View Workspace
Adjust modules: + - ^ v
Start Test Mode
Analyze Images
Found 6 rows
Type here to search
27°C Sunny
17:55
16/07/2022

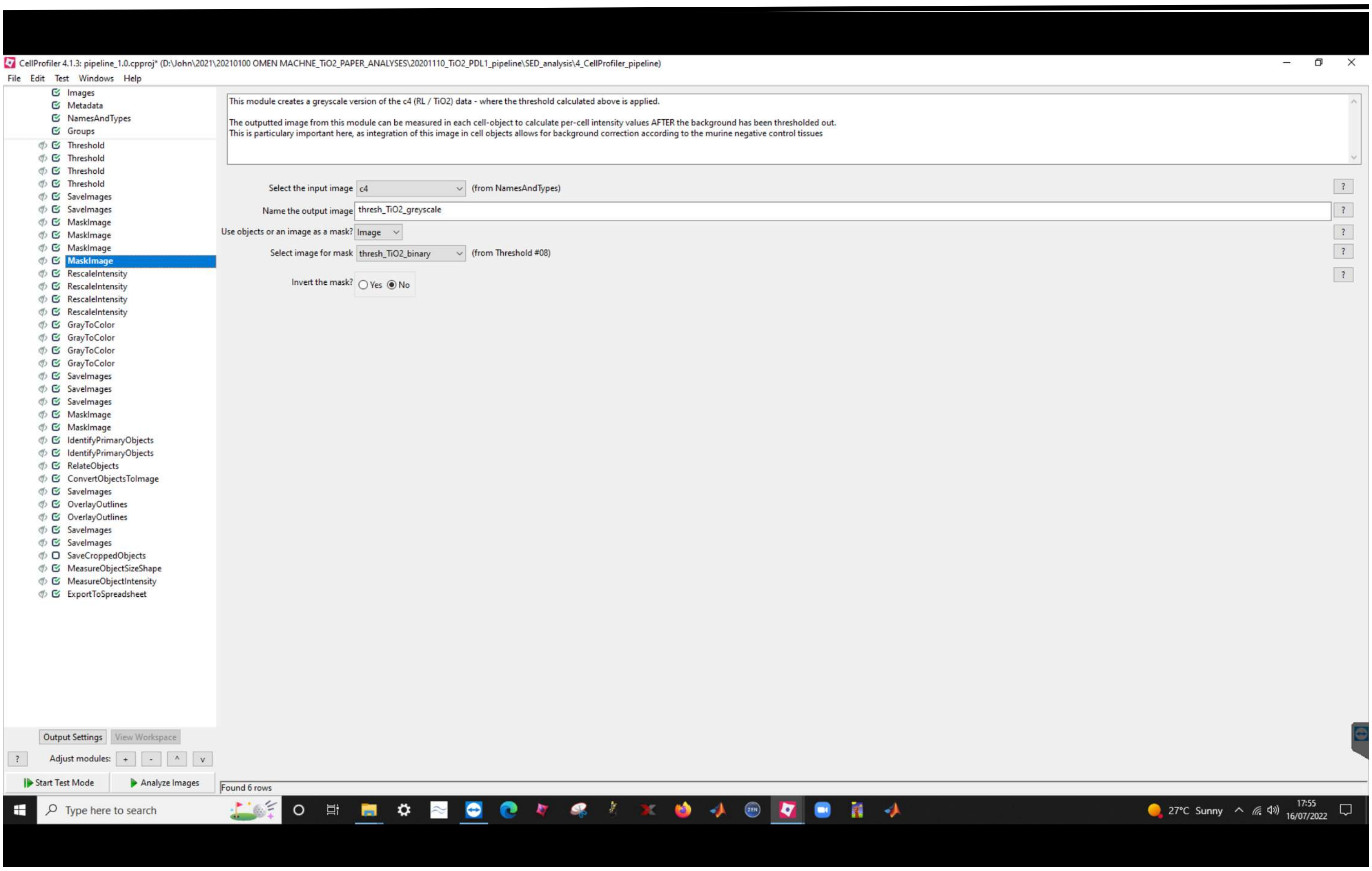
CellProfiler 4.1.3: pipeline_1.0.cpproj* (D:\John\2021\20210100 OMEN MACHNE_TiO2_PAPER_ANALYSES\20201110_TiO2_PDL1_pipeline\SED_analysis\4_CellProfiler_pipeline)
File Edit Test Windows Help
Images
Metadata
NamesAndTypes
Groups
Threshold
Threshold
Threshold
Threshold
SaveImages
SaveImages
MaskImage
MaskImage
MaskImage
MaskImage
RescaleIntensity
RescaleIntensity
RescaleIntensity
RescaleIntensity
GrayToColor
GrayToColor
GrayToColor
GrayToColor
SaveImages
SaveImages
SaveImages
MaskImage
MaskImage
IdentifyPrimaryObjects
IdentifyPrimaryObjects
RelateObjects
ConvertObjectsToImage
SaveImages
OverlayOutlines
OverlayOutlines
SaveImages
SaveImages
SaveCroppedObjects
MeasureObjectSizeShape
MeasureObjectIntensity
ExportToSpreadsheet
This module creates a greyscale version of the c4 (RL / TiO2) data - where the threshold calculated above is applied.
The outputted image from this module can be measured in each cell-object to calculate per-cell intensity values AFTER the background has been thresholded out.
This is particulary important here, as integration of this image in cell objects allows for background correction according to the murine negative control tissues
Select the input image c4 (from NamesAndTypes)
Name the output image thresh_TiO2_greyscale
Use objects or an image as a mask? Image
Select image for mask thresh_TiO2_binary (from Threshold #08)
Invert the mask? Yes No
Output Settings View Workspace
Adjust modules: + - ^ v
Start Test Mode Analyze Images
Found 6 rows
Type here to search
27°C Sunny
17:55
16/07/2022

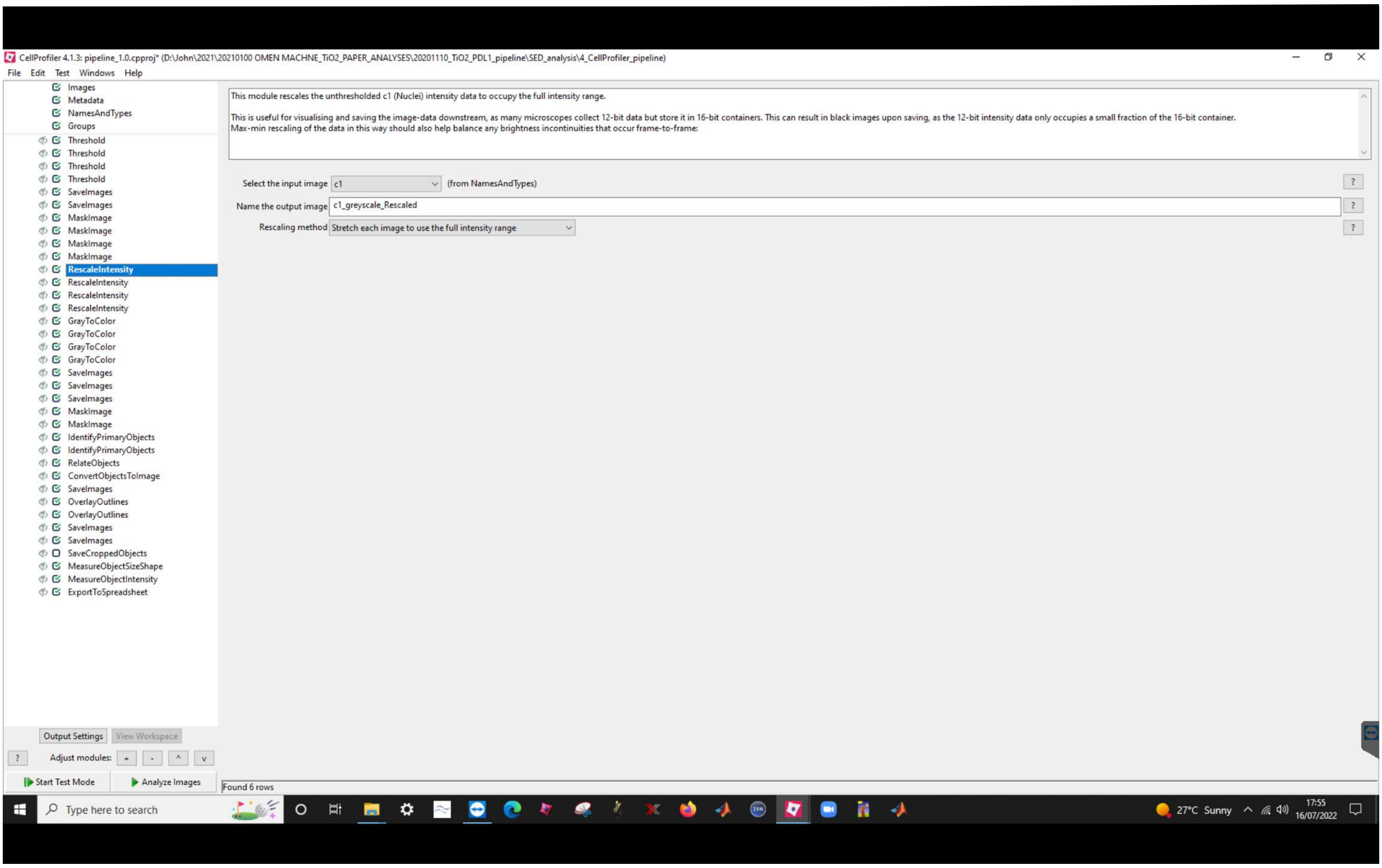
CellProfiler 4.1.3: pipeline_1.0.cpproj* (D:\John\2021\20210100 OMEN MACHNE_TiO2_PAPER_ANALYSES\20201110_TiO2_PDL1_pipeline\SED_analysis\4_CellProfiler_pipeline)
File Edit Test Windows Help
Images
Metadata
NamesAndTypes
Groups
Threshold
Threshold
Threshold
Threshold
SaveImages
SaveImages
MaskImage
MaskImage
MaskImage
MaskImage
RescaleIntensity
RescaleIntensity
RescaleIntensity
RescaleIntensity
GrayToColor
GrayToColor
GrayToColor
GrayToColor
SaveImages
SaveImages
SaveImages
MaskImage
MaskImage
IdentifyPrimaryObjects
IdentifyPrimaryObjects
RelateObjects
ConvertObjectsToImage
SaveImages
OverlayOutlines
OverlayOutlines
SaveImages
SaveImages
SaveCroppedObjects
MeasureObjectSizeShape
MeasureObjectIntensity
ExportToSpreadsheet
This module rescales the unthresholded c1 (Nuclei) intensity data to occupy the full intensity range.
This is useful for visualising and saving the image-data downstream, as many microscopes collect 12-bit data but store it in 16-bit containers. This can result in black images upon saving, as the 12-bit intensity data only occupies a small fraction of the 16-bit container.
Max-min rescaling of the data in this way should also help balance any brightness incontinuities that occur frame-to-frame.
Select the input image c1 (from NamesAndTypes)
Name the output image c1_greyscale_Rescaled
Rescaling method Stretch each image to use the full intensity range
Output Settings
View Workspace
Adjust modules:
Start Test Mode
Analyze Images
Found 6 rows
Type here to search
27°C Sunny
17:55
16/07/2022

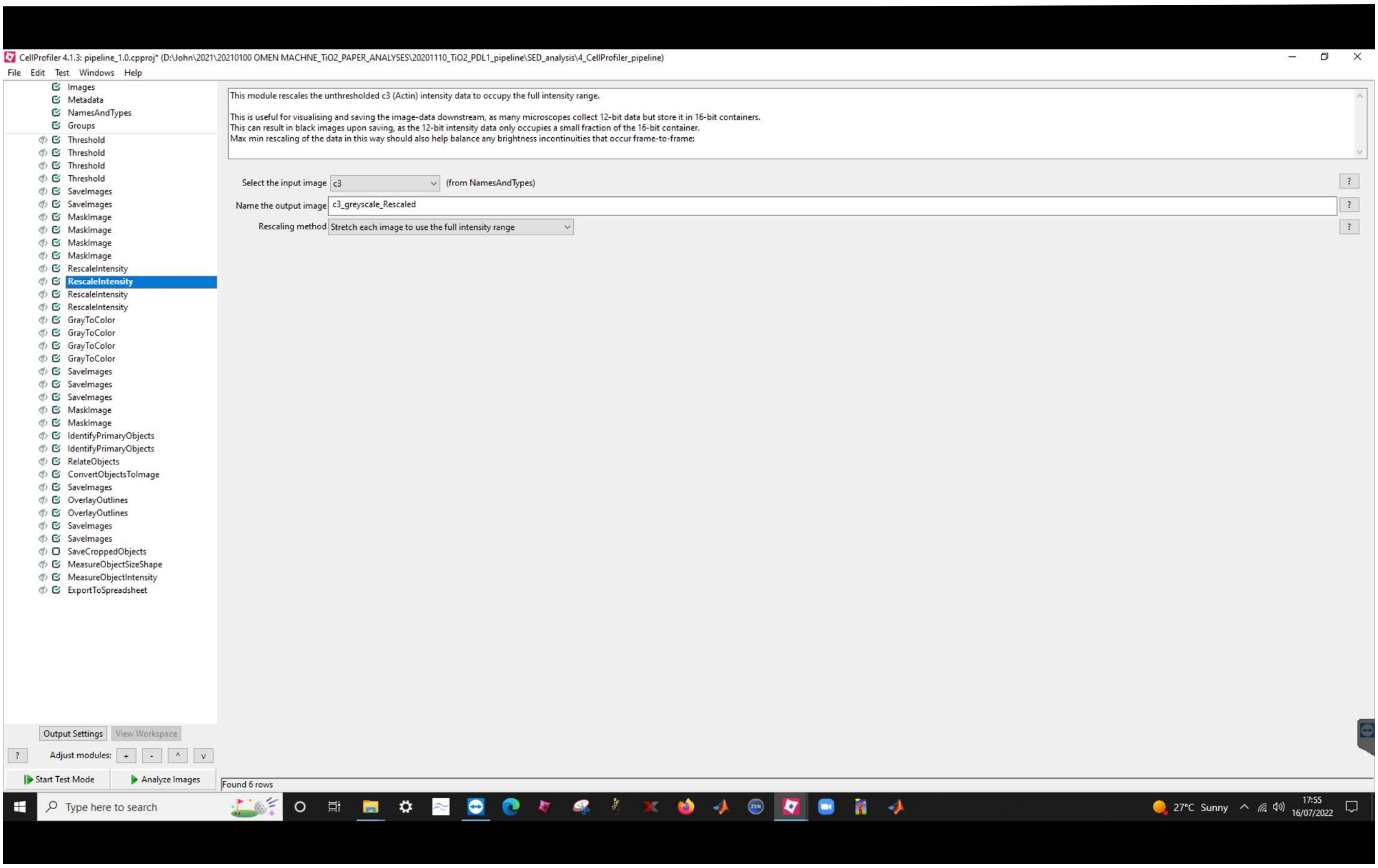
CellProfiler 4.1.3: pipeline_1.0.cpproj* (D:\John\2021\20210100 OMEN MACHNE_TiO2_PAPER_ANALYSES\20201110_TiO2_PDL1_pipeline\SED_analysis\4_CellProfiler_pipeline)
File Edit Test Windows Help
Images
Metadata
NamesAndTypes
Groups
Threshold
Threshold
Threshold
Threshold
SaveImages
SaveImages
MaskImage
MaskImage
MaskImage
MaskImage
RescaleIntensity
RescaleIntensity
RescaleIntensity
RescaleIntensity
GrayToColor
GrayToColor
GrayToColor
GrayToColor
SaveImages
SaveImages
SaveImages
MaskImage
MaskImage
IdentifyPrimaryObjects
IdentifyPrimaryObjects
RelateObjects
ConvertObjectsToImage
SaveImages
OverlayOutlines
OverlayOutlines
SaveImages
SaveImages
SaveCroppedObjects
MeasureObjectSizeShape
MeasureObjectIntensity
ExportToSpreadsheet
This module rescales the unthresholded c3 (Actin) intensity data to occupy the full intensity range.
This is useful for visualising and saving the image-data downstream, as many microscopes collect 12-bit data but store it in 16-bit containers.
This can result in black images upon saving, as the 12-bit intensity data only occupies a small fraction of the 16-bit container.
Max min rescaling of the data in this way should also help balance any brightness incontinuities that occur frame-to-frame:
Select the input image c3 (from NamesAndTypes)
Name the output image c3_greyscale_Rescaled
Rescaling method Stretch each image to use the full intensity range
Output Settings
View Workspace
Adjust modules:
Start Test Mode
Analyze Images
Found 6 rows
Type here to search
27°C Sunny
17:55
16/07/2022

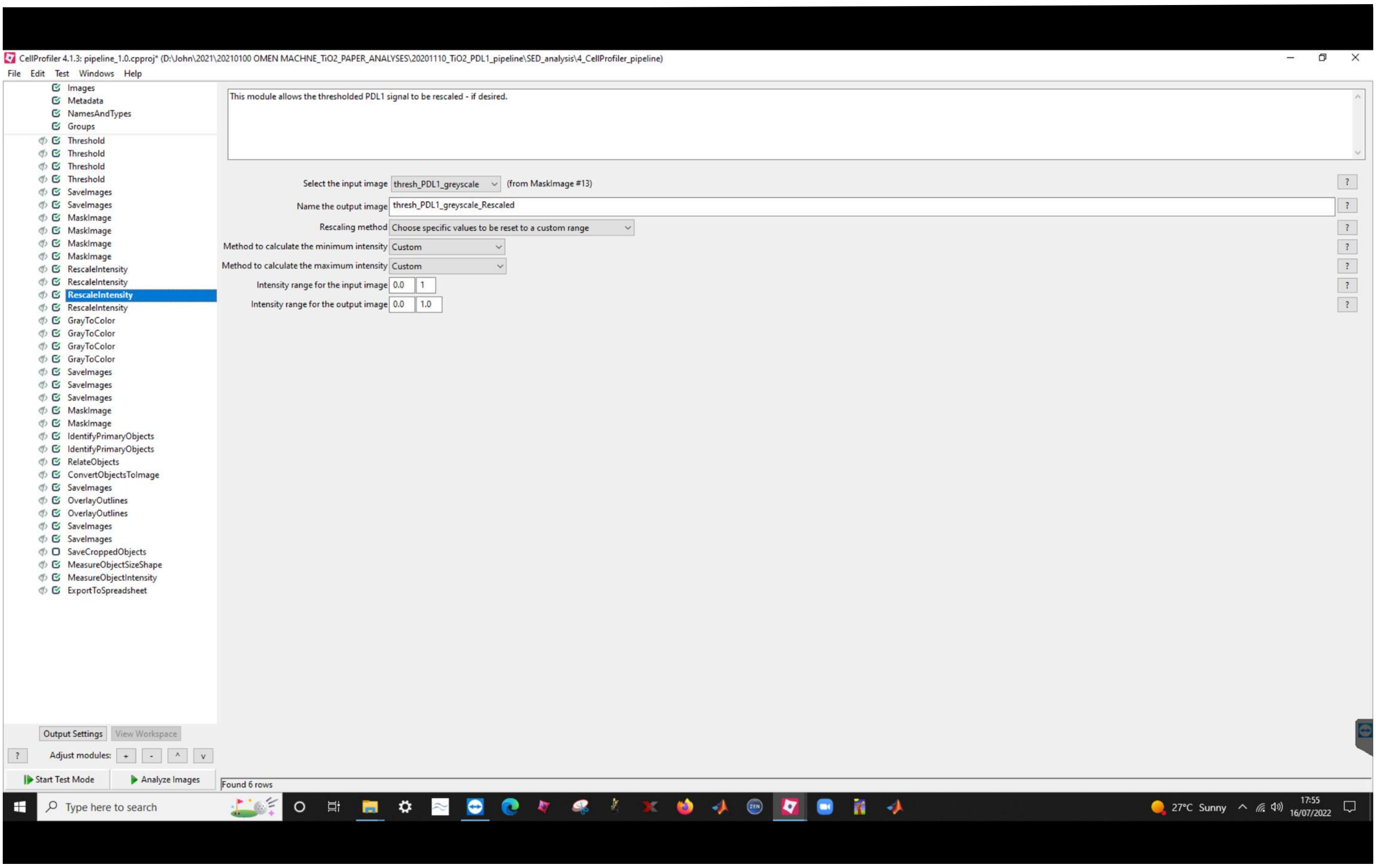
CellProfiler 4.1.3: pipeline_1.0.cpproj* (D:\John\2021\20210100 OMEN MACHNE_TiO2_PAPER_ANALYSES\20201110_TiO2_PDL1_pipeline\SED_analysis\4_CellProfiler_pipeline)
File Edit Test Windows Help
Images
Metadata
NamesAndTypes
Groups
Threshold
Threshold
Threshold
Threshold
SaveImages
SaveImages
MaskImage
MaskImage
MaskImage
MaskImage
RescaleIntensity
RescaleIntensity
RescaleIntensity
RescaleIntensity
GrayToColor
GrayToColor
GrayToColor
GrayToColor
SaveImages
SaveImages
SaveImages
MaskImage
MaskImage
IdentifyPrimaryObjects
IdentifyPrimaryObjects
RelateObjects
ConvertObjectsToImage
SaveImages
OverlayOutlines
OverlayOutlines
SaveImages
SaveImages
SaveCroppedObjects
MeasureObjectSizeShape
MeasureObjectIntensity
ExportToSpreadsheet
This module allows the thresholded PDL1 signal to be rescaled - if desired.
Select the input image thresh_PDL1_greyscale (from MaskImage #13)
Name the output image thresh_PDL1_greyscale_Rescaled
Rescaling method Choose specific values to be reset to a custom range
Method to calculate the minimum intensity Custom
Method to calculate the maximum intensity Custom
Intensity range for the input image 0.0 1
Intensity range for the output image 0.0 1.0
Output Settings
View Workspace
Adjust modules: + - ^ v
Start Test Mode
Analyze Images
Found 6 rows
Type here to search
27°C Sunny
17:55
16/07/2022

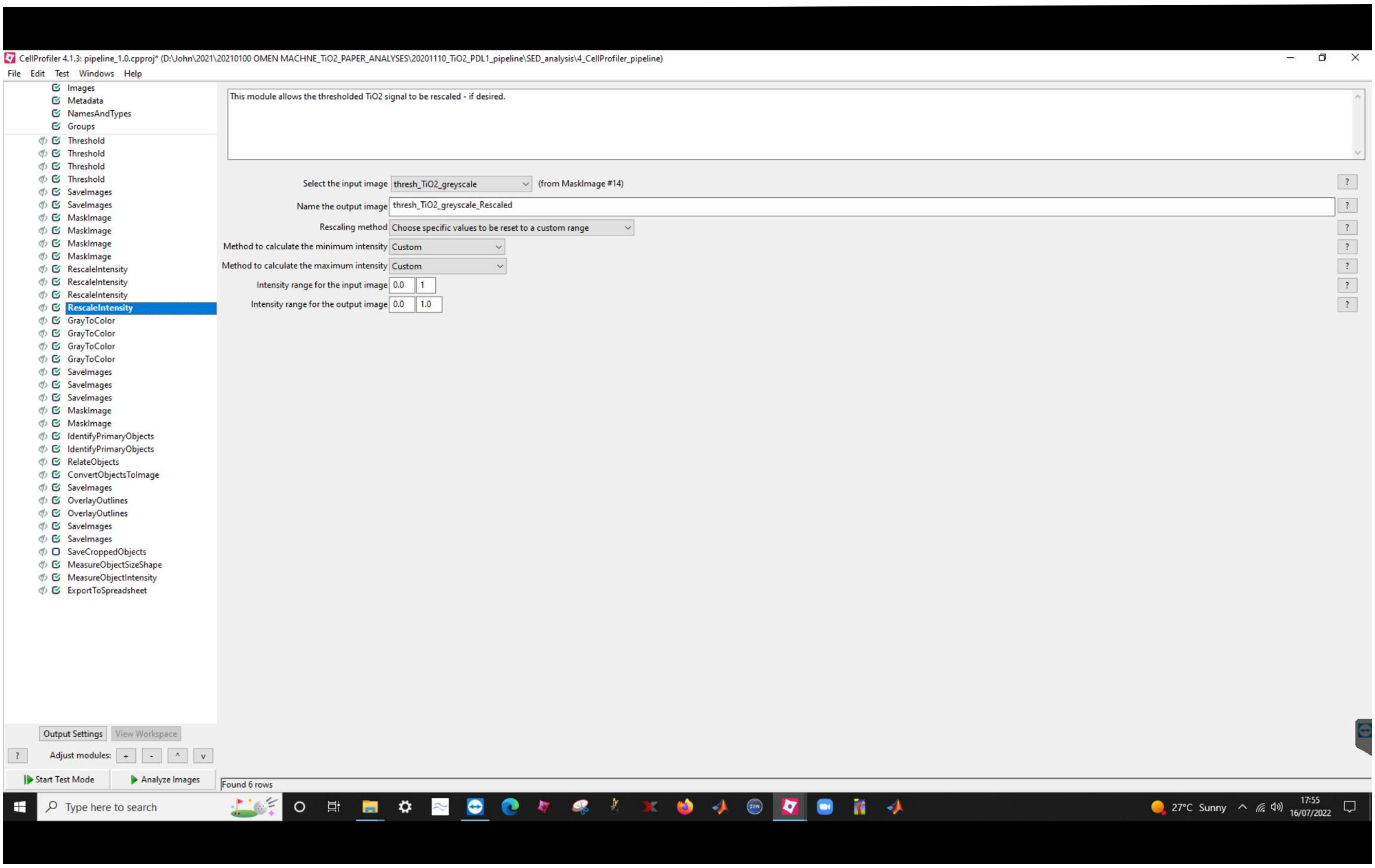
CellProfiler 4.1.3: pipeline_1.0.cpproj* (D:\John\2021\20210100 OMEN MACHNE_TiO2_PAPER_ANALYSES\20201110_TiO2_PDL1_pipeline\SED_analysis\4_CellProfiler_pipeline)
File Edit Test Windows Help
Images
Metadata
NamesAndTypes
Groups
Threshold
Threshold
Threshold
Threshold
SaveImages
SaveImages
MaskImage
MaskImage
MaskImage
MaskImage
RescaleIntensity
RescaleIntensity
RescaleIntensity
RescaleIntensity
GrayToColor
GrayToColor
GrayToColor
GrayToColor
SaveImages
SaveImages
SaveImages
MaskImage
MaskImage
IdentifyPrimaryObjects
IdentifyPrimaryObjects
RelateObjects
ConvertObjectsToImage
SaveImages
OverlayOutlines
OverlayOutlines
SaveImages
SaveImages
SaveCroppedObjects
MeasureObjectSizeShape
MeasureObjectIntensity
ExportToSpreadsheet
This module allows the thresholded TiO2 signal to be rescaled - if desired.
Select the input image thresh_TiO2_greyscale (from MaskImage #14)
Name the output image thresh_TiO2_greyscale_Rescaled
Rescaling method Choose specific values to be reset to a custom range
Method to calculate the minimum intensity Custom
Method to calculate the maximum intensity Custom
Intensity range for the input image 0.0 1
Intensity range for the output image 0.0 1.0
Output Settings View Workspace
Adjust modules: + - ^ v
Start Test Mode
Analyze Images
Found 6 rows
Type here to search
27°C Sunny
17:55
16/07/2022

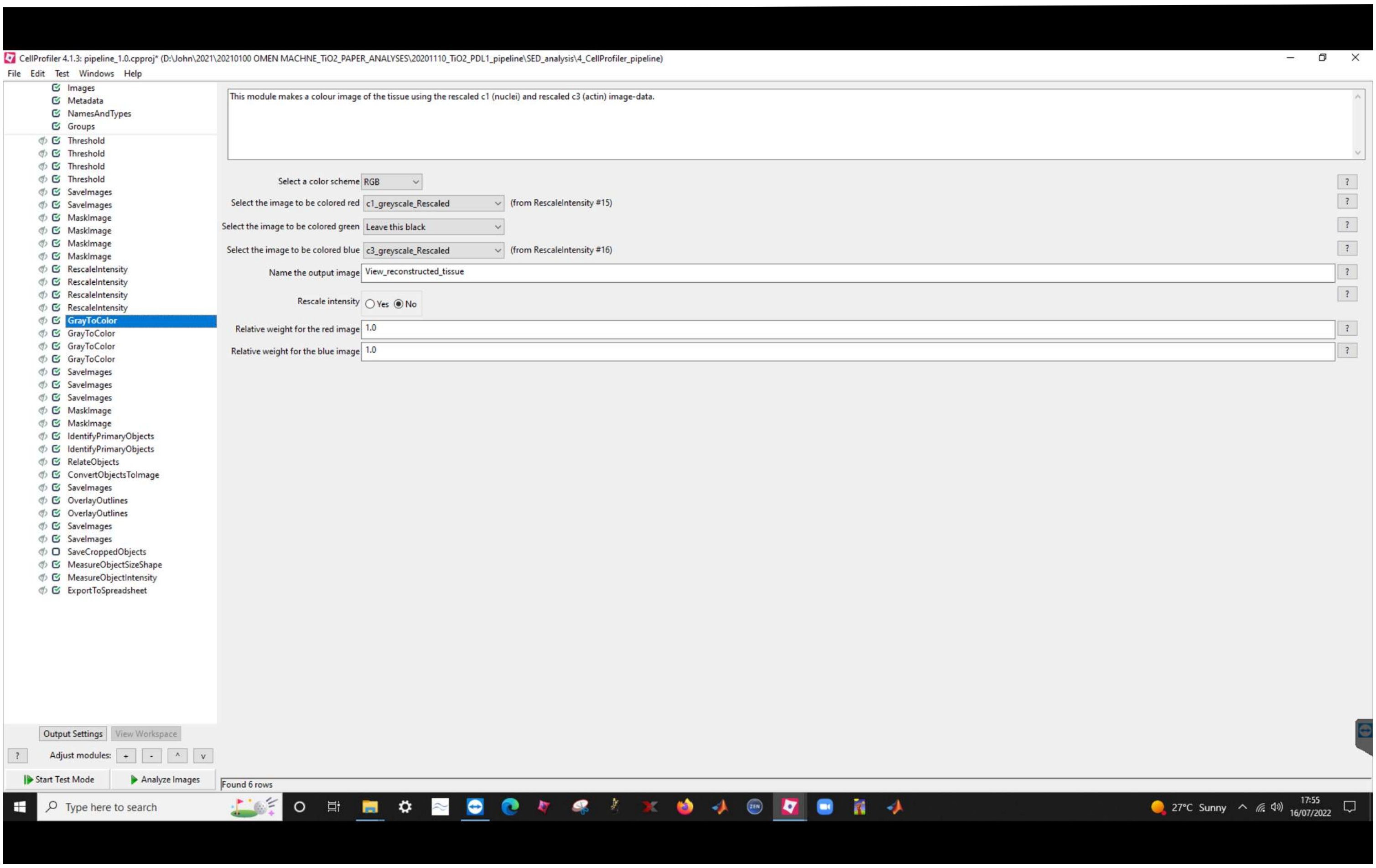
CellProfiler 4.1.3: pipeline_1.0.cpproj* (D:\John\2021\20210100 OMEN MACHNE_TiO2_PAPER_ANALYSES\20201110_TiO2_PDL1_pipeline\SED_analysis\4_CellProfiler_pipeline)
File Edit Test Windows Help
Images
Metadata
NamesAndTypes
Groups
Threshold
Threshold
Threshold
Threshold
SaveImages
SaveImages
MaskImage
MaskImage
MaskImage
MaskImage
RescaleIntensity
RescaleIntensity
RescaleIntensity
RescaleIntensity
GrayToColor
GrayToColor
GrayToColor
GrayToColor
SaveImages
SaveImages
SaveImages
MaskImage
MaskImage
IdentifyPrimaryObjects
IdentifyPrimaryObjects
RelateObjects
ConvertObjectsToImage
SaveImages
OverlayOutlines
OverlayOutlines
SaveImages
SaveImages
SaveCroppedObjects
MeasureObjectSizeShape
MeasureObjectIntensity
ExportToSpreadsheet
This module makes a colour image of the tissue using the rescaled c1 (nuclei) and rescaled c3 (actin) image-data.
Select a color scheme RGB
Select the image to be colored red c1_greyscale_Rescaled (from RescaleIntensity #15)
Select the image to be colored green Leave this black
Select the image to be colored blue c3_greyscale_Rescaled (from RescaleIntensity #16)
Name the output image View_reconstructed_tissue
Rescale intensity Yes No
Relative weight for the red image 1.0
Relative weight for the blue image 1.0
Output Settings View Workspace
Adjust modules: + - ^ v
Start Test Mode
Analyze Images
Found 6 rows
Type here to search
27°C Sunny
17:55
16/07/2022

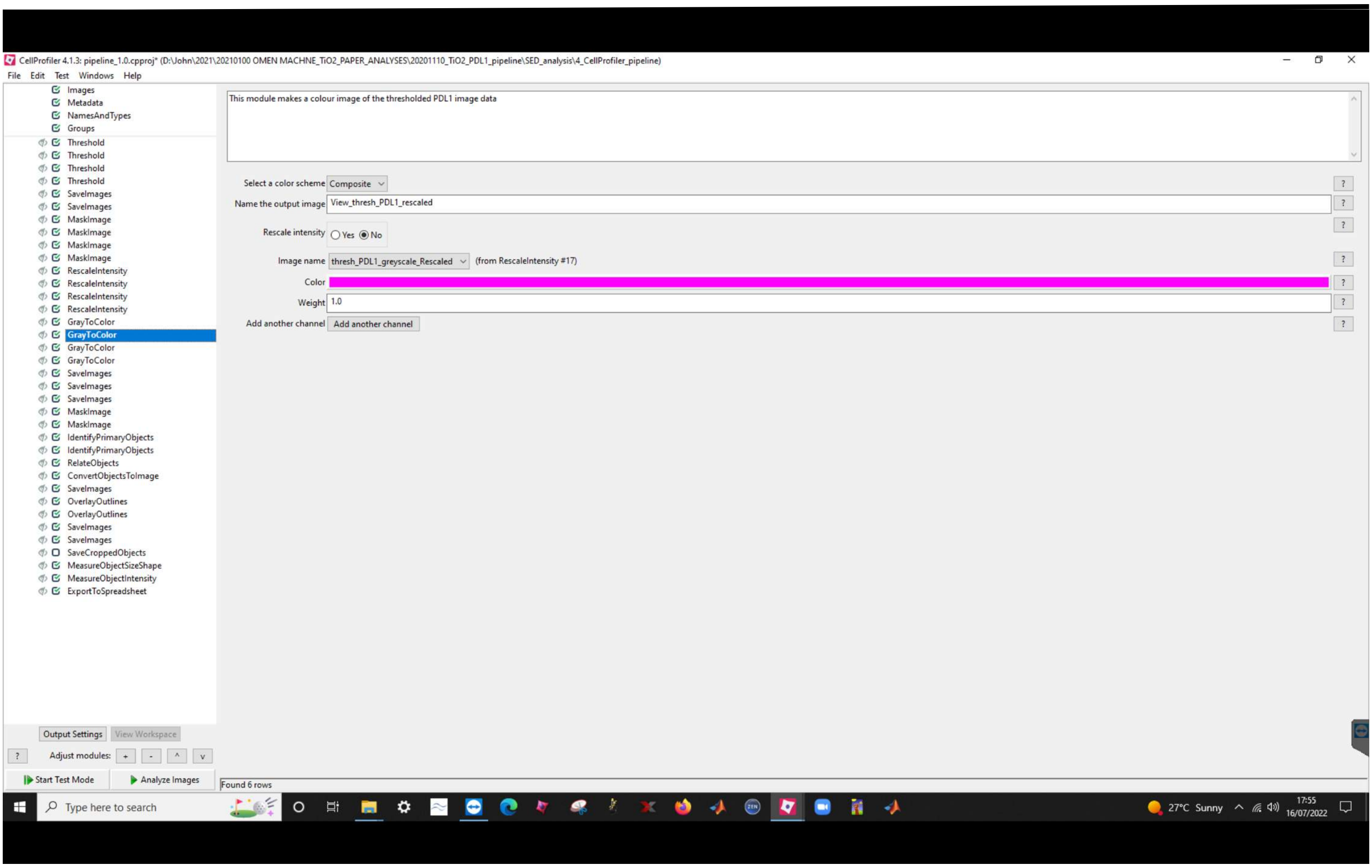
CellProfiler 4.1.3: pipeline_1.0.cpproj* (D:\John\2021\20210100 OMEN MACHNE_TiO2_PAPER_ANALYSES\20201110_TiO2_PDL1_pipeline\SED_analysis\4_CellProfiler_pipeline)
File Edit Test Windows Help
Images
Metadata
NamesAndTypes
Groups
Threshold
Threshold
Threshold
Threshold
SaveImages
SaveImages
MaskImage
MaskImage
MaskImage
MaskImage
RescaleIntensity
RescaleIntensity
RescaleIntensity
RescaleIntensity
GrayToColor
GrayToColor
GrayToColor
GrayToColor
SaveImages
SaveImages
SaveImages
MaskImage
MaskImage
IdentifyPrimaryObjects
IdentifyPrimaryObjects
RelateObjects
ConvertObjectsToImage
SaveImages
OverlayOutlines
OverlayOutlines
SaveImages
SaveImages
SaveCroppedObjects
MeasureObjectSizeShape
MeasureObjectIntensity
ExportToSpreadsheet
This module makes a colour image of the thresholded PDL1 image data
Select a color scheme Composite
Name the output image View_thresh_PDL1_rescaled
Rescale intensity Yes No
Image name thresh_PDL1_greyscale_Rescaled (from RescaleIntensity #17)
Color
Weight 1.0
Add another channel Add another channel
Output Settings View Workspace
Adjust modules: + - ^ v
Start Test Mode
Analyze Images
Found 6 rows
Type here to search
27°C Sunny
17:55
16/07/2022

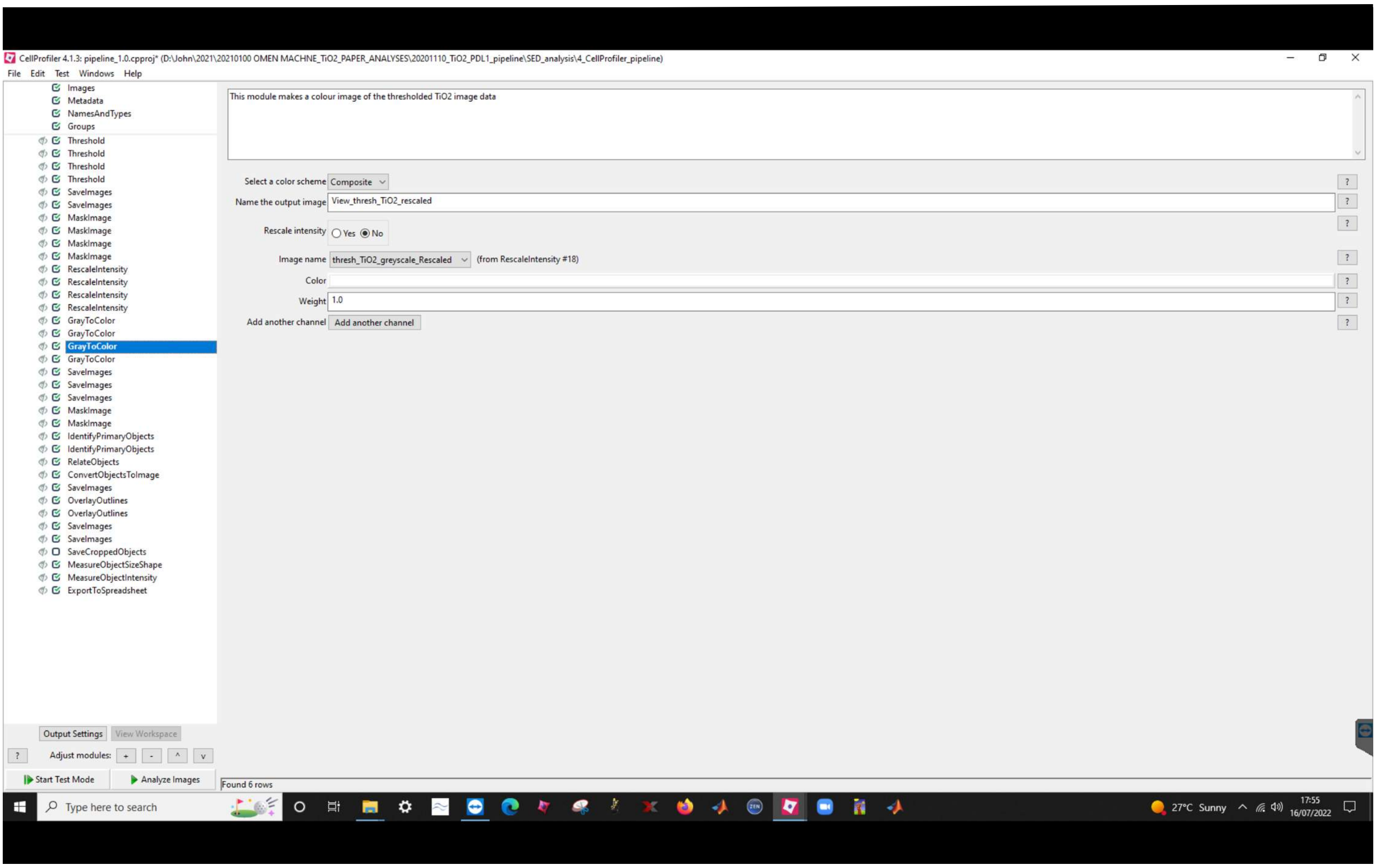
CellProfiler 4.1.3: pipeline_1.0.cpproj* (D:\John\2021\20210100 OMEN MACHNE_TiO2_PAPER_ANALYSES\20201110_TiO2_PDL1_pipeline\SED_analysis\4_CellProfiler_pipeline)
File Edit Test Windows Help
Images
Metadata
NamesAndTypes
Groups
Threshold
Threshold
Threshold
Threshold
SaveImages
SaveImages
MaskImage
MaskImage
MaskImage
MaskImage
RescaleIntensity
RescaleIntensity
RescaleIntensity
RescaleIntensity
GrayToColor
GrayToColor
GrayToColor
GrayToColor
SaveImages
SaveImages
SaveImages
MaskImage
MaskImage
IdentifyPrimaryObjects
IdentifyPrimaryObjects
RelateObjects
ConvertObjectsToImage
SaveImages
OverlayOutlines
OverlayOutlines
SaveImages
SaveImages
SaveCroppedObjects
MeasureObjectSizeShape
MeasureObjectIntensity
ExportToSpreadsheet
This module makes a colour image of the thresholded TiO2 image data
Select a color scheme Composite
Name the output image View_thresh_TiO2_rescaled
Rescale intensity Yes No
Image name thresh_TiO2_greyscale_Rescaled (from RescaleIntensity #18)
Color
Weight 1.0
Add another channel Add another channel
Output Settings View Workspace
Adjust modules: + - ^ v
Start Test Mode
Analyze Images
Found 6 rows
Type here to search
27°C Sunny
17:55
16/07/2022

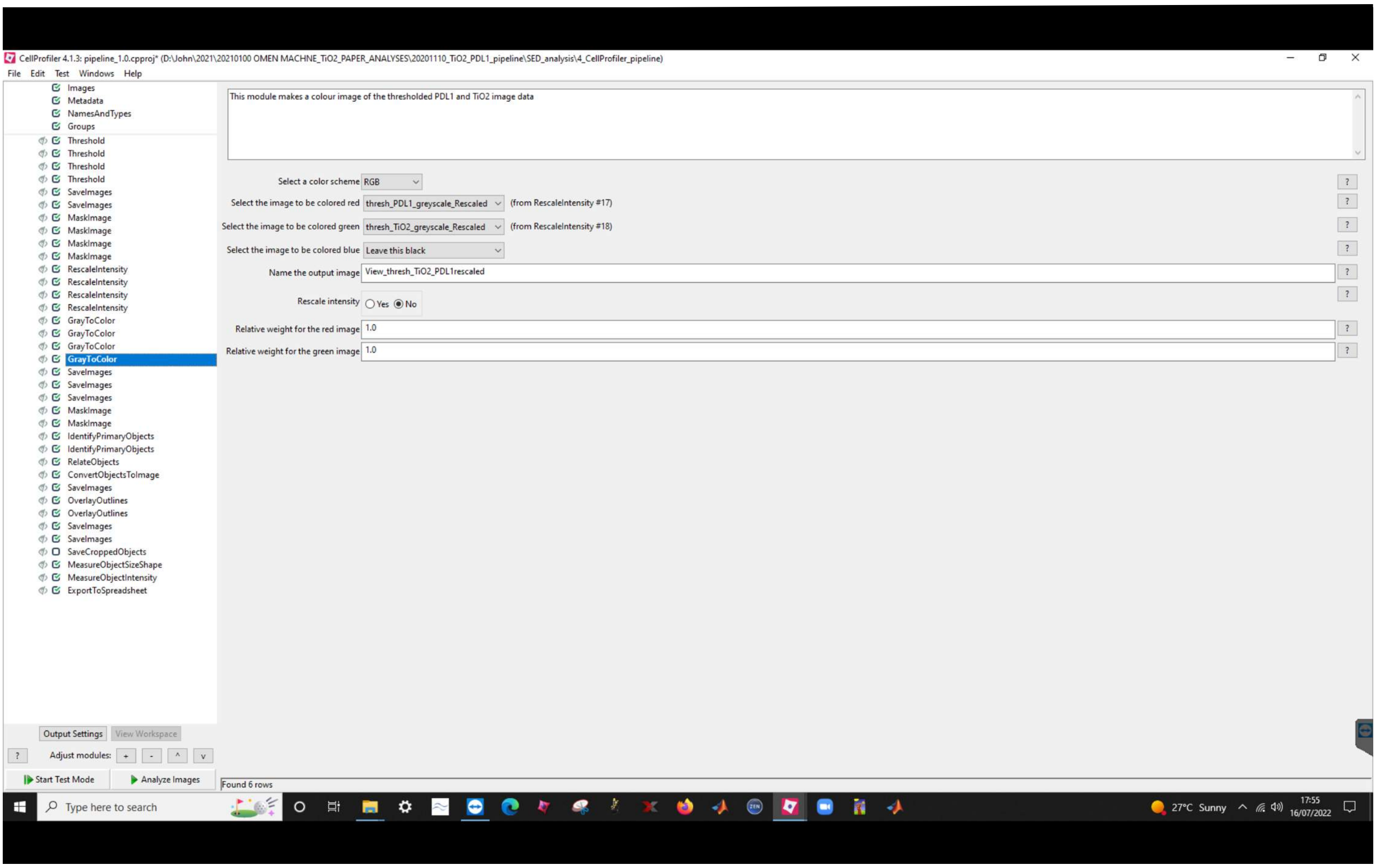
CellProfiler 4.1.3: pipeline_1.0.cpproj* (D:\John\2021\20210100 OMEN MACHNE_TiO2_PAPER_ANALYSES\20201110_TiO2_PDL1_pipeline\SED_analysis\4_CellProfiler_pipeline)
File Edit Test Windows Help
Images
Metadata
NamesAndTypes
Groups
Threshold
Threshold
Threshold
Threshold
SaveImages
SaveImages
MaskImage
MaskImage
MaskImage
MaskImage
RescaleIntensity
RescaleIntensity
RescaleIntensity
RescaleIntensity
GrayToColor
GrayToColor
GrayToColor
GrayToColor
SaveImages
SaveImages
SaveImages
MaskImage
MaskImage
IdentifyPrimaryObjects
IdentifyPrimaryObjects
RelateObjects
ConvertObjectsToImage
SaveImages
OverlayOutlines
OverlayOutlines
SaveImages
SaveImages
SaveCroppedObjects
MeasureObjectSizeShape
MeasureObjectIntensity
ExportToSpreadsheet
This module makes a colour image of the thresholded PDL1 and TiO2 image data
Select a color scheme RGB
Select the image to be colored red thresh_PDL1_greyscale_Rescaled (from RescaleIntensity #17)
Select the image to be colored green thresh_TiO2_greyscale_Rescaled (from RescaleIntensity #18)
Select the image to be colored blue Leave this black
Name the output image View_thresh_TiO2_PDL1rescaled
Rescale intensity Yes No
Relative weight for the red image 1.0
Relative weight for the green image 1.0
Output Settings View Workspace
Adjust modules: + - ^ v
Start Test Mode
Analyze Images
Found 6 rows
Type here to search
27°C Sunny
17:55
16/07/2022

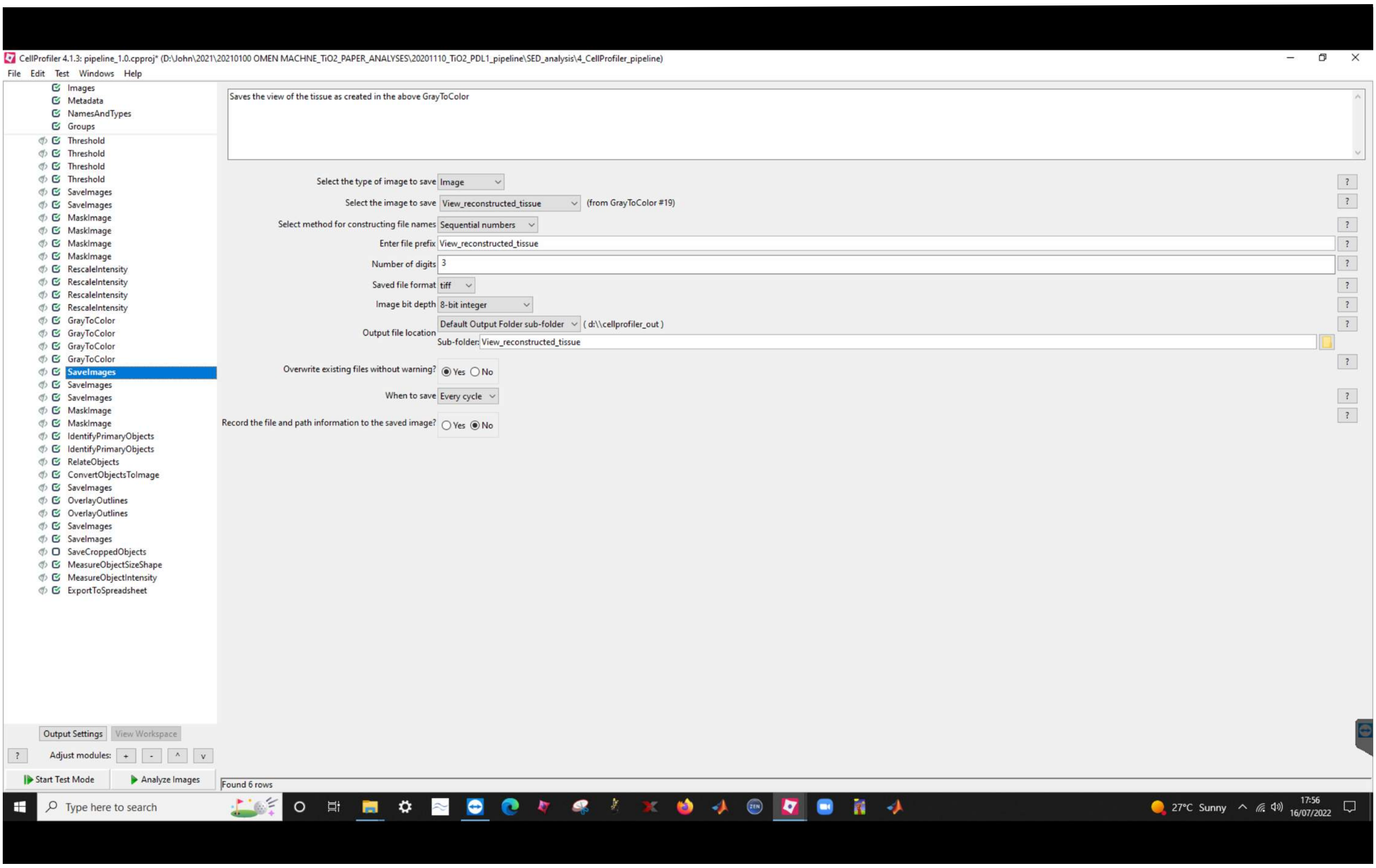
CellProfiler 4.1.3: pipeline_1.0.cpproj* (D:\John\2021\20210100 OMEN MACHNE_TiO2_PAPER_ANALYSES\20201110_TiO2_PDL1_pipeline\SED_analysis\4_CellProfiler_pipeline)
File Edit Test Windows Help
Images
Metadata
NamesAndTypes
Groups
Threshold
Threshold
Threshold
Threshold
SaveImages
SaveImages
MaskImage
MaskImage
MaskImage
MaskImage
RescaleIntensity
RescaleIntensity
RescaleIntensity
RescaleIntensity
GrayToColor
GrayToColor
GrayToColor
GrayToColor
SaveImages
SaveImages
SaveImages
MaskImage
MaskImage
IdentifyPrimaryObjects
IdentifyPrimaryObjects
RelateObjects
ConvertObjectsToImage
SaveImages
OverlayOutlines
OverlayOutlines
SaveImages
SaveImages
SaveCroppedObjects
MeasureObjectSizeShape
MeasureObjectIntensity
ExportToSpreadsheet
Saves the view of the tissue as created in the above GrayToColor
Select the type of image to save Image
Select the image to save View_reconstructed_tissue (from GrayToColor #19)
Select method for constructing file names Sequential numbers
Enter file prefix View_reconstructed_tissue
Number of digits 3
Saved file format tiff
Image bit depth 8-bit integer
Output file location Default Output Folder sub-folder ( d:\\cellprofiler_out )
Sub-folder: View_reconstructed_tissue
Overwrite existing files without warning? Yes No
When to save Every cycle
Record the file and path information to the saved image? Yes No
Output Settings View Workspace
Adjust modules: + - ^ v
Start Test Mode
Analyze Images
Found 6 rows
Type here to search
27°C Sunny
17:56
16/07/2022

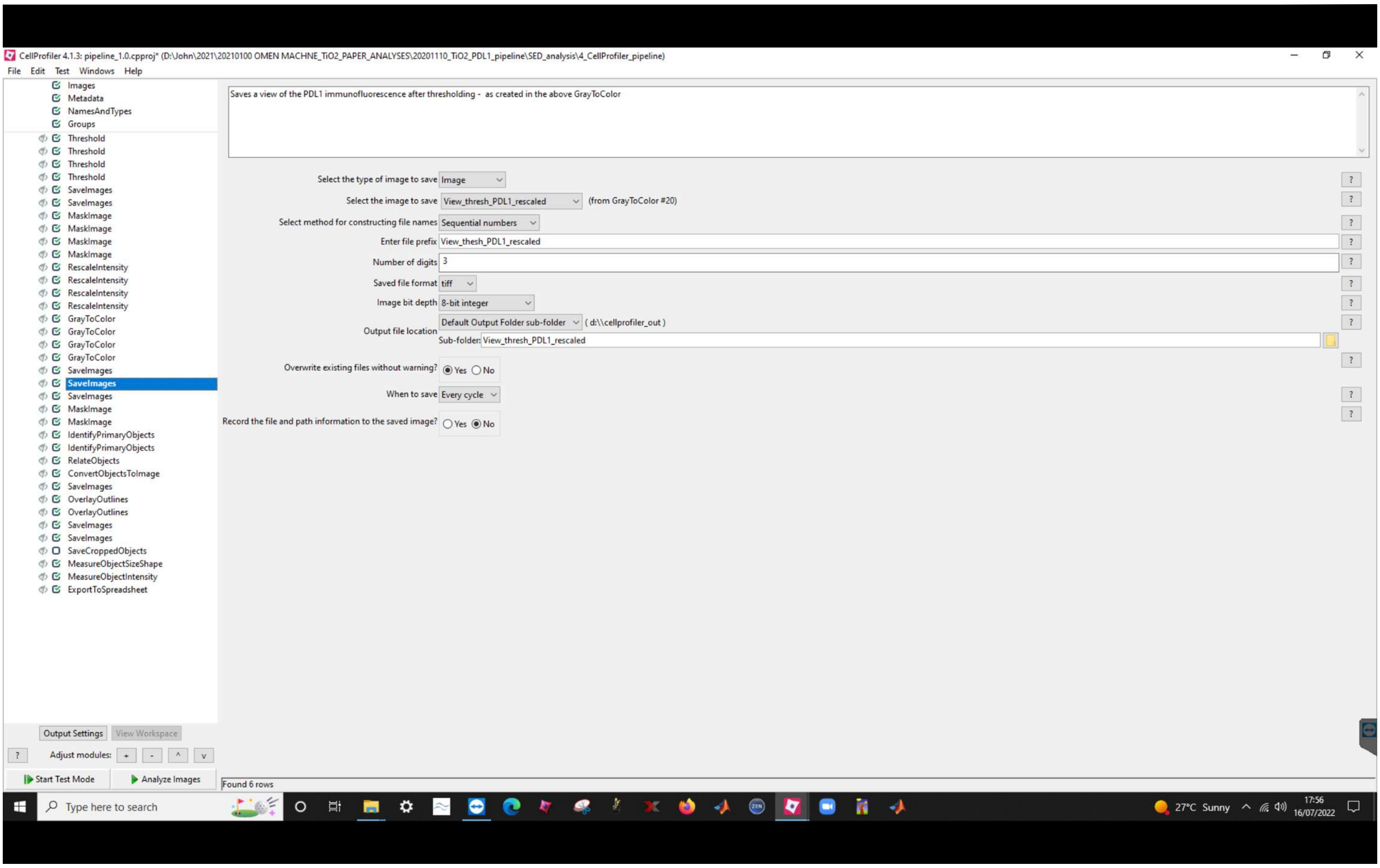
CellProfiler 4.1.3: pipeline_1.0.cpproj* (D:\John\2021\20210100 OMEN MACHNE_TiO2_PAPER_ANALYSES\20201110_TiO2_PDL1_pipeline\SED_analysis\4_CellProfiler_pipeline)
File Edit Test Windows Help
Images
Metadata
NamesAndTypes
Groups
Threshold
Threshold
Threshold
Threshold
SaveImages
SaveImages
MaskImage
MaskImage
MaskImage
MaskImage
RescaleIntensity
RescaleIntensity
RescaleIntensity
RescaleIntensity
GrayToColor
GrayToColor
GrayToColor
GrayToColor
SaveImages
SaveImages
SaveImages
MaskImage
MaskImage
IdentifyPrimaryObjects
IdentifyPrimaryObjects
RelateObjects
ConvertObjectsToImage
SaveImages
OverlayOutlines
OverlayOutlines
SaveImages
SaveImages
SaveCroppedObjects
MeasureObjectSizeShape
MeasureObjectIntensity
ExportToSpreadsheet
Saves a view of the PDL1 immunofluorescence after thresholding - as created in the above GrayToColor
Select the type of image to save Image
Select the image to save View_thresh_PDL1_rescaled (from GrayToColor #20)
Select method for constructing file names Sequential numbers
Enter file prefix View_thesh_PDL1_rescaled
Number of digits 3
Saved file format tiff
Image bit depth 8-bit integer
Output file location Default Output Folder sub-folder ( d:\\cellprofiler_out )
Sub-folder: View_thresh_PDL1_rescaled
Overwrite existing files without warning? Yes No
When to save Every cycle
Record the file and path information to the saved image? Yes No
Output Settings
View Workspace
Adjust modules: + - ^ v
Start Test Mode
Analyze Images
Found 6 rows
Type here to search
27°C Sunny
17:56
16/07/2022

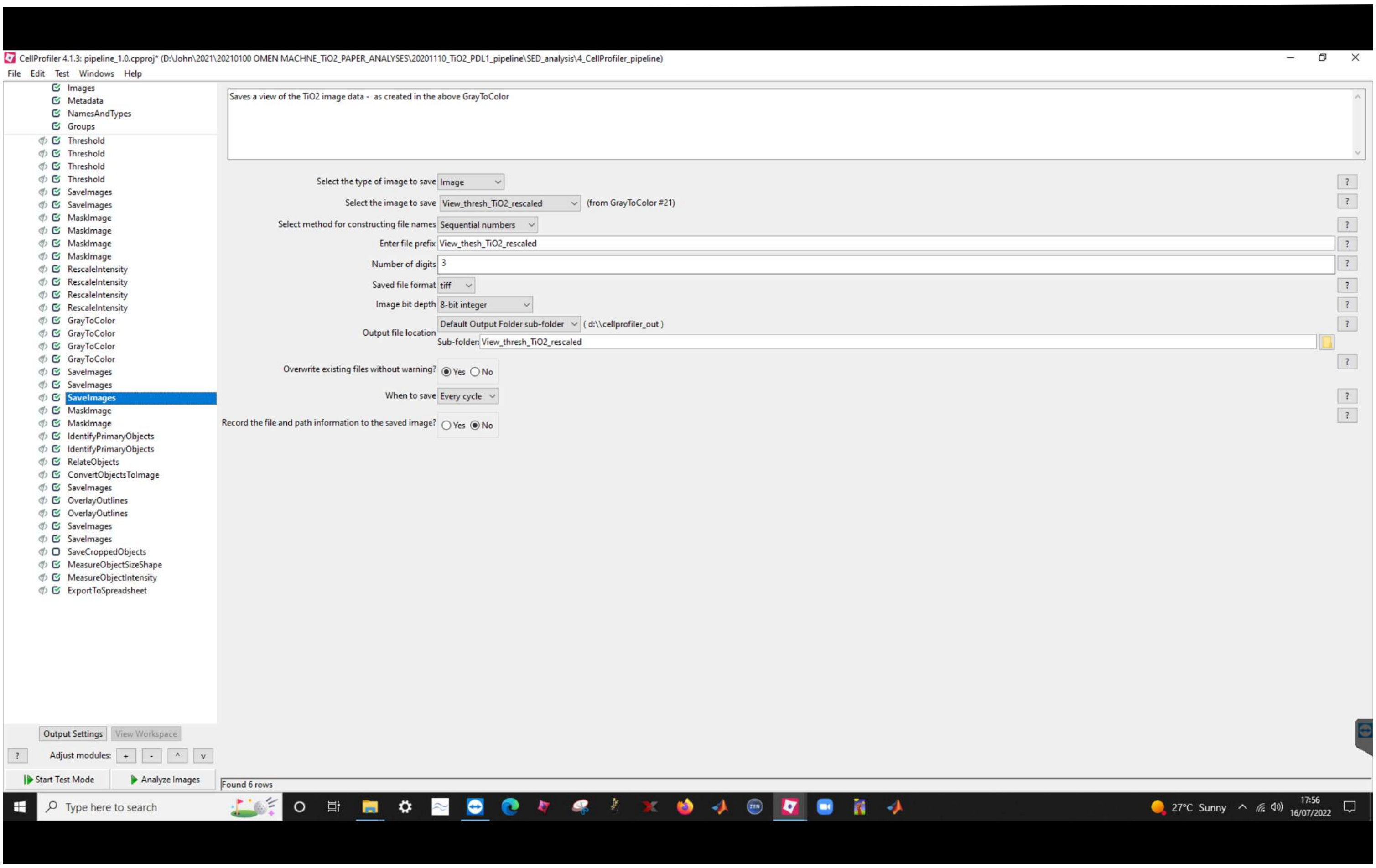
CellProfiler 4.1.3: pipeline_1.0.cpproj* (D:\John\2021\20210100 OMEN MACHNE_TiO2_PAPER_ANALYSES\20201110_TiO2_PDL1_pipeline\SED_analysis\4_CellProfiler_pipeline)
File Edit Test Windows Help
Images
Metadata
NamesAndTypes
Groups
Threshold
Threshold
Threshold
Threshold
SaveImages
SaveImages
MaskImage
MaskImage
MaskImage
MaskImage
RescaleIntensity
RescaleIntensity
RescaleIntensity
RescaleIntensity
GrayToColor
GrayToColor
GrayToColor
GrayToColor
SaveImages
SaveImages
SaveImages
MaskImage
MaskImage
IdentifyPrimaryObjects
IdentifyPrimaryObjects
RelateObjects
ConvertObjectsToImage
SaveImages
OverlayOutlines
OverlayOutlines
SaveImages
SaveImages
SaveCroppedObjects
MeasureObjectSizeShape
MeasureObjectIntensity
ExportToSpreadsheet
Saves a view of the TiO2 image data - as created in the above GrayToColor
Select the type of image to save: Image
Select the image to save: View_thresh_TiO2_rescaled (from GrayToColor #21)
Select method for constructing file names: Sequential numbers
Enter file prefix: View_thesh_TiO2_rescaled
Number of digits: 3
Saved file format: tiff
Image bit depth: 8-bit integer
Output file location: Default Output Folder sub-folder ( d:\\cellprofiler_out )
Sub-folder: View_thresh_TiO2_rescaled
Overwrite existing files without warning? Yes No
When to save: Every cycle
Record the file and path information to the saved image? Yes No
Output Settings
View Workspace
Adjust modules: + - ^ v
Start Test Mode
Analyze Images
Found 6 rows
Type here to search
27°C Sunny
17:56
16/07/2022

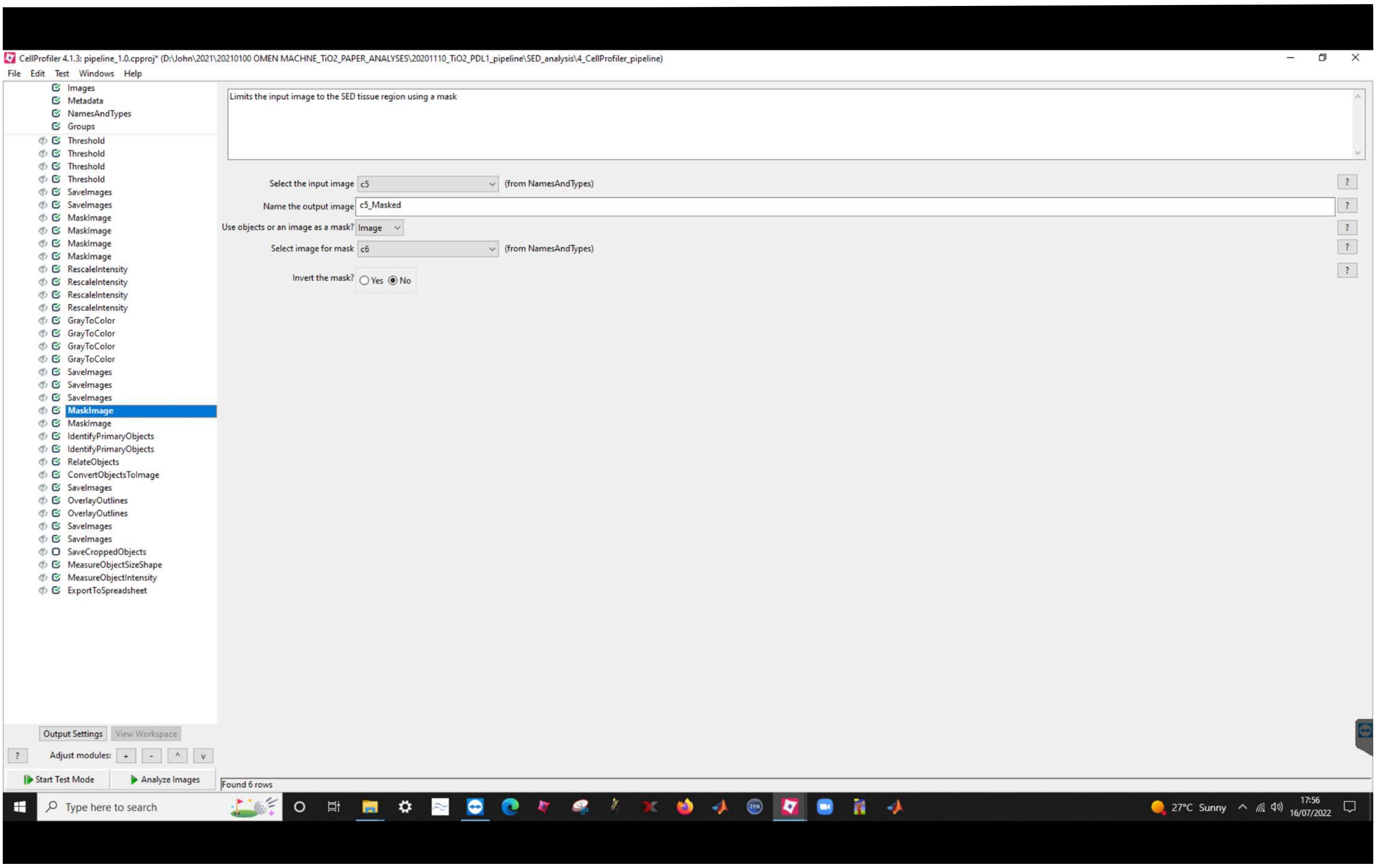
CellProfiler 4.1.3: pipeline_1.0.cpproj* (D:\John\2021\20210100 OMEN MACHNE_TiO2_PAPER_ANALYSES\20201110_TiO2_PDL1_pipeline\SED_analysis\4_CellProfiler_pipeline)
File Edit Test Windows Help
Images
Metadata
NamesAndTypes
Groups
Threshold
Threshold
Threshold
Threshold
SaveImages
SaveImages
MaskImage
MaskImage
MaskImage
MaskImage
RescaleIntensity
RescaleIntensity
RescaleIntensity
RescaleIntensity
GrayToColor
GrayToColor
GrayToColor
GrayToColor
SaveImages
SaveImages
SaveImages
MaskImage
MaskImage
IdentifyPrimaryObjects
IdentifyPrimaryObjects
RelateObjects
ConvertObjectsToImage
SaveImages
OverlayOutlines
OverlayOutlines
SaveImages
SaveImages
SaveCroppedObjects
MeasureObjectSizeShape
MeasureObjectIntensity
ExportToSpreadsheet
Limits the input image to the SED tissue region using a mask
Select the input image c5 (from NamesAndTypes)
Name the output image c5_Masked
Use objects or an image as a mask? Image
Select image for mask c6 (from NamesAndTypes)
Invert the mask? Yes No
Output Settings
View Workspace
Adjust modules: + - ^ v
Start Test Mode
Analyze Images
Found 6 rows
Type here to search
27°C Sunny
17:56
16/07/2022

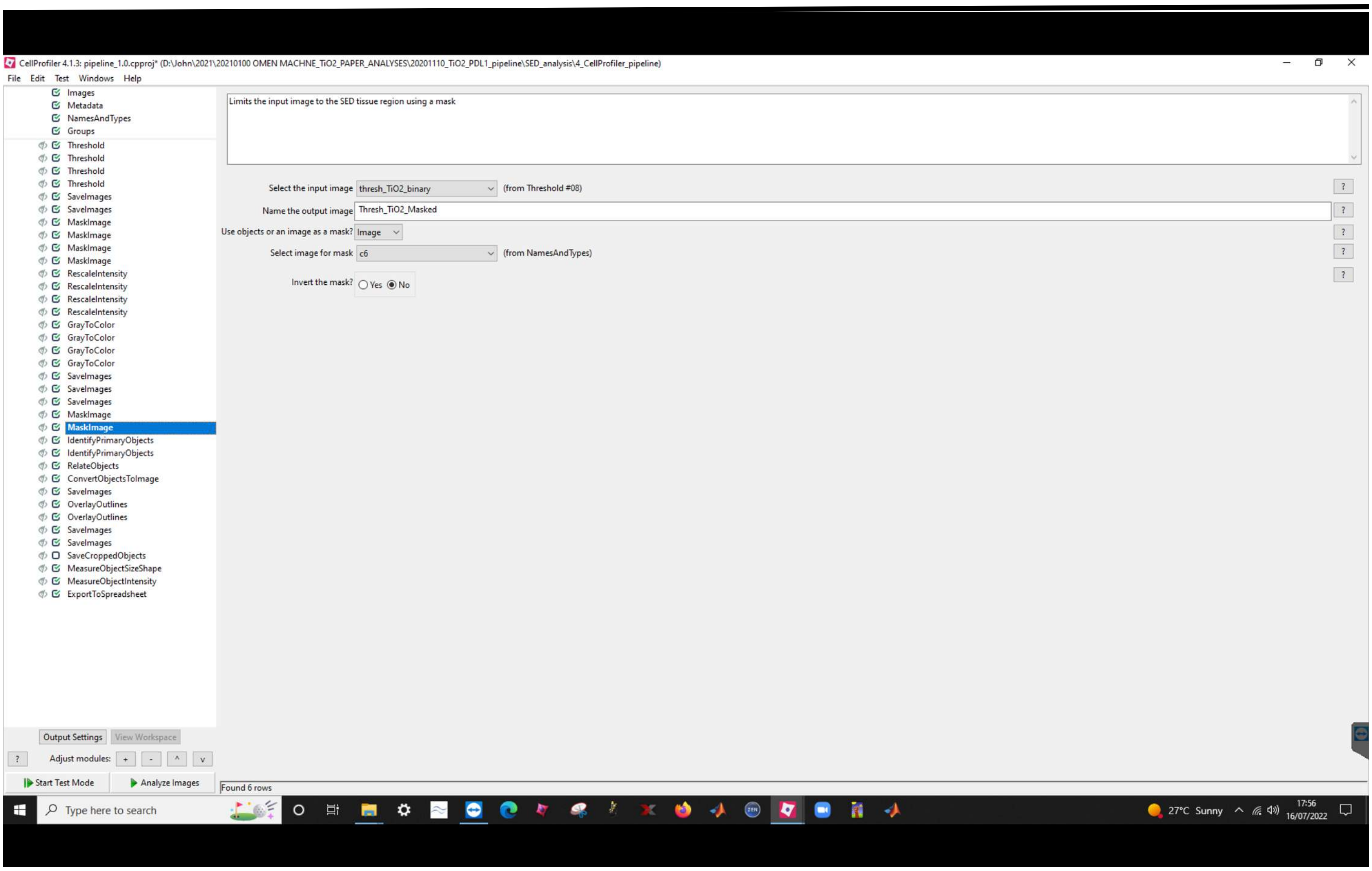
CellProfiler 4.1.3: pipeline_1.0.cpproj* (D:\John\2021\20210100 OMEN MACHNE_TiO2_PAPER_ANALYSES\20201110_TiO2_PDL1_pipeline\SED_analysis\4_CellProfiler_pipeline)
File Edit Test Windows Help
Images
Metadata
NamesAndTypes
Groups
Threshold
Threshold
Threshold
Threshold
SaveImages
SaveImages
MaskImage
MaskImage
MaskImage
MaskImage
RescaleIntensity
RescaleIntensity
RescaleIntensity
RescaleIntensity
GrayToColor
GrayToColor
GrayToColor
GrayToColor
SaveImages
SaveImages
SaveImages
MaskImage
MaskImage
IdentifyPrimaryObjects
IdentifyPrimaryObjects
RelateObjects
ConvertObjectsToImage
SaveImages
OverlayOutlines
OverlayOutlines
SaveImages
SaveImages
SaveCroppedObjects
MeasureObjectSizeShape
MeasureObjectIntensity
ExportToSpreadsheet
Limits the input image to the SED tissue region using a mask
Select the input image thresh_TiO2_binary (from Threshold #08)
Name the output image Thresh_TiO2_Masked
Use objects or an image as a mask? Image
Select image for mask c6 (from NamesAndTypes)
Invert the mask? Yes No
Output Settings
View Workspace
Adjust modules:
Start Test Mode
Analyze Images
Found 6 rows
Type here to search
27°C Sunny
17:56
16/07/2022

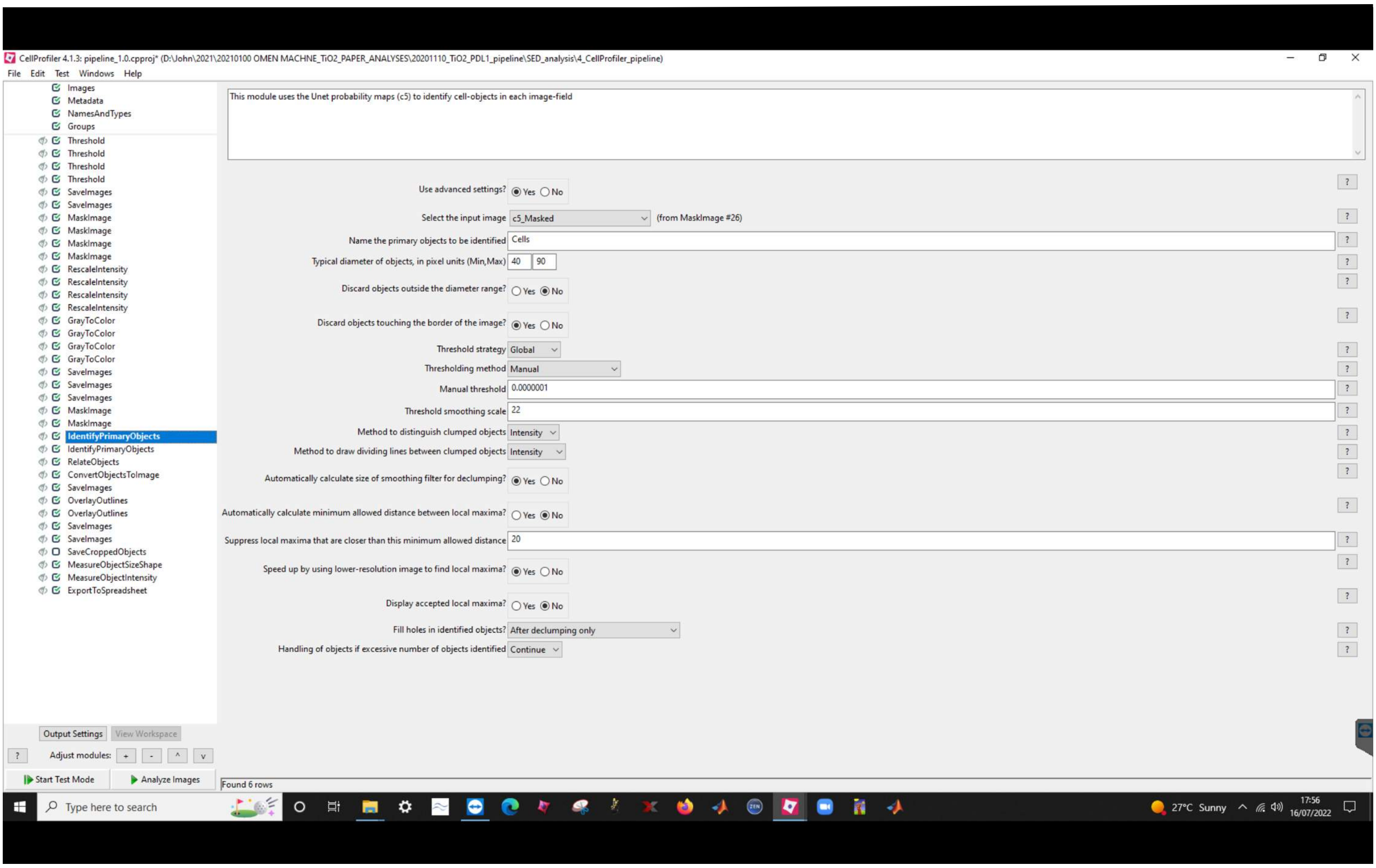
CellProfiler 4.1.3: pipeline_1.0.cpproj* (D:\John\2021\20210100 OMEN MACHNE_TiO2_PAPER_ANALYSES\20201110_TiO2_PDL1_pipeline\SED_analysis\4_CellProfiler_pipeline)
File Edit Test Windows Help
Images
Metadata
NamesAndTypes
Groups
Threshold
Threshold
Threshold
Threshold
SaveImages
SaveImages
MaskImage
MaskImage
MaskImage
MaskImage
RescaleIntensity
RescaleIntensity
RescaleIntensity
RescaleIntensity
GrayToColor
GrayToColor
GrayToColor
GrayToColor
SaveImages
SaveImages
SaveImages
MaskImage
MaskImage
IdentifyPrimaryObjects
IdentifyPrimaryObjects
RelateObjects
ConvertObjectsToImage
SaveImages
OverlayOutlines
OverlayOutlines
SaveImages
SaveImages
SaveCroppedObjects
MeasureObjectSizeShape
MeasureObjectIntensity
ExportToSpreadsheet
This module uses the Unet probability maps (c5) to identify cell-objects in each image-field
Use advanced settings? Yes No
Select the input image c5_Masked (from MaskImage #26)
Name the primary objects to be identified Cells
Typical diameter of objects, in pixel units (Min,Max) 40 90
Discard objects outside the diameter range? Yes No
Discard objects touching the border of the image? Yes No
Threshold strategy Global
Thresholding method Manual
Manual threshold 0.0000001
Threshold smoothing scale 22
Method to distinguish clumped objects Intensity
Method to draw dividing lines between clumped objects Intensity
Automatically calculate size of smoothing filter for declumping? Yes No
Automatically calculate minimum allowed distance between local maxima? Yes No
Suppress local maxima that are closer than this minimum allowed distance 20
Speed up by using lower-resolution image to find local maxima? Yes No
Display accepted local maxima? Yes No
Fill holes in identified objects? After declumping only
Handling of objects if excessive number of objects identified Continue
Output Settings View Workspace
Adjust modules: + - ^ v
Start Test Mode
Analyze Images
Found 6 rows
Type here to search
27°C Sunny
17:56
16/07/2022

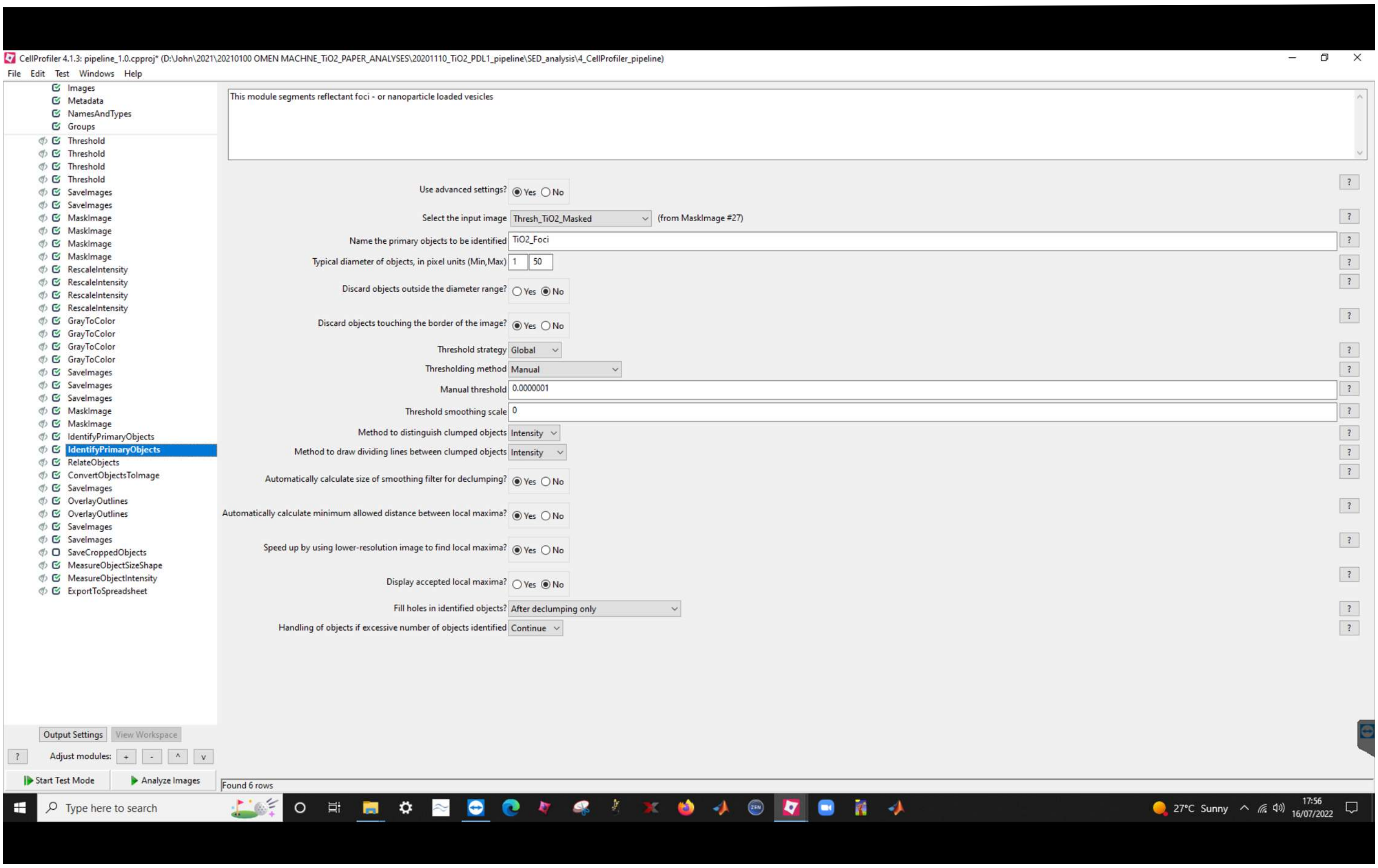
CellProfiler 4.1.3: pipeline_1.0.cpproj* (D:\John\2021\20210100 OMEN MACHNE_TiO2_PAPER_ANALYSES\20201110_TiO2_PDL1_pipeline\SED_analysis\4_CellProfiler_pipeline)
File Edit Test Windows Help
Images
Metadata
NamesAndTypes
Groups
Threshold
Threshold
Threshold
Threshold
SaveImages
SaveImages
MaskImage
MaskImage
MaskImage
MaskImage
RescaleIntensity
RescaleIntensity
RescaleIntensity
RescaleIntensity
GrayToColor
GrayToColor
GrayToColor
GrayToColor
SaveImages
SaveImages
SaveImages
MaskImage
MaskImage
IdentifyPrimaryObjects
IdentifyPrimaryObjects
RelateObjects
ConvertObjectsToImage
SaveImages
OverlayOutlines
OverlayOutlines
SaveImages
SaveImages
SaveCroppedObjects
MeasureObjectSizeShape
MeasureObjectIntensity
ExportToSpreadsheet
This module segments reflectant foci - or nanoparticle loaded vesicles
Use advanced settings? Yes No
Select the input image Thresh_TiO2_Masked (from MaskImage #27)
Name the primary objects to be identified TiO2_Foci
Typical diameter of objects, in pixel units (Min,Max) 1 50
Discard objects outside the diameter range? Yes No
Discard objects touching the border of the image? Yes No
Threshold strategy Global
Thresholding method Manual
Manual threshold 0.0000001
Threshold smoothing scale 0
Method to distinguish clumped objects Intensity
Method to draw dividing lines between clumped objects Intensity
Automatically calculate size of smoothing filter for declumping? Yes No
Automatically calculate minimum allowed distance between local maxima? Yes No
Speed up by using lower-resolution image to find local maxima? Yes No
Display accepted local maxima? Yes No
Fill holes in identified objects? After declumping only
Handling of objects if excessive number of objects identified Continue
Output Settings
View Workspace
Adjust modules:
Start Test Mode
Analyze Images
Found 6 rows
Type here to search
27°C Sunny
17:56
16/07/2022

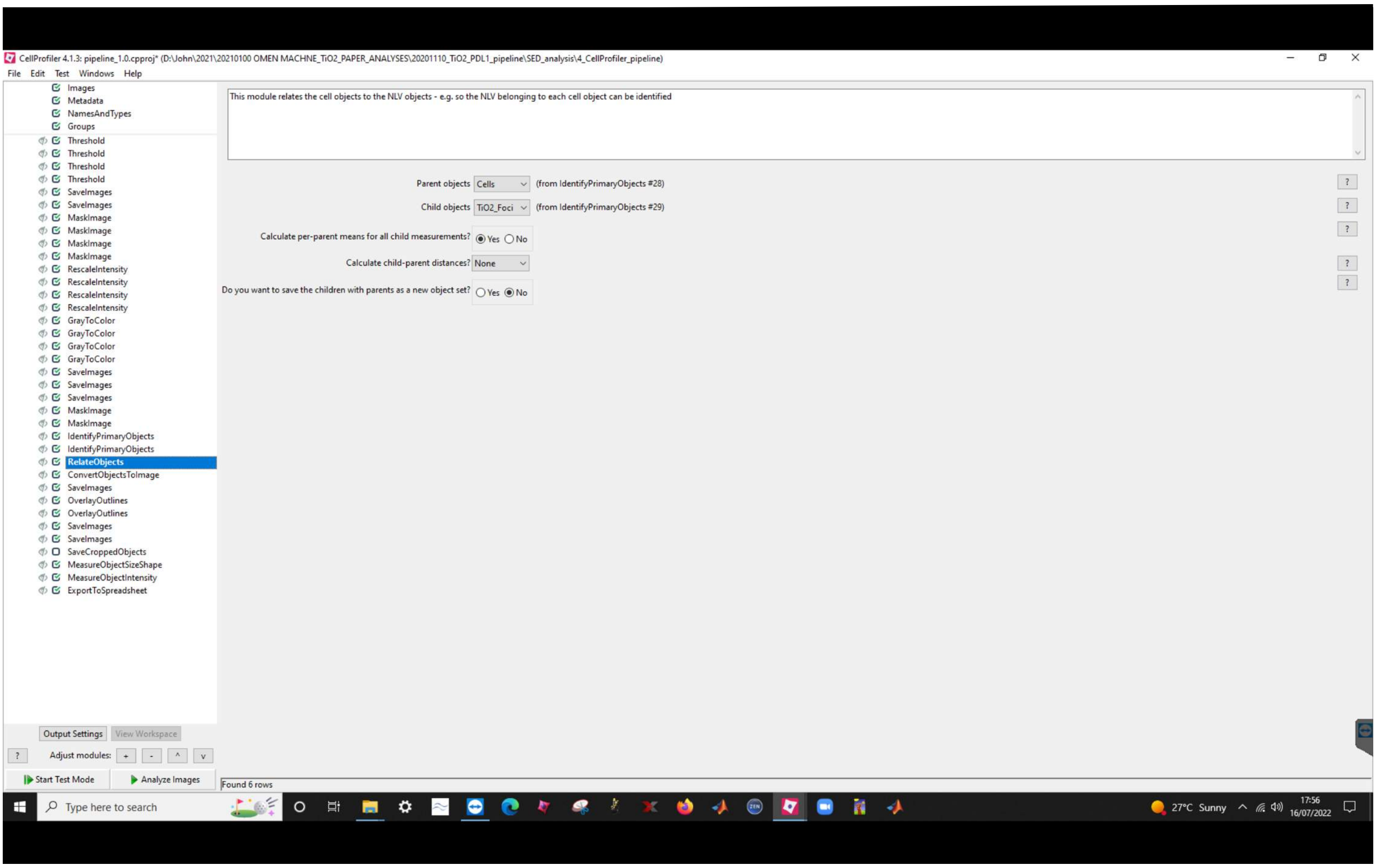
CellProfiler 4.1.3: pipeline_1.0.cpproj* (D:\John\2021\20210100 OMEN MACHNE_TiO2_PAPER_ANALYSES\20201110_TiO2_PDL1_pipeline\SED_analysis\4_CellProfiler_pipeline)
File Edit Test Windows Help
Images
Metadata
NamesAndTypes
Groups
Threshold
Threshold
Threshold
Threshold
SaveImages
SaveImages
MaskImage
MaskImage
MaskImage
MaskImage
RescaleIntensity
RescaleIntensity
RescaleIntensity
RescaleIntensity
GrayToColor
GrayToColor
GrayToColor
GrayToColor
SaveImages
SaveImages
SaveImages
MaskImage
MaskImage
IdentifyPrimaryObjects
IdentifyPrimaryObjects
RelateObjects
ConvertObjectsToImage
SaveImages
OverlayOutlines
OverlayOutlines
SaveImages
SaveImages
SaveCroppedObjects
MeasureObjectSizeShape
MeasureObjectIntensity
ExportToSpreadsheet
This module relates the cell objects to the NLV objects - e.g. so the NLV belonging to each cell object can be identified
Parent objects Cells (from IdentifyPrimaryObjects #28)
Child objects TiO2_Foci (from IdentifyPrimaryObjects #29)
Calculate per-parent means for all child measurements? Yes No
Calculate child-parent distances? None
Do you want to save the children with parents as a new object set? Yes No
Output Settings View Workspace
Adjust modules: + - ^ v
Start Test Mode
Analyze Images
Found 6 rows
Type here to search
27°C Sunny
17:56
16/07/2022

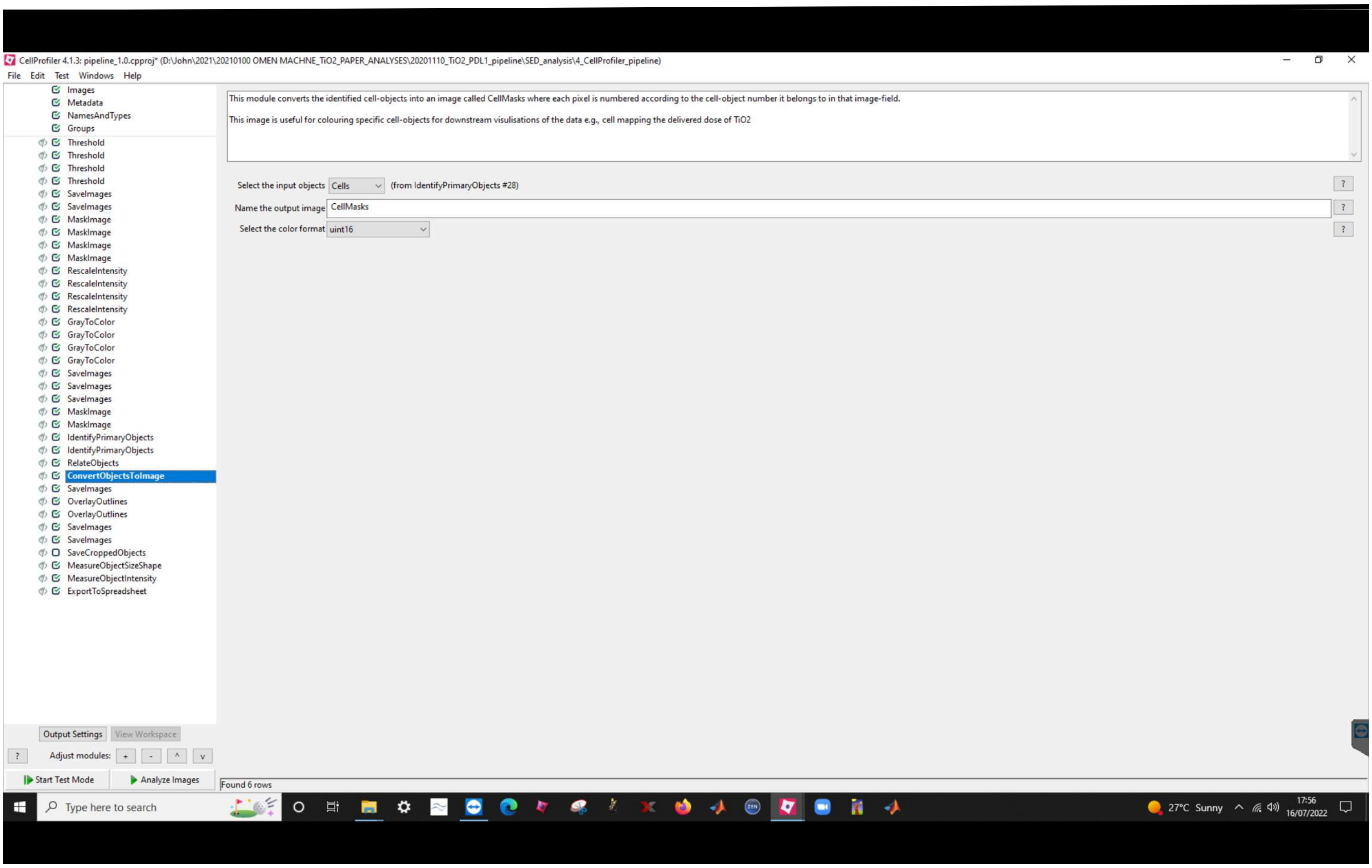
CellProfiler 4.1.3: pipeline_1.0.cpproj* (D:\John\2021\20210100 OMEN MACHNE_TiO2_PAPER_ANALYSES\20201110_TiO2_PDL1_pipeline\SED_analysis\4_CellProfiler_pipeline)
File Edit Test Windows Help
Images
Metadata
NamesAndTypes
Groups
Threshold
Threshold
Threshold
Threshold
SaveImages
SaveImages
MaskImage
MaskImage
MaskImage
MaskImage
RescaleIntensity
RescaleIntensity
RescaleIntensity
RescaleIntensity
GrayToColor
GrayToColor
GrayToColor
GrayToColor
SaveImages
SaveImages
SaveImages
MaskImage
MaskImage
IdentifyPrimaryObjects
IdentifyPrimaryObjects
RelateObjects
ConvertObjectsToImage
SaveImages
OverlayOutlines
OverlayOutlines
SaveImages
SaveImages
SaveCroppedObjects
MeasureObjectSizeShape
MeasureObjectIntensity
ExportToSpreadsheet
This module converts the identified cell-objects into an image called CellMasks where each pixel is numbered according to the cell-object number it belongs to in that image-field.
This image is useful for colouring specific cell-objects for downstream visulisations of the data e.g., cell mapping the delivered dose of TiO2
Select the input objects
Cells
(from IdentifyPrimaryObjects #28)
Name the output image
CellMasks
Select the color format
uint16
Output Settings
View Workspace
Adjust modules:
Start Test Mode
Analyze Images
Found 6 rows
Type here to search
27°C Sunny
17:56
16/07/2022

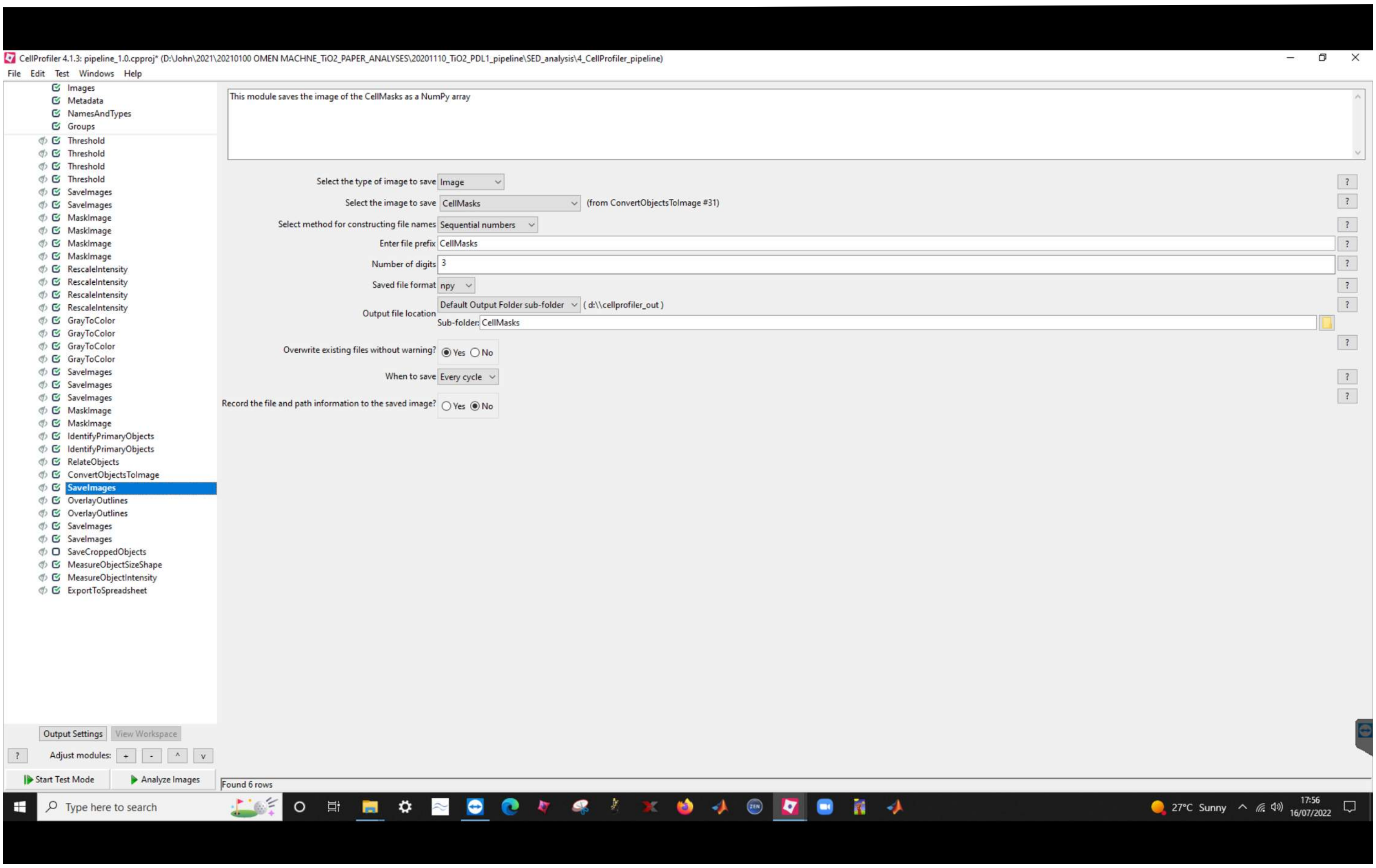
CellProfiler 4.1.3: pipeline_1.0.cpproj* (D:\John\2021\20210100 OMEN MACHNE_TiO2_PAPER_ANALYSES\20201110_TiO2_PDL1_pipeline\SED_analysis\4_CellProfiler_pipeline)
File Edit Test Windows Help
Images
Metadata
NamesAndTypes
Groups
Threshold
Threshold
Threshold
Threshold
SaveImages
SaveImages
MaskImage
MaskImage
MaskImage
MaskImage
RescaleIntensity
RescaleIntensity
RescaleIntensity
RescaleIntensity
GrayToColor
GrayToColor
GrayToColor
GrayToColor
SaveImages
SaveImages
SaveImages
MaskImage
MaskImage
IdentifyPrimaryObjects
IdentifyPrimaryObjects
RelateObjects
ConvertObjectsToImage
SaveImages
OverlayOutlines
OverlayOutlines
SaveImages
SaveImages
SaveCroppedObjects
MeasureObjectSizeShape
MeasureObjectIntensity
ExportToSpreadsheet
This module saves the image of the CellMasks as a NumPy array
Select the type of image to save Image
Select the image to save CellMasks (from ConvertObjectsToImage #31)
Select method for constructing file names Sequential numbers
Enter file prefix CellMasks
Number of digits 3
Saved file format npy
Output file location Default Output Folder sub-folder ( d:\\cellprofiler_out )
Sub-folder: CellMasks
Overwrite existing files without warning? Yes No
When to save Every cycle
Record the file and path information to the saved image? Yes No
Output Settings
View Workspace
Adjust modules: + - ^ v
Start Test Mode
Analyze Images
Found 6 rows
Type here to search
27°C Sunny
17:56
16/07/2022

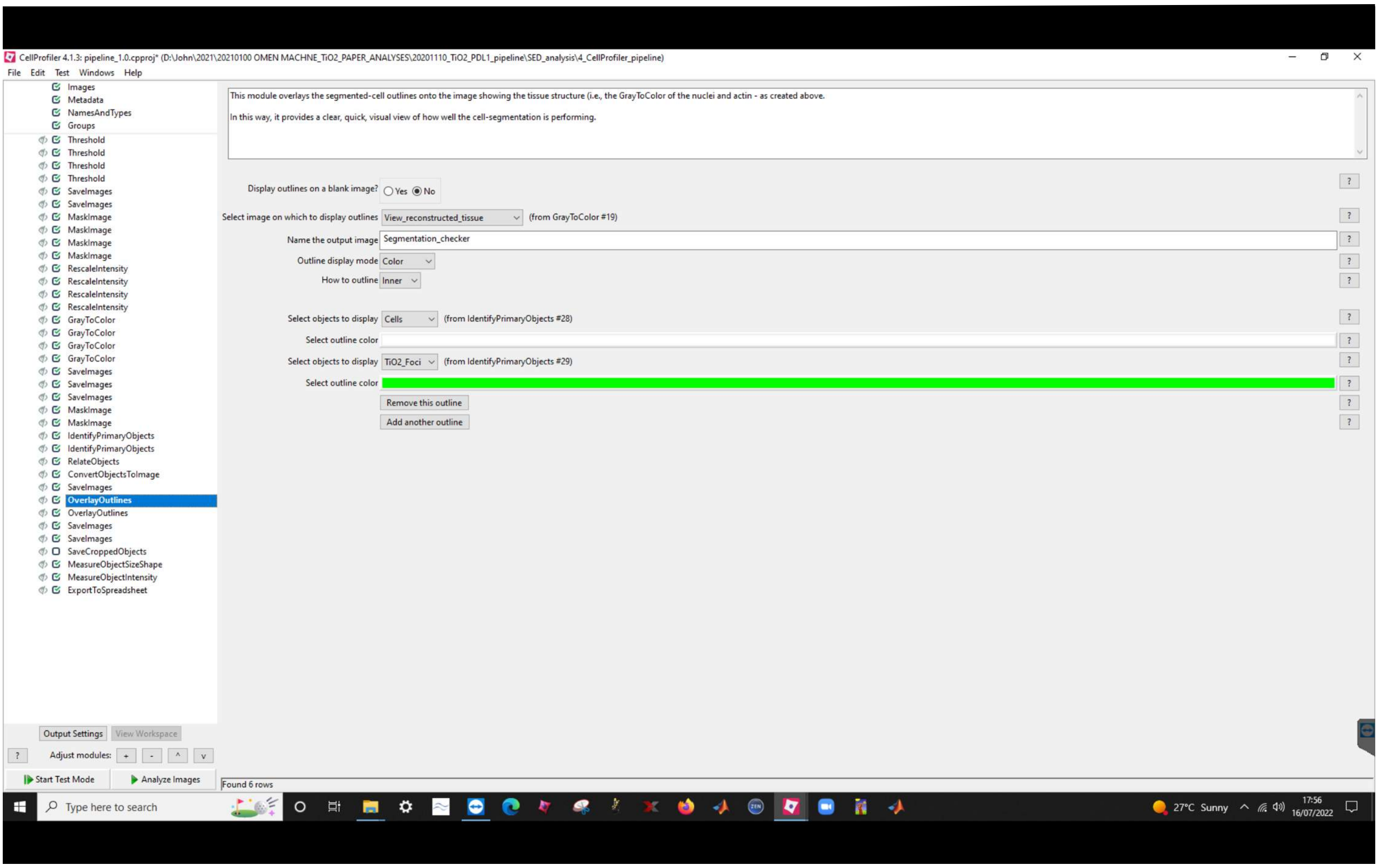
CellProfiler 4.1.3: pipeline_1.0.cpproj* (D:\John\2021\20210100 OMEN MACHNE_TiO2_PAPER_ANALYSES\20201110_TiO2_PDL1_pipeline\SED_analysis\4_CellProfiler_pipeline)
File Edit Test Windows Help
Images
Metadata
NamesAndTypes
Groups
Threshold
Threshold
Threshold
Threshold
SaveImages
SaveImages
MaskImage
MaskImage
MaskImage
MaskImage
RescaleIntensity
RescaleIntensity
RescaleIntensity
RescaleIntensity
GrayToColor
GrayToColor
GrayToColor
GrayToColor
SaveImages
SaveImages
SaveImages
MaskImage
MaskImage
IdentifyPrimaryObjects
IdentifyPrimaryObjects
RelateObjects
ConvertObjectsToImage
SaveImages
OverlayOutlines
OverlayOutlines
SaveImages
SaveImages
SaveCroppedObjects
MeasureObjectSizeShape
MeasureObjectIntensity
ExportToSpreadsheet
This module overlays the segmented-cell outlines onto the image showing the tissue structure (i.e., the GrayToColor of the nuclei and actin - as created above.
In this way, it provides a clear, quick, visual view of how well the cell-segmentation is performing.
Display outlines on a blank image? Yes No
Select image on which to display outlines View_reconstructed_tissue (from GrayToColor #19)
Name the output image Segmentation_checker
Outline display mode Color
How to outline Inner
Select objects to display Cells (from IdentifyPrimaryObjects #28)
Select outline color
Select objects to display TiO2_Foci (from IdentifyPrimaryObjects #29)
Select outline color
Remove this outline
Add another outline
Output Settings
View Workspace
Adjust modules: + - ^ v
Start Test Mode
Analyze Images
Found 6 rows
Type here to search
27°C Sunny
17:56
16/07/2022

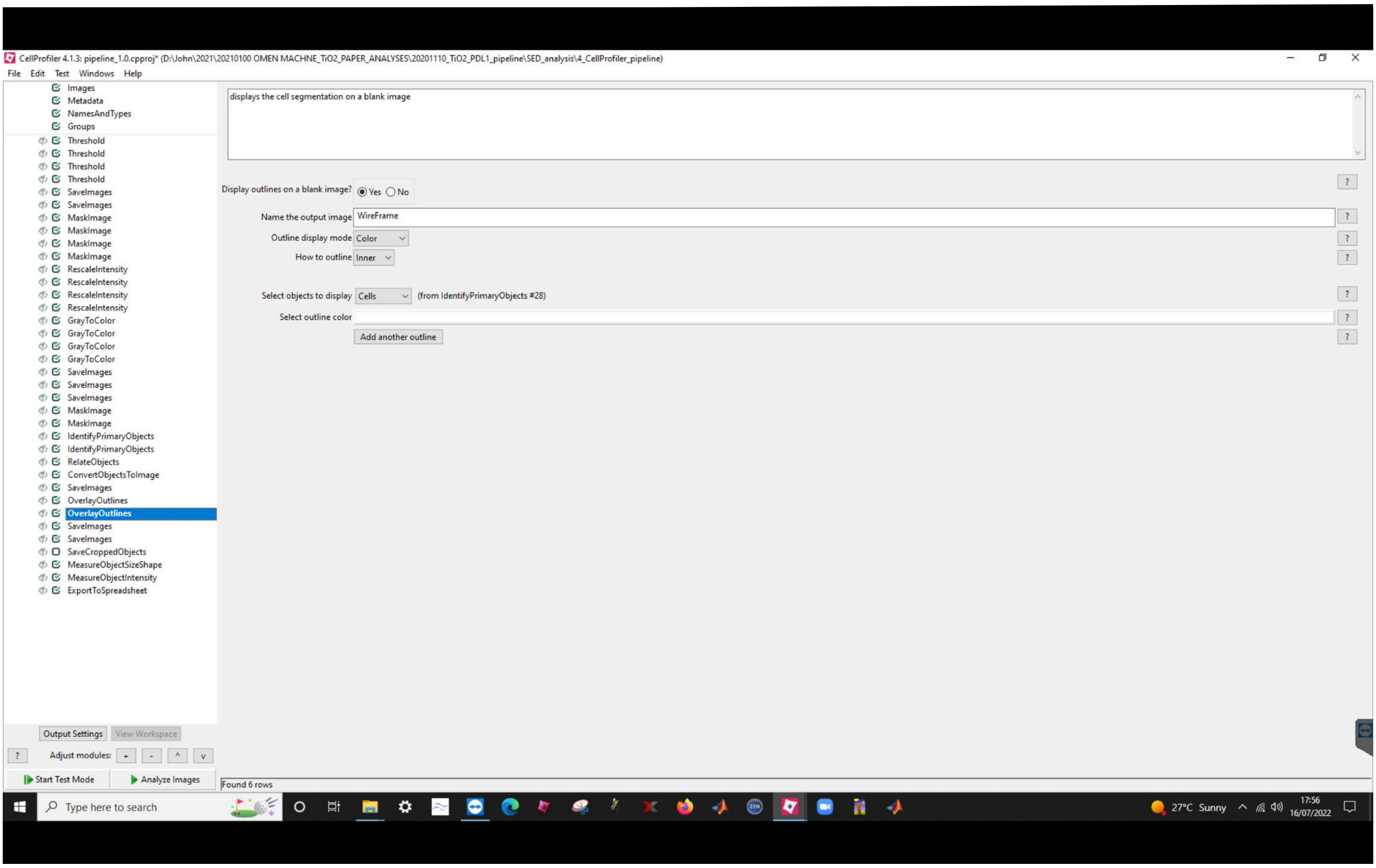
CellProfiler 4.1.3: pipeline_1.0.cpproj* (D:\John\2021\20210100 OMEN MACHNE_TiO2_PAPER_ANALYSES\20201110_TiO2_PDL1_pipeline\SED_analysis\4_CellProfiler_pipeline)
File Edit Test Windows Help
Images
Metadata
NamesAndTypes
Groups
Threshold
Threshold
Threshold
Threshold
SaveImages
SaveImages
MaskImage
MaskImage
MaskImage
MaskImage
RescaleIntensity
RescaleIntensity
RescaleIntensity
RescaleIntensity
GrayToColor
GrayToColor
GrayToColor
GrayToColor
SaveImages
SaveImages
SaveImages
MaskImage
MaskImage
IdentifyPrimaryObjects
IdentifyPrimaryObjects
RelateObjects
ConvertObjectsToImage
SaveImages
OverlayOutlines
OverlayOutlines
SaveImages
SaveImages
SaveCroppedObjects
MeasureObjectSizeShape
MeasureObjectIntensity
ExportToSpreadsheet
displays the cell segmentation on a blank image
Display outlines on a blank image? Yes No
Name the output image WireFrame
Outline display mode Color
How to outline Inner
Select objects to display Cells (from IdentifyPrimaryObjects #28)
Select outline color
Add another outline
Output Settings
View Workspace
Adjust modules: + - ^ v
Start Test Mode
Analyze Images
Found 6 rows
Type here to search
27°C Sunny
17:56
16/07/2022

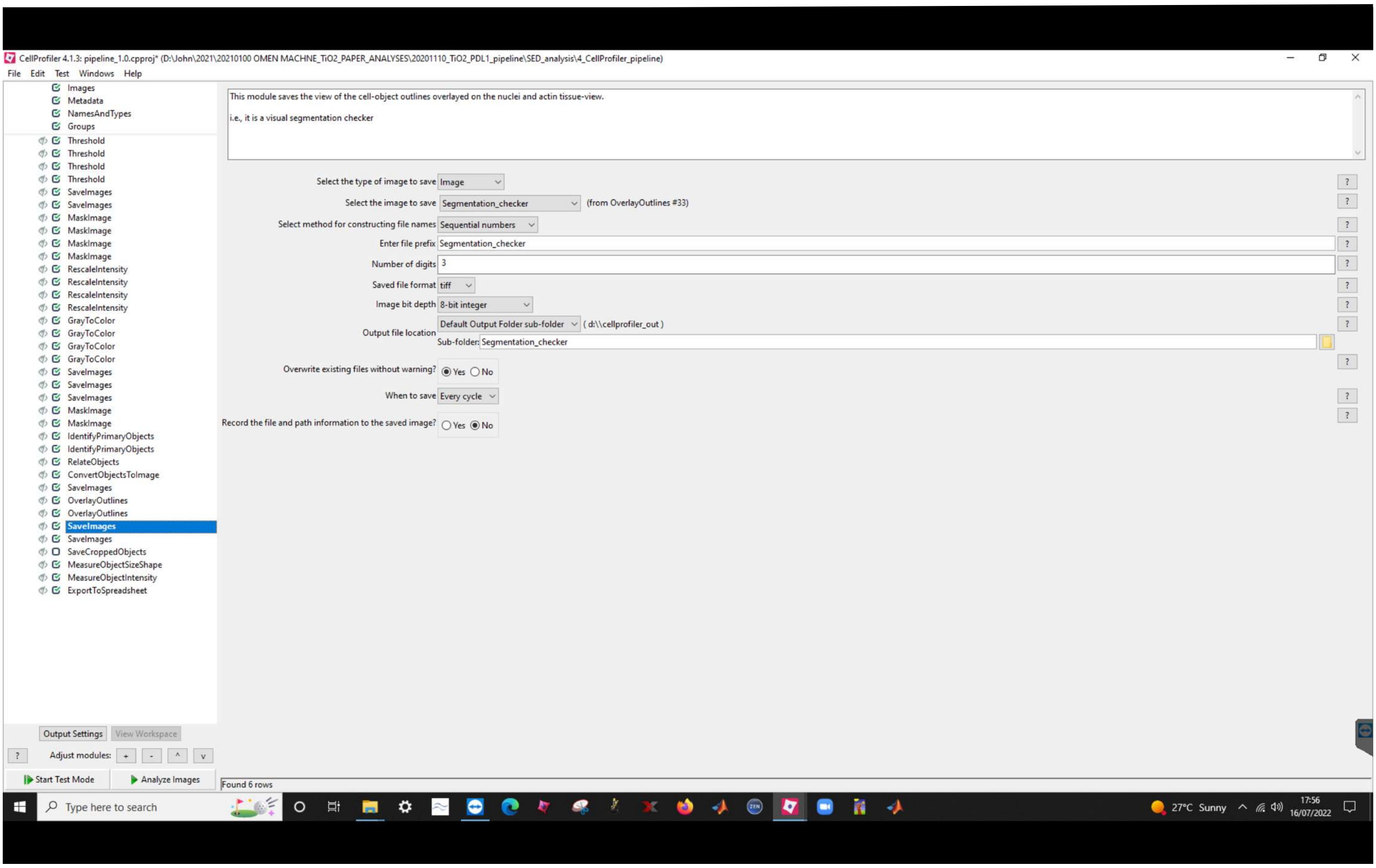
CellProfiler 4.1.3: pipeline_1.0.cpproj* (D:\John\2021\20210100 OMEN MACHNE_TiO2_PAPER_ANALYSES\20201110_TiO2_PDL1_pipeline\SED_analysis\4_CellProfiler_pipeline)
File Edit Test Windows Help
Images
Metadata
NamesAndTypes
Groups
Threshold
Threshold
Threshold
Threshold
SaveImages
SaveImages
MaskImage
MaskImage
MaskImage
MaskImage
RescaleIntensity
RescaleIntensity
RescaleIntensity
RescaleIntensity
GrayToColor
GrayToColor
GrayToColor
GrayToColor
SaveImages
SaveImages
SaveImages
MaskImage
MaskImage
IdentifyPrimaryObjects
IdentifyPrimaryObjects
RelateObjects
ConvertObjectsToImage
SaveImages
OverlayOutlines
OverlayOutlines
SaveImages
SaveImages
SaveCroppedObjects
MeasureObjectSizeShape
MeasureObjectIntensity
ExportToSpreadsheet
This module saves the view of the cell-object outlines overlayed on the nuclei and actin tissue-view.
i.e., it is a visual segmentation checker
Select the type of image to save Image
Select the image to save Segmentation_checker (from OverlayOutlines #33)
Select method for constructing file names Sequential numbers
Enter file prefix Segmentation_checker
Number of digits 3
Saved file format tiff
Image bit depth 8-bit integer
Output file location Default Output Folder sub-folder ( d:\\cellprofiler_out )
Sub-folder: Segmentation_checker
Overwrite existing files without warning? Yes No
When to save Every cycle
Record the file and path information to the saved image? Yes No
Output Settings
View Workspace
Adjust modules:
Start Test Mode
Analyze Images
Found 6 rows
Type here to search
27°C Sunny
17:56
16/07/2022

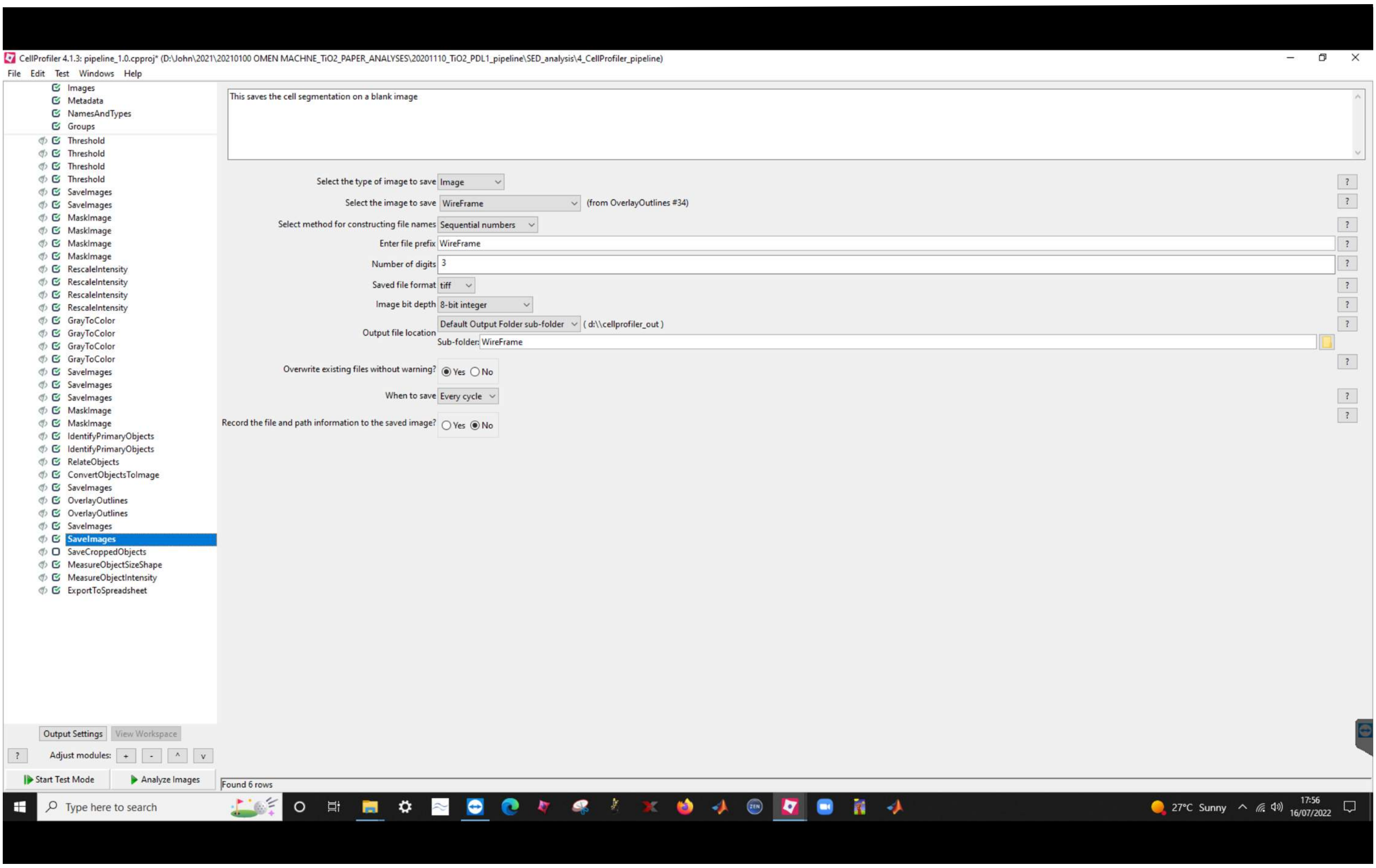
CellProfiler 4.1.3: pipeline_1.0.cpproj* (D:\John\2021\20210100 OMEN MACHNE_TiO2_PAPER_ANALYSES\20201110_TiO2_PDL1_pipeline\SED_analysis\4_CellProfiler_pipeline)
File Edit Test Windows Help
Images
Metadata
NamesAndTypes
Groups
Threshold
Threshold
Threshold
Threshold
SaveImages
SaveImages
MaskImage
MaskImage
MaskImage
MaskImage
RescaleIntensity
RescaleIntensity
RescaleIntensity
RescaleIntensity
GrayToColor
GrayToColor
GrayToColor
GrayToColor
SaveImages
SaveImages
SaveImages
MaskImage
MaskImage
IdentifyPrimaryObjects
IdentifyPrimaryObjects
RelateObjects
ConvertObjectsToImage
SaveImages
OverlayOutlines
OverlayOutlines
SaveImages
SaveImages
SaveCroppedObjects
MeasureObjectSizeShape
MeasureObjectIntensity
ExportToSpreadsheet
This saves the cell segmentation on a blank image
Select the type of image to save Image
Select the image to save WireFrame (from OverlayOutlines #34)
Select method for constructing file names Sequential numbers
Enter file prefix WireFrame
Number of digits 3
Saved file format tiff
Image bit depth 8-bit integer
Output file location Default Output Folder sub-folder ( d:\\cellprofiler_out )
Sub-folder: WireFrame
Overwrite existing files without warning? Yes No
When to save Every cycle
Record the file and path information to the saved image? Yes No
Output Settings
View Workspace
Adjust modules:
Start Test Mode
Analyze Images
Found 6 rows
Type here to search
27°C Sunny
17:56
16/07/2022

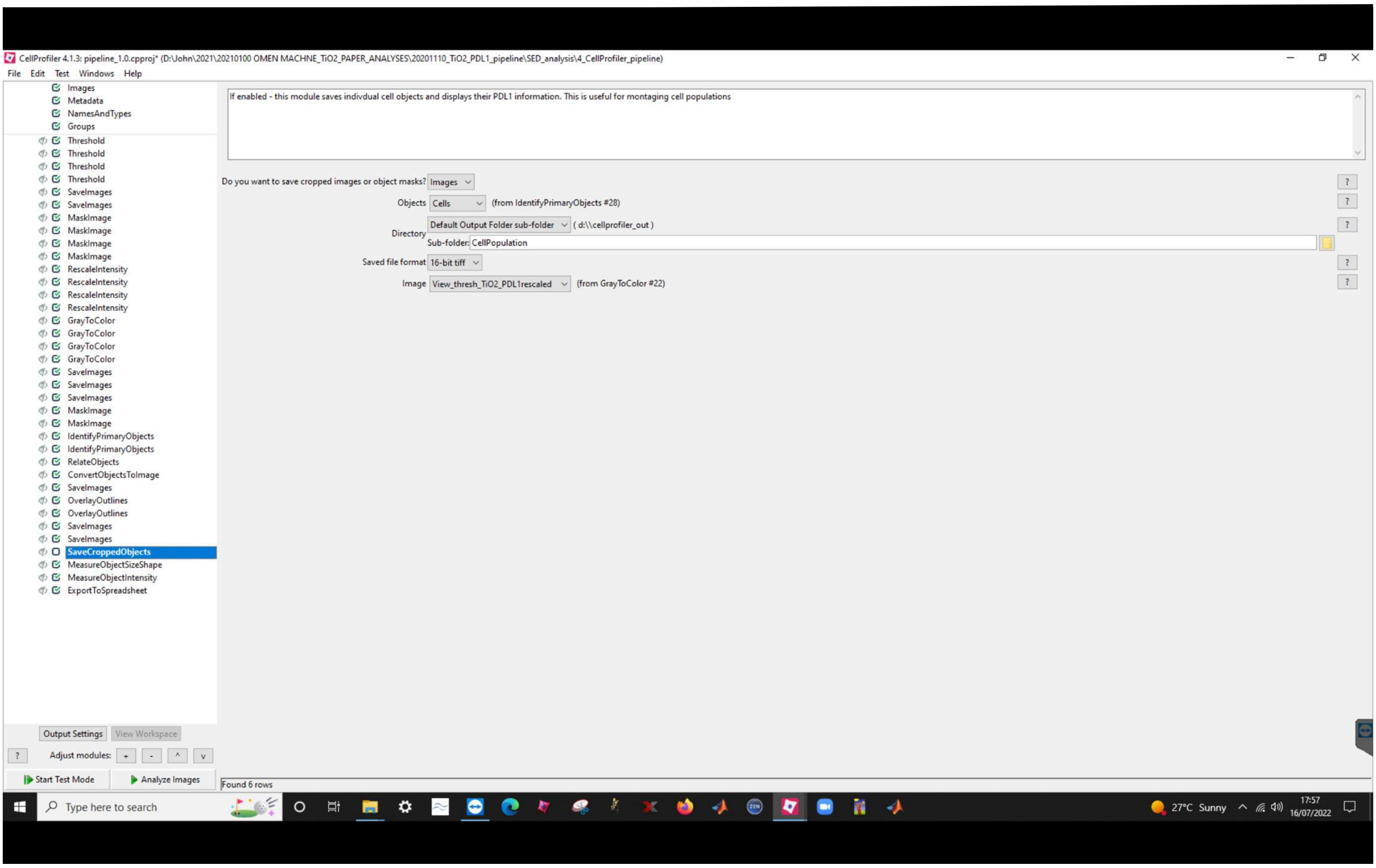
CellProfiler 4.1.3: pipeline_1.0.cpproj* (D:\John\2021\20210100 OMEN MACHNE_TiO2_PAPER_ANALYSES\20201110_TiO2_PDL1_pipeline\SED_analysis\4_CellProfiler_pipeline)
File Edit Test Windows Help
Images
Metadata
NamesAndTypes
Groups
Threshold
Threshold
Threshold
Threshold
SaveImages
SaveImages
MaskImage
MaskImage
MaskImage
MaskImage
RescaleIntensity
RescaleIntensity
RescaleIntensity
RescaleIntensity
GrayToColor
GrayToColor
GrayToColor
GrayToColor
SaveImages
SaveImages
SaveImages
MaskImage
MaskImage
IdentifyPrimaryObjects
IdentifyPrimaryObjects
RelateObjects
ConvertObjectsToImage
SaveImages
OverlayOutlines
OverlayOutlines
SaveImages
SaveImages
SaveCroppedObjects
MeasureObjectSizeShape
MeasureObjectIntensity
ExportToSpreadsheet
If enabled - this module saves indivdual cell objects and displays their PDL1 information. This is useful for montaging cell populations
Do you want to save cropped images or object masks? Images
Objects Cells (from IdentifyPrimaryObjects #28)
Directory Default Output Folder sub-folder ( d:\\cellprofiler_out )
Sub-folder: CellPopulation
Saved file format 16-bit tiff
Image View_thresh_TiO2_PDL1rescaled (from GrayToColor #22)
Output Settings
View Workspace
Adjust modules:
Start Test Mode
Analyze Images
Found 6 rows
Type here to search
27°C Sunny
17:57
16/07/2022

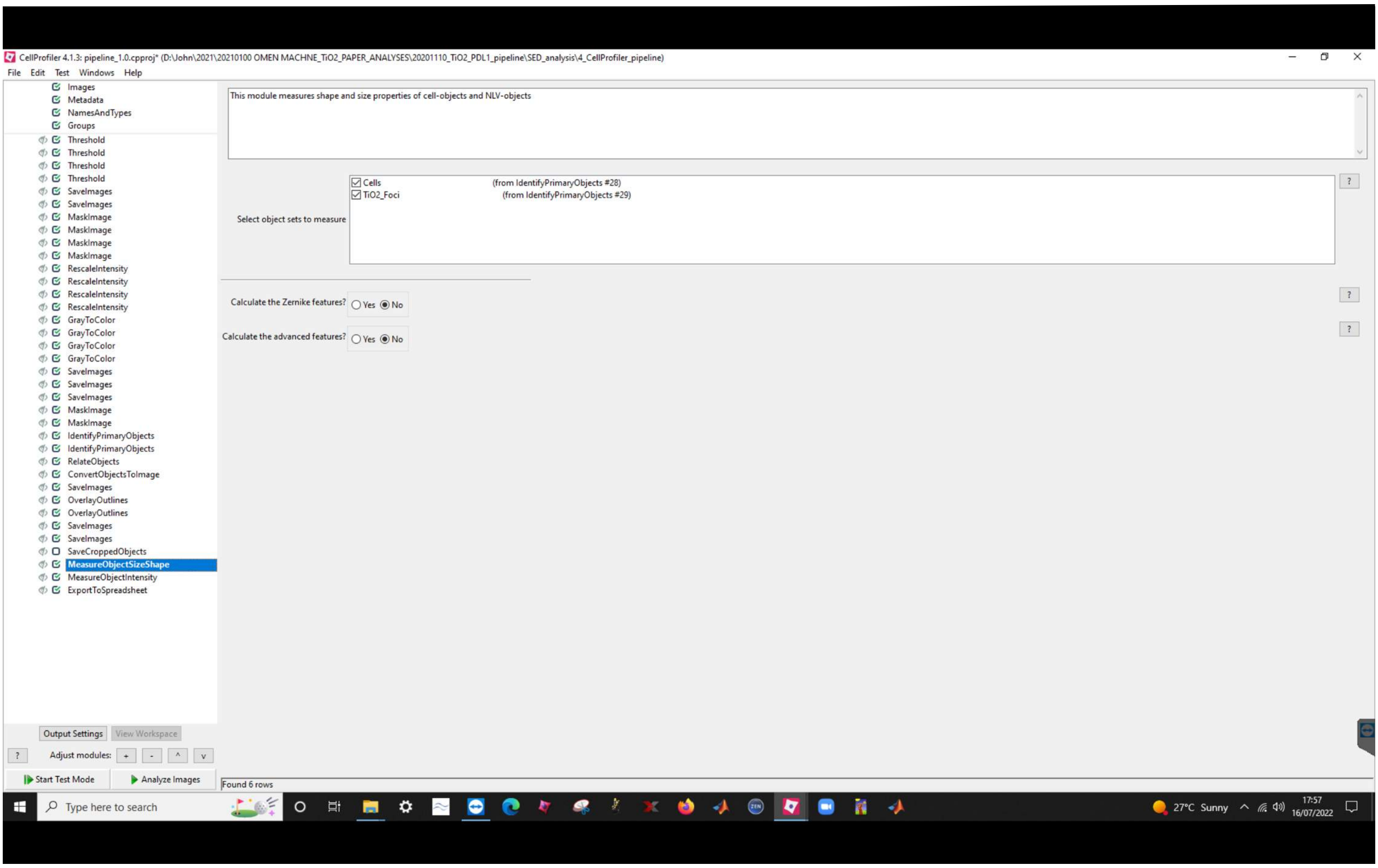
CellProfiler 4.1.3: pipeline_1.0.cpproj* (D:\John\2021\20210100 OMEN MACHNE_TiO2_PAPER_ANALYSES\20201110_TiO2_PDL1_pipeline\SED_analysis\4_CellProfiler_pipeline)
File Edit Test Windows Help
Images
Metadata
NamesAndTypes
Groups
Threshold
Threshold
Threshold
Threshold
SaveImages
SaveImages
MaskImage
MaskImage
MaskImage
MaskImage
RescaleIntensity
RescaleIntensity
RescaleIntensity
RescaleIntensity
GrayToColor
GrayToColor
GrayToColor
GrayToColor
SaveImages
SaveImages
SaveImages
MaskImage
MaskImage
IdentifyPrimaryObjects
IdentifyPrimaryObjects
RelateObjects
ConvertObjectsToImage
SaveImages
OverlayOutlines
OverlayOutlines
SaveImages
SaveImages
SaveCroppedObjects
MeasureObjectSizeShape
MeasureObjectIntensity
ExportToSpreadsheet
This module measures shape and size properties of cell-objects and NLV-objects
Select object sets to measure
Cells (from IdentifyPrimaryObjects #28)
TiO2_Foci (from IdentifyPrimaryObjects #29)
Calculate the Zernike features? Yes No
Calculate the advanced features? Yes No
Output Settings
View Workspace
Adjust modules: + - ^ v
Start Test Mode
Analyze Images
Found 6 rows
Type here to search
27°C Sunny
17:57
16/07/2022

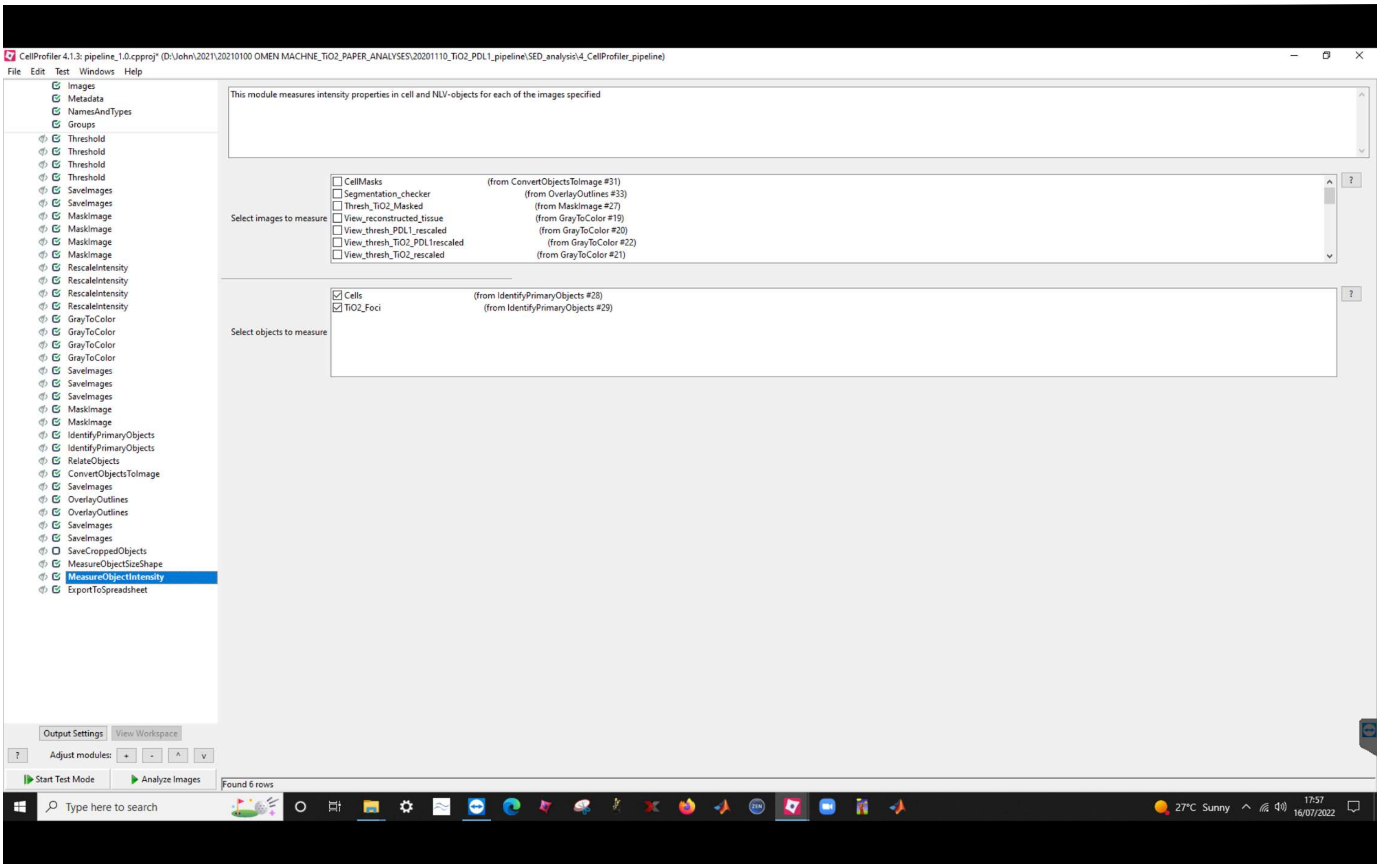
CellProfiler 4.1.3: pipeline_1.0.cpproj* (D:\John\2021\20210100 OMEN MACHNE_TiO2_PAPER_ANALYSES\20201110_TiO2_PDL1_pipeline\SED_analysis\4_CellProfiler_pipeline)
File Edit Test Windows Help
Images
Metadata
NamesAndTypes
Groups
Threshold
Threshold
Threshold
Threshold
SaveImages
SaveImages
MaskImage
MaskImage
MaskImage
MaskImage
RescaleIntensity
RescaleIntensity
RescaleIntensity
RescaleIntensity
GrayToColor
GrayToColor
GrayToColor
GrayToColor
SaveImages
SaveImages
SaveImages
MaskImage
MaskImage
IdentifyPrimaryObjects
IdentifyPrimaryObjects
RelateObjects
ConvertObjectsToImage
SaveImages
OverlayOutlines
OverlayOutlines
SaveImages
SaveImages
SaveCroppedObjects
MeasureObjectSizeShape
MeasureObjectIntensity
ExportToSpreadsheet
This module measures intensity properties in cell and NLV-objects for each of the images specified
Select images to measure
CellMasks (from ConvertObjectsToImage #31)
Segmentation_checker (from OverlayOutlines #33)
Thresh_TiO2_Masked (from MaskImage #27)
View_reconstructed_tissue (from GrayToColor #19)
View_thresh_PDL1_rescaled (from GrayToColor #20)
View_thresh_TiO2_PDL1rescaled (from GrayToColor #22)
View_thresh_TiO2_rescaled (from GrayToColor #21)
Select objects to measure
Cells (from IdentifyPrimaryObjects #28)
TiO2_Foci (from IdentifyPrimaryObjects #29)
Output Settings
View Workspace
Adjust modules:
Start Test Mode
Analyze Images
Found 6 rows
Type here to search
27°C Sunny
17:57
16/07/2022

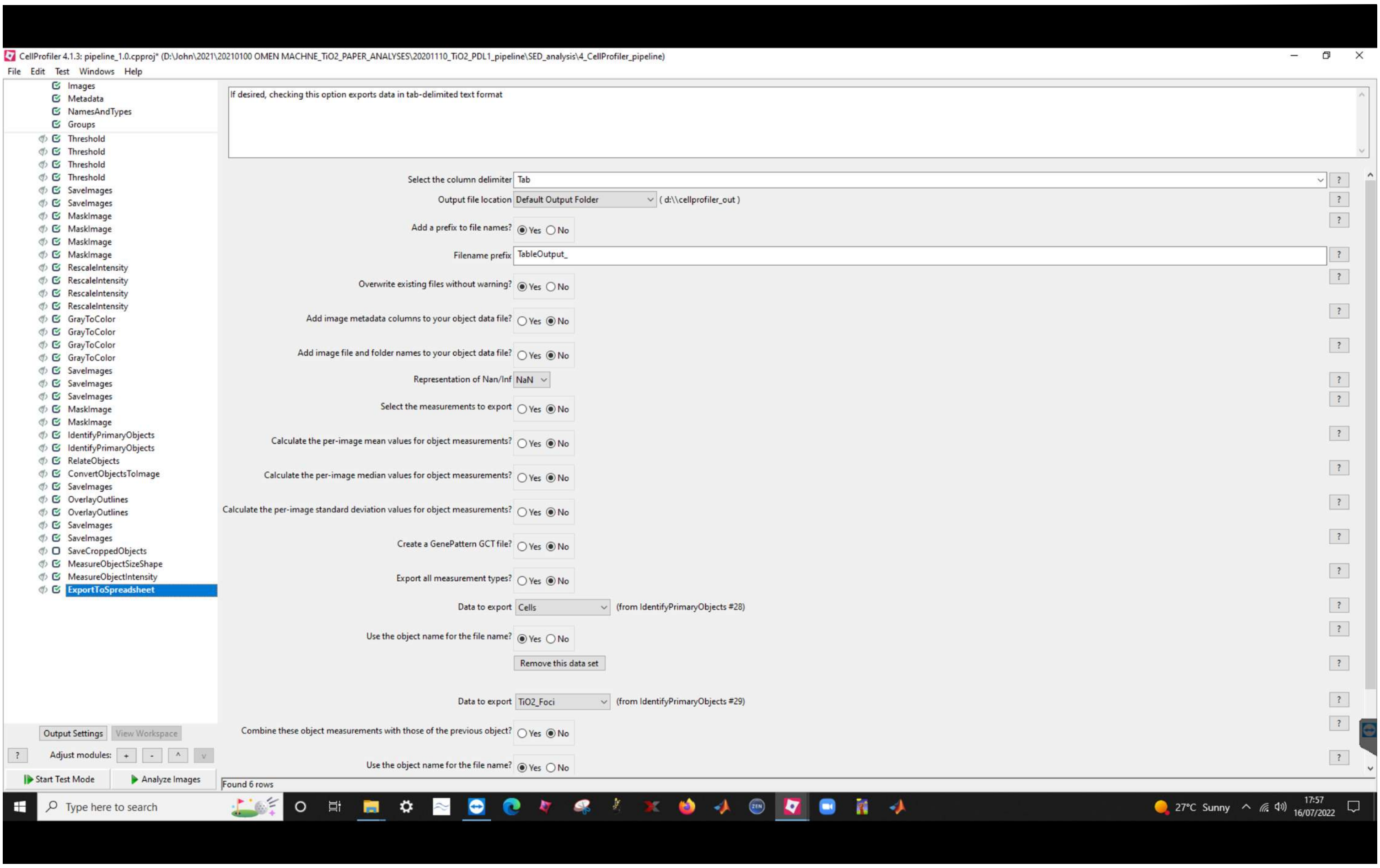
CellProfiler 4.1.3: pipeline_1.0.cpproj* (D:\John\2021\20210100 OMEN MACHNE_TiO2_PAPER_ANALYSES\20201110_TiO2_PDL1_pipeline\SED_analysis\4_CellProfiler_pipeline)
File Edit Test Windows Help
If desired, checking this option exports data in tab-delimited text format
Select the column delimiter Tab
Output file location Default Output Folder ( d:\\cellprofiler_out )
Add a prefix to file names? Yes No
Filename prefix TableOutput_
Overwrite existing files without warning? Yes No
Add image metadata columns to your object data file? Yes No
Add image file and folder names to your object data file? Yes No
Representation of Nan/Inf NaN
Select the measurements to export Yes No
Calculate the per-image mean values for object measurements? Yes No
Calculate the per-image median values for object measurements? Yes No
Calculate the per-image standard deviation values for object measurements? Yes No
Create a GenePattern GCT file? Yes No
Export all measurement types? Yes No
Data to export Cells (from IdentifyPrimaryObjects #28)
Use the object name for the file name? Yes No
Remove this data set
Data to export TiO2_Foci (from IdentifyPrimaryObjects #29)
Combine these object measurements with those of the previous object? Yes No
Use the object name for the file name? Yes No
Images
Metadata
NamesAndTypes
Groups
Threshold
Threshold
Threshold
Threshold
SaveImages
SaveImages
MaskImage
MaskImage
MaskImage
MaskImage
RescaleIntensity
RescaleIntensity
RescaleIntensity
RescaleIntensity
GrayToColor
GrayToColor
GrayToColor
GrayToColor
SaveImages
SaveImages
SaveImages
MaskImage
MaskImage
IdentifyPrimaryObjects
IdentifyPrimaryObjects
RelateObjects
ConvertObjectsToImage
SaveImages
OverlayOutlines
OverlayOutlines
SaveImages
SaveImages
SaveCroppedObjects
MeasureObjectSizeShape
MeasureObjectIntensity
ExportToSpreadsheet
Output Settings
View Workspace
Adjust modules:
Start Test Mode
Analyze Images
Found 6 rows
Type here to search
27°C Sunny
17:57
16/07/2022

**Supplementary Note 3 – Mouse-Human correlative dosimetry pipeline:** This section presents screenshots of the CellProfiler image analysis pipeline used to measure per-cell and TLV-object intensity and shape/size features enabling the correlative mouse-human dosimetry work. The pipeline and sample image-data are available for download from the BioStudies archive accompanying the paper.

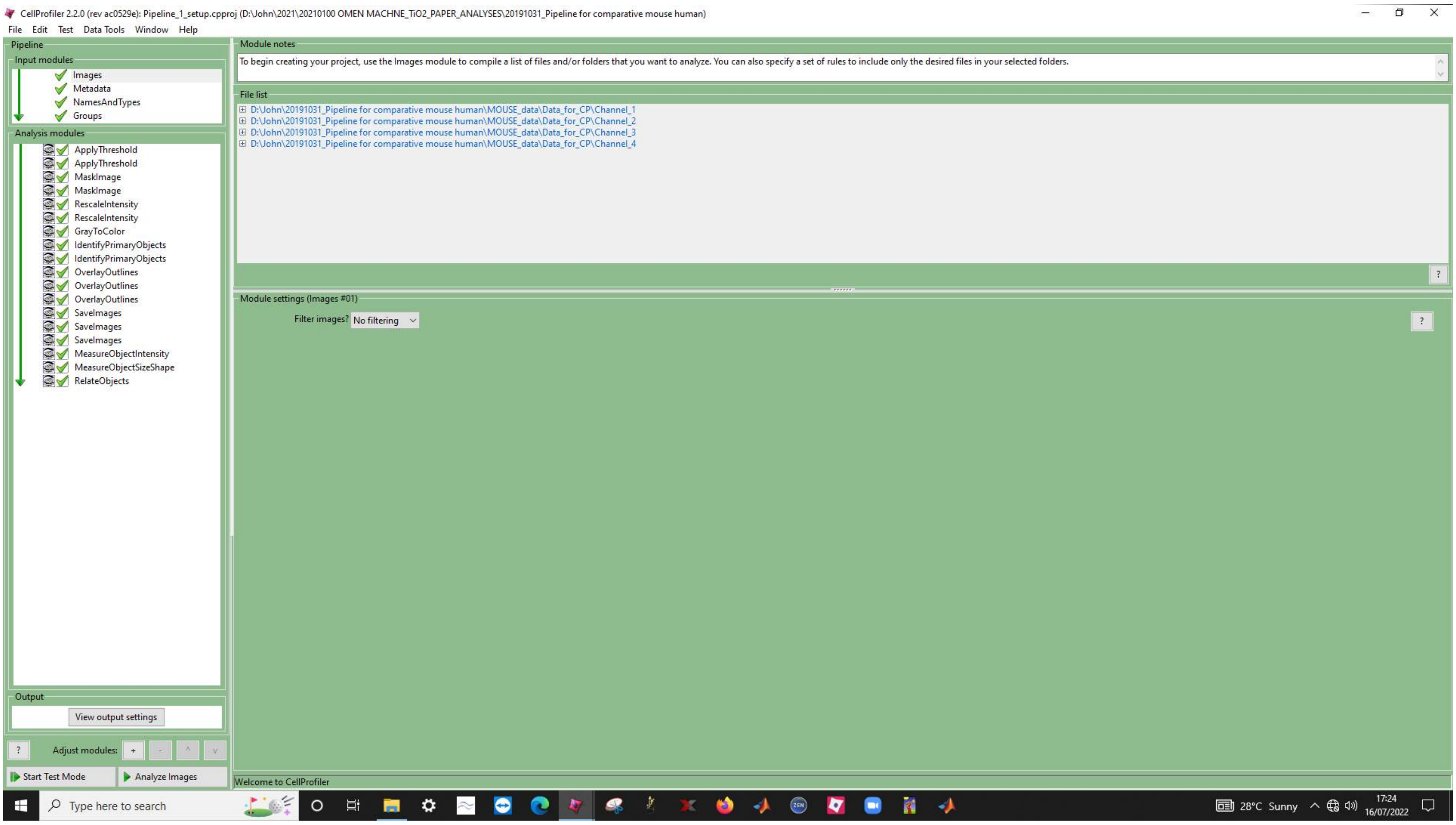

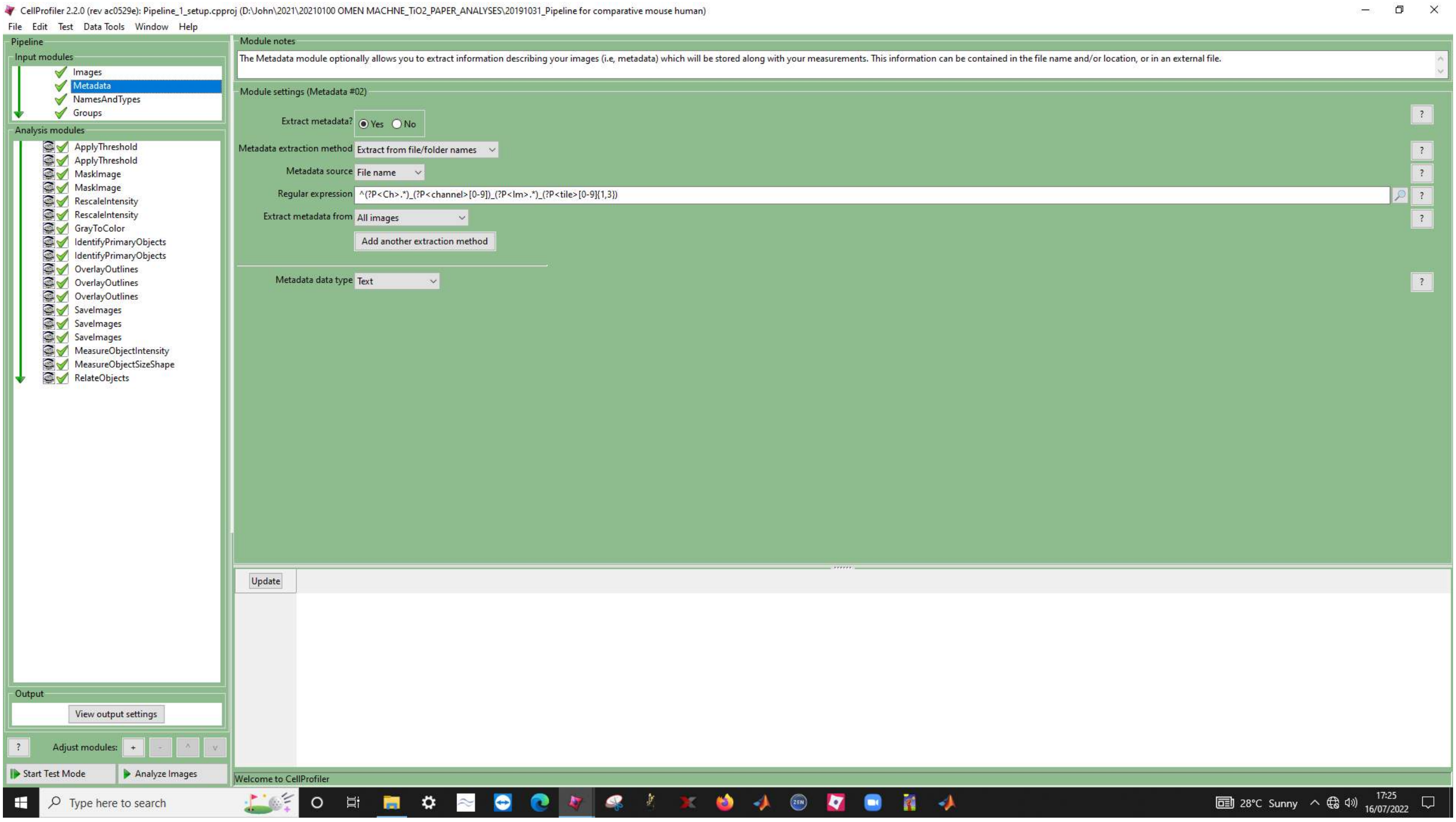
CellProfiler 2.2.0 (rev ac0529e): Pipeline_1_setup.cpproj (D:\John\2021\20210100 OMEN MACHNE_TiO2_PAPER_ANALYSES\20191031_Pipeline for comparative mouse human)
File Edit Test Data Tools Window Help
Pipeline
Input modules
Images
Metadata
NamesAndTypes
Groups
Analysis modules
ApplyThreshold
ApplyThreshold
MaskImage
MaskImage
RescaleIntensity
RescaleIntensity
GrayToColor
IdentifyPrimaryObjects
IdentifyPrimaryObjects
OverlayOutlines
OverlayOutlines
OverlayOutlines
SaveImages
SaveImages
SaveImages
MeasureObjectIntensity
MeasureObjectSizeShape
RelateObjects
Output
View output settings
Adjust modules:
Start Test Mode
Analyze Images
Module notes
The Metadata module optionally allows you to extract information describing your images (i.e, metadata) which will be stored along with your measurements. This information can be contained in the file name and/or location, or in an external file.
Module settings (Metadata #02)
Extract metadata? Yes No
Metadata extraction method Extract from file/folder names
Metadata source File name
Regular expression ^(?P<Ch>.*)_(?P<channel>[0-9])_(?P<Im>.*)_(?P<tile>[0-9]{1,3})
Extract metadata from All images
Add another extraction method
Metadata data type Text
Update
Welcome to CellProfiler
Type here to search
28°C Sunny
17:25
16/07/2022

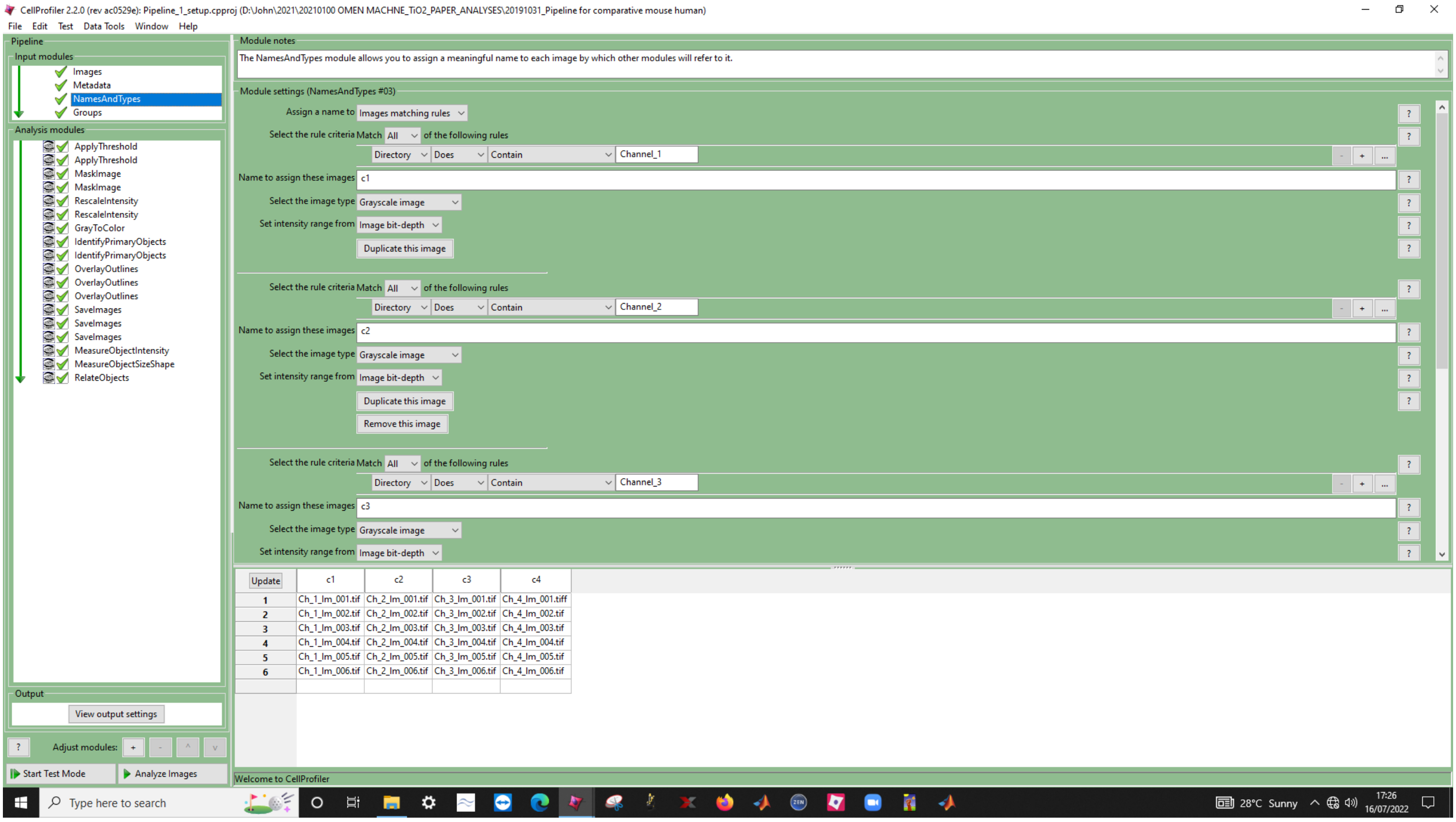
CellProfiler 2.2.0 (rev ac0529e): Pipeline_1_setup.cpproj (D:\John\2021\20210100 OMEN MACHINE_TiO2_PAPER_ANALYSES\20191031_Pipeline for comparative mouse human)
File Edit Test Data Tools Window Help
Pipeline
Input modules
Images
Metadata
NamesAndTypes
Groups
Analysis modules
ApplyThreshold
ApplyThreshold
MaskImage
MaskImage
RescaleIntensity
RescaleIntensity
GrayToColor
IdentifyPrimaryObjects
IdentifyPrimaryObjects
OverlayOutlines
OverlayOutlines
OverlayOutlines
SaveImages
SaveImages
SaveImages
MeasureObjectIntensity
MeasureObjectSizeShape
RelateObjects
Output
View output settings
Adjust modules:
Start Test Mode
Analyze Images
Module notes
The NamesAndTypes module allows you to assign a meaningful name to each image by which other modules will refer to it.
Module settings (NamesAndTypes #03)
Assign a name to Images matching rules
Select the rule criteria Match All of the following rules
Directory Does Contain Channel_1
Name to assign these images c1
Select the image type Grayscale image
Set intensity range from Image bit-depth
Duplicate this image
Select the rule criteria Match All of the following rules
Directory Does Contain Channel_2
Name to assign these images c2
Select the image type Grayscale image
Set intensity range from Image bit-depth
Duplicate this image
Remove this image
Select the rule criteria Match All of the following rules
Directory Does Contain Channel_3
Name to assign these images c3
Select the image type Grayscale image
Set intensity range from Image bit-depth
Update c1 c2 c3 c4
1 Ch_1_Im_001.tif Ch_2_Im_001.tif Ch_3_Im_001.tif Ch_4_Im_001.tiff
2 Ch_1_Im_002.tif Ch_2_Im_002.tif Ch_3_Im_002.tif Ch_4_Im_002.tif
3 Ch_1_Im_003.tif Ch_2_Im_003.tif Ch_3_Im_003.tif Ch_4_Im_003.tif
4 Ch_1_Im_004.tif Ch_2_Im_004.tif Ch_3_Im_004.tif Ch_4_Im_004.tif
5 Ch_1_Im_005.tif Ch_2_Im_005.tif Ch_3_Im_005.tif Ch_4_Im_005.tif
6 Ch_1_Im_006.tif Ch_2_Im_006.tif Ch_3_Im_006.tif Ch_4_Im_006.tif
Welcome to CellProfiler
Type here to search
28°C Sunny
17:26
16/07/2022

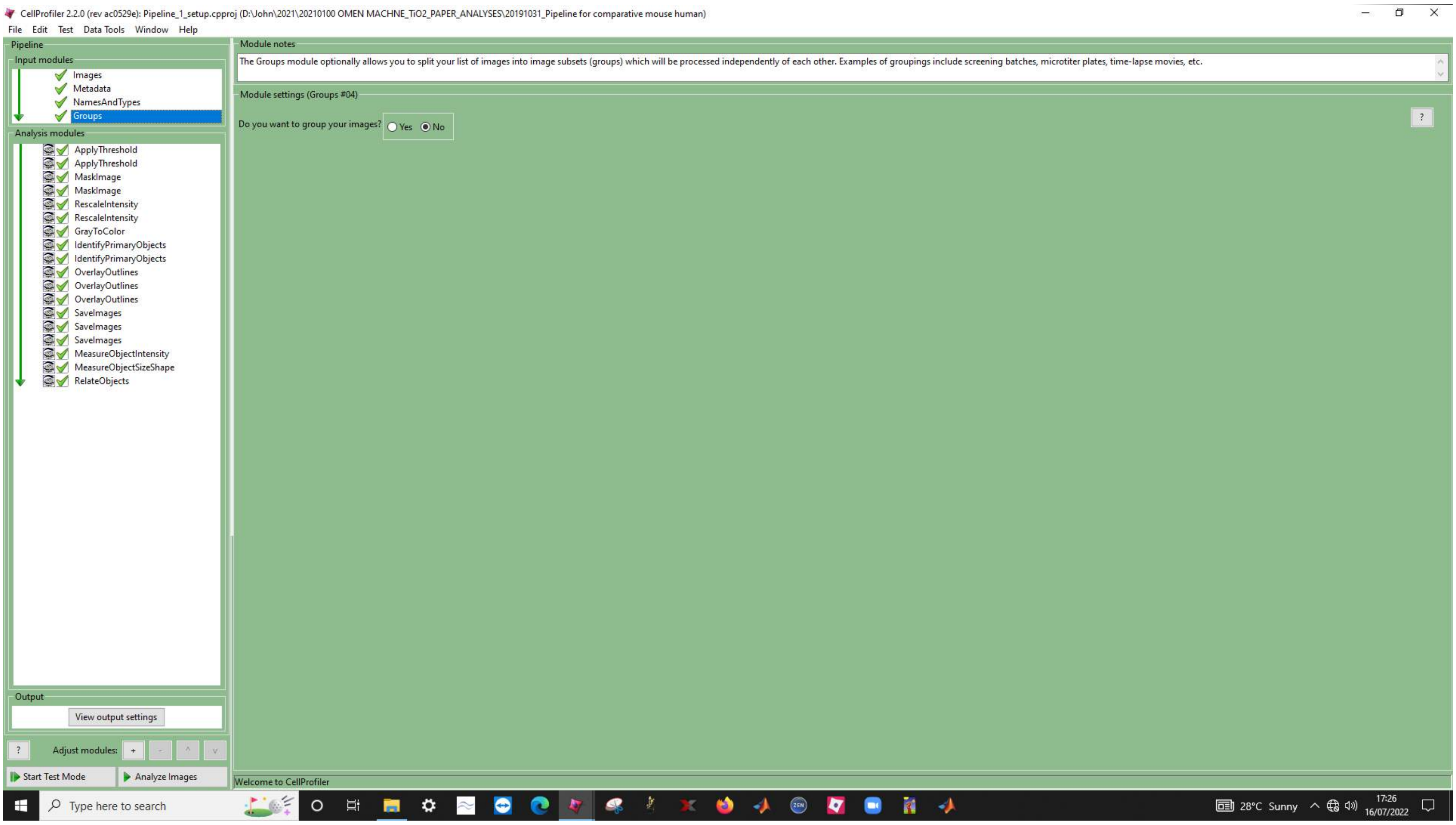
CellProfiler 2.2.0 (rev ac0529e): Pipeline_1_setup.cpproj (D:\John\2021\20210100 OMEN MACHNE_TiO2_PAPER_ANALYSES\20191031_Pipeline for comparative mouse human)
File Edit Test Data Tools Window Help
Pipeline
Input modules
Images
Metadata
NamesAndTypes
Groups
Analysis modules
ApplyThreshold
ApplyThreshold
MaskImage
MaskImage
RescaleIntensity
RescaleIntensity
GrayToColor
IdentifyPrimaryObjects
IdentifyPrimaryObjects
OverlayOutlines
OverlayOutlines
OverlayOutlines
SaveImages
SaveImages
SaveImages
MeasureObjectIntensity
MeasureObjectSizeShape
RelateObjects
Output
View output settings
Adjust modules:
Start Test Mode
Analyze Images
Module notes
The Groups module optionally allows you to split your list of images into image subsets (groups) which will be processed independently of each other. Examples of groupings include screening batches, microtiter plates, time-lapse movies, etc.
Module settings (Groups #04)
Do you want to group your images?
Yes
No
Welcome to CellProfiler
Type here to search
28°C Sunny
17:26
16/07/2022

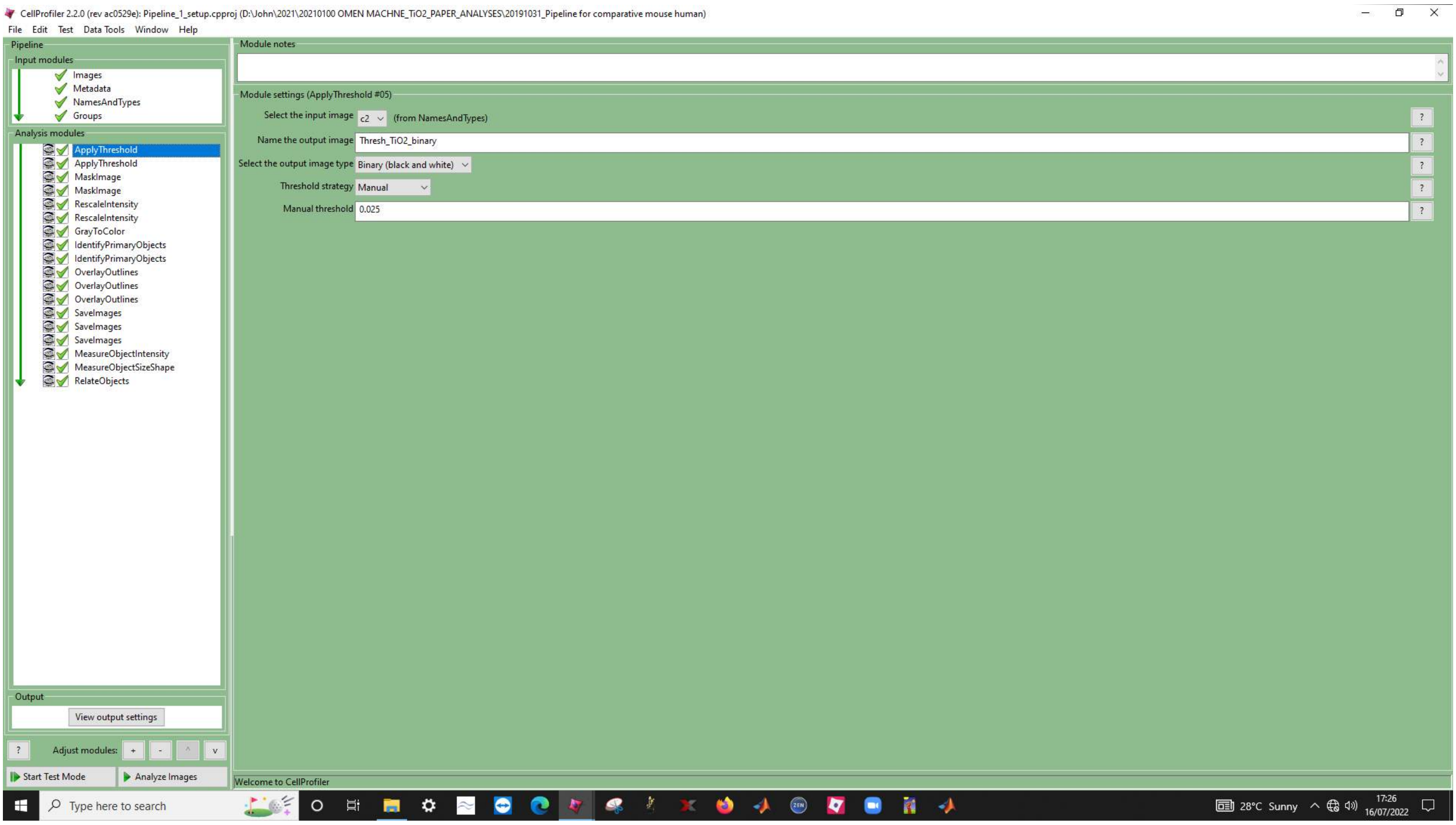
CellProfiler 2.2.0 (rev ac0529e): Pipeline_1_setup.cpproj (D:\John\2021\20210100 OMEN MACHNE_TiO2_PAPER_ANALYSES\20191031_Pipeline for comparative mouse human)
File Edit Test Data Tools Window Help
Pipeline
Input modules
Images
Metadata
NamesAndTypes
Groups
Analysis modules
ApplyThreshold
ApplyThreshold
MaskImage
MaskImage
RescaleIntensity
RescaleIntensity
GrayToColor
IdentifyPrimaryObjects
IdentifyPrimaryObjects
OverlayOutlines
OverlayOutlines
OverlayOutlines
SaveImages
SaveImages
SaveImages
MeasureObjectIntensity
MeasureObjectSizeShape
RelateObjects
Output
View output settings
Adjust modules:
Start Test Mode
Analyze Images
Module notes
Module settings (ApplyThreshold #05)
Select the input image
c2
(from NamesAndTypes)
Name the output image
Thresh_TiO2_binary
Select the output image type
Binary (black and white)
Threshold strategy
Manual
Manual threshold
0.025
Welcome to CellProfiler
Type here to search
28°C Sunny
17:26
16/07/2022

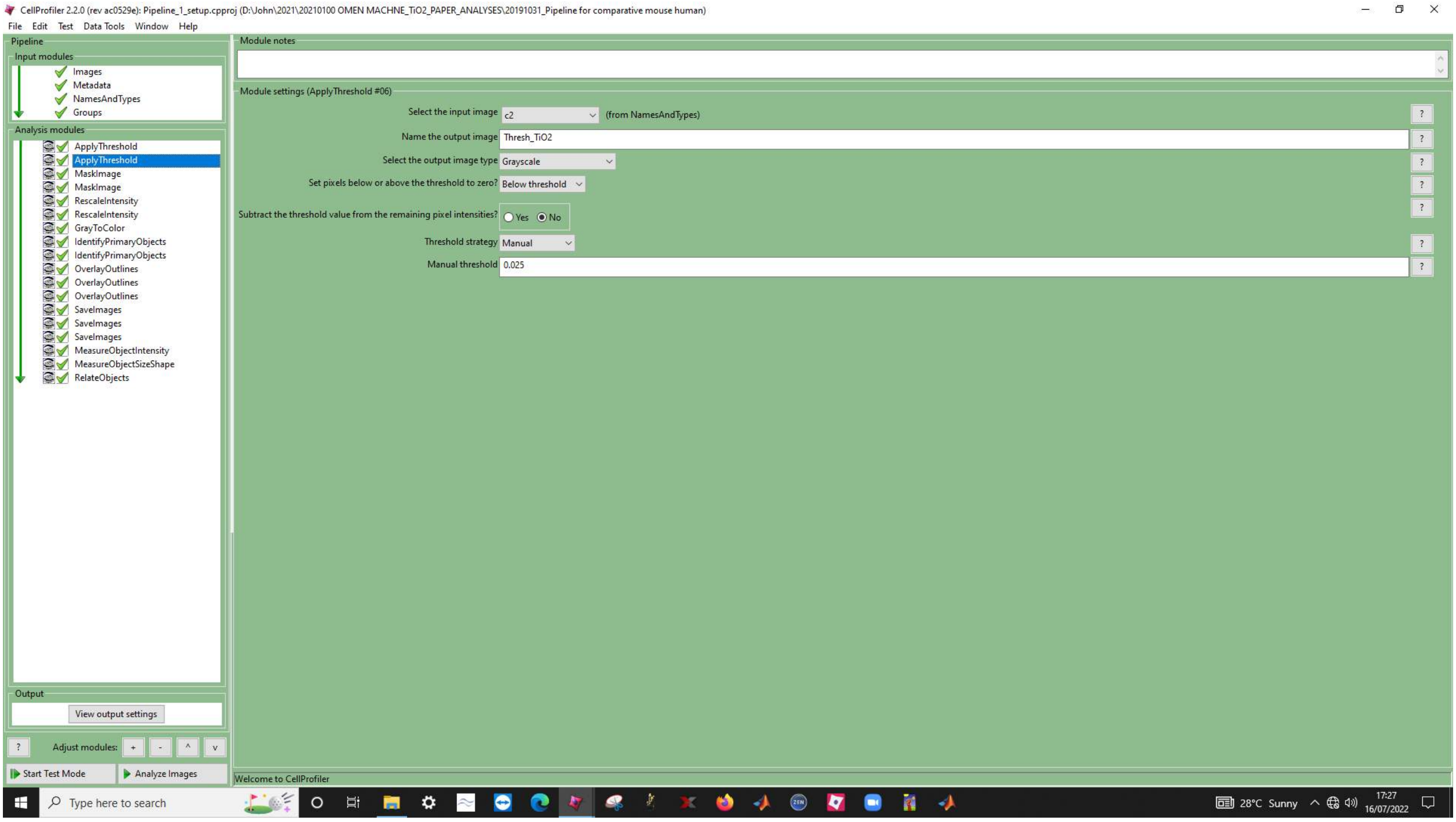
CellProfiler 2.2.0 (rev ac0529e): Pipeline_1_setup.cpproj (D:\John\2021\20210100 OMEN MACHNE_TiO2_PAPER_ANALYSES\20191031_Pipeline for comparative mouse human)
File Edit Test Data Tools Window Help
Pipeline
Input modules
Images
Metadata
NamesAndTypes
Groups
Analysis modules
ApplyThreshold
ApplyThreshold
MaskImage
MaskImage
RescaleIntensity
RescaleIntensity
GrayToColor
IdentifyPrimaryObjects
IdentifyPrimaryObjects
OverlayOutlines
OverlayOutlines
OverlayOutlines
SaveImages
SaveImages
SaveImages
MeasureObjectIntensity
MeasureObjectSizeShape
RelateObjects
Output
View output settings
Adjust modules:
Start Test Mode
Analyze Images
Module notes
Module settings (ApplyThreshold #06)
Select the input image
c2
(from NamesAndTypes)
Name the output image
Thresh_TiO2
Select the output image type
Grayscale
Set pixels below or above the threshold to zero?
Below threshold
Subtract the threshold value from the remaining pixel intensities?
Yes
No
Threshold strategy
Manual
Manual threshold
0.025
Welcome to CellProfiler
Type here to search
28°C Sunny
17:27
16/07/2022

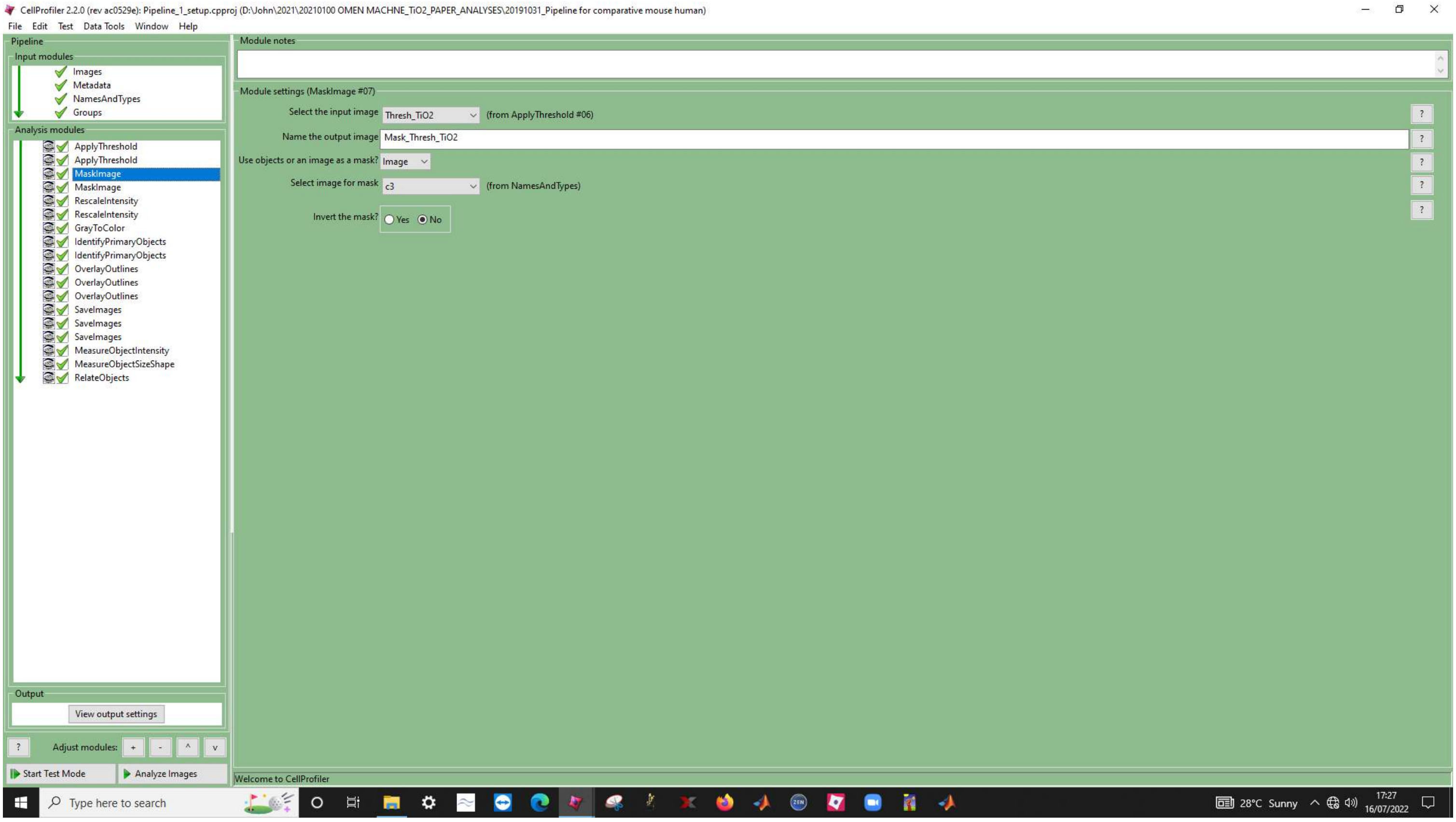
CellProfiler 2.2.0 (rev ac0529e): Pipeline_1_setup.cpproj (D:\John\2021\20210100 OMEN MACHNE_TiO2_PAPER_ANALYSES\20191031_Pipeline for comparative mouse human)
File Edit Test Data Tools Window Help
Pipeline
Input modules
Images
Metadata
NamesAndTypes
Groups
Analysis modules
ApplyThreshold
ApplyThreshold
MaskImage
MaskImage
RescaleIntensity
RescaleIntensity
GrayToColor
IdentifyPrimaryObjects
IdentifyPrimaryObjects
OverlayOutlines
OverlayOutlines
OverlayOutlines
SaveImages
SaveImages
SaveImages
MeasureObjectIntensity
MeasureObjectSizeShape
RelateObjects
Output
View output settings
Adjust modules:
Start Test Mode
Analyze Images
Module notes
Module settings (MaskImage #07)
Select the input image
Thresh_TiO2
(from ApplyThreshold #06)
Name the output image
Mask_Thresh_TiO2
Use objects or an image as a mask?
Image
Select image for mask
c3
(from NamesAndTypes)
Invert the mask?
Yes
No
Welcome to CellProfiler
Type here to search
28°C Sunny
17:27
16/07/2022

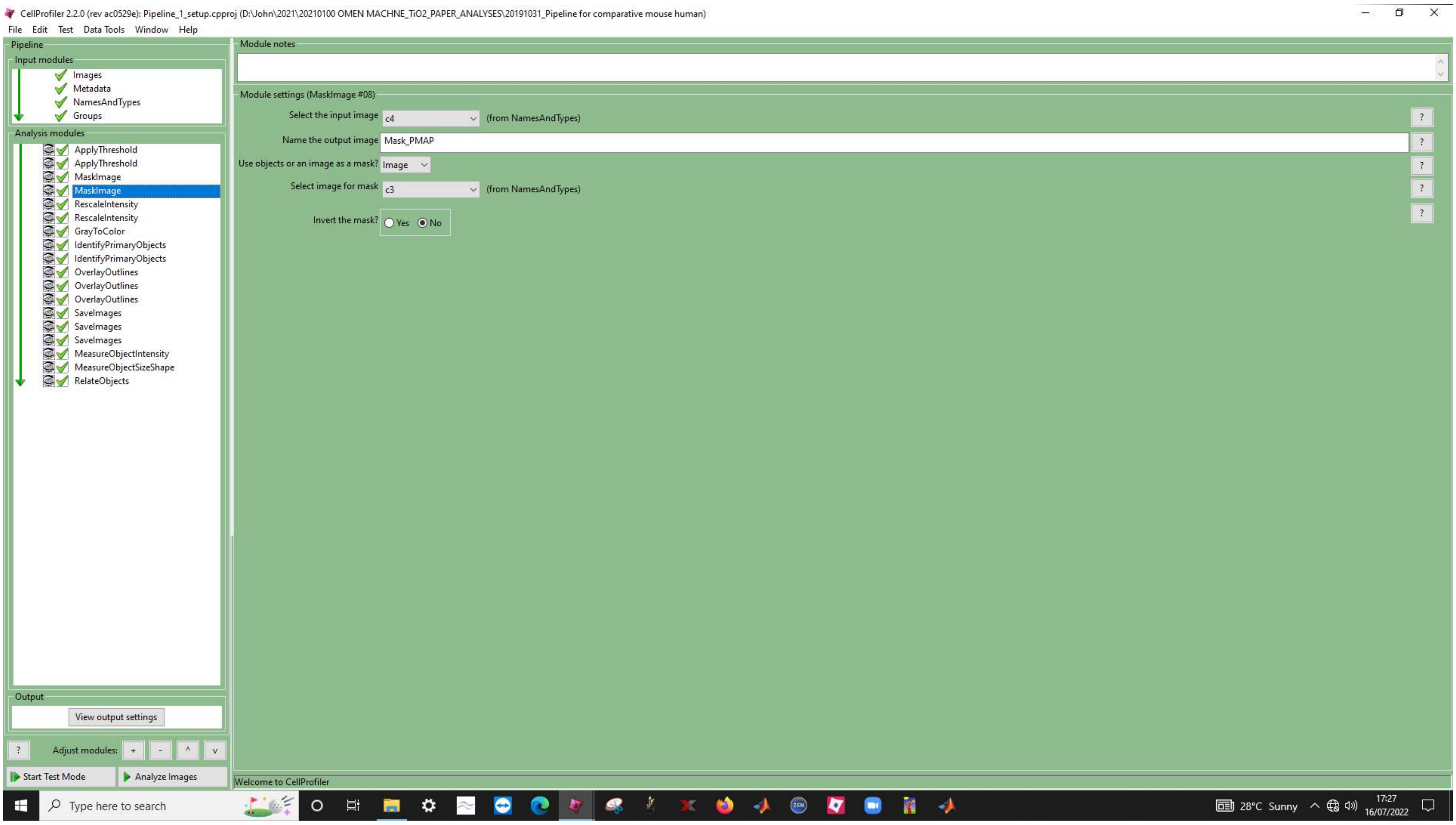
CellProfiler 2.2.0 (rev ac0529e): Pipeline_1_setup.cpproj (D:\John\2021\20210100 OMEN MACHNE_TiO2_PAPER_ANALYSES\20191031_Pipeline for comparative mouse human)
File Edit Test Data Tools Window Help
Pipeline
Input modules
Images
Metadata
NamesAndTypes
Groups
Analysis modules
ApplyThreshold
ApplyThreshold
MaskImage
MaskImage
RescaleIntensity
RescaleIntensity
GrayToColor
IdentifyPrimaryObjects
IdentifyPrimaryObjects
OverlayOutlines
OverlayOutlines
OverlayOutlines
SaveImages
SaveImages
SaveImages
MeasureObjectIntensity
MeasureObjectSizeShape
RelateObjects
Output
View output settings
Adjust modules:
Start Test Mode
Analyze Images
Module notes
Module settings (MaskImage #08)
Select the input image
c4
(from NamesAndTypes)
Name the output image
Mask_PMAP
Use objects or an image as a mask?
Image
Select image for mask
c3
(from NamesAndTypes)
Invert the mask?
Yes
No
Welcome to CellProfiler
Type here to search
28°C Sunny
17:27
16/07/2022

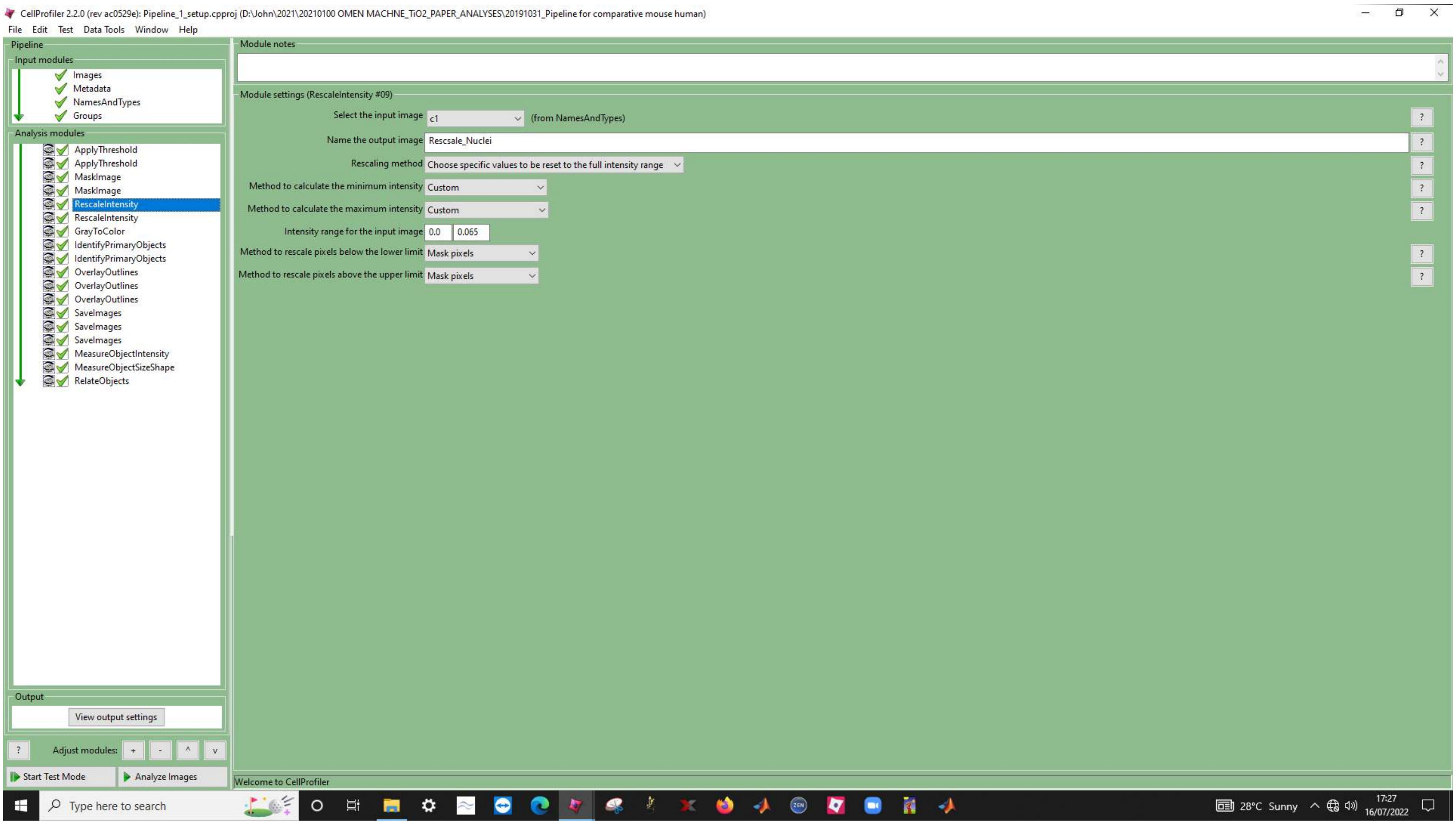
CellProfiler 2.2.0 (rev ac0529e): Pipeline_1_setup.cpproj (D:\John\2021\20210100 OMEN MACHNE_TiO2_PAPER_ANALYSES\20191031_Pipeline for comparative mouse human)
File Edit Test Data Tools Window Help
Pipeline
Input modules
Images
Metadata
NamesAndTypes
Groups
Analysis modules
ApplyThreshold
ApplyThreshold
MaskImage
MaskImage
RescaleIntensity
RescaleIntensity
GrayToColor
IdentifyPrimaryObjects
IdentifyPrimaryObjects
OverlayOutlines
OverlayOutlines
OverlayOutlines
SaveImages
SaveImages
SaveImages
MeasureObjectIntensity
MeasureObjectSizeShape
RelateObjects
Output
View output settings
Adjust modules:
Start Test Mode
Analyze Images
Module notes
Module settings (RescaleIntensity #09)
Select the input image c1 (from NamesAndTypes)
Name the output image Rescsale_Nuclei
Rescaling method Choose specific values to be reset to the full intensity range
Method to calculate the minimum intensity Custom
Method to calculate the maximum intensity Custom
Intensity range for the input image 0.0 0.065
Method to rescale pixels below the lower limit Mask pixels
Method to rescale pixels above the upper limit Mask pixels
Welcome to CellProfiler
Type here to search
28°C Sunny
17:27
16/07/2022

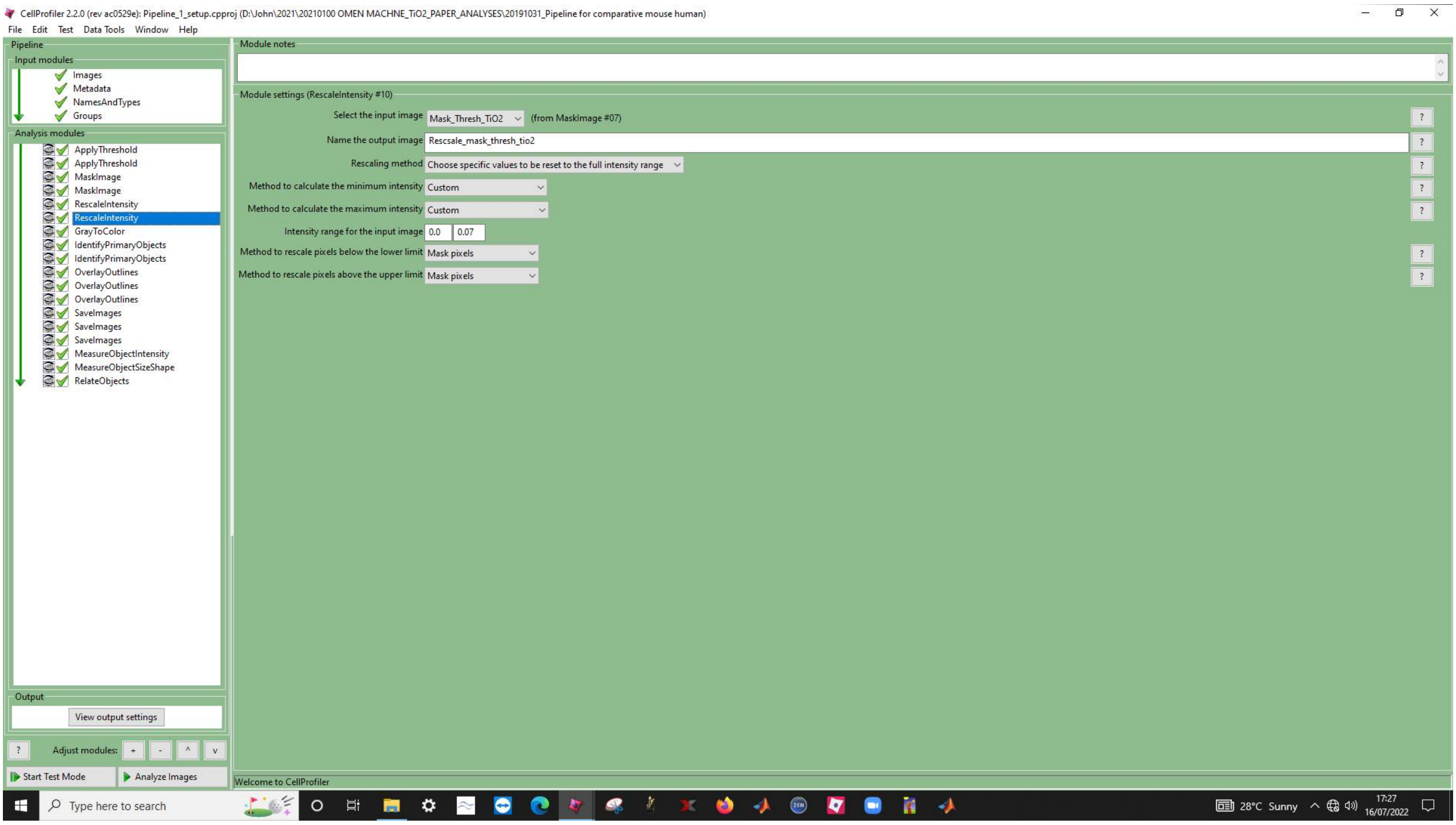
CellProfiler 2.2.0 (rev ac0529e): Pipeline_1_setup.cpproj (D:\John\2021\20210100 OMEN MACHNE_TiO2_PAPER_ANALYSES\20191031_Pipeline for comparative mouse human)
File Edit Test Data Tools Window Help
Pipeline
Input modules
Images
Metadata
NamesAndTypes
Groups
Analysis modules
ApplyThreshold
ApplyThreshold
MaskImage
MaskImage
RescaleIntensity
RescaleIntensity
GrayToColor
IdentifyPrimaryObjects
IdentifyPrimaryObjects
OverlayOutlines
OverlayOutlines
OverlayOutlines
SaveImages
SaveImages
SaveImages
MeasureObjectIntensity
MeasureObjectSizeShape
RelateObjects
Output
View output settings
Adjust modules:
Start Test Mode
Analyze Images
Module notes
Module settings (RescaleIntensity #10)
Select the input image Mask_Thresh_TiO2 (from MaskImage #07)
Name the output image Rescsale_mask_thresh_tio2
Rescaling method Choose specific values to be reset to the full intensity range
Method to calculate the minimum intensity Custom
Method to calculate the maximum intensity Custom
Intensity range for the input image 0.0 0.07
Method to rescale pixels below the lower limit Mask pixels
Method to rescale pixels above the upper limit Mask pixels
Welcome to CellProfiler
Type here to search
28°C Sunny
17:27
16/07/2022

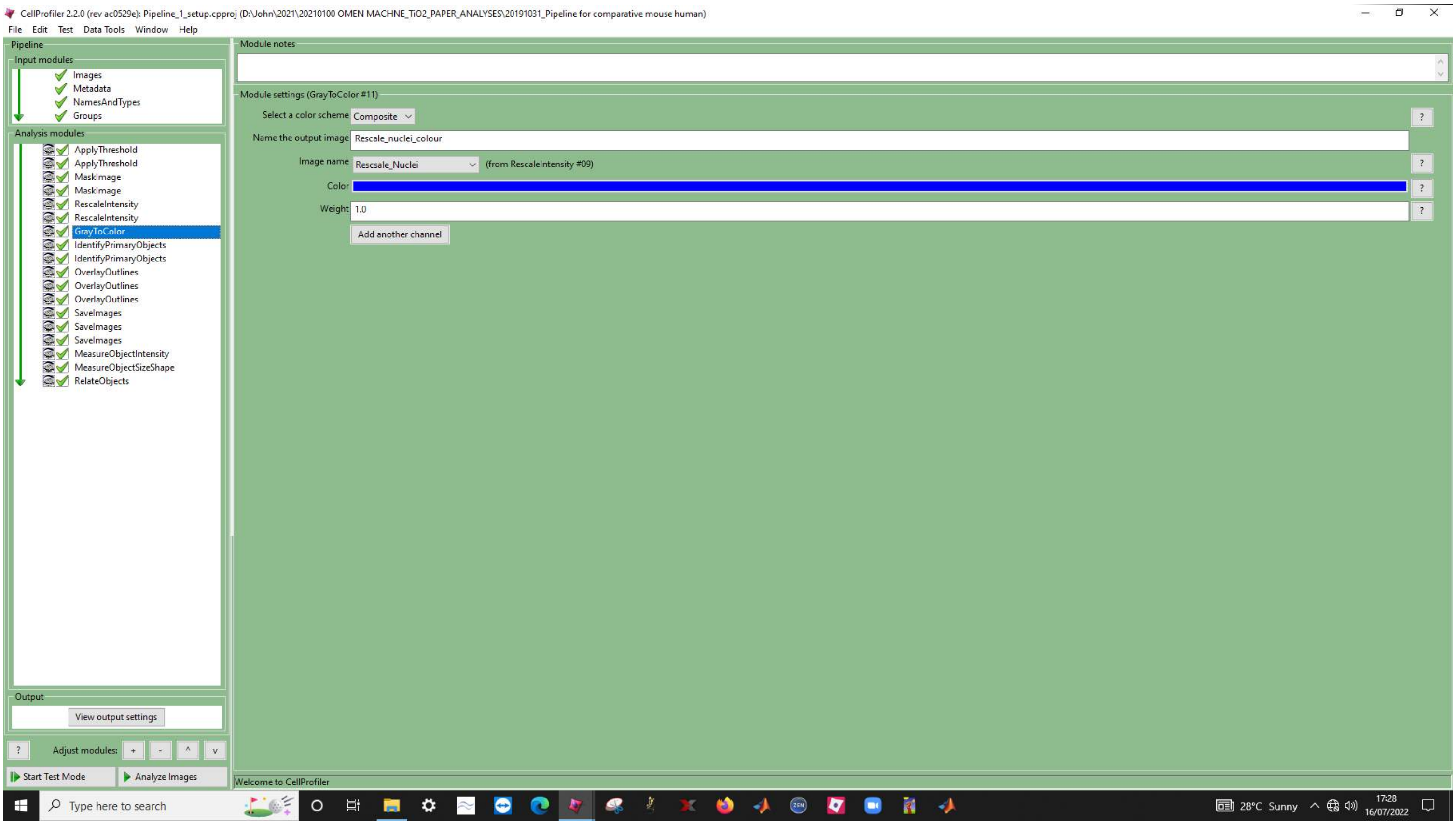

CellProfiler 2.2.0 (rev ac0529e): Pipeline_1_setup.cpproj (D:\John\2021\20210100 OMEN MACHNE_TiO2_PAPER_ANALYSES\20191031_Pipeline for comparative mouse human)
File Edit Test Data Tools Window Help
Pipeline
Input modules
Images
Metadata
NamesAndTypes
Groups
Analysis modules
ApplyThreshold
ApplyThreshold
MaskImage
MaskImage
RescaleIntensity
RescaleIntensity
GrayToColor
IdentifyPrimaryObjects
IdentifyPrimaryObjects
OverlayOutlines
OverlayOutlines
OverlayOutlines
SaveImages
SaveImages
SaveImages
MeasureObjectIntensity
MeasureObjectSizeShape
RelateObjects
Output
View output settings
Adjust modules:
Start Test Mode
Analyze Images
Module notes
Module settings (GrayToColor #11)
Select a color scheme
Composite
Name the output image
Rescale_nuclei_colour
Image name
Rescsale_Nuclei
(from RescaleIntensity #09)
Color
Weight
1.0
Add another channel
Welcome to CellProfiler
Type here to search
28°C Sunny
17:28
16/07/2022

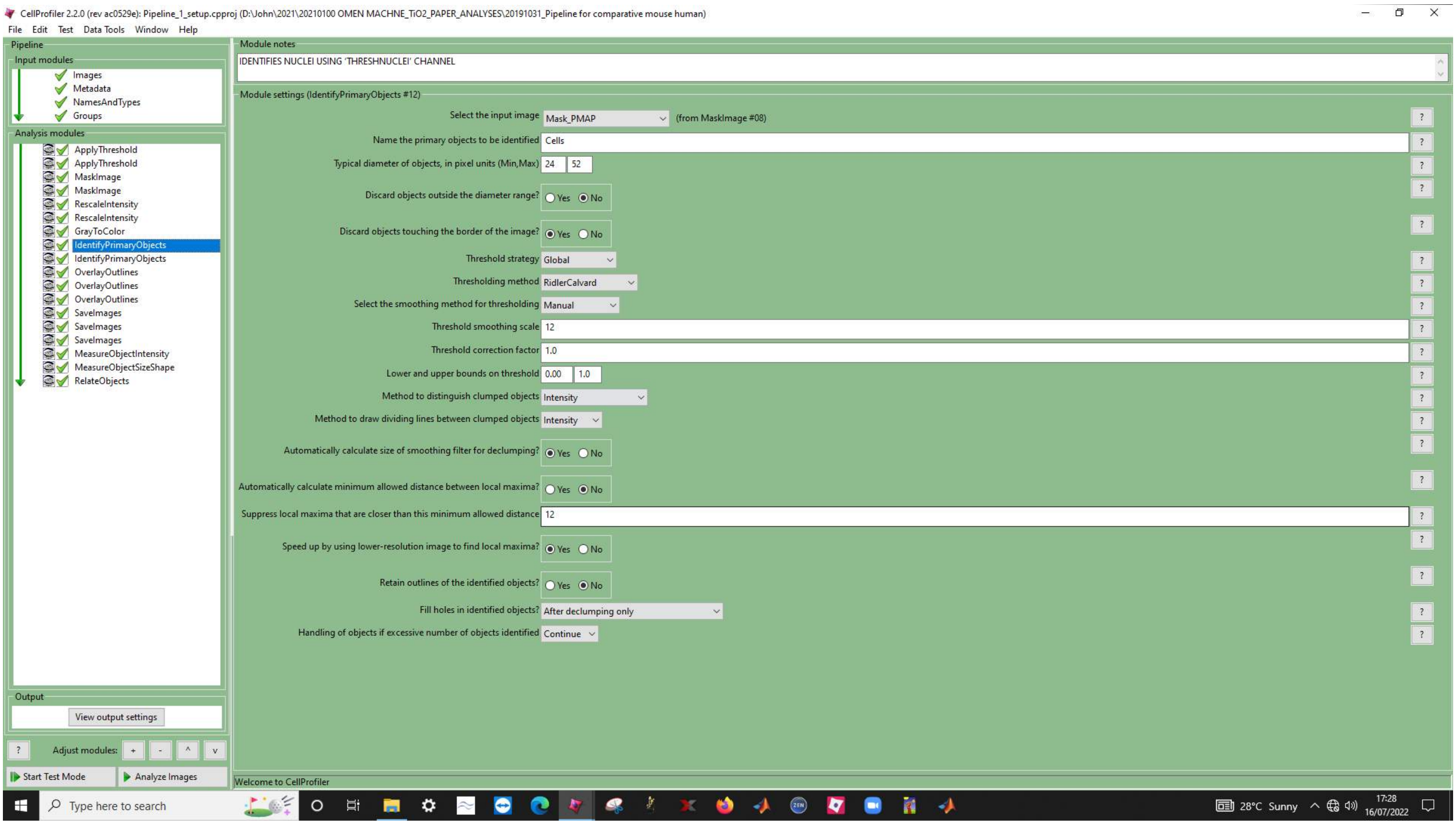
CellProfiler 2.2.0 (rev ac0529e): Pipeline_1_setup.cpproj (D:\John\2021\20210100 OMEN MACHNE_TiO2_PAPER_ANALYSES\20191031_Pipeline for comparative mouse human)
File Edit Test Data Tools Window Help
Pipeline
Input modules
Images
Metadata
NamesAndTypes
Groups
Analysis modules
ApplyThreshold
ApplyThreshold
MaskImage
MaskImage
RescaleIntensity
RescaleIntensity
GrayToColor
IdentifyPrimaryObjects
IdentifyPrimaryObjects
OverlayOutlines
OverlayOutlines
OverlayOutlines
SaveImages
SaveImages
SaveImages
MeasureObjectIntensity
MeasureObjectSizeShape
RelateObjects
Output
View output settings
Adjust modules:
Start Test Mode
Analyze Images
Module notes
IDENTIFIES NUCLEI USING 'THRESHNUCLEI' CHANNEL
Module settings (IdentifyPrimaryObjects #12)
Select the input image Mask_PMAP (from MaskImage #08)
Name the primary objects to be identified Cells
Typical diameter of objects, in pixel units (Min,Max) 24 52
Discard objects outside the diameter range? Yes No
Discard objects touching the border of the image? Yes No
Threshold strategy Global
Thresholding method RidlerCalvard
Select the smoothing method for thresholding Manual
Threshold smoothing scale 12
Threshold correction factor 1.0
Lower and upper bounds on threshold 0.00 1.0
Method to distinguish clumped objects Intensity
Method to draw dividing lines between clumped objects Intensity
Automatically calculate size of smoothing filter for declumping? Yes No
Automatically calculate minimum allowed distance between local maxima? Yes No
Suppress local maxima that are closer than this minimum allowed distance 12
Speed up by using lower-resolution image to find local maxima? Yes No
Retain outlines of the identified objects? Yes No
Fill holes in identified objects? After declumping only
Handling of objects if excessive number of objects identified Continue
Welcome to CellProfiler
Type here to search
28°C Sunny
17:28
16/07/2022

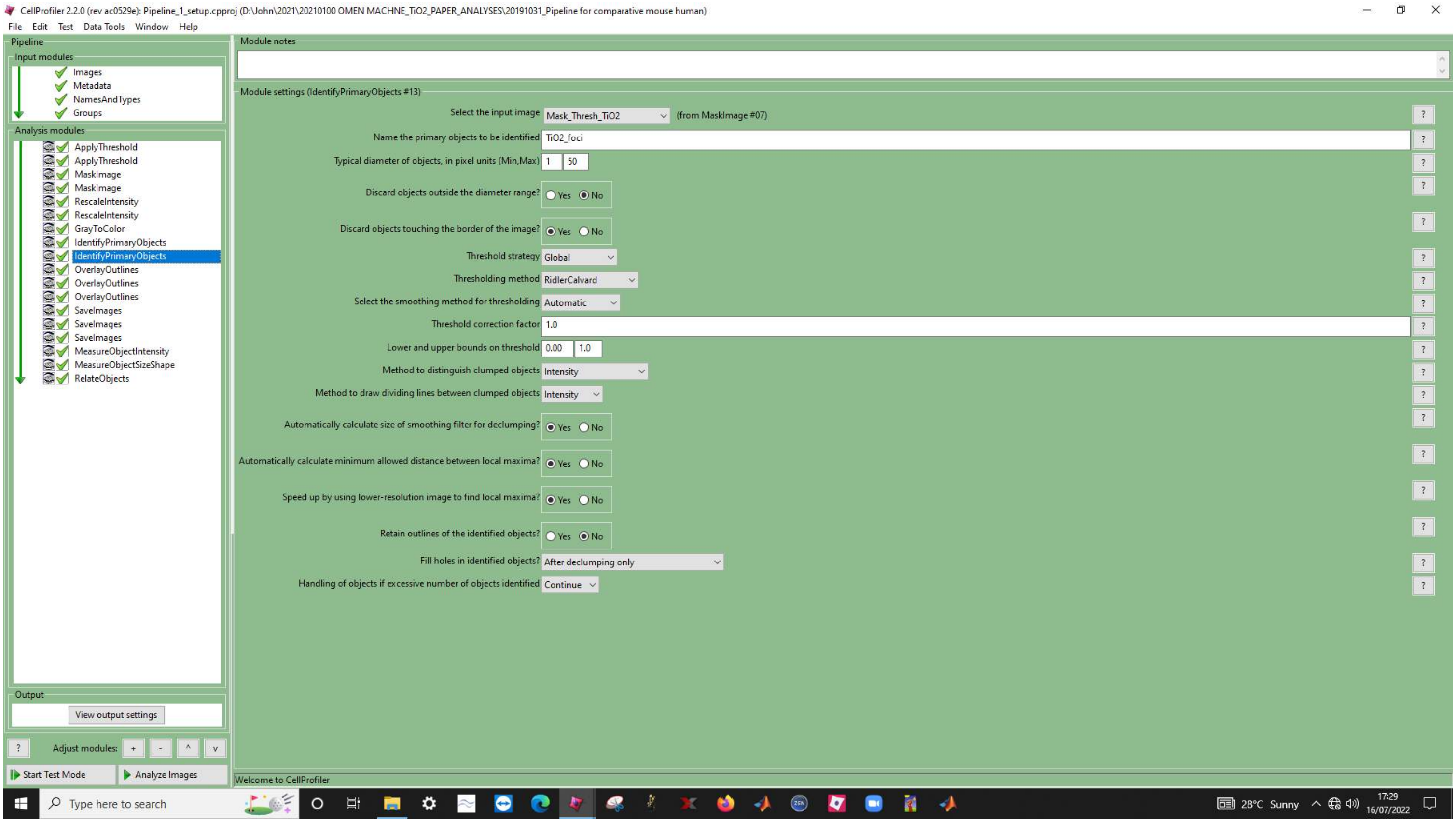
CellProfiler 2.2.0 (rev ac0529e): Pipeline_1_setup.cpproj (D:\John\2021\20210100 OMEN MACHINE_TiO2_PAPER_ANALYSES\20191031_Pipeline for comparative mouse human)
File Edit Test Data Tools Window Help
Pipeline
Input modules
Images
Metadata
NamesAndTypes
Groups
Analysis modules
ApplyThreshold
ApplyThreshold
MaskImage
MaskImage
RescaleIntensity
RescaleIntensity
GrayToColor
IdentifyPrimaryObjects
IdentifyPrimaryObjects
OverlayOutlines
OverlayOutlines
OverlayOutlines
SaveImages
SaveImages
SaveImages
MeasureObjectIntensity
MeasureObjectSizeShape
RelateObjects
Output
View output settings
Adjust modules:
Start Test Mode
Analyze Images
Module notes
Module settings (IdentifyPrimaryObjects #13)
Select the input image Mask_Thresh_TiO2 (from MaskImage #07)
Name the primary objects to be identified TiO2_foci
Typical diameter of objects, in pixel units (Min,Max) 1 50
Discard objects outside the diameter range? Yes No
Discard objects touching the border of the image? Yes No
Threshold strategy Global
Thresholding method RidlerCalvard
Select the smoothing method for thresholding Automatic
Threshold correction factor 1.0
Lower and upper bounds on threshold 0.00 1.0
Method to distinguish clumped objects Intensity
Method to draw dividing lines between clumped objects Intensity
Automatically calculate size of smoothing filter for declumping? Yes No
Automatically calculate minimum allowed distance between local maxima? Yes No
Speed up by using lower-resolution image to find local maxima? Yes No
Retain outlines of the identified objects? Yes No
Fill holes in identified objects? After declumping only
Handling of objects if excessive number of objects identified Continue
Welcome to CellProfiler
Type here to search
28°C Sunny
17:29
16/07/2022

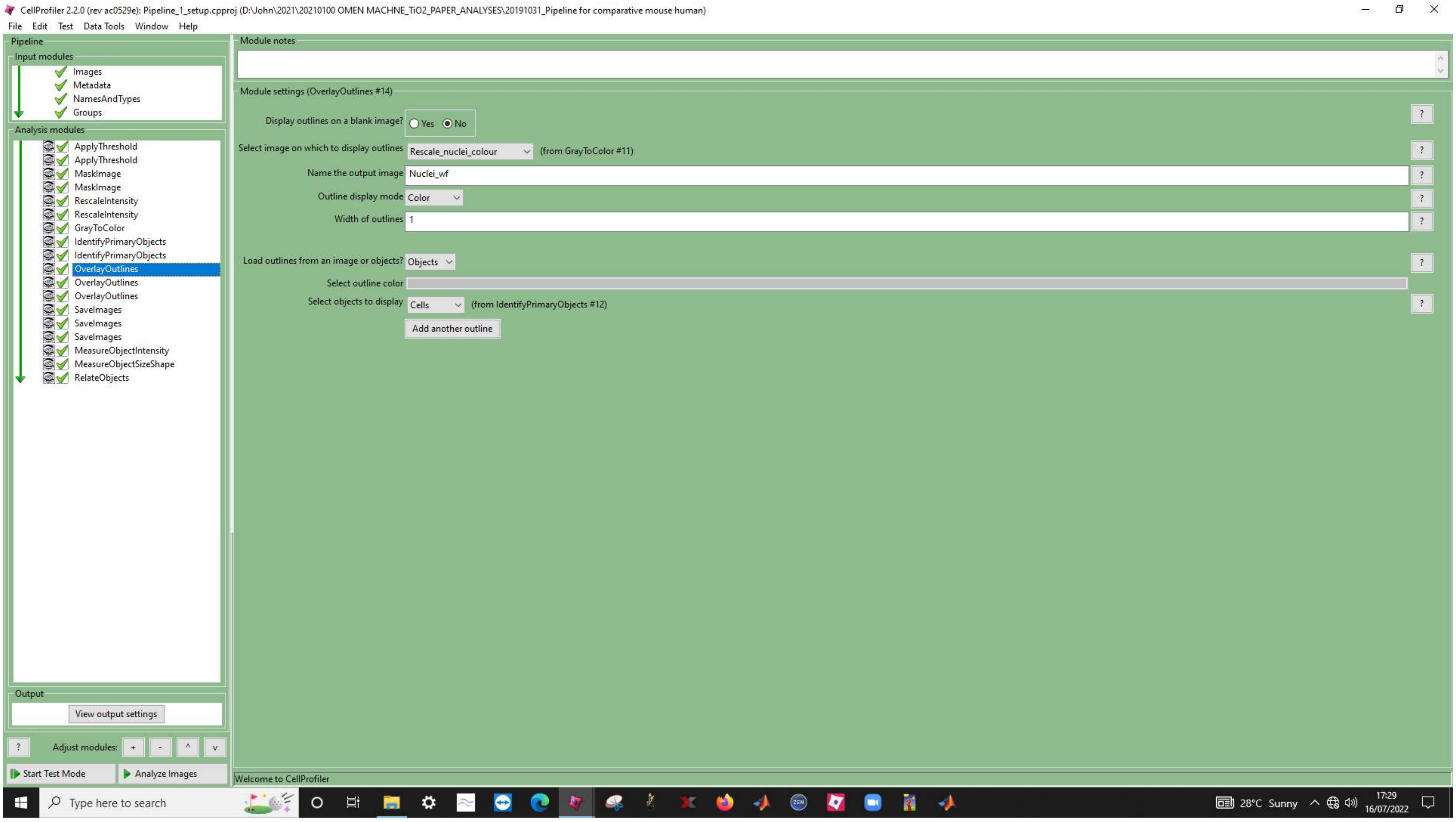
CellProfiler 2.2.0 (rev ac0529e): Pipeline_1_setup.cpproj (D:\John\2021\20210100 OMEN MACHNE_TiO2_PAPER_ANALYSES\20191031_Pipeline for comparative mouse human)
File Edit Test Data Tools Window Help
Pipeline
Input modules
Images
Metadata
NamesAndTypes
Groups
Analysis modules
ApplyThreshold
ApplyThreshold
MaskImage
MaskImage
RescaleIntensity
RescaleIntensity
GrayToColor
IdentifyPrimaryObjects
IdentifyPrimaryObjects
OverlayOutlines
OverlayOutlines
OverlayOutlines
SaveImages
SaveImages
SaveImages
MeasureObjectIntensity
MeasureObjectSizeShape
RelateObjects
Output
View output settings
Adjust modules:
Start Test Mode
Analyze Images
Module notes
Module settings (OverlayOutlines #14)
Display outlines on a blank image? Yes No
Select image on which to display outlines Rescale_nuclei_colour (from GrayToColor #11)
Name the output image Nuclei_wf
Outline display mode Color
Width of outlines 1
Load outlines from an image or objects? Objects
Select outline color
Select objects to display Cells (from IdentifyPrimaryObjects #12)
Add another outline
Welcome to CellProfiler
Type here to search
28°C Sunny
17:29
16/07/2022

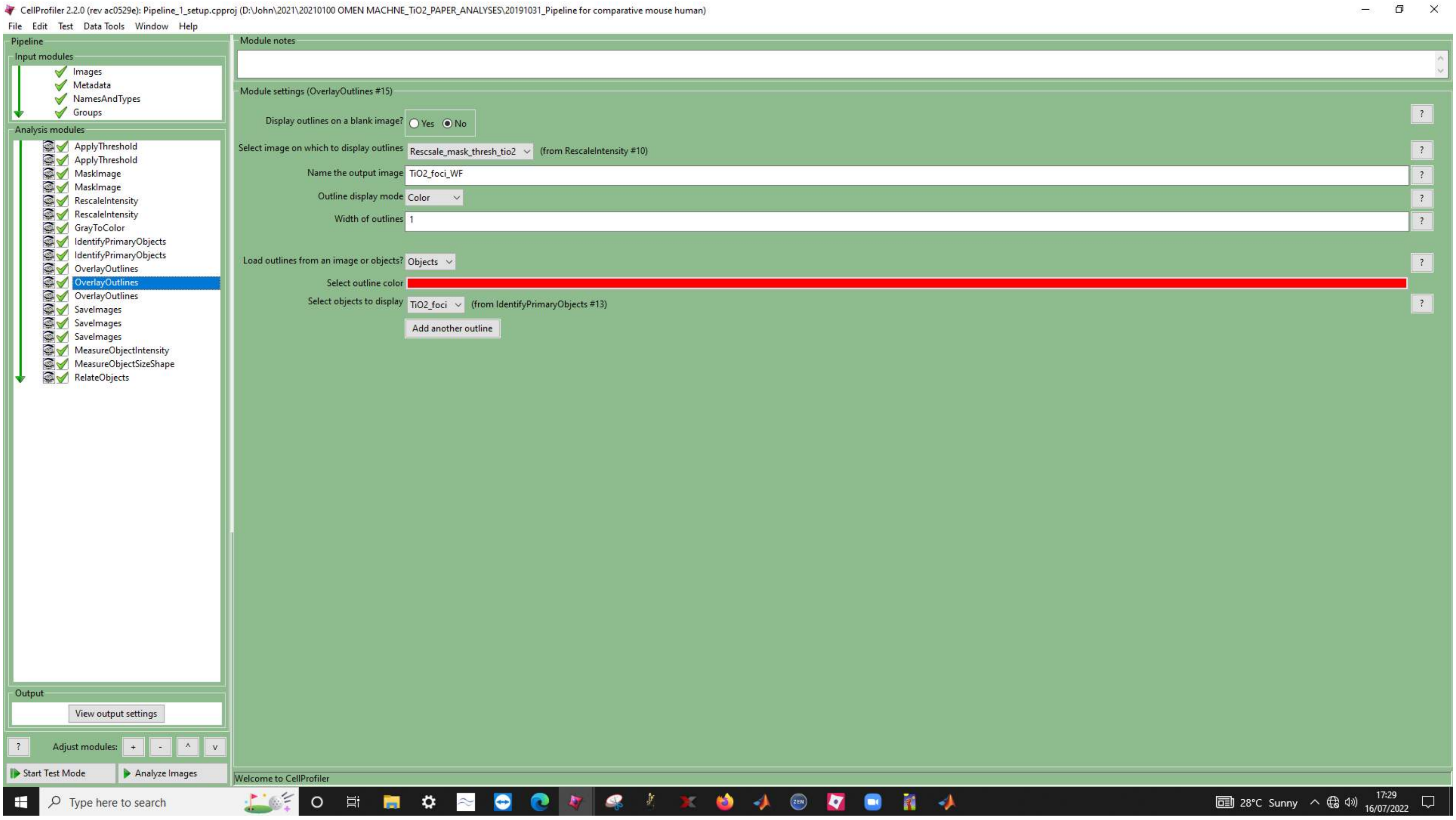
CellProfiler 2.2.0 (rev ac0529e): Pipeline_1_setup.cpproj (D:\John\2021\20210100 OMEN MACHNE_TiO2_PAPER_ANALYSES\20191031_Pipeline for comparative mouse human)
File Edit Test Data Tools Window Help
Pipeline
Input modules
Images
Metadata
NamesAndTypes
Groups
Analysis modules
ApplyThreshold
ApplyThreshold
MaskImage
MaskImage
RescaleIntensity
RescaleIntensity
GrayToColor
IdentifyPrimaryObjects
IdentifyPrimaryObjects
OverlayOutlines
OverlayOutlines
OverlayOutlines
SaveImages
SaveImages
SaveImages
MeasureObjectIntensity
MeasureObjectSizeShape
RelateObjects
Output
View output settings
Adjust modules:
Start Test Mode
Analyze Images
Module notes
Module settings (OverlayOutlines #15)
Display outlines on a blank image? Yes No
Select image on which to display outlines Rescsale_mask_thresh_tio2 (from RescaleIntensity #10)
Name the output image TiO2_foci_WF
Outline display mode Color
Width of outlines 1
Load outlines from an image or objects? Objects
Select outline color
Select objects to display TiO2_foci (from IdentifyPrimaryObjects #13)
Add another outline
Welcome to CellProfiler
Type here to search
28°C Sunny
17:29
16/07/2022

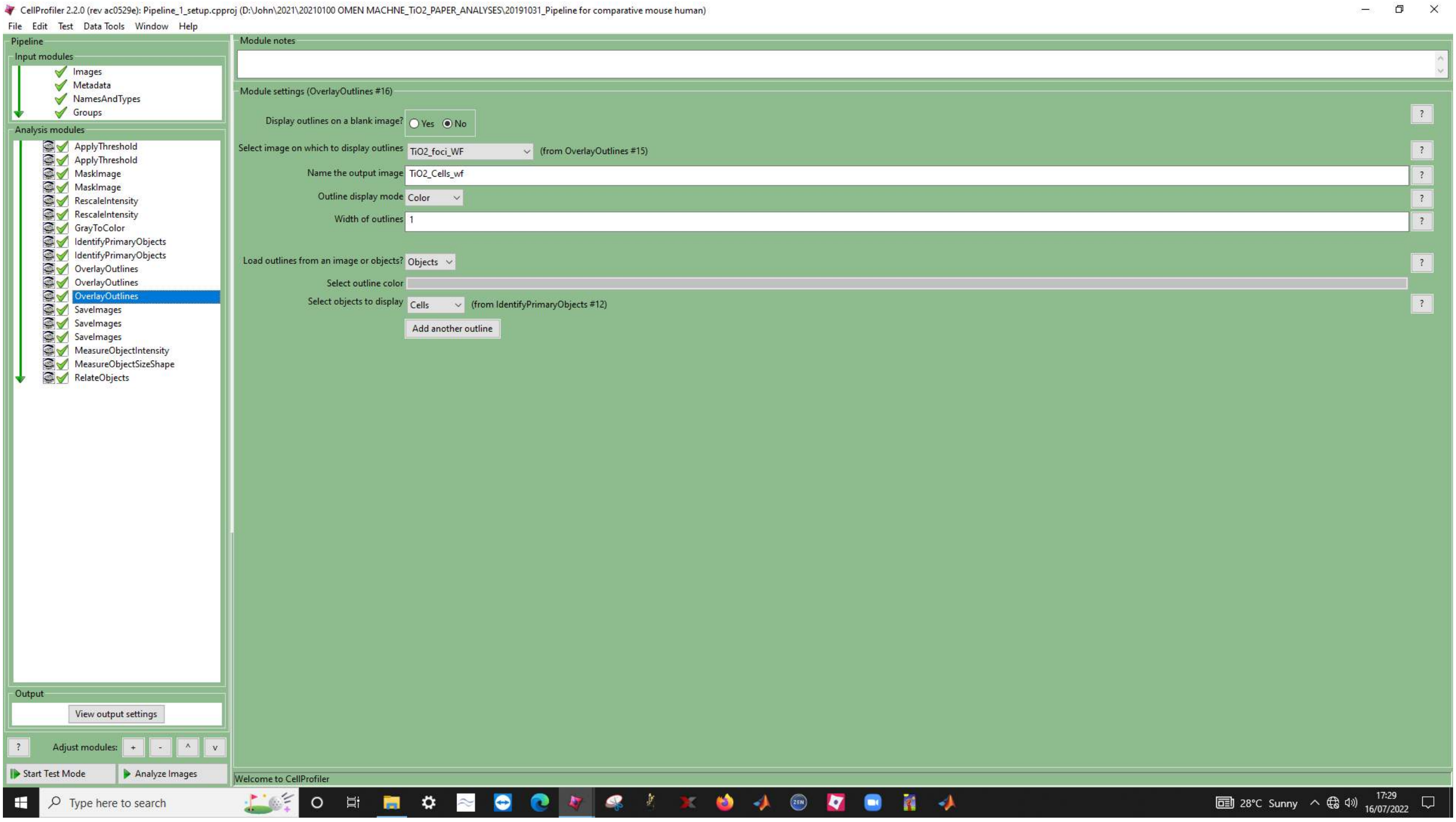
CellProfiler 2.2.0 (rev ac0529e): Pipeline_1_setup.cpproj (D:\John\2021\20210100 OMEN MACHINE_TiO2_PAPER_ANALYSES\20191031_Pipeline for comparative mouse human)
File Edit Test Data Tools Window Help
Pipeline
Input modules
Images
Metadata
NamesAndTypes
Groups
Analysis modules
ApplyThreshold
ApplyThreshold
MaskImage
MaskImage
RescaleIntensity
RescaleIntensity
GrayToColor
IdentifyPrimaryObjects
IdentifyPrimaryObjects
OverlayOutlines
OverlayOutlines
OverlayOutlines
SaveImages
SaveImages
SaveImages
MeasureObjectIntensity
MeasureObjectSizeShape
RelateObjects
Output
View output settings
Adjust modules:
Start Test Mode
Analyze Images
Module notes
Module settings (OverlayOutlines #16)
Display outlines on a blank image? Yes No
Select image on which to display outlines TiO2_foci_WF (from OverlayOutlines #15)
Name the output image TiO2_Cells_wf
Outline display mode Color
Width of outlines 1
Load outlines from an image or objects? Objects
Select outline color
Select objects to display Cells (from IdentifyPrimaryObjects #12)
Add another outline
Welcome to CellProfiler
Type here to search
28°C Sunny
17:29
16/07/2022

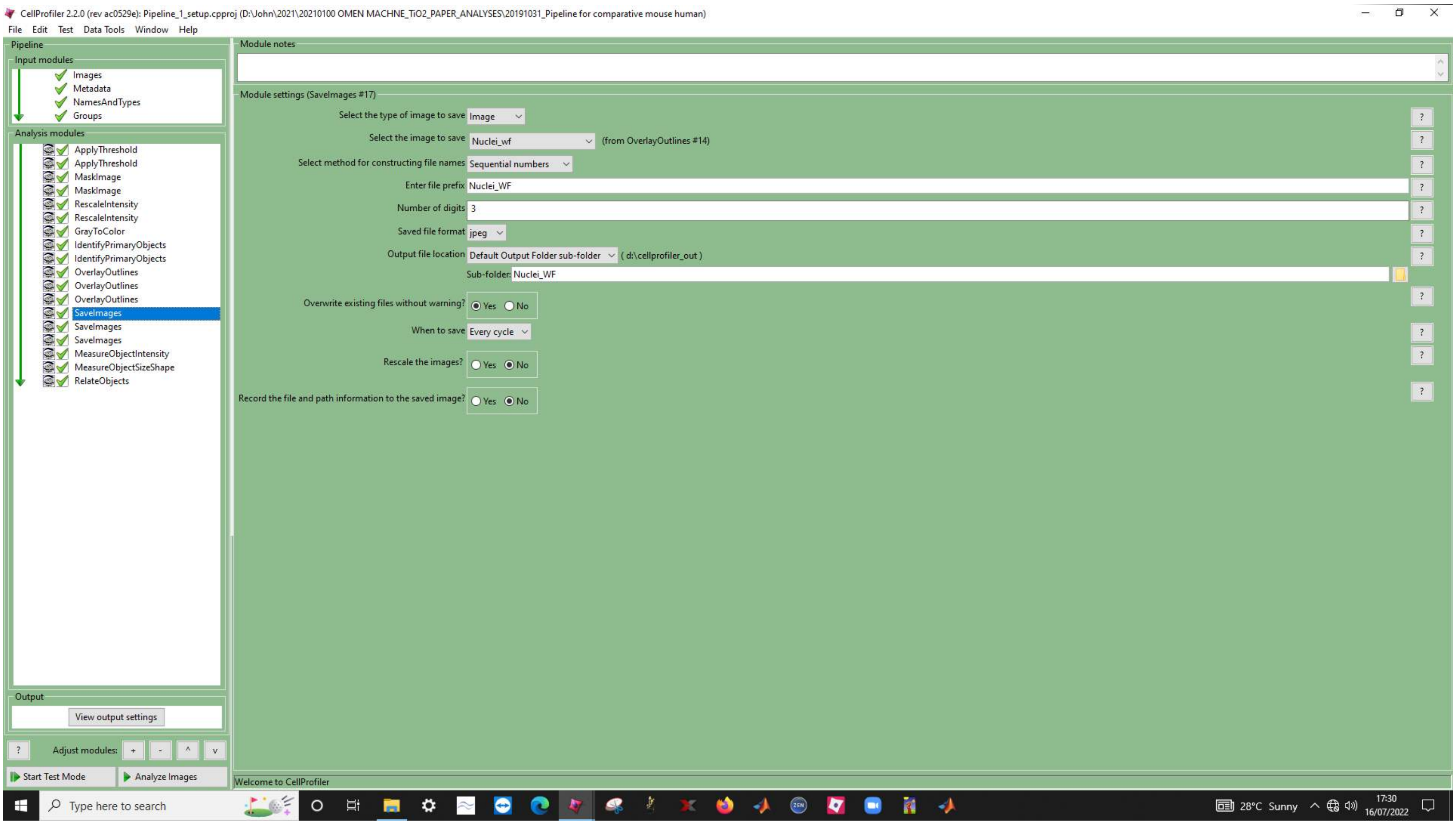
CellProfiler 2.2.0 (rev ac0529e): Pipeline_1_setup.cpproj (D:\John\2021\20210100 OMEN MACHNE_TiO2_PAPER_ANALYSES\20191031_Pipeline for comparative mouse human)
File Edit Test Data Tools Window Help
Pipeline
Input modules
Images
Metadata
NamesAndTypes
Groups
Analysis modules
ApplyThreshold
ApplyThreshold
MaskImage
MaskImage
RescaleIntensity
RescaleIntensity
GrayToColor
IdentifyPrimaryObjects
IdentifyPrimaryObjects
OverlayOutlines
OverlayOutlines
OverlayOutlines
SaveImages
SaveImages
SaveImages
MeasureObjectIntensity
MeasureObjectSizeShape
RelateObjects
Output
View output settings
Adjust modules:
Start Test Mode
Analyze Images
Module notes
Module settings (SaveImages #17)
Select the type of image to save Image
Select the image to save Nuclei_wf (from OverlayOutlines #14)
Select method for constructing file names Sequential numbers
Enter file prefix Nuclei_WF
Number of digits 3
Saved file format jpeg
Output file location Default Output Folder sub-folder ( d:\cellprofiler_out )
Sub-folder: Nuclei_WF
Overwrite existing files without warning? Yes No
When to save Every cycle
Rescale the images? Yes No
Record the file and path information to the saved image? Yes No
Welcome to CellProfiler
Type here to search
28°C Sunny
17:30
16/07/2022

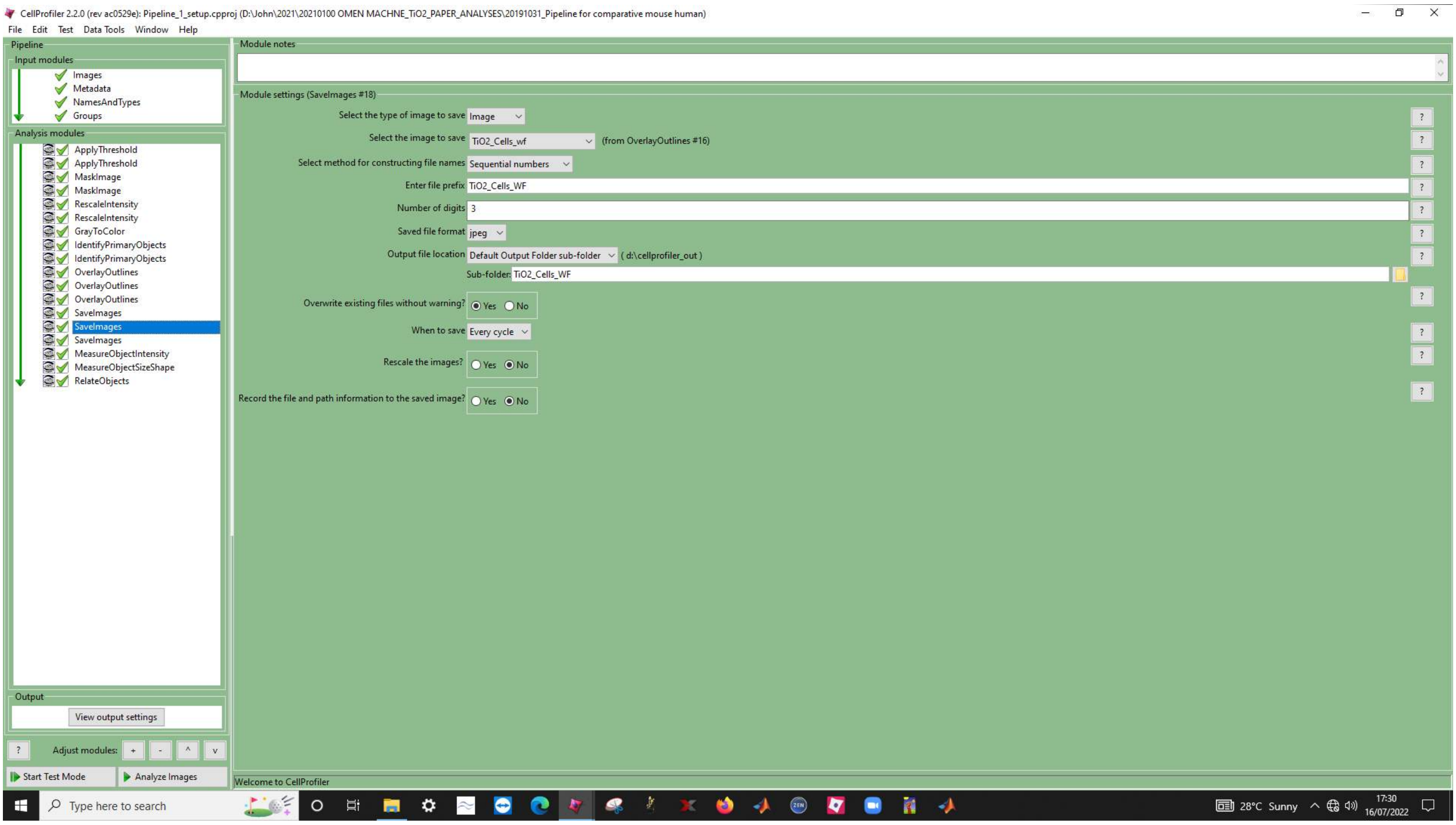

CellProfiler 2.2.0 (rev ac0529e): Pipeline_1_setup.cpproj (D:\John\2021\20210100 OMEN MACHNE_TiO2_PAPER_ANALYSES\20191031_Pipeline for comparative mouse human)
File Edit Test Data Tools Window Help
Pipeline
Input modules
Images
Metadata
NamesAndTypes
Groups
Analysis modules
ApplyThreshold
ApplyThreshold
MaskImage
MaskImage
RescaleIntensity
RescaleIntensity
GrayToColor
IdentifyPrimaryObjects
IdentifyPrimaryObjects
OverlayOutlines
OverlayOutlines
OverlayOutlines
SaveImages
SaveImages
SaveImages
MeasureObjectIntensity
MeasureObjectSizeShape
RelateObjects
Output
View output settings
Adjust modules:
Start Test Mode
Analyze Images
Module notes
Module settings (SaveImages #18)
Select the type of image to save Image
Select the image to save TiO2_Cells_wf (from OverlayOutlines #16)
Select method for constructing file names Sequential numbers
Enter file prefix TiO2_Cells_WF
Number of digits 3
Saved file format jpeg
Output file location Default Output Folder sub-folder ( d:\cellprofiler_out )
Sub-folder: TiO2_Cells_WF
Overwrite existing files without warning? Yes No
When to save Every cycle
Rescale the images? Yes No
Record the file and path information to the saved image? Yes No
Welcome to CellProfiler
Type here to search
28°C Sunny
17:30
16/07/2022

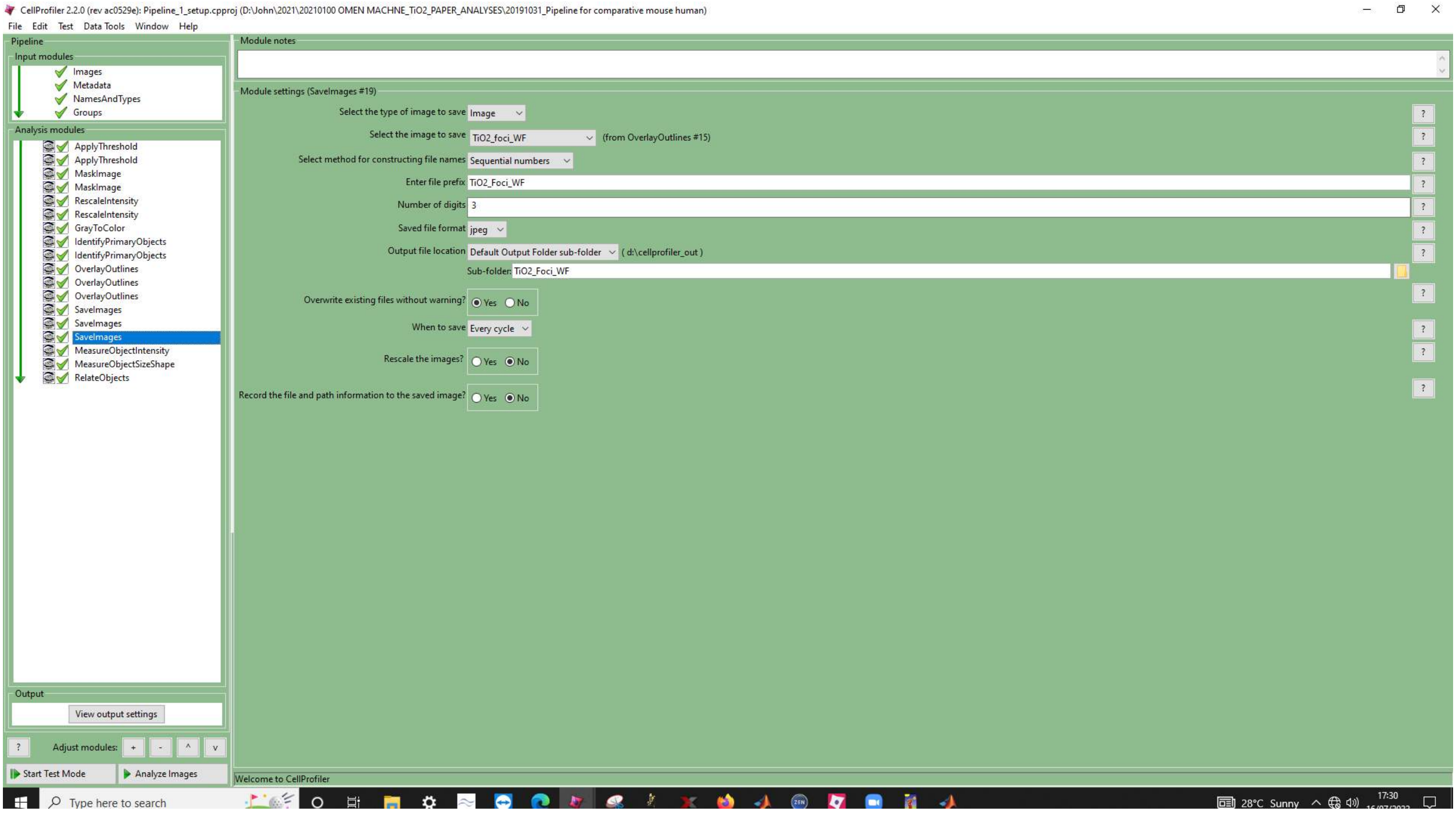
CellProfiler 2.2.0 (rev ac0529e): Pipeline_1_setup.cpproj (D:\John\2021\20210100 OMEN MACHNE_TiO2_PAPER_ANALYSES\20191031_Pipeline for comparative mouse human)
File Edit Test Data Tools Window Help
Pipeline
Input modules
Images
Metadata
NamesAndTypes
Groups
Analysis modules
ApplyThreshold
ApplyThreshold
MaskImage
MaskImage
RescaleIntensity
RescaleIntensity
GrayToColor
IdentifyPrimaryObjects
IdentifyPrimaryObjects
OverlayOutlines
OverlayOutlines
OverlayOutlines
SaveImages
SaveImages
SaveImages
MeasureObjectIntensity
MeasureObjectSizeShape
RelateObjects
Output
View output settings
Adjust modules:
Start Test Mode
Analyze Images
Module notes
Module settings (SaveImages #19)
Select the type of image to save Image
Select the image to save TiO2_foci_WF (from OverlayOutlines #15)
Select method for constructing file names Sequential numbers
Enter file prefix TiO2_Foci_WF
Number of digits 3
Saved file format jpeg
Output file location Default Output Folder sub-folder ( d:\cellprofiler_out )
Sub-folder: TiO2_Foci_WF
Overwrite existing files without warning? Yes No
When to save Every cycle
Rescale the images? Yes No
Record the file and path information to the saved image? Yes No
Welcome to CellProfiler
Type here to search
28°C Sunny
17:30

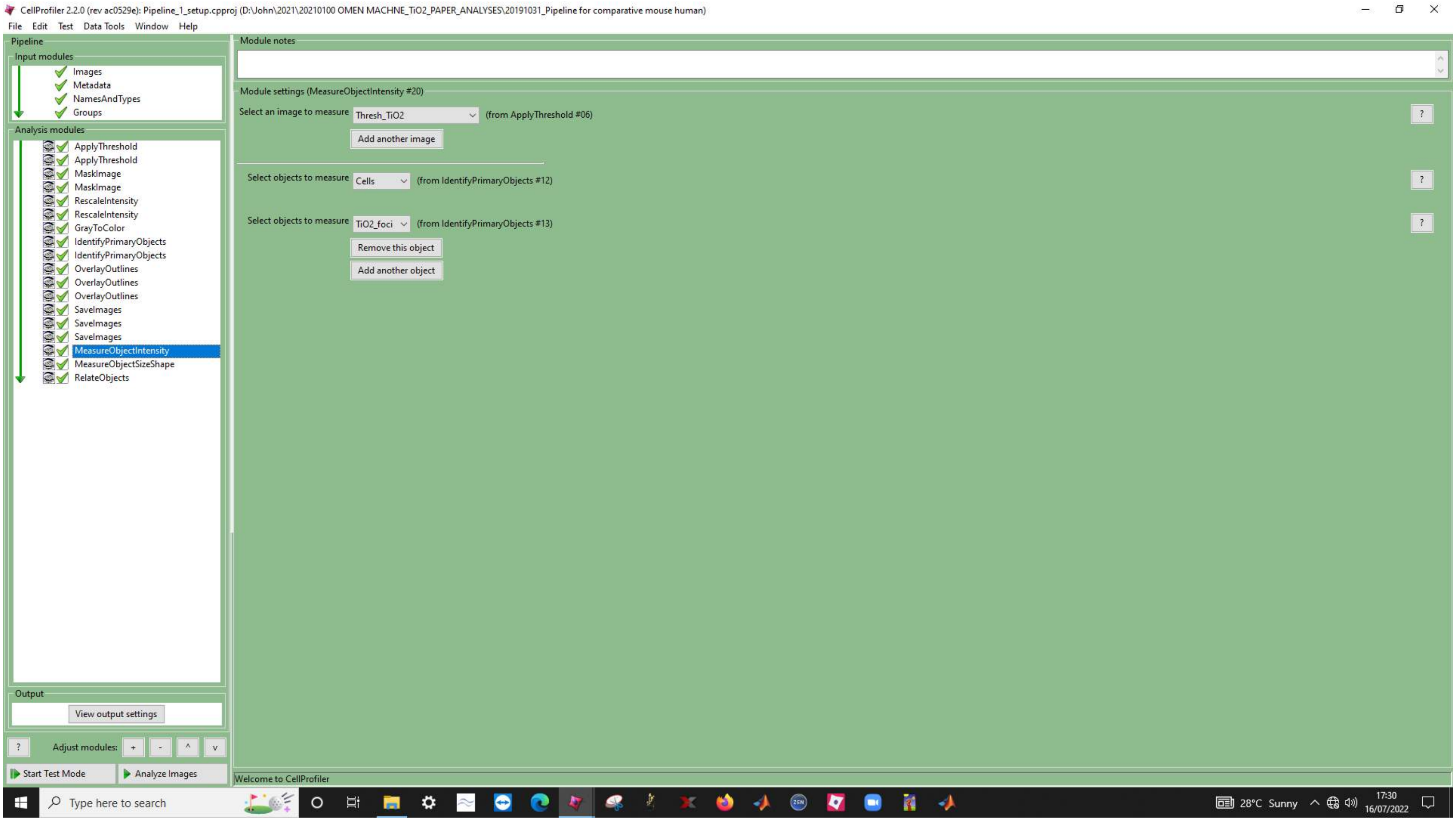
CellProfiler 2.2.0 (rev ac0529e): Pipeline_1_setup.cpproj (D:\John\2021\20210100 OMEN MACHNE_TiO2_PAPER_ANALYSES\20191031_Pipeline for comparative mouse human)
File Edit Test Data Tools Window Help
Pipeline
Input modules
Images
Metadata
NamesAndTypes
Groups
Analysis modules
ApplyThreshold
ApplyThreshold
MaskImage
MaskImage
RescaleIntensity
RescaleIntensity
GrayToColor
IdentifyPrimaryObjects
IdentifyPrimaryObjects
OverlayOutlines
OverlayOutlines
OverlayOutlines
SaveImages
SaveImages
SaveImages
MeasureObjectIntensity
MeasureObjectSizeShape
RelateObjects
Output
View output settings
Adjust modules:
Start Test Mode
Analyze Images
Module notes
Module settings (MeasureObjectIntensity #20)
Select an image to measure
Thresh_TiO2
(from ApplyThreshold #06)
Add another image
Select objects to measure
Cells
(from IdentifyPrimaryObjects #12)
Select objects to measure
TiO2_foci
(from IdentifyPrimaryObjects #13)
Remove this object
Add another object
Welcome to CellProfiler
Type here to search
28°C Sunny
17:30
16/07/2022

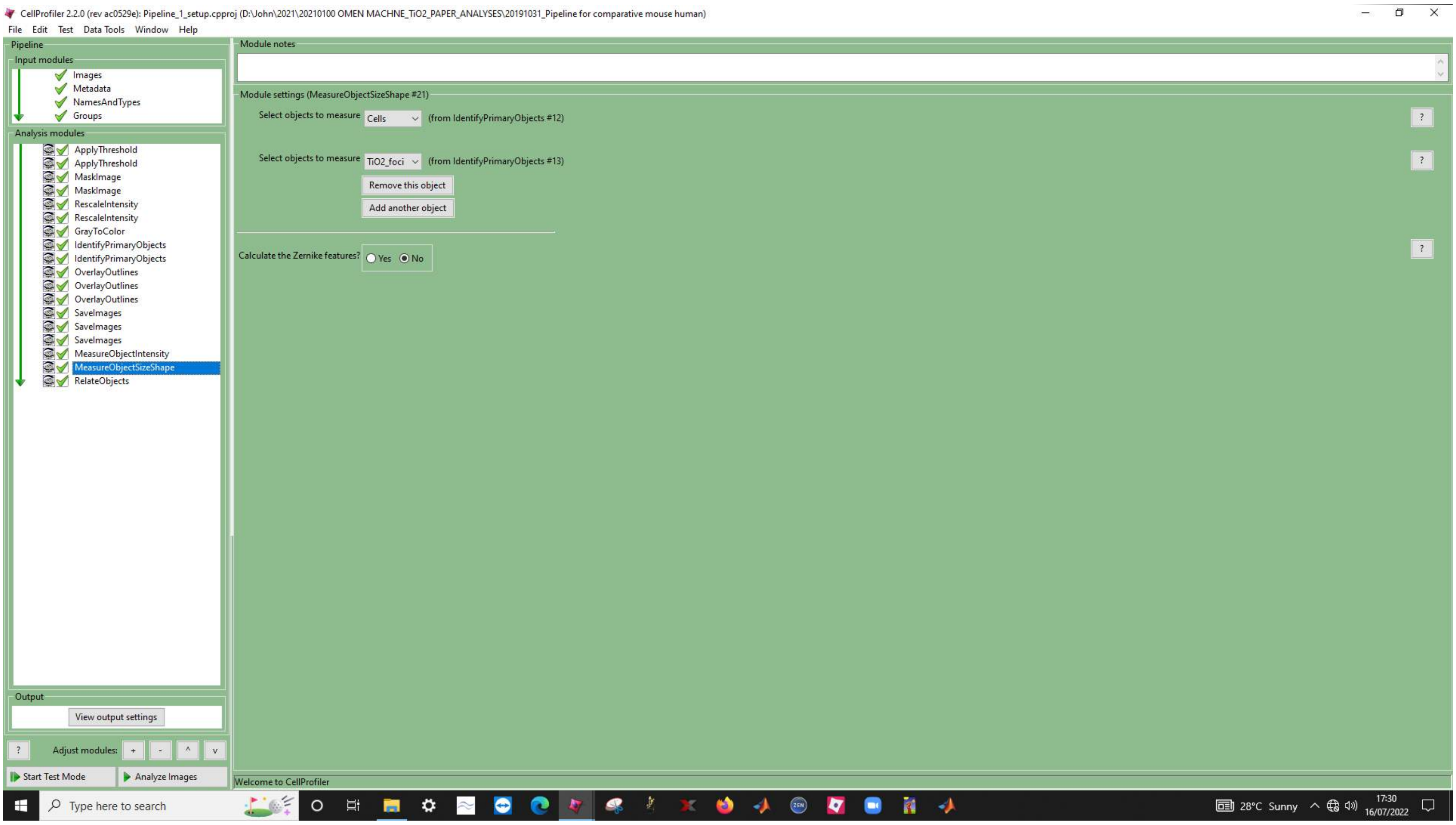
CellProfiler 2.2.0 (rev ac0529e): Pipeline_1_setup.cpproj (D:\John\2021\20210100 OMEN MACHNE_TiO2_PAPER_ANALYSES\20191031_Pipeline for comparative mouse human)
File Edit Test Data Tools Window Help
Pipeline
Input modules
Images
Metadata
NamesAndTypes
Groups
Analysis modules
ApplyThreshold
ApplyThreshold
MaskImage
MaskImage
RescaleIntensity
RescaleIntensity
GrayToColor
IdentifyPrimaryObjects
IdentifyPrimaryObjects
OverlayOutlines
OverlayOutlines
OverlayOutlines
SaveImages
SaveImages
SaveImages
MeasureObjectIntensity
MeasureObjectSizeShape
RelateObjects
Output
View output settings
Adjust modules:
Start Test Mode
Analyze Images
Module notes
Module settings (MeasureObjectSizeShape #21)
Select objects to measure
Cells
(from IdentifyPrimaryObjects #12)
Select objects to measure
TiO2_foci
(from IdentifyPrimaryObjects #13)
Remove this object
Add another object
Calculate the Zernike features?
Yes
No
Welcome to CellProfiler
Type here to search
28°C Sunny
17:30
16/07/2022

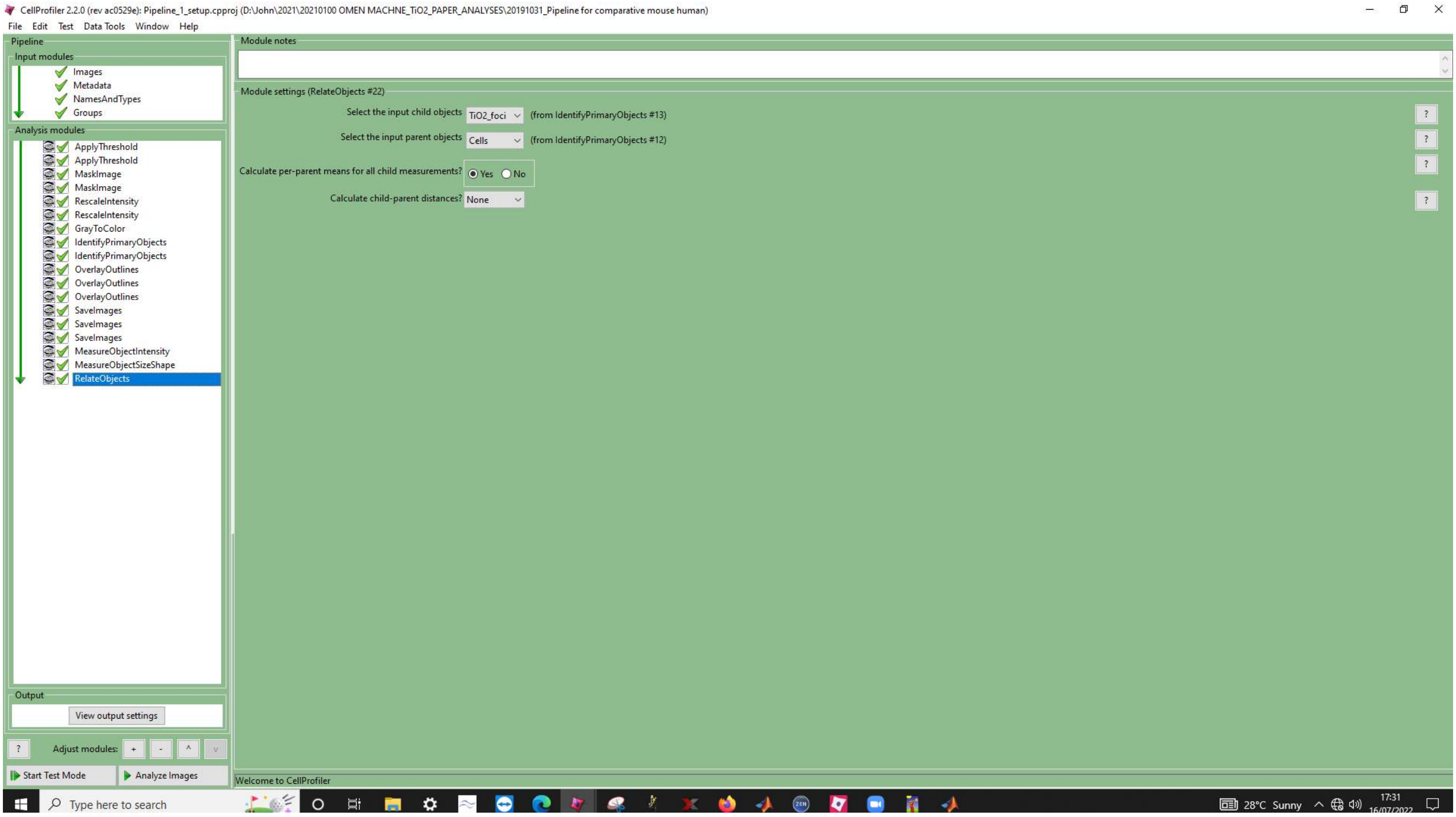
CellProfiler 2.2.0 (rev ac0529e): Pipeline_1_setup.cpproj (D:\John\2021\20210100 OMEN MACHNE_TiO2_PAPER_ANALYSES\20191031_Pipeline for comparative mouse human)
File Edit Test Data Tools Window Help
Pipeline
Input modules
Images
Metadata
NamesAndTypes
Groups
Analysis modules
ApplyThreshold
ApplyThreshold
MaskImage
MaskImage
RescaleIntensity
RescaleIntensity
GrayToColor
IdentifyPrimaryObjects
IdentifyPrimaryObjects
OverlayOutlines
OverlayOutlines
OverlayOutlines
SaveImages
SaveImages
SaveImages
MeasureObjectIntensity
MeasureObjectSizeShape
RelateObjects
Output
View output settings
Adjust modules:
Start Test Mode
Analyze Images
Module notes
Module settings (RelateObjects #22)
Select the input child objects
TiO2_foci
(from IdentifyPrimaryObjects #13)
Select the input parent objects
Cells
(from IdentifyPrimaryObjects #12)
Calculate per-parent means for all child measurements?
Yes
No
Calculate child-parent distances?
None
Welcome to CellProfiler
Type here to search
28°C Sunny
17:31
